## Supplementary material for "Analytical method for reconstructing the stress on a spherical particle from its surface deformation": TeX files: manuscript.tex

\documentclass[aps,pre,twocolumn,10pt,superscriptaddress,nofootinbib,balancelastpage]{revtex4-2} 
\usepackage[latin1]{inputenc}
\usepackage{amsmath,amssymb}
\usepackage{mathrsfs} 
\usepackage[capitalise]{cleveref}
%\usepackage{siunitx}
\usepackage{braket}
\usepackage{calc}
\usepackage{tabularx}
\usepackage{dsfont}
\usepackage{xcolor}
\usepackage{ifthen}
\usepackage[normalem]{ulem}% Fuer Durchstreichungen
\usepackage{graphicx}

\setcounter{secnumdepth}{5}
\setcounter{tocdepth}{5}%
\newcommand{\subsubsubsection}{\paragraph}%
\allowdisplaybreaks

%\newcommand{\crefrangeconjunction}{--}
\newcommand{\crefrangeconjunction}{-}
\newcommand{\creflastconjunction}{, and~}

\newcommand{\Null}{\mathbf{0}}%
\newcommand{\Eins}{\mathds{1}}%
\newcommand{\id}{\operatorname{id}}%
\newcommand{\ii}{\mathrm{i}}%
\newcommand{\R}{\mathbb{R}}%
\newcommand{\C}{\mathbb{C}}%
\newcommand{\rs}{R\hskip1pt'}%
\newcommand{\rss}{R\hskip1pt''}%
\newcommand{\rsss}{R\hskip1pt'''}%
\newcommand{\dif}{\mathrm{d}}%
\newcommand{\tdif}[2]{\frac{\dif#1}{\dif#2}}%
\newcommand{\pdif}[2]{\frac{\partial#1}{\partial#2}}%
\newcommand{\ppdif}[2]{\frac{\partial^{2}#1}{\partial#2^{2}}}%
\newcommand{\Nabla}{\vec{\nabla}}%
\newcommand{\Div}{\operatorname{div}}%
\newcommand{\Laplace}{\boldsymbol{\triangle}}%
\newcommand{\fdif}{\operatorname{\delta}}%
\newcommand{\Fdif}[2]{\frac{\fdif\!#1}{\fdif\!#2}}%
\newcommand{\Rea}{\operatorname{Re}}%
\newcommand{\Ima}{\operatorname{Im}}%
\newcommand{\abs}[1]{\lvert#1\rvert}%
\newcommand{\uu}{\hat{u}}
\newcommand{\nablax}{\Nabla_{\vec{r}}}
\newcommand{\nablaxp}{\Nabla_{\vec{r}'}}
\newcommand{\nablau}{\Nabla_{\uu}}
\newcommand{\nablaup}{\Nabla_{\uu}'}
\newcommand{\norm}[1]{\lVert#1\rVert}%
\newcommand{\rt}{(\vec{r},t)}
\newcommand{\rvs}{(\vec{r},s)}
\newcommand{\rut}{(\vec{r},\uu,t)}
\newcommand{\rput}{(\vec{r}',\uu',t)}
\newcommand{\rpt}{(\vec{r}',t)}
\newcommand{\rus}{(\vec{r},\uu,s)}
\newcommand{\diag}{\operatorname{diag}}%
\newcommand{\tr}{\operatorname{tr}}%
\newcommand{\Tr}{\operatorname{Tr}}%
\newcommand{\ZT}[1]{\textquotedblleft#1\textquotedblright}%

\renewcommand{\vec}[1]{\boldsymbol{#1}}

\newcolumntype{Y}{>{\centering\arraybackslash}X}%     
\newcolumntype{Z}{>{\raggedright\arraybackslash}X}%   

\newlength{\myl}%
\newcommand{\SUM}[2]{{\setlength{\myl}{\widthof{$\displaystyle\sum_{#1}^{#2}$}*\real{0.5}-\widthof{$\displaystyle\sum$}*\real{0.5}}\sum_{#1}^{#2}\;\hspace{-\the\myl}}}% Summen in abgesetzten Gleichungen
\newcommand{\INT}[3]{\settowidth{\myl}{$\displaystyle\int_{#1}^{#2}$}{\int_{#1}^{#2}\;\;\;\hspace{-\the\myl}\dif #3}\,}% Integrale in abgesetzten Gleichungen
\newcommand{\TINT}[3]{\settowidth{\myl}{$\int_{#1}^{#2}$}{\int_{#1}^{#2}\!\ifthenelse{\equal{#1#2}{}}{}{\;\;\;\;\hspace{-\the\myl}}\dif #3}\,}%
\newcommand{\EINT}[3]{\settowidth{\myl}{$\int_{#1}^{#2}$}{\int_{#1}^{#2}\;\;\;\,\hspace{-\the\myl}\dif #3}\,}% Integrale in Exponenten

\newcommand{\Clmn}[3]{c_{#1#2#3}}%
\newcommand{\CIIIDIJa}{c^{(3\mathrm{D})}}%
\newcommand{\CIIIDIJb}[2]{c^{(3\mathrm{D})}_{#1#2}}%
\newcommand{\CIIIDIJc}[4]{c^{(3\mathrm{D})}_{#1#2#3#4}}%
\newcommand{\CIIIDIJd}[6]{c^{(3\mathrm{D})}_{#1#2#3#4#5#6}}%
\newcommand{\WDvarpi}[3]{D^{#1}_{#2#3}}%
\newcommand{\EinrueckabstandI}{\quad\,}% 
\newcommand{\EinrueckabstandII}{\quad\;}% 

\newcommand{\INTOI}{\INT{S_{1}}{}{\Omega}}%
\newcommand{\TINTOI}{\TINT{S_{1}}{}{\Omega}}%
\newcommand{\INTOII}{\INT{S_{2}}{}{\Omega}}%
\newcommand{\TINTOII}{\TINT{S_{2}}{}{\Omega}}%
\newcommand{\ff}[3]{f_{#3}^{(#1\mathrm{D})}}%
\newcommand{\ffp}[3]{f_{#3}^{(#1\mathrm{D})'}}%
\newcommand{\TT}[3]{\mathrm{T}_{#3}^{(#1\mathrm{D})}}%
\newcommand{\fn}[2]{#1(#2)}%

%\newcommand{\New}[1]{\textcolor{green}{#1}}%
\newcommand{\Comment}[1]{\textcolor{magenta}{[#1]}}%
\newcommand{\Alternativformulierung}[1]{\textcolor{violet}{[#1]}}%
\newcommand{\Leacomment}[1]{\textcolor{blue}{[#1]}}%
%\newcommand{\Problem}[1]{\textcolor{red}{#1}}%
%\newcommand{\Remove}[1]{\textcolor{red}{\sout{#1}}}%


\begin{document}
	
\title{Analytical method for reconstructing the stress on a spherical particle from its surface deformation}
\author{Lea Johanna Kr\"uger}
\affiliation{Institute of Theoretical Physics, Center for Soft Nanoscience, University of M\"unster, 48149 M\"unster, Germany}

\author{Michael te Vrugt}
\affiliation{Institute of Theoretical Physics, Center for Soft Nanoscience, University of M\"unster, 48149 M\"unster, Germany}

\author{Stephan Br\"oker}
\affiliation{Institute of Theoretical Physics, Center for Soft Nanoscience, University of M\"unster, 48149 M\"unster, Germany}

\author{Bernhard Wallmeyer}
\affiliation{Centre for Molecular Biology of Inflammation, Institute of Cell Biology, University of M\"unster, 48149 M\"unster, Germany}

\author{Timo Betz}
\affiliation{Third Institute of Physics - Biophysics, University of G\"ottingen, 37077 G\"ottingen, Germany}

\author{Raphael Wittkowski}
\affiliation{Institute of Theoretical Physics, Center for Soft Nanoscience, University of M\"unster, 48149 M\"unster, Germany}


\begin{abstract}
The mechanical forces that cells experience from the tissue surrounding them are crucial for their behavior and development. Experimental studies of such mechanical forces require a method for measuring them. A widely used approach in this context is bead deformation analysis, where spherical particles are embedded into the tissue. The deformation of the particles then allows to reconstruct the mechanical stress acting on them. Existing approaches for this reconstruction are either very time-consuming or not sufficiently general. In this article, we present an analytical approach to this problem based on an expansion in solid spherical harmonics that allows us to find the complete stress tensor describing the stress acting on the tissue. Our approach is based on the linear theory of elasticity and uses an ansatz derived by Love. We clarify the conditions under which this ansatz can be used, making our results useful also for other contexts in which this ansatz is employed. Our method can be applied to arbitrary radial particle deformations and requires a very low computational effort. The usefulness of the method is demonstrated by an application to experimental data. 
\end{abstract}
\maketitle

\section*{Statement of Significance}
Measurements of mechanical forces acting on cells in a tissue are important for understanding the physical behavior of biological systems, but they are also quite challenging. A common strategy is to place a spherical bead inside the tissue and to then reconstruct the mechanical stress from the bead deformation that this stress causes. Here, we introduce a novel analytical method using which this reconstruction can be achieved. This method is significantly faster than numerical approaches and significantly more general than existing analytical techniques, such that it can be expected to find a broad range of applications in mechanobiology.

\section{\label{introduction}Introduction}
\noindent 
A proper description of cell development is essential for many areas of research, including tumor research \cite{tse,Styli,Fiore}, embryogenesis \cite{matre}, and the study of tissue growth \cite{Guillot,Nagle}. The development of cells is influenced by the extracellular matrix (ECM), which is a three-dimensional network of macromolecules surrounding the cells. Biochemical and mechanical properties \cite{trappmann,jean} of the ECM alter the shape and activity of the cells \cite{handorf,Frantz,Discher,Sporrer,Goldmann}, while the macromolecules making up the ECM are produced by the cells themselves. This leads to a feedback loop between the ECM and the cells that usually results in homeostasis. Due to the importance of the ECM, a throughout analysis of its properties is essential for understanding cell development, where homeostasis is disturbed.\\
There has been a significant amount of work on the biochemical properties of the ECM in the past (see Refs.\ \cite{Hay,Valerie,Frantz} for reviews). More recently, experimental advances made it possible to study also the mechanical properties \cite{Dolega,Lee,trappmann} and its influences \cite{Hofmeier, Nagle, Meglio, Mohaghegian2,Biswas,MALMIKAKKADA} on cell development. Measuring the forces inside the ECM is very difficult for a variety of reasons. First, the order of magnitude of the forces inside the tissue is usually in the range of pN to nN \cite{Wallmeyer}. Therefore, measurements have to be sensitive to very small stress differences. Second, the measurements should be performed in vivo, without disturbing the complex mechanisms inside the tissue. An innovative method developed by \citet{Campas} achieves this by exploiting the fact that stress acting on a soft particle results in deformation. Oil droplets are inserted into the tissue, where their deformed shape is measured. Since the mechanical properties of the oil droplets are known, the method allows to reconstruct the stress acting inside the tissue. A remaining challenge is the incompressibility of oil droplets preventing a direct measurement of compressive stresses. The idea was developed further by other researchers \cite{Dolega,Mohagheghian,Traber,Wallmeyer,Lee} who designed new beads whose mechanical properties are optimized for measuring the stress inside the tissue. These beads typically consist of some type of hydrogels, such as polyacrylamide (PAA). One of the main differences between these beads and oil droplets is that the beads are compressible. The general procedure is the same for all approaches where PAA beads are used \cite{Dolega,Mohagheghian,Traber,Wallmeyer,Lee}. One places the PAA beads inside the tissue, where they are deformed due to the stresses acting on them. Then, the deformed shape is measured, usually via confocal microscopy \cite{Wallmeyer}. The last step is the reconstruction of the stress from the shape of the deformed bead. This last step will be the main focus of this article.\\
Using the theory of elasticity, the problem of reconstructing the stress from the shape deformation can be formulated as a differential equation that can be solved with the displacement of the surface as a boundary condition. Numerical approaches to this problem are computationally very expensive, an analytical solution method applicable to arbitrarily deformed spheres, does not exist. In this article, we present an improved analytical solution for the differential equation that allows to compute the stress. To achieve this, an ansatz derived by \citet{Love} (appearing also in Refs.\ \cite{Trefftz,Lurie}) is used. The way this ansatz is constructed leads to several subtleties concerning the harmonicity of the individual terms appearing in the ansatz. These subtleties have not been addressed in existing analytical approaches. In this article, we develop a method that allows using the ansatz for arbitrary radial bead deformations. We demonstrate the applicability of this method by an analysis of experimental data.\\
This article is structured as follows. 
\begin{figure*}[tbh]
\includegraphics[width=\textwidth]{fig1.pdf}
\caption{\label{fig1}Schematic visualization of the method of bead deformation analysis. Step 1: The PAA beads are inserted into the tissue. Step 2: The beads are deformed due to stresses acting in the tissue. Step 3: Using confocal microscopy, the shape of the deformed beads is obtained. Step 4: The stress tensor $\hat{T}$ is reconstructed from the deformed shape.}
\end{figure*}
In section \ref{Theory}, a short introduction to the linear theory of elasticity, including the differential equation central to this article, is provided. Section \ref{current} contains a summary of the current state of research on the stress reconstruction problem. In section \ref{Ansatz}, the ansatz by Love is presented. Additionally, we explain our general approach for solving the differential equation using the ansatz. In section \ref{derivation}, the derivation of the stress tensor using the ansatz is shown. This result is, in section \ref{data}, used to analyze example experimental data. We conclude in section \ref{conc}.
\section{\label{Theory}Theoretical Background}
\noindent The reconstruction of the stress acting on a bead is possible using the theory of elasticity. In this section, the problem is formulated in the language of this theory by deriving a differential equation for the displacement vector $\vec{u}$. Apart from the displacement vector itself, the most important quantities are the stress tensor $\hat{T}$ and the strain tensor $\hat{\epsilon}$. For investigating these quantities, we use the linear theory of elasticity, which assumes that the change of displacement $u_i$ in any direction is small compared to the size of the considered objects:
\begin{align}\label{the:cond:lte}
\Biggl|\frac{\partial u_i}{\partial x_j}\Biggr| \ll 1.
\end{align}
To derive the differential equation for the displacement vector $\vec{u}$, only a handful of fundamental relations of linear elasticity theory are needed \cite{Lurie,Lifschitz,Red}. One is Hooke's law \cite{Lifschitz}
\begin{equation}\label{hooke}
    \hat{T} = 2G\frac{\nu}{1-2\nu} \tr(\hat{\epsilon}) + 2G\hat{\epsilon},
\end{equation}
which linearly connects the stress tensor $\hat{T}$ and the strain tensor $\hat{\epsilon}$. Here, $G$ denotes the shear modulus and $\nu$ Poisson's, both of which characterize the material properties of the deformed body. $\tr((\hat{\epsilon}))$ describes the trace of the strain $\hat{\epsilon}$.
The form \eqref{hooke} of Hooke's law assumes elastic, isotropic, and thermodynamically reversible deformations. Now, the strain tensor $\hat{\epsilon}$ is replaced by its definition in terms of the displacement vector $\vec{u}$. This definition is, in the linear theory of elasticity, given by
\begin{equation}\label{epsilon}
    \hat{\epsilon} = \frac{1}{2}\bigl(\vec{\nabla u}+(\vec{\nabla u})^\text{T}\bigr) = \text{Def}(\vec{u}),
\end{equation}
where Def$(\vec{u})$ is the deformation of the displacement vector field. For the next step, we use the fact that the shape of the beads inside the tissue remains stable, which suggests that static equilibrium is reached. In equilibrium, the divergence of the stress tensor $\vec{\nabla}\cdot\hat{T}$ equals the external forces per unit volume $\rho \vec{K}$, where $\rho$ is the mass density of the elastic body and $\vec{K}$ the force acting on a unit mass. We assume that the external forces are negligible compared to the stress arising from the ECM. Consequently, the divergence of \cref{hooke} is equal to zero. This gives the differential equation
\begin{equation}\label{de}
    \frac{1}{1-2\nu} \vec{\nabla}(\vec{\nabla \cdot u}) + \vec{\nabla} ^2 \vec{u} = 0,
\end{equation}
from which the displacement vector field $\vec{u}$ can be calculated.\\
In experiments, one obtains the shape of the surface of the deformed bead. If the displacement vector on the surface of the bead can be obtained from this information, it can be used as a boundary condition for solving \cref{de}, which in turn allows to obtain the entire displacement field. Inserting the result into \cref{epsilon,hooke} then gives the complete stress tensor $\hat{T}$ and strain tensor $\hat{\epsilon}$.

\section{\label{current}Current state of research}
\noindent There exist a variety of methods for reconstructing the stress from the shape of the deformed bead. These face three main challenges, namely the \textit{computational effort} needed to find the solution of \cref{de}, the \textit{generality of shapes} the method can be applied to, and the \textit{connection of displacement and shape}.\\
The issue of computational effort arises primarily for numerical approaches. Solving the differential equation \eqref{de} or a similar one using the finite element method (FEM) allows to find a solution for arbitrary shapes of the deformed bead \cite{Mohagheghian,Traber}. The downside of the FEM and of numerical approaches in general is the computational effort. Currently, this method takes several days per bead to obtain the result \cite{Vorselen}. Taking into account that understanding the stress inside the ECM requires analyzing a large number of beads, a quicker method to find the stress is required. \\
Alternatively, analytical approaches can be used, which are usually a lot quicker than the FEM. The constraint of many existing analytical approaches is that they make highly simplifying assumptions regarding the shape of the deformed bead. As a result, these methods do not apply to general deformations, but only to those occurring in a specific kind of experimental setup. Examples are the method by \citet{Dolega}, which only applies to hydrostatically compressed beads, and the one by \citet{Lee}, which exclusively describes deformations along one axis.\\
The last issue concerning the connection of displacement $\vec{u}$ and shape arises when the boundary condition for \cref{de} is formulated. The initial state is considered to be a sphere of initial radius $r_0$. Equation \eqref{de} is a differential equation for the displacement $\vec{u}$. The boundary condition should therefore contain information about $\vec{u}$, while the only quantity known from measurement is the shape of the surface of the deformed bead. The connection between the shape of the surface and the displacement of initial the surface $\vec{u}|_{r_0}$ is, however, not unique. In fact, there are infinitely many possibilities for the displacement field on the boundary $\vec{u}|_{r_0}$ that result in the same shape. Often, this problem is solved by assuming that the displacement of the surface is exclusively directed into the radial direction \cite{Dolega, Mohagheghian, Wallmeyer}. Another, more general, possibility was presented by \citet{Vorselen}, who introduced a cost function that minimizes the elastic energy to find the optimal displacement numerically.\\

\section{\label{Ansatz}Ansatz and outline of the approach}
\noindent Here, we introduce an analytical approach where the computational effort is low even though it applies to complicated radial deformations. This is achieved using an ansatz which solves the differential equation \eqref{de} and allows a decomposition of the deformed shape into solid spherical harmonic functions. The main ideas follow Refs. \cite{Wallmeyer,Hofmeier}, where the same ansatz is applied. The ansatz will be analyzed further so that its implications can be understood in more depth, which allows us to use it for a generalized shape of the deformed bead. The connection of shape and displacement $\vec{u}|_{r_0}$ will be assumed to be radial for now.\\
A typical procedure for solving a differential equation of the form \eqref{de} is to use an ansatz that consists exclusively of harmonic functions \cite{Trefftz, Love, Neuber, Papko}. A harmonic is a function $f$ that solves Laplace's equation
\begin{equation}
    \vec{\nabla}^2 f= 0.
\end{equation}
Harmonic functions are useful because their behavior is well understood. In particular, it is known that harmonic functions can be expanded into solid spherical harmonics (SSHs), which are three-dimensional harmonic functions. There are two distinct types of SSHs, regular ($R_n^m$) and irregular ($I_n^m$) ones. They are defined as
\begin{align}\label{eq:SSH}
R_n^m(r, \theta,\phi) &=
 r^n Y_{nm}(\theta,\phi),  \\
I_n^m(r, \theta,\phi) &=\frac{1}{r^{n+1}}Y_{nm}(\theta,\phi),
\end{align} 
where $Y_{nm}$ are spherical harmonic functions. (See Appendix \ref{Ynm} for the definition used in this article.) The difference between regular and irregular SSHs is that regular SSHs diverge for large $r$, whereas irregular SSHs diverge at the origin. \\ The differential equation \eqref{de} can be solved using a general expansion of an ansatz for $\vec{u}$, consisting of harmonic functions, into SSHs. The coefficients of the expansion have to be deduced from the boundary conditions of the problem under consideration. A procedure of this form is useful for the bead deformation analysis because the shape of the deformed bead is usually expanded into spherical harmonics as well. The maximal order of this expansion determines the precision of the measurement.\\
One ansatz for the displacement $\vec{u}$ that solves the differential equation \eqref{de} and consists of harmonic functions only was suggested by \citet{Love} and \citet{Trefftz}. The ansatz is particularly useful for studying the deformation of spherical bodies and is given by
\begin{equation}\label{ansatz}
    \vec{u}(r,\theta,\phi) = \vec{U}(r,\theta,\phi) +(r^2-r_0^2)\vec{\nabla}\psi(r,\theta,\phi),
\end{equation}
where $\psi$ and all components of the vector function $\vec{U}$ are harmonic functions. Moreover, $r_0$ is the radius of the initial spherical body and $r$ is the radial coordinate, where the origin lies in the center of the initial sphere. Equation \eqref{ansatz} shows that the function $\vec{u}$ coincides with $\vec{U}$ on the surface of the initial sphere (i.e., for $r=r_0$). From the ansatz \eqref{ansatz}, a connection of the scalar function $\psi$ and the vector function $\vec{U}$ can be deduced. This relation is based on an expansion of both functions $\psi$ and $\vec{U}$ into SSHs and connects $\Psi_{n-1}$ and $\vec{U}_n$, where $n$ is the order of the corresponding SSH:
\begin{equation}\label{relation}
      \psi_{n-1} = \frac{\vec{\nabla \cdot U}_n}{-2((2-3n)-2\nu(1-2n))}.
\end{equation}
The derivation of \cref{ansatz,relation} is shown in Appendix \ref{Derivationansatz}. This derivation shows that \cref{relation} is only valid if $\vec{U}$ is expanded to  $\vec{U}_n$ in the Cartesian basis. If all Cartesian components of $\vec{U}_n$ are SSHs of order $n$, the same cannot be true for the spherical components of $\vec{U}_n$. In fact, \cref{relation} can only work in the way stated above if $\vec{U}_n$ is a SSH of order $n$ in all of its Cartesian components.\\
Having understood the conditions for the applicability of the ansatz \eqref{ansatz}, we now expand the complete displacement function $\vec{u}$ as given in \cref{ansatz} into SSHs to obtain the general solution of the differential equation \eqref{de}, where the coefficients of the expansion still need to be evaluated. For the expansion of the scalar function $\psi$ and vector function $\vec{U}$, regular SSHs are chosen inside the initial sphere and irregular SSHs outside it. Therefore, the deformation vector $\vec{u}$ does not diverge for large $r$ and is not singular at the origin.
Additionally, the expansion is ascertained to be continuous by the use of appropriate prefactors \cite{Lurie,Wallmeyer}:
\begin{equation}\label{eq:Psi}
\psi= \begin{cases}
\sum\limits_{n=0}^{\infty}\sum\limits_{m=-n}^{n} \frac{r^n}{r_0^n}a_{nm} Y_{nm} &\text{for $r<r_0$,}\\
\sum\limits_{n=0}^{\infty}\sum\limits_{m=-n}^{n} \frac{r_0^{n+1}}{r^{n+1}}a_{nm} Y_{nm} &\text{for $r>r_0$}.
\end{cases}
\end{equation}
\begin{equation}\label{eq:U}
\vec{U}= \begin{cases}
\sum\limits_{n=0}^{\infty}\sum\limits_{m=-n}^{n} \frac{r^n}{r_0^n}\vec{K}_{nm} Y_{nm}  &\text{for $r<r_0$,}\\
\sum\limits_{n=0}^{\infty}\sum\limits_{m=-n}^{n} \frac{r_0^{n+1}}{r^{n+1}}\vec{K}_{nm} Y_{nm} &\text{for $r>r_0$,}
\end{cases}
\end{equation}
What is missing for the full solution for the differential equation \eqref{de} are the scalar coefficients $a_{nm}$ and the vectorial coefficients $\vec{K}_{nm}$. If these coefficients are known, the complete displacement field $\vec{u}(r,\theta,\phi)$ is determined. Inserting the ansatz function \eqref{ansatz} into the definition of the stress tensor \eqref{epsilon} and the result into Hooke's law gives the complete stress tensor $\hat{T}$ in terms of $\psi$ and $\vec{U}$:\\
\begin{multline}\label{stress}
  \hat{T} = 2G\Big(\frac{\nu}{1-2\nu}\left(\vec{\nabla\cdot U} +2\vec{r\cdot \nabla}\psi\right)\vec{\hat{E}}
  +\text{Def}(\vec{U}) \cr + \vec{r}\otimes \vec{\nabla}\psi +(\vec{r}\otimes \vec{\nabla}\psi)^\text{T}
  +(r^2-r_0^2)\vec{\nabla} \otimes \vec{\nabla}\psi\Big).
\end{multline}
Here, $\vec{r}$ is the positional vector, $\vec{\hat{E}}$ is the three dimensional unit tensor and $\otimes$ is the outer product ($\vec{u}\otimes\vec{v})_{ij} = u_iv_j$).
\section{\label{derivation}Radial Deformation and the boundary condition}
To obtain the coefficients $a_{nm}$ and $\vec{K}_{nm}$ that fully determine the stress tensor $\hat{T}$, we require the boundary condition for the displacement vector $\vec{u}|_{r_0}$. The experimental data only provides the shape of the deformed beads. To construct from this information a boundary condition for the displacement vector, we assume that the displacement is only radial. If the radius of the deformed bead at angles $\theta$ and $\phi$ is given by $s(\theta, \phi)$, this deformed shape can be expanded into spherical harmonics in the form
\begin{equation}
     s(\theta,\phi) = \sum\limits_{n=0}^{N}\sum\limits_{m=-n}^{n}s_{nm}Y_{nm},
\end{equation}
where the expansion coefficients $s_{nm}$ are given by
\begin{equation}
    s_{nm} = \int_\Omega \! s(\theta,\phi)Y_{nm}^*(\theta,\phi)\text{d}\Omega.
\end{equation}
We then assume the surface deformation to be
\begin{align}\label{dis}
     \vec{u}(r_0,\theta,\phi) &=(r_0-s(\theta,\phi))\vec{\hat{e}}_r \nonumber \\ &= \sum\limits_{n=0}^{N}\sum\limits_{m=-n}^{n}d_{n,m}Y_{nm} \vec{\hat{e}}_r,
\end{align}
where $r_0$ is the radius of the undeformed bead, $\vec{\hat{e}}_r$ is the unit vector in radial direction and
\begin{equation}\label{eq:dnm}
    d_{n,m} =\begin{cases}
\sqrt{4\pi}r_0-s_{00} & \text{for } n=m=0,\\
-s_{nm} &\text{else}
\end{cases}
\end{equation}
are the coefficients resulting from the radial difference of the initial sphere and the shape of the deformed bead $s(\theta, \phi)$. At this point, the importance of the prerequisites of the ansatz come into play. Because of the way the ansatz \eqref{ansatz} is constructed, the displacement of the surface completely defines the coefficients of the harmonic vector function $\vec{U}$. To use the relation \eqref{relation} for determining the coefficients $a_{nm}$ that define the function $\psi$, we require the harmonic expansion of $\vec{U}$ in terms of Cartesian coordinates.\\
In Cartesian coordinates, the radial unit vector $\vec{\hat{e}}_r$ consists of spherical harmonic functions of order 1:
\begin{equation}
    \vec{\hat{e}}_r =\begin{pmatrix}Y_{1-1}-Y_{11} \\ \ii\left(Y_{1-1}+Y_{11}\right)\\ \sqrt{2}Y_{10}. \end{pmatrix}.
\end{equation}
To obtain the required expansion of $\vec{u}|_{r_0}$, we need to express the product of spherical harmonics $Y_{nm}$ occurring in the Cartesian components in \cref{dis} in harmonic functions. A useful relation for this purpose is \cite{varsh}
\begin{align}\label{var}
     Y_{l_1}^{m_1}Y_{l_2}^{m_2} = &\sqrt{4\pi}\sum\limits_{L = |l_1-l_2|}^{|l_1+l_2|}C_{l_1,m_1,l_2,m_2}^{L,M}C_{l_1,0,l_2,0}^{L,0} \nonumber \\ &\sqrt{\frac{(2l_1+1)(2l_2+1)}{2L+1}}Y_{L}^{M},
\end{align}
where $C_{l_1,m_1,l_2,m_2}$ are Clebsch-Gordan-coefficients and $M=m_1+m_2 $. Using \cref{var}, the boundary condition $\vec{u}|_{r_0}$ is expressed in such a way that the $n$th term of the expansion contains the spherical harmonic of degree $n$ in every Cartesian component. One can then directly express the coefficients of this expansion with the vectorial coefficients $\vec{K}_{nm}$ occurring in \cref{eq:U} in terms of the known coefficients $d_{n,m}$ and the appropriate Clebsch-Gordan-coefficients (Appendix \ref{Knm}). Without transforming the boundary condition \eqref{dis} into Cartesian coordinates, one cannot make proper use of the ansatz \eqref{ansatz}.\\
Having determined the coefficients $\vec{K}_{nm}$, we can construct the harmonic vector function $\vec{U}$ using \cref{eq:U}. To use \cref{relation} and to obtain the coefficients $a_{nm}$ which describe the harmonic scalar function $\psi$, one needs to evaluate the divergence of $\vec{U}$. A general expression for the divergence of $\vec{U}$ can only be obtained if the individual derivatives of the components $\vec{U}$ are expressed as an expansion in SSHs. The derivatives of spherical harmonic functions with respect to Cartesian coordinates are given in Appendix \ref{derivatives}. These allow to express the divergence of $\vec{U}$ in terms of the coefficients $\Vec{K}_{nm}$, which is done in \cref{eq:co:psi2} in Appendix \ref{derivatives}. Applying the method of equating the coefficients to \cref{relation,eq:co:psi2} yields
\begin{equation}\label{anm}
    a_{nm} = \frac{c_{nm}}{2\bigl(2\nu\left(2n-1\right)-3n\bigr)},
\end{equation}
where we introduced the coefficients $c_{nm}$ to keep the expression \eqref{anm} short. The derivation of \cref{anm} and the definition of the coefficients $c_{nm}$ in terms of $\vec{K}_{nm}$ is given in Appendix \ref{derivatives}.\\
Thereby, we have determined the functions $\psi$ and $\vec{U}$ as given in \cref{eq:U,eq:Psi} from the boundary condition \eqref{dis}. We now insert the results for $\psi$ and $\vec{U}$ into \cref{stress} to obtain the full stress tensor:
\begin{widetext}
    \begin{align}\label{fullstress}
     \hat{T}(r,\theta,\phi) = &\sum\limits_{n=0}^{N}\sum\limits_{m=-n}^{n}\frac{2G}{r_0}\Bigl\{ \frac{\nu}{1-2\nu} a_{nm}\Bigl(2r_0\bigl(1-3n+2\nu(2n+1)\bigr)r^n Y_{nm}+2\vec{r}\cdot\vec{\nabla}(r^n Y_{nm})\Bigr)\nonumber \\  &+\Bigl((\vec{K}_{nm}\cdot\vec{e}_r)\vec{e}_r+(\vec{K}_{nm}\cdot\vec{e}_\theta)\vec{e}_\theta+(\vec{K}_{nm}\cdot\vec{e}_\phi)\vec{e}_\phi\Bigr)\otimes \vec{\nabla}(r^nY_{nm})\nonumber \\ &+\vec{\nabla}(r^nY_{nm})\otimes\Bigl((\vec{K}_{nm}\cdot\vec{e}_r)\vec{e}_r+(\vec{K}_{nm}\cdot\vec{e}_\theta)\vec{e}_\theta+(\vec{K}_{nm}\cdot\vec{e}_\phi)\vec{e}_\phi\Bigr)\nonumber \\ &+(r^2+r_0^2)a_{nm}\vec{\nabla}\otimes\vec{\nabla}(r^nY_{nm})\Bigr\},
\end{align}
\end{widetext}
where $\vec{e}_\theta$ is the azimuthal- and $\vec{e}_\phi$ the polar unit vector.
Even though the expression \eqref{fullstress} is rather long, the evaluation for an arbitrarily chosen bead is simple once the tensor has been evaluated up to the desired order. The overall form of the stress tensor remains the same for every bead and depends on the coefficients $a_{nm}$ and $\vec{K}_{nm}$ explicitly. Therefore, the effort of analyzing multiple beads is greatly reduced compared to numerical methods. $a_{nm}$ is obtained by inserting the coefficients $d_{n,m}$ \cref{eq:dnm} which follow from the deformed shape into \cref{eq:cnm} and the resulting coefficients $c_{nm}$ into \cref{relation}. The coefficients $\vec{K}_{nm}$ follow from inserting $d_{n,m}$ into \cref{eq:rad:8,eq:rad:9,eq:rad:10}.Meanwhile, the result allows -- in contrast to previously developed analytical approaches -- to examine beads of arbitrary radial deformation.\\ 
We now analyze the properties of the resulting stress tensor \eqref{stress}. In order to better understand the result, we consider some basic limiting cases. If the deformed bead remains completely spherical, the resulting stress tensor is proportional to the unit tensor $\hat{E}$. The magnitude of the stress is then fully determined by Poisson's ratio $\nu$, the shear modulus $G$, and the difference between the initial radius $r_0$ and the radius of the deformed sphere $r_1$:
\begin{equation}\label{Test:stress:uni}
    \hat{T}_{\text{sphere}} = 8 G \pi\frac{(r_1-r_0)(1+\nu)}{r_0(1-2\nu)} \hat{E}.
\end{equation}
A uniformly deformed bead therefore corresponds to an isotropic stress tensor.\\
Another case that is relevant for many experimental setups is an uniaxial deformation, where the bead is deformed only in one particular direction. Here, we consider as an example a shape function of the deformed bead $s_{\text{uniaxial}}(\phi,\theta)$ that is given as a sum of the initial sphere (described by the zeroth-order spherical harmonic $Y_{00}$, which is a constant) and an additional term containing the spherical harmonic function of order 2 with $m=0$:
\begin{align}\label{soa}
    s_{\text{uniaxial}} &= \sqrt{4\pi}r_0Y_{00} + d_{2,0}Y_{20}  \nonumber \\
    &= s_{\text{initial}}+ d_{2,0}Y_{20}.
\end{align}
A visualization of the initial sphere $s_{\text{initial}} = \sqrt{4\pi}r_0Y_{00}$ and the deformed sphere is given in \cref{fig2}. Using \cref{soa} to determine the coefficients $a_{nm}$ and $K_{nm}$ by \cref{eq:rad:8,eq:rad:9,eq:rad:10,anm} and inserting these into \cref{fullstress}, we obtain a stress tensor which consists of a radial term and an azimuthal term. 
The radial stress is proportional to the radial displacement of the initial sphere compared to the deformed sphere, similar to the case of spherical deformation. The azimuthal stress, on the other hand, depends on the change of the difference of radii along the azimuthal direction. Both components of the stress tensor are plotted on the surface of the initial sphere in \cref{fig2}. \\
For displacements that are not uniaxial, an additional polar term appears which represents the stress acting on the sphere in polar direction. Similar to the azimuthal stress, the polar stress depends on the change of the difference of radii along the azimuthal direction.
\begin{figure*}[tbh]
\includegraphics[width=\textwidth]{fig2.pdf}
\caption{\label{fig2} The case of uniaxial deformation. \textbf{(a)} Shape of the initial sphere of unit radius compared to the deformed shape of the bead as given in \cref{soa} with $d_{2,0} = 0.2$. \textbf{(b)} Radial stress and \textbf{(c)} azimuthal on the surface of the initial sphere, with the arrows indicating the direction and magnitude of the stress.}
\end{figure*}
\section{\label{data}Analyzing experimental data}
\noindent 
So far, we have discussed very simple shapes where only a small number of spherical harmonics is required to describe the shape of the deformed bead. The stress tensor \eqref{stress} derived here is, however, applicable to arbitrary radial bead deformations. To test the result \eqref{stress} for more complicated cases, experimental data is obtained and analyzed. %data from experiments by the group of Timo Betz is obtained and analyzed. 

In the experiments, beads consisting of PAA were marked by fluorescent pigments and then embedded into the tissue, where the shape of their surface was measured using a light-sheet microscope. A detailed description of the experimental setup can be found in section 3.2 of Ref.\ \cite{Wallmeyer}. For this experimental setup, the original size of the beads varies and cannot be determined after the beads are inserted into the tissue. We assume the beads to be incompressible and perfectly spherical. These assumptions are required here to reconstruct the initial shape from the available data. In general, the analytical method developed here is also applicable to compressible beads.\\
The first step of analyzing the data is to numerically fit a spherical harmonic expansion to the data points for the bead's surface. To determine a sufficiently high upper order for fitting while avoiding overfitting, the data points are compared to the surface generated by the fit. The order of the fit is increased until the order $2n_{\mathrm{max}}$, where the fit exactly coincides with the data points. The more data points there are, the higher the order of the fit needs to be to achieve this. To avoid overfitting, we truncate the expansion at order $n_{\mathrm{max}}$. The number of data points differs for the individual beads, implying that $n_{\mathrm{max}}$ differs as well. In \cref{fig3}(a),(d), we show data points and fits for two of the analyzed beads. The maximal order of fitting is $n_{\mathrm{max}}=13$ for the first bead (\cref{fig3}(a)) and $n_{\mathrm{max}}=26$ for the second bead (\cref{fig3}(d)).\\
As a next step, the original size of the beads is determined by evaluating the volume enclosed by the fitted surface. The original bead is now described by a sphere of the same volume. Thereby, a mathematical description of the shape of the original and the deformed particle is known. Using \cref{dis}, we evaluate the displacement $\vec{u}(r,\theta,\phi)|_{r_0}$ of the surface. The coefficients $a_{nm}$ and $\vec{K}_{nm}$ are determined using the procedure explained in section \ref{derivation}. Then, the resulting coefficients are inserted into the expansion of the stress tensor given by \cref{stress}. As a result, we obtain the complete stress tensor for the beads.\\
To visualize some example results, the radial stress $\hat{T}_{rr}$ on the surface of the original beads is plotted for the two beads in \cref{fig3}(c),(f). Additionally, the radial displacement of the surface \eqref{dis} is plotted in comparison to the radial stress (\cref{fig3}(b),(e)).  
The stress is pointing inwards when the bead is compressed and outwards when it is stretched. Therefore, the method introduced in this work allows to reconstruct the stress exerted on radially deformed spherical particles even if the shape of the deformed particle is complicated, as it typically is in experimental contexts.
\begin{figure*}[tbh]
\includegraphics[width=\textwidth]{fig3.pdf}
\caption{\label{fig3} Analysis of experimental data for the deformation of two PAA beads.  \textbf{(a)} Data points and fitted reconstructed surface of bead 1. The order of fitting is $n_{\mathrm{max}} =13$. \textbf{(b)} Initial and deformed surfaces of bead 1. The arrows represent the displacement $\vec{u}$. \textbf{(c)} Radial stress on the surface of the undeformed bead 1. \textbf{(d)-(f)} Like (a)-(c), but for bead 2. The order of fitting is $n_{\mathrm{max}} = 26$ now.}
\end{figure*}
\section{\label{conc}Conclusions}
In this article, a general analytical approach that allows to reconstruct the complete stress tensor for any radially deformed spherical bead from the deformation of its surface was derived. Compared to numerical calculations of the stress tensor, the analytical evaluation is significantly faster. Since the result is given in terms of an expansion into solid spherical harmonics, the overall form of the result to a particular order is the same for all beads, only differing by the coefficients of the expansion. The evaluation of those coefficients, resulting from expansion coefficients of the deformed bead, takes only a few seconds or less on a computer. Moreover, the effort does not increase significantly if more than one bead has to be analyzed, whereas a numerical solution for every single bead can take up to several days \cite{Vorselen}. In future work, one could further extend this method by considering also non-radial deformations \cite{Vorselen}.\\

\section*{Data Availability}
The experimental data analyzed in this study (cf. Fig.\ \ref{fig3}) and the spherical harmonics expansion coefficients used for fitting this data are available online \cite{SI}.

\section*{Author Contributions}
L.J.K.\ performed the derivation. B.W.\ measured the experimental data. L.J.K.\ and S.B.\ analyzed the data and prepared the first version of the figures. L.J.K.\ and M.t.V.\ wrote the first version of the text. T.B.\ and R.W.\ conceived the project and revised the text. M.t.V., T.B., and R.W. revised the figures and supervised the work.

\begin{acknowledgments}
T.B.\ and R.W.\ are funded by the Deutsche Forschungsgemeinschaft (DFG, German Research Foundation) -- Project-IDs BE 6270/2-1 and 433682494 -- SFB 1459.
%R.W.\ is funded by the Deutsche Forschungsgemeinschaft (DFG, German Research Foundation) -- Project-ID 433682494 -- SFB 1459, and T.B.\ is funded by the Deutsche Forschungsgemeinschaft (DFG, German Research Foundation) project BE 6270/2-1.
\end{acknowledgments}

\section*{Declaration of Interest}
There are no conflicts of interest to declare.

\appendix

\section{\label{Derivationansatz}Derivation of \cref{ansatz}}
In this Appendix, we discuss the derivation of \cref{ansatz}. The discussion follows \citet{Love}, but uses a modern notation and focuses on aspects relevant for the present work. 
To solve the differential equation \eqref{de}, a harmonic representation for the complete displacement $\vec{u}(r_,\theta,\phi)$ is derived. \par By evaluating the divergence of \cref{de} and using the fact that the Laplacian $\vec{\nabla}^2 = \vec{\nabla}\cdot \vec{\nabla}$ and the divergence commute, it can be shown that the divergence of $\vec{u}$ is a harmonic function:
\begin{align}\label{eq:love:1}
    0 &= \frac{2-2\nu}{1-2\nu} \vec{\nabla}^2 (\vec{\nabla} \cdot \mathbf{u}).
\end{align}
Harmonic functions can be expanded into solid spherical harmonics. Therefore, $\vec{\nabla}\cdot \vec{u}$ is known to be of the form
\begin{align}\label{eq:love:2}
  \vec{\nabla}\cdot \vec{u} = \sum\limits_{n=0}^{\infty} \Theta_n,
\end{align}
where $\Theta_n$ are SSHs of degree $n$. The differential equation \eqref{de} can then be written in terms of the Laplacian of the displacement $\vec{u}$ and the expansion of $\vec{\nabla}\cdot \vec{u}$:
\begin{align}\label{eq:love:3}
  \vec{\nabla}^2 \vec{u} = \frac{1}{2\nu-1}\sum\limits_{n=0}^{\infty}\vec{\nabla} \Theta_n.
\end{align}
If \cref{de} is written is this form, one can see that it can be solved by finding a particular solution $\vec{u}_p$ for \cref{eq:love:3} and a complementary solution for $\vec{\nabla}^2 \vec{u}_c = 0$. Obviously, the complementary solution $\vec{u}_c$ is also a harmonic function. \\
A particular solution $\vec{u}_p$ to \cref{eq:love:3} is obtained by evaluating the Laplacian of $r^2\vec{\nabla}\Theta_n$. It is used here that the Laplacian of a harmonic function vanishes:
\begin{align}\label{eq:love:4}
  \vec{\nabla}^2 \bigl(r^2 \vec{\nabla} \Theta_n\bigr)
  &= 6 \vec{\nabla}\Theta_n+(4\vec{r}\cdot \vec{\nabla})(\vec{\nabla}\Theta_n)\nonumber\\
  &= 6 \vec{\nabla}\Theta_n+4(n-1)\vec{\nabla}\Theta_n\nonumber\\
  &=(4n+2)\vec{\nabla}\Theta_n.
\end{align}
If we express the harmonic functions $\Theta_n$ in terms of Cartesian coordinates, every component of $\vec{\nabla}\Theta_n$ is also a homogeneous function of degree $n-1$ \cite{Love}. This is used in the second step of \cref{eq:love:4}, where  Euler's theorem on homogeneous functions is applied even though it does not hold for spherical components of SSHs. Therefore, from this point on the derivation does not work for expansions of $\vec{\nabla}\Theta_n$ in curvilinear coordinates. Using \cref{eq:love:4}, a particular solution $\vec{u}_p$ to \cref{eq:love:3} can be constructed:
\begin{align}\label{eq:love:5}
   \vec{u}_p = \frac{1}{2\nu-1}r^2 \sum\limits_{n=0}^{\infty}\frac{1}{2(2n+1)}\vec{\nabla}\Theta_n.
\end{align}
The complete solution for $\vec{u}$ has to satisfy the condition imposed by choosing $\vec{\nabla} \cdot \vec{u} = \sum_{n}\Theta_n$. Using $\vec{u} = \vec{u}_p+\vec{u}_c$, this gives
\begin{align}\label{eq:love:6}
  \vec{\nabla}\cdot \bigl(\vec{u}_p+\vec{u}_c\bigr) &= \sum\limits_{n=0}^{\infty}\Theta_n.
 \end{align}
The particular solution \eqref{eq:love:5}, which is written in terms of $\Theta_n$, is inserted into \cref{eq:love:3}. As regular spherical harmonics of degree $n$ are also homogeneous of degree $n$, Euler's theorem on homogeneous functions can be used: 
\begin{align}\label{eq:love:7}
    \vec{\nabla} \cdot \vec{u}_c 
     &= \sum\limits_{n=0}^{\infty}\Biggl( \frac{3n+1-2\nu(1+2n)}{(2n+1)(1-2\nu)}\Theta_n\Biggr).
\end{align}
As a result, the complete deformation vector $\vec{u}$ can be expressed in terms of the complementary solution $\vec{u}_c$. If $\vec{u}_{c,n}$ is the part of $\vec{u}_c$ which is homogeneous of degree $n$ and therefore consists of SSHs in all of its Cartesian components, the particular solution $\vec{u}_p$ can be written in terms of these harmonic functions:
\begin{align}\label{eq:love:8}
   \vec{u}_p &= \sum\limits_{n=0}^{\infty}r^2\frac{-1}{2[-2\nu(2n+1)+3n+1]}\vec{\nabla}\bigl(\vec{\nabla}\cdot \vec{u}_{c,n+1}\bigr) \nonumber \\ &\equiv \sum\limits_{n=0}^{\infty}r^2M_{n+1}\vec{\nabla}\bigl(\vec{\nabla}\cdot \vec{u}_{c,n+1}\bigr),
\end{align}
where the coefficients $M_n$ are introduced as a shorthand notation:
\begin{equation}\label{eq:love:8_2}
    M_n = \frac{1}{2[2\nu(2n-1)-3n+2]}.
\end{equation}
In the sum \eqref{eq:love:8}, the $n$th term is harmonic of degree $n-1$. The vectors $ \vec{u}_p$ and $ \vec{u}_c$ are added to construct the complete deformation vector $\vec{u}$ in such a way that the $n$th component is harmonic of degree $n$:
\begin{equation}\label{eq:love:9}
    \vec{u} =\sum\limits_{n=0}^{\infty}\bigl( \vec{u}_{c,n}+r^2M_{n+2}\vec{\nabla}(\vec{\nabla}\cdot \vec{u}_{c,n+2})  \bigr).
\end{equation}
The next step is to use the boundary condition for the deformation on the surface, which is specified via an expansion into spherical surface harmonics with coefficients $\vec{A}_n$:
\begin{equation}\label{eq:love:10}
    \vec{u}(r_0) = \sum\limits_{n=0}^{\infty}\vec{A}_n.
\end{equation}
The surface harmonical expansion \eqref{eq:love:10} is expanded again to regular solid harmonics. On the surface $(r=r_0)$, the difference of the derived expression \eqref{eq:love:9} for $\vec{u}$ and the expanded shape of the surface has to vanish:
\begin{equation}\label{eq:love:11}
   \sum\limits_{n=0}^{\infty}\bigg( \vec{u}_{c,n}+r_0^2M_{n+2}\vec{\nabla}\bigl(\vec{\nabla}\cdot \vec{u}_{c,n+2}\bigr)-\frac{r^n}{r_0^n}\vec{A}_n\bigg) = \vec{0}.
\end{equation}
The expression on the left-hand side of \cref{eq:love:11} is a regular harmonic function for $r<r_0$ and vanishes for $r = r_0$. This can only be true if the expression on the left-hand side of \cref{eq:love:11} vanishes for all $r$ and not only on the surface. Thereby, an expression for the $n$th component of $\vec{u}_{c,n}$ is obtained:
\begin{equation}\label{eq:love:12}
  \vec{u}_{c,n}= r_0^2M_{n+2}\vec{\nabla}\bigl(\vec{\nabla}\cdot \vec{u}_{c,n+2}\bigr)+\frac{r^n}{r_0^n}\vec{A}_n.
\end{equation}
Equation \eqref{eq:love:12} is now inserted into the complete expression for the deformation vector $\vec{u}$ given by \cref{eq:love:9}. We can now compare the result
\begin{equation}\label{eq:love:13}
  \vec{u}_{n}= \frac{r^n}{r_0^n}\vec{A}_n+(r^2-r_0^2)M_{n+2}\vec{\nabla}\bigl(\vec{\nabla}\cdot \vec{u}_{c,n+2}\bigr)
\end{equation}
to the ansatz \eqref{ansatz}. Since $r^n\vec{A}_n/r_0^n$ is a harmonic vector function that is equal to the displacement $\vec{u}$ on the surface, the first term on the right-hand side in \cref{eq:love:13} is in agreement with the ansatz \eqref{ansatz}. The expression $\vec{\nabla}\cdot \vec{u}_{c,n+2}$ is a harmonic scalar function of degree $n+1$ if $\vec{u}_{c,n+2}$ is harmonic of degree $n+2$ in its Cartesian components. Consequently, the second term on the right-hand side of \cref{eq:love:13} can be expressed as the gradient of a harmonic scalar function multiplied by $r^2-r_0^2$, which exactly corresponds to the ansatz \eqref{ansatz}.\par
Next, we derive \cref{relation}. To achieve this, we first obtain a connection between the functions $\vec{u}_{c,n}$ and $r^n \vec{A}_n/r_0^n$. These functions can then be expressed with $\psi$ and $\vec{U}$. Taking the divergence of \cref{eq:love:12} and rearranging the result gives
\begin{align}\label{eq:love:14}
   \vec{\nabla}\cdot\vec{u}_{c,n} = \vec{\nabla}\cdot\biggl(\frac{r^n}{r_0^n}\vec{A}_n\biggr).
\end{align}
The $n$th component of the harmonic scalar function $\psi$ is equivalent to $M_{n+1} \vec{\nabla}\cdot \vec{u}_{c,n+1}$. 
 Moreover, the $n$th component of the harmonic vector function $\vec{U}_n$ can be identified with $r^n \vec{A}_n/r_0^n$. Replacing $r^n \vec{A}_n/r_0^n$ and $M_{n+1} \vec{\nabla}\cdot \vec{u}_{c,n+1}$ by $\vec{U}_n$ and $\psi$ in \cref{eq:love:14}, we obtain
\begin{align}\label{eq:love:15}
\vec{\nabla}\cdot\vec{U}_n = \frac{\psi_{n-1}}{M_{n}} ,
\end{align}
which can be solved for $\psi_{n-1}$. Additionally, the definition of $M_n$ as given by \cref{eq:love:8_2} is inserted. This gives
\begin{equation}
    \psi_{n-1} =  \frac{\vec{\nabla}\cdot\vec{U}_n}{2[2\nu(2n-1)-3n+2]},
\end{equation}
 which is equivalent to \cref{relation}.
\section{\label{Ynm} Definition of spherical harmonics}
In the literature, a variety of similar but not equivalent definitions for the spherical harmonic functions are used. Therefore, we here explicitly state the convention used in this article. The spherical harmonics can be defined in terms of the associated Legendre polynomials $P_{l}^{m}$. Here, we use the normalized spherical harmonics \cite{math}
\begin{equation}\label{th:sh:1}
    Y_{l}^{m} = \begin{cases}
    N_{lm}P_{l}^{m}(\cos\theta) \ \text{exp}(\ii m\phi) &  \text{for }m \geq 0,\\
    (-1)^m\bigl[Y_{l}^{m}\bigr]^* & \text{for }m <0
    \end{cases}
\end{equation}
with the polar coordinate $\theta$ ($0 \leq \theta \leq \pi$) and the azimuthal coordinate  $\phi$ ($0 \leq \phi \leq 2\pi$). The factor $N_{lm}$ is given by
\begin{equation}
   N_{lm} =  \sqrt{\frac{2l+1 (l-m)!}{4\pi(l+m)!}}.
\end{equation}
The associated Legendre polynomials $P_{l}^{m}(x)$ are defined by the Legendre polynomials $P_l(x)$ as \cite{math}
\begin{equation}\label{th:sh:2}
    P_{l}^{m}(x) = \begin{cases}
    (1-x^2)^{\frac{m}{2}}\frac{\text{d}^m P_l(x)}{\text{d}x^m} & \text{for }  m \geq 0,\\
    (-1)^m\frac{(l-m)!}{(l+m)!}P_{l}^{m}(x) & \text{for } m <0,
    \end{cases}
\end{equation}
which can be evaluated using Rodrigue's formula \cite{math}
\begin{equation}\label{th:sh:3}
P_l(x) = \frac{1}{2^ll!}\frac{\text{d}^l}{\text{d}x^l}(x^2-1)^l.
\end{equation}

\begin{widetext}

\section{\label{Knm} Transforming $\vec{U}$ into Cartesian coordinates}
\noindent Here, we express the coefficients $\vec{K}_{nm}$ in terms of the coefficients $d_{n,m}$ from the boundary condition and the Clebsch-Gordan coefficients, where the $\vec{\hat{e}}_x,\vec{\hat{e}}_y$ and $\vec{\hat{e}}_z$ are the unit vectors in $x,y$ and $z$ direction respectively:
\begin{align}\label{eq:rad:8}
 \vec{K}_{nm}\cdot \vec{\hat{e}}_x &=  \sqrt{\frac{2\pi}{3}}\Biggl[\bigl(d_{n,m+1}C_{n,m+1,1,-1}^{n,m}-d_{n,m-1}C_{n,m-1,1,-1}^{n,m}\bigr)C_{n,0,1,0}^{n,0}\sqrt{12\pi}\nonumber \\ &\quad +\bigl(d_{n-1,m+1}C_{n-1,m+1,1,-1}^{n,m}-d_{n-1,m-1}C_{n-1,m-1,1,-1}^{n,m}\bigr)C_{n-1,0,1,0}^{n,0}\sqrt{\frac{12\pi(2n-1)}{2n+1}}\nonumber \\ &\quad+\bigl(d_{n+1,m+1}C_{n+1,m+1,1,-1}^{n,m}-d_{n+1,m-1}C_{n+1,m-1,1,-1}^{n,m}\bigr)C_{n+1,0,1,0}^{n,0}\sqrt{\frac{12\pi(2n+3)}{2n+1}}\Biggr],
\end{align}
\begin{align}\label{eq:rad:9}
 \vec{K}_{nm}\cdot \vec{\hat{e}}_y &=  \ii\sqrt{\frac{2\pi}{3}}\Biggl[\bigl(d_{n,m+1}C_{n,m+1,1,-1}^{n,m}+d_{n,m-1}C_{n,m-1,1,-1}^{n,m}\bigr)C_{n,0,1,0}^{n,0}\sqrt{12\pi}\nonumber \\ &\quad+\bigl(d_{n-1,m+1}C_{n-1,m+1,1,-1}^{n,m}+d_{n-1,m-1}C_{n-1,m-1,1,-1}^{n,m}\bigr)C_{n-1,0,1,0}^{n,0}\sqrt{\frac{12\pi(2n-1)}{2n+1}}\nonumber \\ &\quad+\bigl(d_{n+1,m+1}C_{n+1,m+1,1,-1}^{n,m}+d_{n+1,m-1}C_{n+1,m-1,1,-1}^{n,m}\bigr)C_{n+1,0,1,0}^{n,0}\sqrt{\frac{12\pi(2n+3)}{2n+1}}\Biggr],
\end{align}
\begin{align}\label{eq:rad:10}
 \vec{K}_{nm}\cdot \vec{\hat{e}}_z &=  2\sqrt{\frac{\pi}{3}}\Biggl[d_{n,m}C_{n,m,1,0}^{n,m}C_{n,0,1,0}^{n,0}\sqrt{12\pi}+d_{n-1,m}C_{n-1,m,1,0}^{n,m}C_{n-1,0,1,0}^{n,0}\sqrt{\frac{12\pi(2n-1)}{2n+1}}\nonumber \\ &\quad+d_{n+1,m}C_{n+1,m,1,0}^{n,m}C_{n+1,0,1,0}^{n,0}\sqrt{\frac{12\pi(2n+3)}{2n+1}}\Biggr].
\end{align}
\section{\label{derivatives} Derivatives of SSHs with respect to Cartesian coordinates.}
\noindent The derivatives of SSHs with respect to Cartesian coordinates can be derived by using various recurrence relations for Legendre polynomials and associated Legendre polynomials. The derivatives are given by
\begin{align}\label{eq:derivatives}
    \frac{\partial\bigl( r^nY_n^m(\theta,\phi)\bigr)}{\partial x} &= \frac{N_{n,m}}{2}\Biggl[\frac{Y_{n-1,m+1}}{N_{n-1,m-1}}-(n+m)(m-1+n)\frac{Y_{n-1,m-1}}{N_{n-1,m-1}}\Biggr],\\ 
    \label{eq:derivatives2}\frac{\partial\bigl( r^nY_n^m(\theta,\phi)\bigr)}{\partial y} &= \frac{N_{n,m}}{2}i\Biggl[-\frac{Y_{n-1,m+1}}{N_{n-1,m-1}}-(n+m)(m-1+n)\frac{Y_{n-1,m-1}}{N_{n-1,m-1}}\Biggr],\\
    \label{eq:derivatives3}
\frac{\partial\bigl( r^nY_n^m(\theta,\phi)\bigr)}{\partial z} &=\frac{N_{n,m}}{N_{n-1,m}}(n+m)Y_{n-1}^m .
\end{align}
Using the derivatives \eqref{eq:derivatives}-\eqref{eq:derivatives3}, the divergence of $\vec{U}_n$ can be easily expressed as an expansion into SSHs, where SSHs of the same order are grouped together:
\begin{align}\label{eq:co:psi2}
 \vec{\nabla}\cdot\vec{U}_n &= \sum\limits_{m=-n}^{n}\frac{r^{n-1}Y_{n-1,m}}{r_0^nN_{n-1,m}}\frac{1}{2}\Biggl[-(m+n)(m+n+1)N_{n,m+1}\bigl(K_{x,n,m+1}+\ii K_{y,n,m+1}\bigr)\nonumber \\ &\quad + N_{n,m-1}\bigl(K_{x,n,m-1}-\ii K_{y,n,m-1}\bigr)+2N_{n,m}(n+m)K_{z,n,m}\Biggr] \nonumber \\
  &\equiv \sum\limits_{m=-n+1}^{n-1}\frac{r^{n-1}Y_{n-1,m}}{r_0^{n-1}} c_{nm},
\end{align}
where the coefficients $c_{nm}$ are defined as:
\begin{align}\label{eq:cnm}
    c_{nm} = \frac{1}{2N_{n-1,m}}\Biggl[&-(m+n)(m+n+1)N_{n,m+1}\bigl(K_{x,n,m+1}+\ii K_{y,n,m+1}\bigr)\nonumber \\ &+ N_{n,m-1}\bigl(K_{x,n,m-1}-\ii K_{y,n,m-1}\bigr)+2N_{n,m}(n+m)K_{z,n,m}\Biggr].
\end{align}
From this expansion, one can find the coefficients $a_{nm}$ defining the harmonic scalar function $\psi$ as given in \cref{anm}. 
\end{widetext}
% \clearpage

\bibliography{refs}
\end{document}
