## Supplementary material for "Analytical method for reconstructing the stress on a spherical particle from its surface deformation": TeX files: refs.bib

%% Literatursammlung vom 22.11.2019 von MtV und RW

@STRING{Aa={Aachen}}
@STRING{AaA={Astronomy and Astrophysics}}
@STRING{AAftAoS={American Association for the Advancement of Science}} 
@STRING{AAoPT={American Association of Physics Teachers}} 
@STRING{ACS={American Chemical Society}}
@STRING{ACIE={Angewandte Chemie International Edition}}
@STRING{ACMToMS={ACM Transactions on Mathematical Software}} 
@STRING{AdPD={Annalen der Physik}} 
@STRING{AdPF={Annales de Physique}}
@STRING{AdvM={Advanced Materials}}
@STRING{AfCM={Association for Computing Machinery}}
@STRING{AFM={Advanced Functional Materials}}
@STRING{AfRMaA={Archive for Rational Mechanics and Analysis}} 
@STRING{AiAM={Advances in Applied Mechanics}}
@STRING{AIoP={American Institute of Physics}}
@STRING{AiP={Advances in Physics}}
@STRING{AJ={Astronomical Journal}}
@STRING{AJoP={American Journal of Physics}}
@STRING{AJoTMaH={American Journal of Tropical Medicine and Hygiene}}
@STRING{AphyJ={Astrophysical Journal}}
@STRING{AJSS={Astrophysical Journal Supplement Series}}
@STRING{AM={Acta Mechanica}}
@STRING{AMatS={American Mathematical Society}}
@STRING{AMatST={American Mathematical Society Translations}}
@STRING{AMS={American Meteorological Society}}
@STRING{AMST={Amsterdam}}
@STRING{AMT={Applied Materials Today}}
@STRING{AoNS={Archive of Numerical Software}}
@STRING{AoP={Annals of Physics}}
@STRING{AotNYAoS={Annals of the New York Academy of Sciences}} 
@STRING{AP={Academic Press}} 
@STRING{APhy={Astroparticle Physics}}
@STRING{AphysJ={Astrophysical Journal}}
@STRING{AphysJL={Astrophysical Journal Letters}}
@STRING{APL={Applied Physics Letters}}
@STRING{APS={American Physical Society}}
@STRING{AR={Annual Reviews}} 
@STRING{ARes={Anticancer Research}} 
@STRING{ARoAaA={Annual Review of Astronomy and Astrophysics}} 
@STRING{ARoCMP={Annual Review of Condensed Matter Physics}}
@STRING{ARoFM={Annual Review of Fluid Mechanics}}
@STRING{ARoPC={Annual Review of Physical Chemistry}}
@STRING{ASfM={American Society for Microbiology}} 
@STRING{ASoTMaH={American Society of Tropical Medicine and Hygiene}}
@STRING{ASoZ={American Society of Zoologists}} 
@STRING{AV={Akademie Verlag}}
@STRING{AVGuP={Akademische Verlagsgesellschaft Geest \& Portig}}
@STRING{AW={Addison Wesley}} 
@STRING{AZ={American Zoologist}}
@STRING{BAMS={Bulletin American Meteorological Society}}
@STRING{BddcG={Berichte der deutschen chemischen Gesellschaft}}
@STRING{BER={Berlin}}
@STRING{BH={Butterworth-Heinemann}}
@STRING{Bio={Biopolymers}}
@STRING{BiuZ={Biologie in unserer Zeit}}
@STRING{BJ={Biophysical Journal}} 
@STRING{BM={Biomedical Microdevices}}
@STRING{BRA={Braunschweig}}
@STRING{BrWi={Braunschweig, Wiesbaden}}
@STRING{CAM={Cambridge}}
@STRING{CaPS={Colloid \& Polymer Science}}
@STRING{CaQG={Classical and Quantum Gravity}}
@STRING{CaSA:PaEA={Colloids and Surfaces A: Physicochemical and Engineering Aspects}}
@STRING{CB={Current Biology}}
@STRING{CellP={Cell Press}}
@STRING{CES={Chemical Engineering Science}}
@STRING{ChemP={Chemical Physics}}
@STRING{CHI={Chichester}}
@STRING{CiMS={Communications in Mathematical Sciences}}
@STRING{CJoP={Czechoslovak Journal of Physics}}
@STRING{CM={Computational Mathematics}}
@STRING{CMES={CMES-Computer Modelling in Engineering \& Sciences}}
@STRING{CMJ={College Mathematics Journal}}
@STRING{CM:NTaA={Computational Mechanics: New Trends and Applications}}
@STRING{CMP={Condensed Matter Physics}}
@STRING{CMS={Computational Materials Science}}
@STRING{COiCaIS={Current Opinion in Colloid \& Interface Science}}
@STRING{CoM={Chemistry of Materials}}
@STRING{CoPaAM={Communications on Pure and Applied Mathematics}}
@STRING{CP={Clarendon Press}}
@STRING{CPL={Chemical Physics Letters}}
@STRING{CR={Chemical Reviews}}
@STRING{CRCP={CRC Press}}
@STRING{CRdlAdS={Comptes Rendus de l'Acad{\'e}mie des Sciences}}
@STRING{CRdlAdS:SII={Comptes Rendus de l'Acad{\'e}mie des Sciences - Series II}}
@STRING{CSR={Chemical Society Reviews}}
@STRING{CtPP={Contributions to Plasma Physics}}
@STRING{CUP={Cambridge University Press}} 
@STRING{DANSSSR={Doklady Akademii Nauk SSSR}}
@STRING{DOR={Dordrecht}} 
@STRING{DotFS={Discussions of the Faraday Society}} 
@STRING{DP={Dover Publications}} 
@STRING{EAP={Elsevier Academic Publishers}}
@STRING{EBJ={European Biophysics Journal}} 
@STRING{EDPS={EDP Sciences}}
@STRING{EFD={Elementary Fluid Dynamics}}
@STRING{EI={Elsevier Insights}}
@STRING{EiF={Experiments in Fluids}}
@STRING{EJoP={European Journal of Physics}}
@STRING{eL={eLife}}
@STRING{eLSP={eLife Sciences Publications}}
@STRING{ELS={Elsevier}}
@STRING{ELSAP={Elsevier Academic Press}}
@STRING{EPJ={European Physical Journal}}
@STRING{EPJB={European Physical Journal B}}
@STRING{EPJE={European Physical Journal E}}
@STRING{EPJE:SMaBP={European Physical Journal E: Soft Matter and Biological Physics}}
@STRING{EPJST={European Physical Journal Special Topics}}
@STRING{EPL={Europhysics Letters}}
@STRING{EPS={European Physical Society}}
@STRING{ES={Elsevier Science}}
@STRING{EST={Elsevier Science \& Technology}}
@STRING{FaM={Frankfurt am Main}}
@STRING{FAoS={French Academy of Sciences}}
@STRING{FDotCS={Faraday Discussions of the Chemical Society}}
@STRING{FS={Faraday Society}}
@STRING{FSaE={Fusion Science and Engineering}}
@STRING{FSaT={Fusion Science and Technology}}
@STRING{FT={Fusion Technology}}
@STRING{GRL={Geophysical Research Letters}}
@STRING{GTV={Georg Thieme Verlag}}
@STRING{HAP={Harwood Academic Publishers}} 
@STRING{HD={Verlag Harri Deutsch}} 
@STRING{ICARUS={Icarus}}
@STRING{ICP={Imperial College Press}}
@STRING{IEEEJoQE={IEEE Journal of Quantum Electronics}} 
@STRING{IEEEMS={IEEE Magnetics Society}}
@STRING{IEEEToM={IEEE Transactions on Magnetics}}
@STRING{IF={Interface Focus}}
@STRING{IfCMP={Institute for Condensed Matter Physics}}
@STRING{IIoAR={International Institute of Anticancer Research}}
@STRING{IJfNMiE={International Journal for Numerical Methods in Engineering}}
@STRING{IJoCSaE={International Journal of Computational Science and Engineering}}
@STRING{IJoES={International Journal of Engineering Science}}
@STRING{IJoMPC={International Journal of Modern Physics C}}
@STRING{IJoQC={International Journal of Quantum Chemistry}}
@STRING{INCB={Il Nuovo Cimento B (1971-1996)}}
@STRING{IndP={Inderscience Publishers}}
@STRING{IoP={Institute of Physics}}
@STRING{IoPP={Institute of Physics Publishing}}
@STRING{IP={International Press}}
@STRING{IPS={Italian Physical Society}}
@STRING{JdP={Journal de Physique}} 
@STRING{JdPC={Journal de Physique Colloques}}
@STRING{JdPeLR={Journal de Physique et Le Radium}}
@STRING{JdPII={Journal de Physique II}} 
@STRING{JdPL={Journal de Physique Lettres}}
@STRING{JJoAP={Japanese Journal of Applied Physics}}
@STRING{JoAP={Journal of Applied Physics}}
@STRING{JoB={Journal of Bacteriology}}
@STRING{JoBC={Journal of Biological Chemistry}}
@STRING{JoBNMR={Journal of Biomolecular NMR}}
@STRING{JoCaAP={Journal of Cosmology and Astroparticle Physics}}
@STRING{JoCaIS={Journal of Colloid and Interface Science}} 
@STRING{JoChemP={Journal of Chemical Physics}} 
@STRING{JoCompP={Journal of Computational Physics}}
@STRING{JoCR={Journal of Controlled Release}}
@STRING{JoCS={Journal of Colloid Science}}
@STRING{JoEaTP={Journal of Experimental and Theoretical Physics}}
@STRING{JoEaTPL={Journal of Experimental and Theoretical Physics Letters}}
@STRING{JoEM={Journal of Engineering Mathematics}}
@STRING{JoFM={Journal of Fluid Mechanics}}
@STRING{JoGR={Journal of Geophysical Research}}
@STRING{JoLTP={Journal of Low Temperature Physics}}
@STRING{JoMaM={Journal of Micromechanics and Microengineering}}
@STRING{JoMaMM={Journal of Magnetism and Magnetic Materials}}
@STRING{JoMC={Journal of Materials Chemistry}}
@STRING{JoMP={Journal of Mathematical Physics}}
@STRING{JoMR={Journal of Magnetic Resonance}}
@STRING{JoMS={Journal of Materials Science}}
@STRING{JoN={Journal of Nanobiotechnology}}
@STRING{JoNM={Journal of Nuclear Materials}}
@STRING{JoNNFM={Journal of Non-Newtonian Fluid Mechanics}} 
@STRING{JoNO={Journal of Neuro-Oncology}}
@STRING{JoP={Journal of Physics}}
@STRING{JoPA={Journal of Physics A}} 
@STRING{JoPA:MaG={Journal of Physics A: Mathematical and General}}
@STRING{JoPA:MaT={Journal of Physics A: Mathematical and Theoretical}}
@STRING{JoPB={Journal of Physics B}}
@STRING{JoPC={Journal of Physics C}}
@STRING{JoPChem={Journal of Physical Chemistry}}
@STRING{JoPChemB={Journal of Physical Chemistry B}}
@STRING{JoP:CS={Journal of Physics: Conference Series}}
@STRING{JoPC:SSP={Journal of Physics C: Solid State Physics}}
@STRING{JoP:CM={Journal of Physics: Condensed Matter}}
@STRING{JoPD={Journal of Physics D}}
@STRING{JoPE={Journal of Physics E}}
@STRING{JoPF={Journal of Physics F}}
@STRING{JoPG={Journal of Physics G}}
@STRING{JoPG:NaPP={Journal of Physics G: Nuclear and Particle Physics}}
@STRING{JoPP={Journal of Plasma Physics}} 
@STRING{JoPSaE={Journal of Petroleum Science and Engineering}}
@STRING{JoR={Journal of Rheology}}
@STRING{JoSM:TaE={Journal of Statistical Mechanics: Theory and Experiment}}
@STRING{JotACS={Journal of the American Chemical Society}}
@STRING{JotAS={Journal of the Atmospheric Sciences}}
@STRING{JoTB={Journal of Theoretical Biology}}
@STRING{JotCSFT={Journal of the Chemical Society, Faraday Transactions}}
@STRING{JotPSoJ={Journal of the Physical Society of Japan}}
@STRING{JotSfIaAM={Journal of the Society for Industrial and Applied Mathematics}}
@STRING{JotSfID={Journal of the Society for Information Display}}
@STRING{JSoAP={Japan Society of Applied Physics}}
@STRING{KAP={Kluwer Academic Publishers}}
@STRING{LANG={Langmuir}}
@STRING{LC={Liquid Crystals}}
@STRING{LoaC={Lab on a Chip}}
@STRING{LON={London}}
@STRING{LSdJdPeLR={La Soci\'{e}t\'{e} du Journal de Physique et Le Radium}}
@STRING{MAC={Macromolecules}}
@STRING{MaN={Microfluidics and Nanofluidics}}
@STRING{MAoA={Mathematical Association of America}}
@STRING{MCaLC={Molecular Crystals and Liquid Crystals}}
@STRING{MCaLCL={Molecular Crystals and Liquid Crystals Letters}}
@STRING{MD={Marcel Dekker}}
@STRING{MNotRAS={Monthly Notices of the Royal Astronomical Society}}
@STRING{MP={Molecular Physics}}
@STRING{MRC={Macromolecular Rapid Communications}}
@STRING{MT={Materials Today}}
@STRING{NANO={ACS Nano}}
@STRING{NAoS={National Academy of Sciences}}
@STRING{NAT={Nature}}
@STRING{NC={Nature Chemistry}} 
@STRING{NCommun={Nature Communications}} 
@STRING{NdAdWiG={Nachrichten der Akademie der Wissenschaften in G{\"o}ttingen}}
@STRING{NdKGdWzG={Nachrichten der K{\"o}niglichen Gesellschaft der Wissenschaften zu G{\"o}ttingen}}
@STRING{NF={Nuclear Fusion}} 
@STRING{NH={North-Holland Publishing Company}} 
@STRING{NJoP={New Journal of Physics}}
@STRING{NL={Nano Letters}} 
@STRING{NM={Nature Materials}} 
@STRING{NP={Nature Physics}}
@STRING{NPG={Nature Publishing Group, a division of Macmillan Publishers Limited}}
@STRING{NT={Nano Today}}
@STRING{NY={New York}}
@STRING{NYAoS={New York Academy of Sciences}}
@STRING{OE={Optics Express}}
@STRING{OSoA={Optical Society of America}}
@STRING{OSP={Oxford Science Publications}}
@STRING{OUP={Oxford University Press}}
@STRING{OX={Oxford}}
@STRING{PA={Physica A}}
@STRING{PaCoL={Physics and Chemistry of Liquids}}
@STRING{PAR={Paris}}
@STRING{PA:SMaiA={Physica A: Statistical Mechanics and its Applications}}
@STRING{PB={Physica B}}
@STRING{PB:CM={Physica B: Condensed Matter}}
@STRING{PBP={Perseus Books Publishing}}
@STRING{PC={Physica C}}
@STRING{PCCP={Physical Chemistry Chemical Physics}}
@STRING{PD={Physica D}} 
@STRING{PD:NP={Physica D: Nonlinear Phenomena}} 
@STRING{PHY={Physics}}
@STRING{PiCaPS={Progress in Colloid and Polymer Science}}
@STRING{PJ={Physik Journal}}
@STRING{PLA={Physics Letters A}}
@STRING{PLoS={Public Library of Science}}
@STRING{PM={Physicalia Magazine}}
@STRING{PoASaT={Problems of Atomic Science and Technology}}
@STRING{PoF={Physics of Fluids}}
@STRING{PoFB={Physics of Fluids B}}
@STRING{POne={PLOS ONE}}
@STRING{PoP={Physics of Plasmas}}
@STRING{PotNAoS={Proceedings of the National Academy of Sciences U.S.A.}}
@STRING{PoTP={Progress of Theoretical Physics}}
@STRING{PotRNAoAaSaA={Proceedings of the Royal Netherlands Academy of Arts and Sciences at Amsterdam}}
@STRING{PotRSoL={Proceedings of the Royal Society of London}}
@STRING{PotRSoLA:CPoaMaPC={Proceedings of the Royal Society of London A: Containing Papers of a Mathematical and Physical Character}}
@STRING{PotRSoLSACPoaMaPC={Proceedings of the Royal Society of London. Series A, Containing Papers of a Mathematical and Physical Character}}
@STRING{PP={Pleiades Publishing}}
@STRING{PPaCF={Plasma Physics and Controlled Fusion}}
@STRING{PR={Physical Review}}
@STRING{PRA={Physical Review A}}
@STRING{PRB={Physical Review B}}
@STRING{PRC={Physical Review C}}
@STRING{PRD={Physical Review D}}
@STRING{PRE={Physical Review E}}
@STRING{PRF={Physical Review Fluids}}
@STRING{PRL={Physical Review Letters}}
@STRING{PRS={Physics Reports}}
@STRING{PS={Physica Scripta}}
@STRING{PSoJ={Physical Society of Japan}}
@STRING{PST={Physica Scripta T}}
@STRING{PT={Physics Today}}
@STRING{PTotRSA:MPaES={Philosophical Transactions of the Royal Society A: Mathematical, Physical and Engineering Sciences}}
@STRING{PU={Physics Uspekhi}} 
@STRING{QJoMaAM={Quarterly Journal of Mechanics and Applied Mathematics}}
@STRING{QRoB={Quarterly Reviews of Biophysics}}
@STRING{RA={Rheologica Acta}}
@STRING{RMS={Russian Mathematical Surveys}}
@STRING{RNAoAaS={Royal Netherlands Academy of Arts and Sciences}}
@STRING{RoGaSP={Reviews of Geophysics and Space Physics}}
@STRING{RoMP={Reviews of Modern Physics}}
@STRING{RoMPB={Reviews of Modern Physics B}}
@STRING{RoMPL={Reviews of Modern Physics Letters}}
@STRING{RoPiP={Reports on Progress in Physics}}
@STRING{RS={Royal Society}}
@STRING{RSCP={RSC Publishing}}
@STRING{RSoC={Royal Society of Chemistry}}
@STRING{RSoLPSA={Royal Society of London Proceedings Series A}}
@STRING{RSoLPTSA={Royal Society of London Philosophical Transactions Series A}}
@STRING{SA={Science Advances}}
@STRING{SaToAM={Science and Technology of Advanced Materials}}
@STRING{SAV={Spektrum Akademischer Verlag}}
@STRING{SCI={Science}}
@STRING{SfIaAM={Society for Industrial and Applied Mathematics}}
@STRING{SfIaCB={Society for Integrative and Comparative Biology}}
@STRING{SfID={Society for Information Display}}
@STRING{SIN={Singapore}}
@STRING{SM={Soft Matter}}
@STRING{SMALL={Small}}
@STRING{SPR={Springer-Verlag}}
@STRING{SPSS={Soviet Physics - Solid State}}
@STRING{SPU={Soviet Physics Uspekhi}}
@STRING{SR={Scientific Reports}}
@STRING{SS={Springer Spektrum}}
@STRING{STU={Stuttgart}} 
@STRING{SV={Shaker Verlag}}
@STRING{TEU={B. G. Teubner}}
@STRING{TF={Taylor \& Francis}}
@STRING{TotFS={Transactions of the Faraday Society}}
@STRING{TSP={Tech Science Press}}
@STRING{USSRAoS={USSR Academy of Sciences}}
@STRING{VAC={Vacuum}}
@STRING{VdZfN={Verlag der Zeitschrift f{\"u}r Naturforschung}}
@STRING{VIE={Vieweg Verlag}}
@STRING{VvWE={Verlag von Wilhelm Engelmann}}
@STRING{WEI={Weinheim}}
@STRING{WIL={John Wiley \& Sons}}
@STRING{WileyP={Wiley Periodicals, Inc.}}
@STRING{WILEYVCH={WILEY-VCH Verlag GmbH \& Co. KGaA}}
@STRING{WSP={World Scientific Publishing}}
@STRING{WT={Wiley \& Teubner}}
@STRING{YHaKU={Yukawa Hall \& Kyoto University}}
@STRING{ZEiTF={Zhurnal \'{E}ksperimental'no\u{i} i Teoretichesko\u{i} Fiziki}}
@STRING{ZfN={Zeitschrift f{\"u}r Naturforschung}}
@STRING{ZfNTA={Zeitschrift f{\"u}r Naturforschung Teil A}}
@STRING{ZfP={Zeitschrift f{\"u}r Physik}}
@STRING{ZfPA:HaN={Zeitschrift f{\"u}r Physik A: Hadrons and Nuclei}}
@STRING{ZfPB:CM={Zeitschrift f{\"u}r Physik B: Condensed Matter}} 
@STRING{ZfPC={Zeitschrift f{\"u}r Physikalische Chemie}}
@STRING{ZfpCSuV={Zeitschrift f{\"u}r physikalische Chemie, St{\"o}chiometrie und Verwandtschaftslehre}}
%STRING{={}}
@article{Nagle,
author = {Nagle, I and Delort, Florence and H\'enon, Sylvie and Wilhelm, Claire and Batonnet-Pichon, Sabrina and Reffay, Myriam},
year = {2022},
month = {12},
title = {The importance of intermediate filaments in the shape maintenance of myoblast model tissues},
volume = {11},
journal = {eLife},
pages = {e76409}
}
@article{Meglio,
author = {Meglio, Ilaria and Trushko, Anastasiya and Guillamat, Pau and Blanch-Mercader, Carles and Abuhattum, Shada and Roux, Aur\'elien},
year = {2022},
month = {08},
pages = {111227},
title = {Pressure and curvature control of the cell cycle in epithelia growing under spherical confinement},
volume = {40},
journal = {Cell Reports}
}
@article{Discher,
author = {Discher, D E and Janmey, Pand  Wang, Y L},
year = {2005},
month = {11},
title = {Tissue cells feel and respond to the stiffness of their substrate},
volume = {310},
journal = {Science},
pages = {1139-1143}
}
@article{Sporrer,
author = {Sp\"orrer, M. and Kah, D. and Gerum, R. and Reischl, B. and Huraskin, D.and Dessalles, C. A. and Fabry, B. },
year = {2021},
title = {The Desmin Mutation {R349P} Increases Contractility and Fragility Of Stem Cell-generated Muscle Micro-tissues},
volume = {3},
number ={48},
journal = {Neuropathology and Applied Neurobiology},
pages = {e12784}
}
@article{Goldmann,
author = {Goldmann, Wolfgang H},
title = {Mechanotransduction in cells},
journal = {Cell Biology International},
volume = {36},
number = {6},
pages = {567-570},
doi = {https://doi.org/10.1042/CBI20120071},
year = {2012}
}

@article{Valerie,
author = {Valerie S. LeBleu and Brian MacDonald and Raghu Kalluri},
title ={Structure and Function of Basement Membranes},
journal = {Experimental Biology and Medicine},
volume = {232},
number = {9},
pages = {1121-1129},
year = {2007}
}

% doi = {10.3181/0703-MR-72},

URL = { 
        https://doi.org/10.3181/0703-MR-72
    
},
eprint = { 
        https://doi.org/10.3181/0703-MR-72
}

 @BOOK{Lurie,
   title        = {Theory of Elasticity},
   author       = {I. A. Lurie},
   year         = {2005},
   publisher    = SPR,
   address      = {Berlin Heidelberg New York},
   isbn         = {3-540-24556-1 }
}
%    pages        = {1050},

@BOOK{Lifschitz,
   title        = {Theory of Elasticity},
   author       = {L. D. Landau and E. M. Lifshitz},
   year         = {1970},
   publisher    = {Pergamon Press},
   volume       = {7},
   address      = {Oxford},
   isbn         = {978-0750626330}
}
%   pages        = {165},

@incollection{Trefftz,
   booktitle        = {Mechanik der Elastischen {K}\"orper},
   editor       = {G. Angenheister},
   year         = {1928},
   publisher    = {Springer-Verlag},
   address      = {Heidelberg},
   isbn         = {978-0-511-48036-2},
   title={Mathematische {E}lastizit\"atstheorie},
   author = {Trefftz, E.},
   pages ={47-140}
}


@BOOK{Red,
   title        = {An Introduction to Continuum Mechanics},
   author       = {J. N. Reddy},
   year         = {2008},
   publisher    = {Cambridge University Press},
   address      = {Cambridge}
}

@BOOK{DeB,
   title        = {Handbook of Linear and Partial Differential Equations for Engineers and Scientists},
   author       = {Andrei D. Polyanin},
   year         = {2002},
   publisher    = {Chapman and Hall/CRC},
   pages        = {667},
   address      = {Florida},
   isbn         = {ISBN 1-58488-299-9}

}
@BOOK{math,
   title        = {Mathematical Methods for Physics and Engineering},
   author       = {K. F. Riley and M. P. Hobson and S. J. Bence},
   year         = {2006},
   publisher    = {Cambridge University Press},
   address      = {Cambridge},
   isbn         = {978-0-521-67971-8}

}
% pages        = {1333},
@article{Dolega,
   author       = {Dolega, M. and Delarue, M. and Ingremeau, F. and Prost, J and Delon, A and Cappello, G},
   title        = {Cell-like pressure sensors reveal increase of mechanical stress towards the core of multicellular spheroids under compression},
   journal      = NCommun,
   publisher    = NPG,
   volume       = {8},
   pages       = {14056},
   year         = {2017}
}
@article{Mohagheghian,
   author       = {Mohagheghian, E. and Luo, J. and Chen, J. and G Chaudhary and J Chen and J Sun and R. H. Ewoldt and N Wang},
   title        = {Quantifying compressive forces between living cell layers and within tissues using elastic round microgels},
   journal      = NCommun,
   publisher    = NPG,
   volume       = {9},
   number       = {1878},
   year         = {2018}
}
@BOOK{Love,
   title        = {A Treatise on the Mathematical Theory of Elasticity},
   author       = {Love, A. E. H.},
   year         = {1892},
   publisher    = {Cambridge University Press},
   address      = {Cambridge}
}  
%    pages        = {348},

@BOOK{Shankar,
   title        = {Principles of Quantum Mechanics},
   author       = {R. Shankar},
   year         = {1994},
   publisher    = {Springer},
   pages        = {676},
   address      = {Connecticut},
   isbn         = {978-0-306-44790-7},
}   
@BOOK{Hay,
   title        = {Cell Biology of Extracellular Matrix},
   author       = {Hay, E. D.},
   year         = {1991},
   publisher    = {Springer},
   address      = {New York},
   isbn         = {978-0-306-43951-3}
}   
%    pages        = {468},

@BOOK{Reddy2,
   title        = {An Introduction to the Finite Element Method},
   author       = {J. N. Reddy},
   year         = {1993},
   publisher    = {MCGraw-Hill, Inc},
   pages        = {684},
   isbn         = {0-07-051355-4},
}   
@BOOK{wood,
   title        = {Nonlinear Continuum Mechanics for Finite Element Analysis},
   author       = {J. Bonet and R. D. Wood},
   year         = {1997},
   publisher    = {Cambridge University press},
   pages        = {318},
   isbn         = {978-0-521-83870-2},
} 
@BOOK{varsh,
   title        = {Quantum Theory of Angular Momentum},
   author       = {D. A. Varshalovich and A. N. Moskalev and V. K. Khersonskii},
   year         = {1988},
  address={Singapore},
   publisher    = {World Scientific},
   isbn         = {978-9971-5-0107-5 }
} 
%    pages        = {528},
@article{Vorselen,
   author       = {Vorselen, D. and Wang, Y. and  {de Jesus}, M. M. and Shah, P K and Footer, M J and Huse, M and Cai, W and Theriot, J A},
   title        = {Microparticle traction force microscopy reveals subcellular force exertion patterns in immune cell-target interactions},
   journal      = NCommun,
   publisher    = NPG,
   volume       = {11},
   pages      = {20},
   year         = {2020}
}
@article{Biswas,
  author={Biswas, Arikta and Ng, Boon Heng and Prabhakaran, Vinod S/O and Chan, Chii Jou},   
	 title={Squeezing the eggs to grow: The mechanobiology of mammalian folliculogenesis},      
	journal={Frontiers in Cell and Developmental Biology},      
	volume={10},           
	year={2022},   
pages={1038107}   
}
% 	  URL={https://www.frontiersin.org/articles/10.3389/fcell.2022.1038107},       
%DOI={10.3389/fcell.2022.1038107},  

@article{MALMIKAKKADA,
title = {Adhesion strength between cells regulate nonmonotonic growth by a biomechanical feedback mechanism},
journal = {Biophysical Journal},
volume = {121},
number = {19},
pages = {3719-3729},
year = {2022},
issn = {0006-3495},
doi = {https://doi.org/10.1016/j.bpj.2022.04.032},
url = {https://www.sciencedirect.com/science/article/pii/S0006349522003605},
author = {Abdul N. Malmi-Kakkada and Sumit Sinha and Xin Li and D. Thirumalai},

}
@article{
Mohaghegian2,
author = {Erfan Mohagheghian  and Junyu Luo  and F. Max Yavitt  and Fuxiang Wei  and Parth Bhala  and Kshitij Amar  and Fazlur Rashid  and Yuzheng Wang  and Xingchen Liu  and Chenyang Ji  and Junwei Chen  and David P. Arnold  and Zhen Liu  and Kristi S. Anseth  and Ning Wang },
title = {Quantifying stiffness and forces of tumor colonies and embryos using a magnetic microrobot},
journal = {Science Robotics},
volume = {8},
number = {74},
pages = {eadc9800},
year = {2023},
doi = {10.1126/scirobotics.adc9800},
URL = {https://www.science.org/doi/abs/10.1126/scirobotics.adc9800}}
@article{Lee,
   author       = {Lee, W. and Kalashnikov, N. and Mok, S. and Ruba Halaoui and Elena Kuzmin and Andrew J. Putnam and Shuichi Takayama and Morag Park and Luke McCaffrey and Ruogang Zhao and Richard L. Leask and Christopher Moraes},
   title        = {Dispersible hydrogel force sensors reveal patterns of solid mechanical stress in multicellular spheroid cultures},
   journal      = NCommun,
   publisher    = NPG,
   volume       = {10},
   pages       = {144},
   year         = {2019}
}
@article{Traber,
   author       = {Tr\"aber, N. and Uhlmann, K. and Girardo S. and others},
   title        = {Polyacrylamide Bead Sensors for in vivo Quantification of Cell-Scale Stress in Zebrafish Development},
   journal      = SR,
   publisher    = NPG,
   volume       = {9},
   pages      = {17031},
   year         = {2019}
}
@article{Handorf,
   author       = {Andrew M Handorf and Yaxian Zhou and Matthew A Halanski and Wan-Ju Li},
   title        = {Tissue Stiffness Dictates Development, Homeostasis, and Disease Progressiont},
   journal      = {Organogenesis},
   publisher    = {Landes Bioscience},
   volume       = {11},
   pages={1-15},
   year         = {2015}
}
@article{Frantz,
   author       = {Christian Frantz and Kathleen M. Stewart and Valerie M. Weaver},
   title        = {The extracellular matrix at a glance},
   journal      = {Journal of Cell Science},
   publisher    = {The Company of Biologists Ltd},
   volume       = {123},
   pages        = {4195-4200},
   year         = {2010}
}
@article{Juhl,
   author       = {Juhl, O. J., Buettmann, E. G., Friedman, M. A. and others},
   title        = {Update on the effects of microgravity on the musculoskeletal system},
   journal      = {npj Microgravity },
   publisher    = {The Company of Biologists Ltd},
   volume       = {7,28},
   year         = {2021}
}
@article{Sugimura,
    author = {K Sugimura and Y Bella\"{i}che and F Graner},
    title = {Robustness of force and stress inference in an epithelial tissue},
    journal= {Annu Int Conf IEEE Eng Med Biol Soc},
    year = {2013},
}

@article{Courant,
    author = {R. Courant},
    title = {VARIATIONAL METHODS FOR THE SOLUTION OF PROBLEMS OF EQUILIBRIUM AND VIBRATIONS },
    journal= {Bull. Amer. Math. Soc. },
    volume = {49(1)},
    pages = {1-23},
    year = {1943},
    month = JAN
}
@article{Turner,
    author = {Turner, M. J. and Clough, R. W. and Martin, H. C. and Topp, L. J.},
    title = {Stiffness and Deflection Analysis of Complex Structures  },
    journal= {Journal of the Aeronautical Sciences },
    volume = {23(9)},
    pages = {805-823},
    year = {1956},
    month = SEP
}
@article{Campas,
    author = {Ottger Camp\`{a}s and Tadanori Mammoto and Sean Hasso and R A Sperling and D {O'Connell} and A G Bischof and R Maas and D A Weitz and L Mahadevan and D E Ingber},
    title = {Quantifying cell-generated mechanical forces within living 
    embryonic tissues},
    journal= NM,
    publisher = NPG,
    volume = {11},
    pages = {183-189},
    year = {2014},
}

@phdthesis{Wallmeyer,
  author       = {Wallmeyer, Bernhard},
  title        = {Collective cell migration in embryogenesis as a wetting phenomenon governed by hydrodynamics of active liquids},
  school      = {University of M\"unster},
  year         = {2019},
  month        = JUN
}

@article{Mott_2013,
	author = {Mott, P. H. and Roland, C. M.},
	title = {Limits to Poissons ratio in isotropic materialsgeneral result for arbitrary deformation},
	journal = {Physica Scripta},
	year = {2013},
	month = {apr},
	publisher = {IOP Publishing},
	volume = {87},
	number = {5},
	pages = {055404},
}   
@article{Mongera,
	author = {Mongera, A. and Rowghanian, P. and Gustafson, H. J. and Shelton, E. and Kealhofer, D. A. and Carn, E. K. and Serwane, F. and Lucio, A. A. and Giammona, J. and Camp\`{a}s, O.},
	title = {A fluid-to-solid jamming transition underlies vertebrate body axis elongation.},
	journal = NAT,
	year = {2018},
	month = {sep},
	publisher = {Springer},
	volume = {561},
	pages = {401-405},
}   
@article{Fiore,
	author = {Fiore, V. F. and Krajnc, M. and Quiroz, F. G. and others},
	title = {Mechanics of a multilayer epithelium instruct tumour architecture and function},
	journal = NAT,
	year = {2020},
	month = {sep},
	publisher = {Springer},
	volume = {585},
	pages = {433-439},
}   	
@article{Wang,
   title={Spherical harmonics method for computing the image stress due to a spherical void},
   volume={126},
   ISSN={0022-5096},
   journal={Journal of the Mechanics and Physics of Solids},
   publisher={Elsevier BV},
   author={Yifan Wang and Xiaohan Zhang and Cai, Wei},
   year={2019},
   month={May},
   pages={151–167}
}
@article{Neuber,
   author={H. Neuber},
   title={Ein neuer {A}nsatz zur {L}\"osung r\"aumlicher {P}robleme der {E}lastizit\"atstheorie},
   volume={14},
   journal={Journal of Applied Mathematics and Mechanics},
   year={1934},
   pages={203-212}
}
@article{Rivlin,
   author={Rivlin, R. S.},
   title={Large elastic deformations of isotropic materials IV. further developments of the general theory},
   volume={241},
   journal={Philosophical Transactions of the Royal Society of London. Series A, Mathematical and Physical Sciences},
   year={1948},
   pages={379-397}
}
@article{Papko,
   author={P. F. Papkovich},
   title={Solution G\'en\'erale des \'equations differentielles fondamentales d'\'elasticit\'e exprim\'ee par trois fonctions harmoniques},
   volume={195},
   journal=CRdlAdS,
   year={1932},
   pages={513-515}
}
@Manual{ram2010,
    title = {Mathematica 8.0},
    author = {{Wolfram Research Inc.}},
    year = {2010},
    address = {http://www.wolfram.com},
    note = {checked on 10.09.2021}
  }

@article {Hofmeier,
article_type = {journal},
title = {Global and local tension measurements in biomimetic skeletal muscle tissues reveals early mechanical homeostasis},
author = {Hofemeier, Arne D and Limon, Tamara and Muenker, Till Moritz and Wallmeyer, Bernhard and Jurado, Alejandro and Afshar, Mohammad Ebrahim and Ebrahimi, Majid and Tsukanov, Roman and Oleksiievets, Nazar and Enderlein, Jörg and Gilbert, Penney M and Betz, Timo},
editor = {Walczak, Aleksandra M and Bassereau, Patricia and Bassereau, Patricia and Chang, Natasha},
volume = 10,
year = 2021,
month = {jan},
pub_date = {2021-01-18},
pages = {e60145},
citation = {eLife 2021;10:e60145},
doi = {10.7554/eLife.60145},
url = {https://doi.org/10.7554/eLife.60145},
journal = {eLife},
issn = {2050-084X},
publisher = {eLife Sciences Publications, Ltd},
}

@Manual{Mat_defSh,
    title = {SphericalHarmonicY-Wolfram},
    author = {{Wolfram Research Inc.}},
    year = {2021},
    address = {https://reference.wolfram.com/language/ref/SphericalHarmonicY.html},
    note = {checked on 11.09.2021}
  }  
  
@Manual{SHReals,
    title = {Definition Real Spherical harmonics},
    author = {{SHTools}},
    year = {2021},
    address = {https://shtools.github.io/SHTOOLS/real-spherical-harmonics.html},
    note = {checked on 10.09.2021}
}

@Manual{SHExpand,
    title = {SHEpandLSQ()},
    author = {SHTools},
    year = {2021},
    address = {https://shtools.oca.eu/shtools/public/pyshexpandlsq.html},
    note = {checked on 10.09.2021}    
  }  

@ARTICLE{WittkowskiSS2009,
    author    = {Wittkowski, R. and Schelin, A.~B. and Spatschek, K.~H.},
    title     = {Mean motion in stochastic plasmas with a space-dependent diffusion coefficient},
    journal   = CtPP,
    publisher = WILEYVCH,
    address   = WEI,
    number    = {1-2},
    volume    = {49},
    pages     = {55-69},
    month     = MAR,
    year      = {2009}
}
%doi          = {10.1002/ctpp.200910009}

@BOOK{Abdullaev2006,
  title        = {Construction of Mappings for Hamiltonian Systems and Their Applications},
  publisher    = SPR,
  year         = {2006},
  author       = {S.~S. Abdullaev},
  volume       = {691},
  month        = APR,
  series       = {Lecture Notes in Physics},
  address      = BER,
  pages        = {379},
  isbn         = {3-540-30915-2}
}

@book{HairerLW2006,
    author     = {Hairer, Ernst and Lubich, Christian and Wanner, Gerhard},
    title      = {Geometric Numerical Integration: Structure-Preserving Algorithms for Ordinary Differential Equations},
    series     = CM,
    publisher  = SPR,
    address    = BER,
    month      = FEB,
    year       = {2006},
    pages      = {644},
    isbn       = {3-540-30663-3}
}

@book{Kroener1997,
    author     = {Kr{\"o}ner, Dietmar},
    title      = {Numerical Schemes for Conservation Laws},
    publisher  = WT,
    address    = {Chichester \& Stuttgart},
    year       = {1997},
    pages      = {508},
    isbn       = {3-519-02720-8}
}

@BOOK{Burgers1974,
   author    = {Burgers, J.~M.},
   title     = {The nonlinear diffusion equation},
   publisher = {Reidel},
   address   = DOR,
   year      = {1974}
}
 
@BOOK{Hackbusch1993,
   author    = {W. Hackbusch},
   title     = {Iterative L{\"o}sung gro\ss{}er schwachbesetzter Gleichungssysteme},
   publisher = TEU,
   address   = STU,
   pages     = {404},
   year      = {1993},
   isbn      = {3-519-12372-X}
}

@BOOK{Schwarz1993,
   author    = {Schwarz, H.~R.},
   title     = {Numerische Mathematik},
   publisher = TEU,
   address   = STU,
   year      = {1993}
}

@article{BevanP2000,
   author       = {Michael~A. Bevan and Dennis~C. Prieve},
   title        = {Hindered diffusion of colloidal particles very near to a wall: Revisited},
   publisher    = AIoP,
   year         = {2000},
   journal      = JoChemP,
   volume       = {113},
   number       = {3},
   pages        = {1228-1236}
}

@ARTICLE{Smoluchowski1906,
  author       = {von~Smoluchowski, M.},
  title        = {Zur kinetischen {T}heorie der {B}rownschen {M}olekularbewegung und der {S}uspensionen},
  journal      = AdPD,
  publisher    = WILEYVCH,
  year         = {1906},
  volume       = {326},
  number       = {14},
  pages        = {756-780},
  month        = NOV
}
% doi = {10.1002/andp.19063261405} address = WEI
% url = {http://adsabs.harvard.edu/abs/1906AnP...326..756V}

@article{Smoluchowski1916,
   author       = {von~Smoluchowski, M.},
   title        = {{\"U}ber {B}rownsche {M}olekularbewegung unter {E}inwirkung {\"a}u{\ss}erer {K}r{\"a}fte und deren {Z}usammenhang mit der verallgemeinerten {D}iffusionsgleichung},
   journal      = AdPD,
   volume       = {353},
   number       = {24},
   pages        = {1103-1112},
   year         = {1916},
   publisher    = WILEYVCH,
   month        = JAN
}
% doi = {10.1002/andp.19163532408} 
% address      = WEI
% url = {http://adsabs.harvard.edu/abs/1915AnP...353.1103S} 

@ARTICLE{Smoluchowski1916a,
  author       = {von~Smoluchowski, M.},
  title        = {{D}rei {V}ortr{\"a}ge {\"u}ber {D}iffusion, {B}rownsche {B}ewegung und {K}oagulation von {K}olloidteilchen},
  journal      = ZfP,
  publisher    = SPR,
  year         = {1916},
  volume       = {17},
  number       = {???},
  pages        = {557-585},
  month        = {???}
}
% doi = {}
% url = {http://adsabs.harvard.edu/abs/1916ZPhy...17..557S}

@ARTICLE{Einstein1905c,
  author       = {Einstein, A.},
  title        = {{\"U}ber die von der molekularkinetischen {T}heorie der {W}{\"a}rme geforderte {B}ewegung von in ruhenden {F}l{\"u}ssigkeiten suspendierten {T}eilchen},
  journal      = AdPD,
  publisher    = WILEYVCH,
  year         = {1905},
  volume       = {322},
  number       = {8},
  pages        = {549-560},
  month        = JUN
}
% doi = {10.1002/andp.19053220806}
% url = {http://adsabs.harvard.edu/abs/1905AnP...322..549E}

@ARTICLE{Einstein1906,
  author       = {Einstein, A.},
  title        = {{E}ine neue {B}estimmung der {M}olek{\"u}ldimensionen},
  journal      = AdPD,
  publisher    = WILEYVCH,
  year         = {1906},
  volume       = {324},
  number       = {2},
  pages        = {289-306},
  month        = FEB
}
% doi = {10.1002/andp.19063240204}
% url = {http://adsabs.harvard.edu/abs/1906AnP...324..289E}

@article{Einstein2005,
   author       = {Einstein, A.},
   title        = {{E}ine neue {B}estimmung der {M}olek{\"u}ldimensionen},
   journal      = AdPD,
   volume       = {14},
   number       = {1},
   year         = {2005},
   pages        = {229-247},
   publisher    = WILEYVCH,
   address      = WEI
}

@ARTICLE{Einstein1906a,
  author       = {Einstein, A.},
  title        = {{Z}ur {T}heorie der {B}rownschen {B}ewegung},
  journal      = AdPD,
  publisher    = WILEYVCH,
  year         = {1906},
  volume       = {324},
  number       = {2},
  pages        = {371-381},
  month        = FEB
}
% doi = {10.1002/andp.19063240208}
% url = {http://adsabs.harvard.edu/abs/1906AnP...324..371E}

@BOOK{Stoecker2007, 
   editor       = {Horst St{\"o}cker},
   title        = {Taschenbuch der Physik: Formeln, Tabellen, {\"U}ber"-sich"-ten},
   address      = FaM,
   edition      = {5, korrigierte Auflage, Nachdruck},
   publisher    = HD,
   year         = {2007}
} 

@BOOK{Gerthsen2002,
   author       = {Gerthsen, Christian and Meschede, Dieter},
   title        = {Physik},
   series       = {Springer Lehrbuch},
   publisher    = SPR,
   address      = {Berlin, Heidelberg, New York},
   pages        = {1288},
   year         = {2002},
   edition      = {21},
   isbn         = {3-540-42024-X}
}

@BOOK{BrockhausNWuT2003,
   title        = {Naturwissenschaft und Technik}, 
   series       = {Der Brockhaus},
   publisher    = {F. A. Brockhaus \& Spektrum Akademischer Verlag},
   address      = {Mannheim \& Heidelberg},
   year         = {2003},
   author       = {Wei\ss{}, Joachim and Winkenbach, Marion and {et~al.}},  
   volume       = {1-3},
   pages        = {2259},
   isbn         = {3-7653-1065-4}
}   

@BOOK{WalzEtAl2003,
   title        = {Lexikon der Mathematik},
   author       = {Walz, Guido and {et~al.}},
   year         = {2003},
   publisher    = SAV,
   volume       = {1-6},
   pages        = {2674},
   address      = {Heidelberg},
   isbn         = {3-827-41159-9}
}

@article{OttinoW2004,
   author       = {Ottino, Julio~M. and Wiggins, Stephen},
   title        = {Applied physics: designing optimal micromixers},
   journal      = SCI,
   publisher    = AAftAoS,
   volume       = {305},
   number       = {5683},
   pages        = {485-486},
   year         = {2004}
}

@ARTICLE{LambR1998,
   author       = {{Jeroen~S.~W.} Lamb and {John~A.~G.} Roberts},
   title        = {Time-reversal symmetry in dynamical systems: a survey},
   journal      = PD,
   publisher    = ELS,
   volume       = {112},
   pages        = {1-39},
   year         = {1998}
}

@ARTICLE{DittrichKOS2000,
   author       = {T. Dittrich and R. Ketzmerick and M.-F. Otto and H. Schanz},
   title        = {Classical and quantum transport in deterministic Hamiltonian ratchets},
   journal      = AdPD,
   publisher    = WILEYVCH,
   address      = Berlin,
   volume       = {9},
   number       = {9-10},
   pages        = {755-763},
   year         = {2000}
}

@ARTICLE{FlachYZ2000,
   author       = {S. Flach and O. Yevtushenko and Y. Zolotaryuk},
   title        = {Directed Current due to Broken Time-Space Symmetry},
   journal      = PRL,
   publisher    = APS,
   volume       = {84},
   number       = {11},
   pages        = {2358-2361},
   month        = MAR,
   year         = {2000}
}

@BOOK{Roempp1999,
   author       = {R{\"o}mpp, Hermann and {et~al.}},
   editor       = {J{\"u}rgen Falbe and Manfred Regitz},
   title        = {R{\"o}mpp kompakt Basislexikon Chemie},
   year         = {1999},
   publisher    = GTV,
   volume       = {1-4},
   address      = {Stuttgart, New York},
   edition      = {10},
   pages        = {3038}
}

@BOOK{Beilstein1975,
   author       = {Beilstein, {Friedrich~Konrad} and {et~al.}},
   title        = {Beilsteins Handbuch der Organischen Chemie},
   editor       = {{Beilstein-Institut f{\"u}r Literatur der Organischen Chemie}},
   publisher    = SPR,
   address      = {Berlin, New York},
   edition      = {4},
   year         = {1975},
   isbn         = {0-387-12021-1}
}

@BOOK{Lide2008,
   title        = {CRC Handbook of Chemistry and Physics},
   edition      = {89},
   author       = {Lide, {David~P.} and {et~al.}},
   publisher    = CRCP,
   month        = JUN,
   year         = {2000}
}

@book{BronsteinSMM2006,
   author       = {Bronstein, {Ilja~N.} and Semendjajew, {Konstantin~A.} and Musiol, {Gerhard} and M{\"u}hlig, {Heiner}},
   title        = {Taschenbuch der Mathematik},
   edition      = {6, Nachdruck},
   publisher    = HD,
   address      = FaM,
   pages        = {1200},
   isbn         = {3-8171-2016-8},
   year         = {2006}
}

@BOOK{LangP2005,
   title        = {Mathematische Methoden in der Physik},
   author       = {Lang, Christian~B. and Pucker, Norbert},
   edition      = {2},
   publisher    = SAV,
   address      = {M{\"u}nchen},
   pages        = {713},
   isbn         = {3-8274-1558-6},
   year         = {2005},
   month        = AUG
}

@BOOK{GriebelDN1995,
   author       = {Griebel, Michael and Dornseifer, Thomas and Neunhoeffer, Tilman},
   title        = {Numerische Simulation in der Str{\"o}mungsmechanik: Eine praxisorientierte Einf{\"u}hrung},
   publisher    = VIE,
   series       = {Lehrbuch Scientific Computing},
   year         = {1995},
   address      = BrWi,
   pages        = {219},
   isbn         = {3-528-06761-6}
}

 @article{PeacemanR1955,
   title        = {The Numerical Solution of Parabolic and Elliptic Differential Equations},
   author       = {Peaceman, {D.~W.} and {Rachford, Jr.}, {H.~H.}},
   journal      = JotSfIaAM,
   volume       = {3},
   number       = {1},
   month        = MAR,
   pages        = {28-41},
   publisher    = SfIaAM, 
   year         = {1955}
}
%  url          = {http://www.jstor.org/stable/2098834}
%  issn         = {03684245}

@ARTICLE{CourantIR1952,
   author       = {Courant, Richard and Isaacson, Eugene and Rees, Mina},
   title        = {On the solution of nonlinear hyperbolic differential equations by finite differences},
   journal      = CoPaAM,
   volume       = {5},
   number       = {3},
   pages        = {243-255},
   year         = {1952},
   publisher    = WileyP
}
% doi          = {10.1002/cpa.3160050303},
% url          = {http://dx.doi.org/10.1002/cpa.3160050303}

@BOOK{Hackbusch1985,
   author       = {Hackbusch, Wolfgang},
   title        = {Multi-grid methods and applications},
   publisher    = SPR,
   year         = {1985},
   pages        = {377},
   edition      = {2},
   isbn         = {3-540-12761-5}
}

@article{CrankN1947,
   author       = {Crank, J. and Nicolson, P.},
   title        = {A practical method for numerical evaluation of solutions of partial differential equations of the heat-conduction type},
   journal      = {Proceedings of the Cambridge Philosophical Society},
   volume       = {43},
   pages        = {50-67},
   year         = {1947}
}
    
@article{Thom1933,
   author       = {Thom, A.},
   title        = {The flow past circular cylinders at low speeds},
   journal      = PotRSoLA:CPoaMaPC,
   publisher    = RS,
   year         = {1933},
   month        = SEP,
   volume       = {141},
   number       = {845},
   pages        = {651-669}
}

@article{SchmidtRvBD2008,
  title={Non-equilibrium sedimentation of colloids: confocal microscopy and {B}rownian dynamics simulations},
  author={Schmidt, Matthias and Royall, C Patrick and van Blaaderen, Alfons and Dzubiella, Joachim},
  journal=JoP:CM,
  volume={20},
  number={49},
  pages={494222},
  year={2008},
  publisher={IOP Publishing}
}

@BOOK{Wesseling1992,
   author       = {Wesseling, P.},
   title        = {An introduction to multigrid methods},
   publisher    = WILEYVCH,
   address      = NY,
   year         = {1992},
   pages        = {291},
   isbn         = {0-471-93083-0}
}

@article{WiseWL2009,
  title={An energy-stable and convergent finite-difference scheme for the phase field crystal equation},
  author={Wise, Steven M and Wang, Cheng and Lowengrub, John S},
  journal={SIAM Journal on Numerical Analysis},
  volume={47},
  number={3},
  pages={2269--2288},
  year={2009},
  publisher={SIAM}
}


@MISC{BinnigR1982,  
   author       = {Binnig, Gerd and Rohrer, Heinrich},
   title        = {Scanning tunneling microscope},
   howpublished = {United States Patent 4343993},
   month        = AUG,
   year         = {1982},
   note         = {International Business Machines Corporation; International Classification: G01N 2300}
}

@article{BinnigRGW1982,
   author       = {Binnig, G. and Rohrer, H. and Gerber, {\relax Ch.} and Weibel, E.},
   title        = {Tunneling through a controllable vacuum gap},
   publisher    = AIoP,
   year         = {1982},
   journal      = APL,
   volume       = {40},
   number       = {2},
   pages        = {178-180}
}   
%  url = {http://link.aip.org/link/?APL/40/178/1}
%  doi = {10.1063/1.92999}

@article{BinnigQG1986,
   title       = {Atomic Force Microscope},
   author      = {Binnig, G. and Quate, {C.~F.} and Gerber, {\relax Ch.}},
   journal     = PRL,
   volume      = {56},
   number      = {9},
   pages       = {930-933},
   year        = {1986},
   month       = MAR,
   publisher   = APS
}
%  doi = {10.1103/PhysRevLett.56.930}

@ARTICLE{Kolmogorov1954,
   author      = {Kolmogorov, A.~N.},
   title       = {Conservation of conditionally periodic motions for a small change in the Hamiltonian},
   journal     = DANSSSR,
   publisher   = USSRAoS,
   year        = {1954},
   volume      = {98},
   pages       = {527-530}
}

@ARTICLE{Moser1962,
   author      = {Moser, J.},
   title       = {On invariant curves of area-preserving mappings of an annulus},
   journal     = NdAdWiG,
   year        = {1962},
   volume      = {II. Mathematisch-physikalische Klasse},
   pages       = {1-20}
}

@ARTICLE{Arnold1963,
   author      = {Arnol'd, V.~I.},
   title       = {Proof of a theorem of A. N. Kolmogorov on the invariance of quasi-periodic motions under small perturbations of the Hamiltonian},
   journal     = RMS,
   publisher   = IoPP,
   year        = {1963},
   month       = JAN,
   volume      = {18},
   number      = {5},
   pages       = {9-36}
}
%  doi = {10.1070/RM1963v018n05ABEH004130}
%  url = {http://stacks.iop.org/0036-0279/18/9}

@BOOK{Tabor1989,
   author      = {Tabor, Michael},
   title       = {Chaos and Integrability in Nonlinear Dynamics: An Introduction},
   publisher   = WILEYVCH,
   year        = {1989},
   month       = JAN,
   address     = NY,
   edition     = {1},
   pages       = {384},
   isbn        = {0-471-82728-2}
}

@ARTICLE{Meiss1992,
  title        = {Symplectic maps, variational principles, and transport},
  author       = {Meiss, J.~D.},
  journal      = RoMP,
  publisher    = APS,
  volume       = {64},
  number       = {3},
  pages        = {795-848},
  year         = {1992},
  month        = JUL,
  publisher    = APS
}
% doi = {10.1103/RevModPhys.64.795}
% url = {http://link.aps.org/doi/10.1103/RevModPhys.64.795}

@ARTICLE{Noether1918,
  author       = {Noether, E.},
  title        = {Invariante Variationsprobleme},
  journal      = NdKGdWzG,
  volume       = {II. Mathematisch-physikalische Klasse},
  year         = {1918},
  month        = SEP,
  pages        = {235-257}
}
 
@BOOK{Rasband1997,
  author       = {Rasband, S.~Neil},
  title        = {Chaotic Dynamics of Nonlinear Systems},
  series       = {Wiley Professional Paperback Series},
  publisher    = WIL,
  year         = {1997},
  month        = FEB,
  address      = NY,
  pages        = {230},
  isbn         = {0-471-18434-9}
}

@INBOOK{Rasband1997ThePoincareMap,
  author       = {Rasband, S.~Neil},
  title        = {Chaotic Dynamics of Nonlinear Systems},
  chapter      = {5.3: The Poincar{\'e} Map},
  series       = {Wiley Professional Paperback Series},
  publisher    = WIL,
  year         = {1997},
  month        = FEB,
  address      = NY,
  pages        = {92-95},
  isbn         = {0-471-18434-9}
}
  
@BOOK{GoldstonR1998,
  author       = {Goldston, Robert~J. and Rutherford, Paul~H.},
  title        = {Plasmaphysik: Eine Ein"-f{\"u}h"-rung},
  publisher    = VIE,
  year         = {1998},
  edition      = {1},
  series       = {Lehrbuch Physik},
  address      = BrWi,
  pages        = {388},
  isbn         = {3-528-06884-1}
}

@ARTICLE{Taylor1969,
  author       = {Taylor, J.~B.},
  title        = {Investigation of Charged Particle Invariants},
  journal      = {UKAEA Culham Laboratory Progress Report},
  year         = {1969},
  volume       = {CLM-PR 12}
}

@BOOK{dAlembert1997,
  author       = {{d'Alembert}, J.},
  title        = {Abhandlung {\"u}ber Dynamik},
  publisher    = HD,
  year         = {1997},
  volume       = {106},
  edition      = {2},
  series       = {Ostwalds Klassiker},
  address      = FaM,
  pages        = {210},
  isbn         = {3-8171-3106-8}
}

@BOOK{Doppler2000,
  author       = {Doppler, {\relax Ch.}},
  title        = {Schriften aus der Fr{\"u}hzeit der Astrophysik},
  publisher    = HD,
  year         = {2000},
  volume       = {161},
  edition      = {2},
  series       = {Ostwalds Klassiker},
  address      = FaM,
  pages        = {194},
  isbn         = {3-8171-3161-7}
}

@BOOK{Friedmann2006,
  author       = {Friedmann, A.},
  title        = {Die Welt als Raum und Zeit},
  publisher    = HD,
  year         = {2006},
  volume       = {287},
  edition      = {3},
  series       = {Ostwalds Klassiker},
  address      = FaM,
  pages        = {233},
  isbn         = {3-8171-3416-8}
}

@BOOK{EinsteinvS1997,
  author       = {Einstein, A. and von~Smoluchowski, M.},
  title        = {Untersuchungen {\"u}ber die Theorie der Brownschen Bewegung - Abhandlung {\"u}ber die Brownsche Bewegung und verwandte Erscheinungen},
  publisher    = HD,
  year         = {1997},
  volume       = {199},
  edition      = {3},
  series       = {Ostwalds Klassiker},
  address      = FaM,
  pages        = {252},
  isbn         = {3-8171-3207-2}
}

@BOOK{CarnotMC2003,
  author       = {Carnot, S. and Mayer, R. and Clausius, R.},
  title        = {Betrachtungen {\"u}ber die bewegende Kraft des Feuers - {\"U}ber die bewegende Kraft der W{\"a}rme - Die Mechanik der W{\"a}rme},
  publisher    = HD,
  year         = {2003},
  volume       = {37},
  edition      = {1},
  series       = {Ostwalds Klassiker},
  address      = FaM,
  pages        = {227},
  isbn         = {3-8171-3411-3}
}

@BOOK{Boltzmann2000,
  author       = {Boltzmann, L.},
  title        = {Entropie und Wahrscheinlichkeit},
  publisher    = HD,
  year         = {2000},
  volume       = {286},
  edition      = {305},
  series       = {Ostwalds Klassiker},
  address      = FaM,
  pages        = {305},
  isbn         = {3-8171-3286-7}
}

@BOOK{Maxwell2008,
  author       = {Maxwell, J.~C.},
  title        = {{\"U}ber Faradays Kraftlinien - {\"U}ber physikalische Kraftlinien},
  publisher    = HD,
  year         = {2008},
  volume       = {69},
  edition      = {4},
  series       = {Ostwalds Klassiker},
  address      = FaM,
  pages        = {276},
  isbn         = {3-8171-3424-3}
}

@ARTICLE{ElderKHG2002,
  title     = {Modeling elasticity in crystal growth},
  author    = {Elder, K.~R. and Katakowski, Mark and Haataja, Mikko and Grant, Martin},
  journal   = PRL,
  volume    = {88},
  number    = {24},
  pages     = {245701},
  year      = {2002},
  month     = JUN,
  publisher = APS
}
% doi       = {10.1103/PhysRevLett.88.245701}

@ARTICLE{ElderG2004,
  title     = {Modeling elastic and plastic deformations in nonequilibrium processing using phase field crystals},
  author    = {Elder, K.~R. and Grant, Martin},
  journal   = PRE,
  volume    = {70},
  number    = {5},
  pages     = {051605},
  year      = {2004},
  month     = NOV,
  publisher = APS
}
% doi       = {10.1103/PhysRevE.70.051605}

@ARTICLE{Emmerich2009,
  author    = {Emmerich, H.},
  title     = {Phase-field modelling for metals and colloids and nucleation therein - an overview},
  journal   = JoP:CM,
  publisher = IoPP,
  volume    = {21},
  number    = {46},
  pages     = {464103},
  year      = {2009},
  month     = NOV
}
% doi       = {10.1088/0953-8984/21/46/464103}
% url       = {http://adsabs.harvard.edu/abs/2009JPCM...21T4103E}

@ARTICLE{YuLV2009,
  title     = {Phase-field modeling of anomalous spiral step growth on Si(001) surface},
  author    = {Yu, {Yan-Mei} and Liu, {Bang-Gui} and Voigt, Axel},
  journal   = PRB,
  volume    = {79},
  number    = {23},
  pages     = {235317},
  year      = {2009},
  month     = JUN,
  publisher = APS
}
% doi       = {10.1103/PhysRevB.79.235317}

@ARTICLE{JaatinenAEAN2009,
  title     = {Thermodynamics of bcc metals in phase-field-crystal models},
  author    = {Jaatinen, A. and Achim, C.~V. and Elder, K.~R. and Ala-Nissila, T.},
  journal   = PRE,
  volume    = {80},
  number    = {3},
  pages     = {031602},
  year      = {2009},
  month     = SEP,
  publisher = APS
}
% doi       = {10.1103/PhysRevE.80.031602}

@ARTICLE{TegzeGTPJANP2009,
  title     = {Diffusion-controlled anisotropic growth of stable and metastable crystal polymorphs in the phase-field crystal model},
  author    = {Tegze, G. and Gr{\'a}n{\'a}sy, L. and T{\'o}th, G.~I. and Podmaniczky, F. and Jaatinen, A. and {Ala-Nissila}, T. and Pusztai, T.},
  journal   = PRL,
  volume    = {103},
  number    = {3},
  pages     = {035702},
  year      = {2009},
  month     = JUL,
  publisher = APS
}
% doi       = {10.1103/PhysRevLett.103.035702}
% url       = {http://adsabs.harvard.edu/abs/2009PhRvL.103c5702T}

@ARTICLE{MellenthinKP2008,
  title     = {Phase-field crystal study of grain-boundary premelting},
  author    = {Mellenthin, Jesper and Karma, Alain and Plapp, Mathis},
  journal   = PRB,
  volume    = {78},
  number    = {18},
  pages     = {184110},
  year      = {2008},
  month     = NOV,
  publisher = APS
}
% doi       = {10.1103/PhysRevB.78.184110}

@ARTICLE{McKennaGV2009,
   author = {McKenna, I.~M. and Gururajan, M.~P. and Voorhees, P.~W.},
    title = {Phase field modeling of grain growth: effect of boundary thickness, triple junctions, misorientation, and anisotropy},
  journal = JoMS,
     year = 2009,
    month = JAN,
   volume = 44,
    pages = {2206-2217}
}
%     doi = {10.1007/s10853-008-3196-7},
%  adsurl = {http://adsabs.harvard.edu/abs/2009JMatS..44.2206M},
% adsnote = {Provided by the SAO/NASA Astrophysics Data System}

@ARTICLE{HuangE2008,
  title     = {Mesoscopic and microscopic modeling of island formation in strained film epitaxy},
  author    = {Huang, {Zhi-Feng} and Elder, K.~R.},
  journal   = PRL,
  volume    = {101},
  number    = {15},
  pages     = {158701},
  year      = {2008},
  month     = OCT,
  publisher = APS
}
% doi       = {10.1103/PhysRevLett.101.158701}

@ARTICLE{WuV2009,
  title     = {Stress-induced morphological instabilities at the nanoscale examined using the phase field crystal approach},
  author    = {Wu, {Kuo-An} and Voorhees, Peter~W.},
  journal   = PRB,
  volume    = {80},
  number    = {12},
  pages     = {125408},
  year      = {2009},
  month     = SEP,
  publisher = APS
}
% doi       = {10.1103/PhysRevB.80.125408}

@ARTICLE{Evans1979,
author      = {Evans, R.},
title       = {The nature of the liquid-vapour interface and other topics in the statistical mechanics of non-uniform, classical fluids},
  journal   = AiP,
  publisher = TF,
  year      = {1979},
  month     = MAR,
  volume    = {28},
  pages     = {143-200}
}
% doi = {10.1080/00018737900101365},
% adsurl = {http://adsabs.harvard.edu/abs/1979AdPhy..28..143E},

@ARTICLE{Singh1991,
  author    = {Singh, Yashwant},
  title     = {Density-functional theory of freezing and properties of the ordered phase},
  journal   = PRS,
  publisher = ELS,
  volume    = {207},
  number    = {6},
  pages     = {351-444},
  year      = {1991},
  month     = SEP
} 
% issn      = {0370-1573},
% doi       = {10.1016/0370-1573(91)90097-6},
% url       = {http://adsabs.harvard.edu/abs/1991PhR...207..351S}

@ARTICLE{Loewen1994a,
  author    = {L{\"o}wen, Hartmut},
  title     = {Melting, freezing and colloidal suspensions},
  journal   = PRS,
  publisher = ELS,
  volume    = {237},
  number    = {5},
  pages     = {249-324},
  year      = {1994},
  month     = FEB
} 
% issn      = {0370-1573},
% doi       = {10.1016/0370-1573(94)90017-5},
% url       = {http://adsabs.harvard.edu/abs/1994PhR...237..249L},

@ARTICLE{vanTeeffelenBVL2009,
  title     = {Derivation of the phase-field-crystal model for colloidal solidification},
  author    = {van~Teeffelen, Sven and Backofen, Rainer and Voigt, Axel and L{\"o}wen, Hartmut},
  journal   = PRE,
  volume    = {79},
  number    = {5},
  pages     = {051404},
  year      = {2009},
  month     = MAY,
  publisher = APS
} 
% doi       = {10.1103/PhysRevE.79.051404}

@ARTICLE{RamakrishnanY1979,
  title     = {First-principles order-parameter theory of freezing},
  author    = {Ramakrishnan, T.~V. and Yussouff, M.},
  journal   = PRB,
  volume    = {19},
  number    = {5},
  pages     = {2775-2794},
  year      = {1979},
  month     = MAR,
  publisher = APS
}
% doi       = {10.1103/PhysRevB.19.2775}

@article{FangS2001,
  title={Kinetics and thermodynamics of protein adsorption: a generalized molecular theoretical approach},
  author={Fang, Fang and Szleifer, Igal},
  journal=BJ,
  volume={80},
  number={6},
  pages={2568--2589},
  year={2001},
  publisher={Elsevier}
}


@ARTICLE{RosenfeldSLT1997,
  title     = {Fun\-da\-men\-tal-mea\-sure free-energy density functional for hard spheres: dimensional crossover and freezing},
  author    = {Rosenfeld, Y. and Schmidt, M. and L{\"o}wen, H. and Tarazona, P.},
  journal   = PRE,
  volume    = {55},
  number    = {4},
  pages     = {4245-4263},
  year      = {1997},
  month     = APR,
  publisher = APS
}
% doi       = {10.1103/PhysRevE.55.4245}

@ARTICLE{RothELK2002,
  author   = {Roth, R. and Evans, R. and Lang, A. and Kahl, G.},
  title    = {Fundamental measure theory for hard-sphere mixtures revisited: the {White} {Bear} version},
  journal  = JoP:CM,
  volume   = {14},
  number   = {46},
  pages    = {12063-12078},
  year     = {2002},
  month    = NOV
} 
% doi      = {10.1088/0953-8984/14/46/313}
% url      = {http://stacks.iop.org/0953-8984/14/i=46/a=313}

@ARTICLE{HansenGoosM2009,
  title     = {Fundamental measure theory for inhomogeneous fluids of nonspherical hard particles},
  author    = {{Hansen-Goos}, Hendrik and Mecke, Klaus},
  journal   = PRL,
  volume    = {102},
  number    = {1},
  pages     = {018302},
  year      = {2009},
  month     = JAN,
  publisher = APS
} 
% doi       = {10.1103/PhysRevLett.102.018302}

@ARTICLE{LoewenBW1989,
  author    = {L{\"o}wen, H. and Beier, T. and Wagner, H.},
  title     = {Van der {W}aals theory of surface melting},
  journal   = EPL,
  publisher = EPS,
  volume    = {9},
  number    = {8},
  pages     = {791-796},
  year      = {1989},
  month     = AUG
} 
% doi       = {10.1209/0295-5075/9/8/009}
% url       = {http://adsabs.harvard.edu/abs/1989EL......9..791L}

@ARTICLE{OhnesorgeLW1991,
  title     = {Density-functional theory of surface melting},
  author    = {Ohnesorge, R. and L{\"o}wen, H. and Wagner, H.},
  journal   = PRA,
  volume    = {43},
  number    = {6},
  pages     = {2870-2878},
  year      = {1991},
  month     = MAR,
  publisher = APS
} 
% doi       = {10.1103/PhysRevA.43.2870}

@ARTICLE{Lutsko2006,
  author    = {Lutsko, James~F.},
  title     = {First principles derivation of {G}inzburg-{L}andau free energy models for crystalline systems},
  journal   = PA,
  publisher = ELS,
  volume    = {366},
  number    = {1},
  pages     = {229-242},
  year      = {2006},
  month     = JUL
} 
% issn      = {0378-4371},
% doi       = {10.1016/j.physa.2005.10.044},
% url       = {http://adsabs.harvard.edu/abs/2006PhyA..366..229L}

@ARTICLE{ElderPBSG2007,
  title     = {Phase-field crystal modeling and classical density functional theory of freezing},
  author    = {Elder, K.~R. and Provatas, Nikolas and Berry, Joel and Stefanovic, Peter and Grant, Martin},
  journal   = PRB,
  volume    = {75},
  number    = {6},
  pages     = {064107},
  year      = {2007},
  month     = FEB,
  publisher = APS
} 
% doi       = {10.1103/PhysRevB.75.064107}

@ARTICLE{MarconiT1999,
  author    = {{Marini Bettolo Marconi}, Umberto and Tarazona, Pedro},
  title     = {Dynamic density functional theory of fluids},
  publisher = AIoP,
  year      = {1999},
  journal   = JoChemP,
  volume    = {110},
  number    = {16},
  pages     = {8032-8044},
  month     = APR 
}
% doi       = {10.1063/1.478705}
% url       = {http://adsabs.harvard.edu/abs/1999JChPh.110.8032M}

@ARTICLE{ArcherE2004,
  author    = {Archer, A.~J. and Evans, R.},
  title     = {Dynamical density functional theory and its application to spinodal decomposition},
  publisher = AIoP,
  year      = {2004},
  journal   = JoChemP,
  volume    = {121},
  number    = {9},
  pages     = {4246-4254},
  month     = SEP
}
% doi       = {10.1063/1.1778374}
% url       = {http://adsabs.harvard.edu/abs/2004JChPh.121.4246A}

@ARTICLE{EspanolL2009,
  author    = {Espa{\~n}ol, Pep and L{\"o}wen, Hartmut},
  title     = {Derivation of dynamical density functional theory using the projection operator technique},
  publisher = AIoP,
  year      = {2009},
  journal   = JoChemP,
  volume    = {131},
  number    = {24},
  pages     = {244101},
  month     = DEC
}
% eid       = {244101}
% doi       = {10.1063/1.3266943}
% url       = {http://adsabs.harvard.edu/abs/2009JChPh.131x4101E}

@ARTICLE{PrielerHLVHE2009,
  author    = {Prieler, R. and Hubert, J. and Li, D. and Verleye, D. and Haberkern, R. and Emmerich, H.},
  title     = {An anisotropic phase-field crystal model for heterogeneous nucleation of ellipsoidal colloids},
  journal   = JoP:CM,
  volume    = {21},
  number    = {46},
  pages     = {1-7},
  year      = {2009}
}
% url       = {http://stacks.iop.org/0953-8984/21/i=46/a=464110}

@ARTICLE{BolhuisF1997,
  author    = {Bolhuis, Peter and Frenkel, Daan},
  title     = {Tracing the phase boundaries of hard spherocylinders},
  publisher = AIoP,
  year      = {1997},
  journal   = JoChemP,
  volume    = {106},
  number    = {2},
  pages     = {666-687},
  month     = JAN
}
% doi       = {10.1063/1.473404}
% url       = {http://link.aip.org/link/?JCP/106/666/1},

@ARTICLE{RexWL2007,
  title     = {Dynamical density functional theory for anisotropic colloidal particles},
  author    = {Rex, M. and Wensink, H.~H. and L{\"o}wen, H.},
  journal   = PRE,
  volume    = {76},
  number    = {2},
  pages     = {021403},
  year      = {2007},
  month     = AUG,
  publisher = APS
} 
% doi       = {10.1103/PhysRevE.76.021403}

@ARTICLE{WensinkL2008,
  title     = {Aggregation of self-propelled colloidal rods near confining walls},
  author    = {Wensink, H.~H. and L{\"o}wen, H.},
  journal   = PRE,
  volume    = {78},
  number    = {3},
  pages     = {031409},
  year      = {2008},
  month     = SEP,
  publisher = APS
}
% doi       = {10.1103/PhysRevE.78.031409}

@ARTICLE{Loewen1994b,
  title     = {Brownian dynamics of hard spherocylinders},
  author    = {L{\"o}wen, Hartmut},
  journal   = PRE,
  volume    = {50},
  number    = {2},
  pages     = {1232-1242},
  year      = {1994},
  month     = AUG,
  publisher = APS
}
% doi       = {10.1103/PhysRevE.50.1232}

@ARTICLE{FrenkelMMT1984,
  title     = {Phase diagram of a system of hard ellipsoids},
  author    = {Frenkel, D. and Mulder, B.~M. and McTague, J.~P.},
  journal   = PRL,
  volume    = {52},
  number    = {4},
  pages     = {287-290},
  year      = {1984},
  month     = JAN,
  publisher = APS
}
% doi       = {10.1103/PhysRevLett.52.287}

@ARTICLE{Loewen1994c,
  title     = {Interaction between charged rodlike colloidal particles},
  author    = {L{\"o}wen, Hartmut},
  journal   = PRL,
  volume    = {72},
  number    = {3},
  pages     = {424-427},
  year      = {1994},
  month     = JAN,
  publisher = APS
}
% doi       = {10.1103/PhysRevLett.72.424}

@ARTICLE{Loewen1994d,
   author = {L{\"o}wen, H.},
    title = {Charged rodlike colloidal suspensions: an ab initio approach},
  journal = JoChemP,
     year = {1994},
    month = MAY,
   volume = {100},
    pages = {6738-6749}
}
%     doi = {10.1063/1.467034},
%  adsurl = {http://adsabs.harvard.edu/abs/1994JChPh.100.6738L},

@ARTICLE{KirchhoffLK1996,
  title     = {Dynamical correlations in suspensions of charged rodlike macromolecules},
  author    = {Kirchhoff, {\relax Th.} and L{\"o}wen, H. and Klein, R.},
  journal   = PRE,
  volume    = {53},
  number    = {5},
  pages     = {5011-5022},
  year      = {1996},
  month     = MAY,
  publisher = APS
} 
% doi       = {10.1103/PhysRevE.53.5011}

@ARTICLE{CleaverCAN1996,
  title     = {Extension and generalization of the Gay-Berne potential},
  author    = {Cleaver, Douglas~J. and Care, Christopher~M. and Allen, Michael~P. and Neal, Maureen~P.},
  journal   = PRE,
  volume    = {54},
  number    = {1},
  pages     = {559-567},
  year      = {1996},
  month     = JUL,
  publisher = APS
} 
% doi       = {10.1103/PhysRevE.54.559}

@book{KimK2013,
  title={Microhydrodynamics: Principles and Selected Applications},
  author={Kim, S. and Karrila, S.J.},
  series={Dover Civil and Mechanical Engineering},
  year={2013},
  publisher={Dover Publications}
}

@ARTICLE{FukunagaTD2004,
   author   = {Fukunaga, H. and Takimoto, J.-I. and Doi, M.},
    title   = {Molecular dynamics simulation study on the phase behavior of the Gay-Berne model with a terminal dipole and a flexible tail},
  journal   = JoChemP,
     year   = {2004},
    month   = APR,
   volume   = {120},
    pages   = {7792-7800}
} 
%     doi   = {10.1063/1.1687681},
%  adsurl   = {http://adsabs.harvard.edu/abs/2004JChPh.120.7792F},

@ARTICLE{MuccioliZ2006,
   author = {Muccioli, L. and Zannoni, C.},
    title = {A deformable {G}ay {B}erne model for the simulation of liquid crystals and soft materials},
  journal = CPL,
     year = {2006},
    month = MAY,
   volume = {423},
    pages = {1-6}
} 
%     doi = {10.1016/j.cplett.2006.03.022},
%  adsurl = {http://adsabs.harvard.edu/abs/2006CPL...423....1M},

@ARTICLE{PoniewierskiH1988,
  title     = {Density-functional theory for nematic and smectic-A ordering of hard spherocylinders},
  author    = {Poniewierski, A. and Ho\l{}yst, R.},
  journal   = PRL,
  volume    = {61},
  number    = {21},
  pages     = {2461-2464},
  numpages  = {3},
  year      = {1988},
  month     = NOV,
  publisher = APS
} 
% doi       = {10.1103/PhysRevLett.61.2461}

@ARTICLE{GrafL1999,
  author    = {Graf, Hartmut and L{\"o}wen, Hartmut},
  title     = {Density functional theory for hard spherocylinders: phase transitions in the bulk and in the presence of external fields},
  journal   = JoP:CM,
  publisher = IoPP,
  volume    = {11},
  number    = {6},
  pages     = {1435-1452},
  year      = {1999},
  month     = FEB
}
% doi       = {10.1088/0953-8984/11/6/008}
% url       = {http://stacks.iop.org/0953-8984/11/i=6/a=008}

@ARTICLE{MukherjeePB2001,
   author = {Mukherjee, P.~K. and Pleiner, H. and Brand, H.~R.},
    title = {Simple Landau model of the smectic-A-isotropic phase transition},
  journal = EPJE,
publisher = EDPS,
     year = {2001},
    month = MAR,
   volume = {4},
    pages = {293-297}
}
%     doi = {10.1007/s101890170111},
%  adsurl = {http://adsabs.harvard.edu/abs/2001EPJE....4..293M},

@BOOK{deGennesP1995,
  author    = {de~Gennes, P.-G. and Prost, J.},
  title     = {The Physics of Liquid Crystals},
  series    = {International Series of Monographs on Physics},
  volume    = {83},
  edition   = {2},
  publisher = OUP,
  address   = OX,
  month     = APR,
  year      = {1995},
  pages     = {597},
  isbn      = {0-198-51785-8}
} 
% ean       = {9780198517856}

@ARTICLE{TasinkevychSPTdG2002,
   author = {Tasinkevych, M. and Silvestre, N.~M. and Patr{\'i}cio, P. and {Telo da Gama}, M.~M.},
    title = {Colloidal interactions in two-dimensional nematics},
  journal = EPJE,
publisher = EDPS,
     year = {2002},
    month = NOV,
   volume = {9},
   number = {4},
    pages = {341-347}
}
%     doi = {10.1140/epje/i2002-10087-y},
%  adsurl = {http://adsabs.harvard.edu/abs/2002EPJE....9..341T},

@ARTICLE{BrandP1980,
  author    = {Brand, H.~R. and Pleiner, H.},
  title     = {Nonlinear reversible hydrodynamics of liquid crystals and crystals},
  journal   = JdP,
  year      = {1980},
  volume    = {41},
  number    = {6},
  pages     = {553-564},
} 
% doi       = {10.1051/jphys:01980004106055300},
% url       = {http://dx.doi.org/10.1051/jphys:01980004106055300}

@ARTICLE{BrandP1987,
  title     = {Nonlinear effects in the electrohydrodynamics of uniaxial nematic liquid crystals},
  author    = {Brand, Helmut~R. and Pleiner, Harald},
  journal   = PRA,
  volume    = {35},
  number    = {7},
  pages     = {3122-3127},
  year      = {1987},
  month     = APR,
  publisher = APS
}
% doi       = {10.1103/PhysRevA.35.3122}

@ARTICLE{vanRoijBMF1995,
  title     = {Transverse interlayer order in lyotropic smectic liquid crystals},
  author    = {van~Roij, Ren{\'e} and Bolhuis, Peter and Mulder, Bela and Frenkel, Daan},
  journal   = PRE,
  volume    = {52},
  number    = {2},
  pages     = {R1277-R1280},
  numpages  = {3},
  year      = {1995},
  month     = AUG,
  publisher = APS
} 
% doi       = {10.1103/PhysRevE.52.R1277}

@ARTICLE{LiuM1997,
  author    = {Liu, C. and Muthukumar, M.},
  title     = {Annihilation kinetics of liquid crystal defects},
  publisher = AIoP,
  year      = {1997},
  journal   = JoChemP,
  volume    = {106},
  number    = {18},
  pages     = {7822-7828},
  month     = MAY
} 
% doi       = {10.1063/1.473740}
% url       = {http://link.aip.org/link/?JCP/106/7822/1},

@ARTICLE{BechhoeferLT1991,
  title     = {Dynamical mechanism for the formation of metastable phases},
  author    = {Bechhoefer, John and L{\"o}wen, Hartmut and Tuckerman, Laurette~S.},
  journal   = PRL,
  volume    = {67},
  number    = {10},
  pages     = {1266-1269},
  year      = {1991},
  month     = SEP,
  publisher = APS
} 
% doi       = {10.1103/PhysRevLett.67.1266}

@ARTICLE{Loewen2010,
  author       = {L{\"o}wen, Hartmut},
  title        = {A phase-field-crystal model for liquid crystals},
  journal      = JoP:CM,
  publisher    = IoP,
  year         = {2010},
  volume       = {22},
  number       = {36},
  pages        = {364105},
  month        = MAR
}
% doi = { 10.1088/0953-8984/22/36/364105}
% url = {http://stacks.iop.org/0953-8984/22/i=36/a=364105}

@article{DonevFVE2014,
  title={A reversible mesoscopic model of diffusion in liquids: from giant fluctuations to {F}ick’s law},
  author={Donev, Aleksandar and Fai, Thomas G and Vanden-Eijnden, Eric},
  journal={Journal of Statistical Mechanics: Theory and Experiment},
  volume={2014},
  number={4},
  pages={P04004},
  year={2014},
  publisher={IOP Publishing}
}

@book{Zeh1989,
    title={The physical basis of the direction of time},
    author ={Zeh, Hans-Dieter},
    year={1989},
    publisher={Springer Verlag},
    address ={Berlin Heidelberg}
}

@article{AlandLV2011,
  title={A continuum model of colloid-stabilized interfaces},
  author={Aland, Sebastian and Lowengrub, John and Voigt, Axel},
  journal=PoF,
  volume={23},
  number={6},
  pages={062103},
  year={2011},
  publisher={AIP}
}

@ARTICLE{ParrishEK1999,
  author       = {{Parrish}, J.~K. and {Edelstein-Keshet}, L.},
  title        = {Complexity, pattern, and evolutionary trade-offs in animal aggregation},
  journal      = SCI,
  publisher    = AAftAoS,
  year         = {1999},
  volume       = {284},
  number       = {5411},
  pages        = {99-101},
  month        = APR
}
% doi = {10.1126/science.284.5411.99}
% url = {http://adsabs.harvard.edu/abs/1999Sci...284...99P}

@ARTICLE{KruseJJPS2005,
  author       = {{Kruse}, K. and {Joanny}, J.~F. and {J{\"u}licher}, F. and {Prost}, J. and {Sekimoto}, K.},
  title        = {Generic theory of active polar gels: a paradigm for cytoskeletal dynamics},
  journal      = EPJE,
  publisher    = EDPS,
  year         = {2005},
  volume       = {16},
  number       = {1},
  pages        = {5-16},
  month        = JAN
}
% doi = {10.1140/epje/e2005-00002-5}
% url = {http://adsabs.harvard.edu/abs/2005EPJE...16....5K}

@ARTICLE{GiomiML2008,
  author       = {{Giomi}, L. and {Marchetti}, M.~C. and {Liverpool}, T.~B.},
  title        = {Complex spontaneous flows and concentration banding in active polar films},
  journal      = PRL,
  publisher    = APS,
  year         = {2008},
  volume       = {101},
  number       = {19},
  pages        = {198101},
  month        = NOV
}
% doi = {10.1103/PhysRevLett.101.198101}
% url = {http://adsabs.harvard.edu/abs/2008PhRvL.101s8101G}

@ARTICLE{BrandPS2011,
  author       = {Brand, H. and Pleiner, H. and Sven\v{s}ek, D.},
  title        = {Macroscopic behavior of systems with an axial dynamic preferred direction},
  journal      = EPJE,
  publisher    = EDPS,
  year         = {2011},
  volume       = {34},
  number       = {11},
  pages        = {1-10},
  month        = NOV
}
% doi = {10.1140/epje/i2011-11128-2}
% url = {http://dx.doi.org/10.1140/epje/i2011-11128-2}

@ARTICLE{WittkowskiLB2010,
  author       = {Wittkowski, Raphael and L{\"o}wen, Hartmut and Brand, Helmut~R.},
  title        = {Derivation of a three-dimensional phase-field-crystal model for liquid crystals from density functional theory},
  journal      = PRE,
  publisher    = APS,
  year         = {2010},
  volume       = {82},
  number       = {3},
  pages        = {031708},
  month        = SEP
}
% doi = {10.1103/PhysRevE.82.031708}
% url = {http://pre.aps.org/abstract/PRE/v82/i3/e031708}

@article{ChandraB1990,
  title={Collective orientational relaxation in a dense liquid of ellipsoidal molecules},
  author={Chandra, Amalendu and Bagchi, Biman},
  journal={Physica A: Statistical Mechanics and its Applications},
  volume={169},
  number={2},
  pages={246--262},
  year={1990},
  publisher={Elsevier}
}

@article{ChandraB1989,
  title={A molecular theory of collective orientational relaxation in pure and binary dipolar liquids},
  author={Chandra, Amalendu and Bagchi, Biman},
  journal=JoChemP,
  volume={91},
  number={3},
  pages={1829--1842},
  year={1989},
  publisher={AIP}
}

@BOOK{Chandrasekhar1992,
  author       = {Chandrasekhar, S.},
  title        = {Liquid crystals},
  publisher    = CUP,
  year         = {1992},
  month        = NOV,
  address      = CAM,
  edition      = {2},
  pages        = {480},
  isbn         = {0-521-41747-3}
}

@article{ZhongCD2015,
  title={Sessile nanofluid droplet drying},
  author={Zhong, Xin and Crivoi, Alexandru and Duan, Fei},
  journal={Advances in Colloid and Interface Science},
  volume={217},
  pages={13--30},
  year={2015},
  publisher={Elsevier}
}

@article{HaatajaGL2010,
  title={Classical density functional theory methods in soft and hard matter},
  author={Haataja, Mikko and Gr{\'a}n{\'a}sy, L{\'a}szl{\'o} and L{\"o}wen, Hartmut},
  journal=JoP:CM,
  volume={22},
  number={36},
  pages={360301},
  year={2010},
  publisher={IOP Publishing}
}

@article{Tozzini2009,
  title={Multiscale modeling of proteins},
  author={Tozzini, Valentina},
  journal={Accounts of Chemical Research},
  volume={43},
  number={2},
  pages={220--230},
  year={2009},
  publisher={ACS Publications}
}

@ARTICLE{Wetter1985,
  author       = {Wetter, Carl},
  title        = {Die Fl{\"u}ssigkristalle des Tabakmosaikvirus},
  journal      = BiuZ,
  publisher    = WIL,
  year         = {1985},
  volume       = {15},
  number       = {3},
  pages        = {81-89},
  month        = JUN
}
% doi = {10.1002/biuz.19850150307}
% url = {http://dx.doi.org/10.1002/biuz.19850150307}

@ARTICLE{VollrathK2001,
  author       = {Vollrath, Fritz and Knight, David~P.},
  title        = {Liquid crystalline spinning of spider silk},
  journal      = NAT,
  publisher    = NPG,
  year         = {2001},
  volume       = {410},
  number       = {6828},
  pages        = {541-548},
  month        = MAR
}
% doi = {10.1038/35069000}
% url = {http://dx.doi.org/10.1038/35069000}

@MISC{Luk1981,
   author       = {Luk, Catherine~M.},
   title        = {Liquid crystal temperature indicator for measuring human body temperature},
   howpublished = {United States Patent 4302971},
   month        = DEC,
   year         = {1981}
}

@ARTICLE{PrasadKSJWK2005,
  author       = {Prasad, Veena and Kang, {Shin-Woong} and Suresh, K.~A. and Joshi, Leela and Wang, Qingbing and Kumar, Satyendra},
  title        = {Thermotropic Uniaxial and Biaxial Nematic and Smectic Phases in Bent-Core Mesogens},
  journal      = JotACS,
  publisher    = ACS,
  year         = {2005},
  volume       = {127},
  number       = {49},
  pages        = {17224-17227},
  month        = APR
}
% doi = {10.1021/ja052769n}
% url = {http://pubs.acs.org/doi/pdf/10.1021/ja052769n}

@article{Nakajima1958,
  title={On quantum theory of transport phenomena: steady diffusion},
  author={Nakajima, Sadao},
  year={1958},
  journal=PoTP,
  volume={20},
  number={6},
  pages={948-959},
  publisher=OUP
}

@ARTICLE{MadsenDNS2004,
  author       = {Madsen, L.~A. and Dingemans, T.~J. and Nakata, M. and Samulski, E.~T.},
  title        = {Thermotropic Biaxial Nematic Liquid Crystals},
  journal      = PRL,
  publisher    = APS,
  year         = {2004},
  volume       = {92},
  number       = {14},
  pages        = {145505},
  month        = APR
}
% doi = {10.1103/PhysRevLett.92.145505}
% url = {https://link.aps.org/doi/10.1103/PhysRevLett.92.145505}

@article{Liu2016,
  title={Development of 3-dimensional time-dependent density functional theory and its application to gas diffusion in nanoporous materials},
  author={Liu, Yu},
  journal={Physical Chemistry Chemical Physics},
  volume={18},
  number={19},
  pages={13158--13163},
  year={2016},
  publisher={Royal Society of Chemistry}
}

@article{MarconiM2014,
  title={Kinetic Density Functional Theory: A microscopic approach to fluid mechanics},
  author={{Marini Bettolo Marconi}, Umberto and Melchionna, Simone},
  journal={Communications in Theoretical Physics},
  volume={62},
  number={4},
  pages={596},
  year={2014},
  publisher={IOP Publishing}
}

@article{NunesGAdG2018,
  title={Field-driven dynamical demixing of binary mixtures},
  author={Nunes, Andr{\'e} S and Gupta, Akshat and Ara{\'u}jo, Nuno AM and da Gama, Margarida M Telo},
  journal={Molecular Physics},
  volume={116},
  number={21-22},
  pages={3224--3230},
  year={2018},
  publisher={Taylor \& Francis}
}

@article{Klapp2016,
  title={Collective dynamics of dipolar and multipolar colloids: From passive to active systems},
  author={Klapp, Sabine HL},
  journal={Current Opinion in Colloid \& Interface Science},
  volume={21},
  pages={76--85},
  year={2016},
  publisher={Elsevier}
}

@article{LiebchenL2018,
  title={Modelling chemotaxis of microswimmers: from individual to collective behavior},
  author={Liebchen, B and L{\"o}wen, H},
  journal={preprint},
  volume={arXiv:1802.07933},
  year={2018}
}

@article{GrawitterS2018,
  title={Feedback control of photoresponsive fluid interfaces},
  author={Grawitter, Josua and Stark, Holger},
  journal=SM,
  volume={14},
  number={10},
  pages={1856--1869},
  year={2018},
  publisher={The Royal Society of Chemistry}
}

@article{LiuL2018,
  title={Time-dependent density functional theory for fluid diffusion in graphene oxide membranes/graphene membranes},
  author={Liu, Yu and Liu, Honglai},
  journal={Chemical Engineering Science},
  volume={188},
  pages={150--157},
  year={2018},
  publisher={Elsevier}
}

@inproceedings{SanchezRM2013,
  title={Irreversible equilibration and aging in glass-forming liquids},
  author={S{\'a}nchez-D{\'\i}az, LE and Ram{\'\i}rez-Gonz{\'a}lez, PE and Medina-Noyola, M},
  booktitle={AIP Conference Proceedings},
  volume={1518},
  number={1},
  pages={28--37},
  year={2013},
  organization={AIP}
}

@incollection{ArcherWTK2016,
  title={Generation of defects and disorder from deeply quenching a liquid to form a solid},
  author={Archer, Andrew J and Walters, Morgan C and Thiele, Uwe and Knobloch, Edgar},
  booktitle={Mathematical Challenges in a New Phase of Materials Science},
  pages={1--26},
  year={2016},
  publisher={Springer}
}

@incollection{GernertLLK2016,
  title={Feedback control of colloidal transport},
  author={Gernert, Robert and Loos, Sarah AM and Lichtner, Ken and Klapp, Sabine HL},
  booktitle={Control of Self-Organizing Nonlinear Systems},
  pages={375--392},
  year={2016},
  publisher={Springer}
}

@article{LoosGK2019,
  title={Non-reciprocal hidden degrees of freedom: a unifying perspective on memory, feedback, and activity},
  author={Loos, Sarah A M and Hermann, Simon M and Klapp, Sabine HL},
  journal={arXiv:1910.08372},
  year={2019}
}

@article{InoueY2014,
  title={Effects of interactions between particles on dynamics in microrheology},
  author={Inoue, Masao and Yoshimori, Akira},
  journal={Journal of Molecular Liquids},
  volume={200},
  pages={81--84},
  year={2014},
  publisher={Elsevier}
}

@article{Emmerich2011,
  title={Phase-field models for colloidal systems: First steps, perspectives and open challenges},
  author={Emmerich, H},
  journal={Current Opinion in Solid State and Materials Science},
  volume={15},
  number={3},
  pages={83--86},
  year={2011},
  publisher={Elsevier}
}

@article{OkamotoO2016,
  title={Density functional theory of gas--liquid phase separation in dilute binary mixtures},
  author={Okamoto, Ryuichi and Onuki, Akira},
  journal=JoP:CM,
  volume={28},
  number={24},
  pages={244012},
  year={2016},
  publisher={IOP Publishing}
}

@article{LiQK2018,
  title={Direct discretization method for the {C}ahn--{H}illiard equation on an evolving surface},
  author={Li, Yibao and Qi, Xuelin and Kim, Junseok},
  journal={Journal of Scientific Computing},
  volume={77},
  number={2},
  pages={1147--1163},
  year={2018},
  publisher={Springer}
}

@article{Yoshimori2007,
  title={A molecular theory of large-solute diffusion},
  author={Yoshimori, A},
  journal={Condensed Matter Physics},
  year={2007},
  publisher={Інститут фізики конденсованих систем НАН України}
}

@article{StuhlmullerEdlHS2018,
  title={Structural nonequilibrium forces in driven colloidal systems},
  author={Stuhlm{\"u}ller, Nico CX and Eckert, Tobias and {de las Heras}, Daniel and Schmidt, Matthias},
  journal=PRL,
  volume={121},
  number={9},
  pages={098002},
  year={2018},
  publisher={APS}
}

@article{LiNZG2018,
  title={Phase field crystal modeling of grain boundary structures and growth in polycrystalline graphene},
  author={Li, Jiaoyan and Ni, Bo and Zhang, Teng and Gao, Huajian},
  journal={Journal of the Mechanics and Physics of Solids},
  volume={120},
  pages={36--48},
  year={2018},
  publisher={Elsevier}
}

@article{QiS2017b,
  title={Hybrid particle-continuum simulations coupling {B}rownian dynamics and local dynamic density functional theory},
  author={Qi, Shuanhu and Schmid, Friederike},
  journal=SM,
  volume={13},
  number={43},
  pages={7938--7947},
  year={2017},
  publisher={Royal Society of Chemistry}
}

@article{NunesAT2016,
  title={Self-assembly of colloidal bands driven by a periodic external field},
  author={Nunes, Andr{\'e} S and Ara{\'u}jo, Nuno AM and Telo da Gama, Margarida M},
  journal=JoChemP,
  volume={144},
  number={3},
  pages={034902},
  year={2016},
  publisher={AIP Publishing}
}

@article{YeDTZM2017,
  title={Diffusive dynamics of polymer chains in an array of nanoposts},
  author={Ye, Yi and Du, Zhongjie and Tian, Ming and Zhang, Liqun and Mi, Jianguo},
  journal={Physical Chemistry Chemical Physics},
  volume={19},
  number={1},
  pages={380--387},
  year={2017},
  publisher={Royal Society of Chemistry}
}

@article{KikkinidesM2015,
  title={Dynamic density functional theory with hydrodynamic interactions: Theoretical development and application in the study of phase separation in gas-liquid systems},
  author={Kikkinides, ES and Monson, PA},
  journal=JoChemP,
  volume={142},
  number={9},
  pages={094706},
  year={2015},
  publisher={AIP Publishing}
}

@article{HughesTA2015,
  title={Liquid drops on a surface: Using density functional theory to calculate the binding potential and drop profiles and comparing with results from mesoscopic modelling},
  author={Hughes, Adam P and Thiele, Uwe and Archer, Andrew J},
  journal=JoChemP,
  volume={142},
  number={7},
  pages={074702},
  year={2015},
  publisher={AIP Publishing}
}

@article{JouCL1999,
  title={Extended irreversible thermodynamics revisited (1988-98)},
  author={Jou, David and Casas-V\'{a}zquez, Jose and Lebon, Georgy},
  journal={Reports on Progress in Physics},
  volume={62},
  number={7},
  pages={1035},
  year={1999},
  publisher={IOP Publishing}
}

@article{HughesBBRB2012,
  title={Extended hydrodynamic approach to quantum-classical nonequilibrium evolution. II. Application to nonpolar solvation},
  author={Hughes, Keith H and Baxter, Sean N and Bousquet, David and Ramanathan, Padmanaban and Burghardt, Irene},
  journal=JoChemP,
  volume={136},
  number={1},
  pages={014102},
  year={2012},
  publisher={AIP}
}

@article{BierA2013,
  title={Nonequilibrium interfaces in colloidal fluids},
  author={Bier, Markus and Arnold, Daniel},
  journal=PRE,
  volume={88},
  number={6},
  pages={062307},
  year={2013},
  publisher={APS}
}

@article{HoellLM2018,
  title={Particle-scale statistical theory for hydrodynamically induced polar ordering in microswimmer suspensions},
  author={Hoell, Christian and L{\"o}wen, Hartmut and Menzel, Andreas M},
  journal=JoChemP,
  volume={149},
  number={14},
  pages={144902},
  year={2018},
  publisher={AIP Publishing}
}

@inproceedings{diZS2019,
  title={A massively parallel infrastructure for adaptive multiscale simulations: modeling {RAS} initiation pathway for cancer},
  author={Di Natale, Francesco and Bhatia, Harsh and Carpenter, Timothy S and Neale, Chris and Schumacher, Sara Kokkila and Oppelstrup, Tomas and Stanton, Liam and Zhang, Xiaohua and Sundram, Shiv and Scogland, Thomas R.~W. and others},
  booktitle={Supercomputing: The International Conference for High Performance Computing, Networking, Storage, and Analysis (To Appear). ACM},
  year={2019}
}

@article{CasselmanDM2015,
  title={Modelling the dynamics of condensation and evaporation of fluids in three-dimensional slit pores},
  author={Casselman, Joshua A and Desouza, Anish and Monson, Peter A},
  journal=MP,
  volume={113},
  number={9-10},
  pages={1250--1260},
  year={2015},
  publisher={Taylor \& Francis}
}

@article{JuniperZSBALD2017,
  title={Dynamic mode locking in a driven colloidal system: experiments and theory},
  author={Juniper, Michael PN and Zimmermann, Urs and Straube, Arthur V and Besseling, Rut and Aarts, Dirk GAL and L{\"o}wen, Hartmut and Dullens, Roel PA},
  journal=NJoP,
  volume={19},
  number={1},
  pages={013010},
  year={2017},
  publisher={IOP Publishing}
}

@article{Kawasaki2006b,
  title={Interpolation of stochastic and deterministic reduced dynamics},
  author={Kawasaki, Kyozi},
  journal={Physica A: Statistical Mechanics and its Applications},
  volume={362},
  number={2},
  pages={249--260},
  year={2006},
  publisher={Elsevier}
}

@article{BiswasC2007,
  title={A novel approach to solvation time scale in nonpolar solvents via instability of solvent density modes},
  author={Biswas, R and Chakrabarti, J},
  journal={Journal of Physical Chemistry B},
  volume={111},
  number={49},
  pages={13743--13747},
  year={2007},
  publisher={ACS Publications}
}

@article{Magnico2012,
  title={Ion size effects on electric double layers and ionic transport through ion-exchange membrane systems},
  author={Magnico, Pierre},
  journal={Journal of Membrane Science},
  volume={415},
  pages={412--423},
  year={2012},
  publisher={Elsevier}
}

@article{QiS2017,
  title={Dynamic density functional theories for inhomogeneous polymer systems compared to {B}rownian dynamics simulations},
  author={Qi, Shuanhu and Schmid, Friederike},
  journal={Macromolecules},
  volume={50},
  number={24},
  pages={9831--9845},
  year={2017},
  publisher={ACS Publications}
}

@article{ArcherM2011,
  title={On the interplay between sedimentation and phase separation phenomena in two-dimensional colloidal fluids},
  author={Archer, Andrew J and Malijevsk{\`y}, Alexandr},
  journal=MP,
  volume={109},
  number={7-10},
  pages={1087--1099},
  year={2011},
  publisher={Taylor \& Francis}
}

@article{Onsager1931,
  title={Reciprocal relations in irreversible processes. I.},
  author={Onsager, Lars},
  journal=PR,
  volume={37},
  number={4},
  pages={405},
  year={1931},
  publisher={APS}
}

@article{Onsager1931b,
  title={Reciprocal relations in irreversible processes. II.},
  author={Onsager, Lars},
  journal=PR,
  volume={38},
  number={12},
  pages={2265},
  year={1931},
  publisher={APS}
}

@ARTICLE{Onsager1949,
  author       = {Onsager, L.},
  title        = {The effects of shape on the interaction of colloidal particles},
  journal      = AotNYAoS,
  publisher    = NYAoS,
  year         = {1949},
  volume       = {51},
  number       = {4},
  pages        = {627-659},
  month        = MAY
}
% doi = {10.1111/j.1749-6632.1949.tb27296.x}
% url = {http://adsabs.harvard.edu/abs/1949NYASA..51..627O}

@ARTICLE{VroegeL1992,
  author       = {Vroege, G.~J. and Lekkerkerker, H.~N.~W.},
  title        = {Phase transitions in lyotropic colloidal and polymer liquid crystals},
  journal      = RoPiP,
  publisher    = IoPP,
  year         = {1992},
  volume       = {55},
  number       = {8},
  pages        = {1241-1309},
  month        = AUG
}
% doi = {10.1088/0034-4885/55/8/003}
% url = {http://adsabs.harvard.edu/abs/1992RPPh...55.1241V}

@ARTICLE{MaierS1958,
  author       = {Maier, W. and Saupe, A.},
  title        = {Eine einfache molekulare Theorie des nematischen kristallinfl{\"u}ssigen Zustandes},
  journal      = ZfNTA,
  publisher    = VdZfN,
  year         = {1958},
  volume       = {13},
  pages        = {564-566},
  month        = JUL
}
% url = {http://adsabs.harvard.edu/abs/1958ZNatA..13..564M}

@ARTICLE{MaierS1959,
  author       = {Maier, W. and Saupe, A.},
  title        = {Eine einfache molekularstatistische Theorie der nematischen kristallinfl{\"u}ssigen Phase. Teil I},
  journal      = ZfNTA,
  publisher    = VdZfN,
  year         = {1959},
  volume       = {14},
  pages        = {882-889},
  month        = OCT
}
% url = {http://adsabs.harvard.edu/abs/1959ZNatA..14..882M}

@ARTICLE{MaierS1960,
  author       = {Maier, W. and Saupe, A.},
  title        = {Eine einfache molekularstatistische Theorie der nematischen kristallinfl{\"u}ssigen Phase. Teil II},
  journal      = ZfNTA,
  publisher    = VdZfN,
  year         = {1960},
  volume       = {15},
  pages        = {287-292},
  month        = APR
}
% url = {http://adsabs.harvard.edu/abs/1960ZNatA..15..287M}

@ARTICLE{McMillan1971,
  author       = {McMillan, W.~L.},
  title        = {Simple molecular model for the smectic A phase of liquid crystals},
  journal      = PRA,
  publisher    = APS,
  year         = {1971},
  volume       = {4},
  number       = {3},
  pages        = {1238-1246},
  month        = SEP
}
% doi = {10.1103/PhysRevA.4.1238}
% url = {http://adsabs.harvard.edu/abs/1971PhRvA...4.1238M}

@ARTICLE{Ericksen1959,
  author       = {Ericksen, J.~L.},
  title        = {Anisotropic fluids},
  journal      = AfRMaA,
  publisher    = SPR,
  year         = {1959},
  volume       = {4},
  number       = {1},
  pages        = {231-237},
  month        = JAN
}
% doi = {10.1007/BF00281389}
% url = {http://adsabs.harvard.edu/abs/1959ArRMA...4..231E}

@ARTICLE{Ericksen1961,
  author       = {Ericksen, J.~L.},
  title        = {Conservation laws for liquid crystals},
  journal      = JoR,
  publisher    = AIoP,
  year         = {1961},
  volume       = {5},
  number       = {1},
  pages        = {23-34},
  month        = MAR
}
% doi = {10.1122/1.548883}
% url = {http://adsabs.harvard.edu/abs/1961JRheo...5...23E}

@ARTICLE{Leslie1966,
  author       = {Leslie, F.~M.},
  title        = {Some constitutive equations for anisotropic fluids},
  journal      = QJoMaAM,
  publisher    = OUP,
  year         = {1966},
  volume       = {19},
  number       = {3},
  pages        = {357-370},
  month        = APR
}
% doi = {10.1093/qjmam/19.3.357}
% url = {http://qjmam.oxfordjournals.org/content/19/3/357.full.pdf+html}

@ARTICLE{deGennes1971,
  author       = {de~Gennes, P.-G.},
  title        = {Short range order effects in the isotropic phase of nematics and cholesterics},
  journal      = MCaLC,
  publisher    = TF,
  year         = {1971},
  volume       = {12},
  number       = {3},
  pages        = {193-214},
  month        = FEB
}
% doi = {10.1080/15421407108082773}
% url = {http://dx.doi.org/10.1080/15421407108082773}

@BOOK{LandauL2008,
  author       = {Landau, Lev~D. and Lifschitz, Evgenij~M.},
  title        = {{S}tatistische {P}hysik {I}},
  publisher    = HD,
  year         = {2008},
  volume       = {5},
  series       = {{L}ehrbuch der theoretischen {P}hysik},
  address      = FaM,
  edition      = {8},
  month        = DEC,
  pages        = {517},
  isbn         = {978-3-817-11330-9}
}

@BOOK{LandauL1996,
  author       = {Landau, Lev~D. and Lifshitz, Evgenij~M.},
  title        = {Statistical Physics {I}},
  publisher    = BH,
  year         = {1996},
  volume       = {5},
  series       = {Landau and Lifshitz: Course of Theoretical Physics},
  address      = OX,
  edition      = {3},
  month        = AUG,
  pages        = {544},
  isbn         = {978-0-750-63372-7}
}
% ISBN-10: 0-750-63372-7

@BOOK{LandauL1991,
  author       = {Landau, Lev~D. and Lifschitz, Evgenij~M.},
  title        = {Hydrodynamik},
  publisher    = AV,
  year         = {1991},
  volume       = {6},
  series       = {{L}ehrbuch der theoretischen {P}hysik},
  address      = BER,
  edition      = {5},
  month        = DEC,
  pages        = {683},
  isbn         = {3-055-00070-6}
}

@BOOK{LandauL1987,
  author       = {Landau, Lev~D. and Lifshitz, Evgenij~M.},
  title        = {Fluid Mechanics},
  publisher    = BH,
  year         = {1987},
  volume       = {6},
  series       = {Landau and Lifshitz: Course of Theoretical Physics},
  address      = OX,
  edition      = {2},
  month        = AUG,
  pages        = {552},
  isbn         = {978-0-750-62767-2}
}
% ISBN-10: 0-750-62767-0

@BOOK{LandauL1991a,
  author       = {Landau, Lev~D. and Lifschitz, Evgenij~M.},
  title        = {Elastizit{\"a}tstheorie},
  publisher    = HD,
  year         = {1991},
  volume       = {7},
  series       = {{L}ehrbuch der theoretischen {P}hysik},
  address      = FaM,
  edition      = {7},
  month        = JAN,
  pages        = {223},
  isbn         = {3-817-11332-3}
}
% EAN: 9783817113323

@BOOK{LandauL2005,
  author       = {Landau, Lev~D. and Lifshitz, Evgenij~M.},
  title        = {Theory of Elasticity},
  publisher    = BH,
  year         = {2005},
  volume       = {7},
  series       = {Landau and Lifshitz: Course of Theoretical Physics},
  address      = OX,
  edition      = {3},
  month        = JAN,
  pages        = {195},
  isbn         = {978-0-750-62633-0}
}
% ISBN-10: 0-750-62633-X

@BOOK{LandauL1992,
  author       = {Landau, Lev~D. and Lifschitz, Evgenij~M.},
  title        = {{S}tatistische {P}hysik {II}},
  publisher    = HD,
  year         = {1992},
  volume       = {9},
  series       = {{L}ehrbuch der theoretischen {P}hysik},
  address      = FaM,
  edition      = {4},
  month        = JAN,
  pages        = {404},
  isbn         = {3-817-11334-X}
}

@BOOK{LandauL1990,
  author       = {Landau, Lev~D. and Lifschitz, Evgenij~M.},
  title        = {{P}hysikalische {K}inetik},
  publisher    = HD,
  year         = {1990},
  volume       = {10},
  series       = {{L}ehrbuch der theoretischen {P}hysik},
  address      = FaM,
  edition      = {2},
  month        = DEC,
  pages        = {480},
  isbn         = {978-3-817-11335-4}
}

@BOOK{LifshitzP1981,
  author       = {Lifshitz, E.~M. and Pitaevskii, L.~P.},
  title        = {Physical Kinetics},
  publisher    = BH,
  year         = {1981},
  volume       = {10},
  series       = {Landau and Lifshitz: Course of Theoretical Physics},
  address      = OX,
  edition      = {1},
  month        = JAN,
  pages        = {625},
  isbn         = {978-0-750-62635-4}
}
% ISBN-10: 0-7506-2635-6

@INCOLLECTION{Fix1983,
  author       = {Fix, G.~J.},
  title        = {Phase Field Models for Free Boundary Problems},
  booktitle    = {Free Boundary Problems: Theory and Applications},
  publisher    = {Pitman},
  year         = {1983},
  editor       = {Fasano, A. and Primicerio, M.},
  volume       = {2},
  series       = {Research Notes in Mathematics Series},
  pages        = {580-589},
  address      = {Boston}
}

@INCOLLECTION{Langer1986,
  author       = {Langer, J.~S.},
  title        = {Models of Pattern Formation in First?Order Phase Transitions},
  booktitle    = {Directions in Condensed Matter Physics},
  publisher    = WSP,
  year         = {1986},
  editor       = {Grinstein, G. and Mazenko, G.},
  volume       = {1},
  series       = {World Scientific Series on Directions in Condensed Matter Physics},
  pages        = {165-186},
  address      = SIN,
  isbn         = {9-971-97842-3}
}

@ARTICLE{HoughEtAl2009,
  author       = {Hough, L.~E. and Spannuth, M. and Nakata, M. and Coleman, D.~A. and Jones, C.~D. and Dantlgraber, G. and Tschierske, C. and Watanabe, J. and K{\"o}rblova, E. and Walba, D.~M. and Maclennan, J.~E. and Glaser, M.~A. and Clark, N.~A.},
  title        = {Chiral Isotropic Liquids from Achiral Molecules},
  journal      = SCI,
  publisher    = AAftAoS,
  year         = {2009},
  volume       = {325},
  number       = {5939},
  pages        = {452-456},
  month        = JUL
}
% doi = {10.1126/science.1170028}
% url = {http://adsabs.harvard.edu/abs/2009Sci...325..452H}

@BOOK{GreinerNS1993,
  author       = {Greiner, Walter and Neise, Ludwig and St{\"o}cker, Horst},
  title        = {{T}hermodynamik und {S}tatistische {M}echanik},
  publisher    = HD,
  year         = {1993},
  month        = JAN,
  volume       = {9},
  series       = {{T}heoretische {P}hysik},
  address      = FaM,
  edition      = {2},
  pages        = {579},
  isbn         = {3-817-11262-9}
}

@BOOK{Greiner2002,
  author       = {Greiner, Walter},
  title        = {{K}lassische {E}lektrodynamik},
  publisher    = HD,
  year         = {2002},
  month        = OCT,
  volume       = {3},
  series       = {{T}heoretische {P}hysik},
  address      = FaM,
  edition      = {6},
  pages        = {544},
  isbn         = {3-817-11660-8}
}

@BOOK{Greiner2005,
  author       = {Greiner, Walter},
  title        = {{Q}uantenmechanik: {E}inf{\"u}hrung},
  publisher    = HD,
  year         = {2005},
  month        = DEC,
  volume       = {4},
  series       = {{T}heoretische {P}hysik},
  address      = FaM,
  edition      = {6},
  pages        = {494},
  isbn         = {3-817-11765-5}
}

@BOOK{Greiner2009,
  author       = {Greiner, Walter},
  title        = {Quantum Mechanics: An Introduction},
  publisher    = SPR,
  year         = {2009},
  month        = FEB,
  address      = BER,
  edition      = {4},
  pages        = {510},
  isbn         = {3-540-67458-6}
}

@BOOK{GreinerM2005,
  author       = {Greiner, Walter and M{\"u}ller, Berndt},
  title        = {{Q}uantenmechanik: {S}ymmetrien},
  publisher    = HD,
  year         = {2005},
  month        = JAN,
  volume       = {5},
  series       = {{T}heoretische {P}hysik},
  address      = FaM,
  edition      = {4},
  pages        = {510},
  isbn         = {3-817-11616-0}
}

@BOOK{Dhont1996,
  author       = {Dhont, Jan~K.~G.},
  title        = {An Introduction to Dynamics of Colloids},
  publisher    = ES,
  year         = {1996},
  volume       = {2},
  series       = {Studies in Interface Science},
  address      = {Amsterdam},
  edition      = {1},
  month        = JUN,
  pages        = {642},
  isbn         = {0-444-82009-4}
}

@BOOK{HansenMD2009,
  author       = {Hansen, Jean-Pierre and McDonald, Ian~R.},
  title        = {Theory of Simple Liquids: with Applications to Soft Matter},
  publisher    = ELSAP,
  year         = {2009},
  address      = OX,
  edition      = {4th},
  month        = AUG,
  pages        = {636},
  isbn         = {0-123-87032-1}
}
% EAN=978 0 123 87032 2

@BOOK{BarratH2003,
  author       = {Barrat, Jean-Louis and Hansen, Jean-Pierre},
  title        = {Basic Concepts for Simple and Complex Liquids},
  publisher    = CUP,
  year         = {2003},
  address      = CAM,
  edition      = {1},
  month        = MAR,
  pages        = {308},
  isbn         = {0-521-78344-5}
}

@ARTICLE{Oseen1933,
  author       = {Oseen, C.~W.},
  title        = {The theory of liquid crystals},
  journal      = TotFS,
  publisher    = FS,
  number       = {140},
  year         = {1933},
  volume       = {29},
  pages        = {883-899}
}
% doi = {10.1039/TF9332900883}

@ARTICLE{Zoecher1933,
  author       = {Z{\"o}cher, H.},
  title        = {The effect of a magnetic field on the nematic state},
  journal      = TotFS,
  publisher    = FS,
  number       = {140},
  year         = {1933},
  volume       = {29},
  pages        = {945-957}
}
% doi = {10.1039/TF9332900945}

@ARTICLE{Frank1958,
  author       = {Frank, F.~C.},
  title        = {On the theory of liquid crystals},
  journal      = DotFS,
  publisher    = FS,
  year         = {1958},
  volume       = {25},
  number       = {1},
  pages        = {19-28},
  month        = FEB
}
% doi = {10.1039/DF9582500019}
% url = {http://pubs.rsc.org/en/Content/ArticleLanding/1958/DF/DF9582500019}

@BOOK{Grossmann1988,
  author       = {Gro\ss{}mann, Siegfried},
  title        = {Funktionalanalysis im Hinblick auf Anwendungen in der Physik},
  publisher    = {AULA-Verlag},
  year         = {1988},
  series       = {Studien-Text},
  address      = {Wiesbaden},
  edition      = {4},
  pages        = {317},
  isbn         = {3-891-04479-8}
}

@BOOK{GrayG1984,
  author       = {Gray, Christopher~G. and Gubbins, Keith~E.},
  title        = {Theory of Molecular Fluids: Fundamentals},
  publisher    = OUP,
  year         = {1984},
  volume       = {1},
  series       = {International Series of Monographs on Chemistry 9},
  address      = OX,
  edition      = {1},
  month        = DEC,
  pages        = {626},
  isbn         = {0-198-55602-0}
}

@BOOK{GrayGJ2011,
  author       = {Gray, Christopher~G. and Gubbins, Keith~E. and Joslin, Christopher~G.},
  title        = {Theory of Molecular Fluids: Applications},
  publisher    = OUP,
  year         = {2011},
  volume       = {2},
  series       = {International Series of Monographs on Chemistry 10},
  address      = OX,
  edition      = {1},
  month        = SEP,
  pages        = {784}, 
  isbn         = {0-198-55621-7}
}
% 978-0-198-55621-3

@BOOK{ArfkenW2005,
  author       = {Arfken, George~B. and Weber, Hans~J.},
  title        = {Mathematical Methods for Physicists},
  publisher    = ELS,
  year         = {2005},
  address      = OX,
  edition      = {6},
  month        = AUG,
  isbn         = {0-120-59876-0}
}
%   pages        = {1182},

@BOOK{Hobson1955,
  author       = {Hobson, Ernest~W.},
  title        = {The Theory of Spherical and Ellipsoidal Harmonics},
  publisher    = {Chelsea Publishing Company},
  year         = {1955},
  address      = NY,
  edition      = {1},
  month        = JUN,
  pages        = {500},
  isbn         = {0-82-840104-7}
}
 
@ARTICLE{Kalf1995,
  author       = {Kalf, Hubert},
  title        = {On the Expansion of a Function in Terms of Spherical Harmonics in Arbitrary Dimensions},
  journal      = {Bulletin of the Belgian Mathematical Society},
  publisher    = {Belgian Mathematical Society},
  year         = {1995},
  volume       = {2},
  number       = {4},
  pages        = {361-380}
}
% url = {http://www.emis.de/journals/BBMS/Bulletin/bul954/KALF.PDF}

@BOOK{MacRobertS1967,
  author       = {MacRobert, Thomas~Murray and Sneddon, Ian~Naismith},
  title        = {Spherical Harmonics: An Elementary Treatise on Harmonic Functions, with Applications},
  publisher    = {Pergamon Press},
  year         = {1967},
  volume       = {98},
  series       = {International series of monographs in pure and applied mathematics},
  address      = OX,
  edition      = {3},
  pages        = {349}
}

@BOOK{Weisstein2002,
  author       = {Weisstein, Eric~W.},
  title        = {CRC Concise Encyclopedia of Mathematics},
  publisher    = {Chapman \& Hall/CRC},
  year         = {2002},
  address      = {Boca Raton, Florida},
  edition      = {2},
  month        = DEC,
  pages        = {3252},
  isbn         = {1-584-88347-2}
}

@BOOK{Messiah1999,
  author       = {Messiah, Albert},
  title        = {Quantum Mechanics},
  publisher    = DP,
  year         = {1999},
  address      = NY,
  edition      = {1},
  month        = JUL,
  pages        = {1152},
  isbn         = {0-486-40924-4}
}

@ARTICLE{TonerTR2005,
  author       = {Toner, J. and Tu, Y. and Ramaswamy, S.},
  title        = {Hydrodynamics and phases of flocks},
  journal      = {Annals of Physics},
  publisher    = ELS,
  year         = {2005},
  volume       = {318},
  number       = {1},
  pages        = {170-244},
  month        = JUL
}
% doi = {10.1016/j.aop.2005.04.011}
% url = {http://adsabs.harvard.edu/abs/2005AnPhy.318..170T}

@article{TonerT1995,
  title={Long-range order in a two-dimensional dynamical {XY} model: how birds fly together},
  author={Toner, John and Tu, Yuhai},
  journal=PRL,
  volume={75},
  number={23},
  pages={4326},
  year={1995},
  publisher={APS}
}

@ARTICLE{LaugaP2009,
  author       = {Lauga, E. and Powers, T.~R.},
  title        = {The hydrodynamics of swimming microorganisms},
  journal      = {Reports on Progress in Physics},
  publisher    = IoP,
  year         = {2009},
  volume       = {9},
  number       = {72},
  pages        = {096601},
  month        = SEP
}
% doi = {10.1088/0034-4885/72/9/096601}
% url = {http://adsabs.harvard.edu/abs/2009RPPh...72i6601L}

@ARTICLE{Ramaswamy2010,
  author       = {Ramaswamy, Sriram},
  title        = {The mechanics and statistics of active matter},
  journal      = ARoCMP,
  publisher    = AR,
  year         = {2010},
  volume       = {1},
  number       = {1},
  pages        = {323-345},
  month        = AUG
}
% doi = {10.1146/annurev-conmatphys-070909-104101}
% url = {http://www.annualreviews.org/doi/abs/10.1146/annurev-conmatphys-070909-104101}

@INCOLLECTION{Loewen2010W,
  author       = {L{\"o}wen, Hartmut},
  title        = {Density Functional Theory for Inhomogeneous Fluids II: Statics, Dynamics, and Applications},
  booktitle    = {3rd Warsaw School of Statistical Physics},
  publisher    = {Warsaw University Press},
  year         = {2010},
  editor       = {Cichocki, B. and Napi{\'o}rkowski, M. and Piasecki, J.},
  type         = {chapter},
  chapter      = {3},
  pages        = {87-121},
  address      = {Warsaw, Poland},
  isbn         = {978-83-235-0602-7}
}

@ARTICLE{AnnaratoneEtAl2001,
  author       = {Annaratone, B.~M. and Khrapak, A.~G. and Ivlev, A.~V. and S{\"o}llner, G. and Bryant, P. and S{\"u}tterlin, R. and Konopka, U. and Yoshino, K. and Zuzic, M. and Thomas, H.~M. and Morfill, G.~E.},
  title        = {Levitation of cylindrical particles in the sheath of an rf plasma},
  journal      = PRE,
  publisher    = APS,
  year         = {2001},
  volume       = {63},
  number       = {3},
  pages        = {036406},
  month        = MAR
}
% doi = {10.1103/PhysRevE.63.036406}
% url = {http://adsabs.harvard.edu/abs/2001PhRvE..63c6406A}

@ARTICLE{IvlevKKAMY2003,
  author       = {Ivlev, A.~V. and Khrapak, A.~G. and Khrapak, S.~A. and
Annaratone, B.~M. and Morfill, G. and Yoshino, K.},
  title        = {Rodlike particles in gas discharge plasmas: theoretical model},
  journal      = PRE,
  publisher    = APS,
  year         = {2003},
  volume       = {68},
  number       = {2},
  pages        = {026403},
  month        = AUG
}
% doi = {10.1103/PhysRevE.68.026403}
% url = {http://adsabs.harvard.edu/abs/2003PhRvE..68b6403I}

@ARTICLE{BorciaBB2008,
  author       = {Borcia, R. and Borcia, I.~D. and Bestehorn, M.},
  title        = {Drops on an arbitrarily wetting substrate: A phase field description},
  journal      = PRE,
  publisher    = APS,
  year         = {2008},
  volume       = {78},
  number       = {6},
  pages        = {066307},
  month        = DEC
}
% doi = {10.1103/PhysRevE.78.066307}
% url = {http://adsabs.harvard.edu/abs/2008PhRvE..78f6307B}

@ARTICLE{AlandLV2010,
  author       = {Aland, S. and Lowengrub, J. and Voigt, A.},
  title        = {Two-phase flow in complex geometries: A diffuse domain approach},
  journal      = CMES,
  publisher    = TSP,
  year         = {2010},
  volume       = {57},
  number       = {1},
  pages        = {77-107},
  month        = FEB
}
% doi = {}
% url = {http://www.techscience.com/cmes/2010/v57n1_index.html}

@inproceedings{Thomas1927,
  title={The calculation of atomic fields},
  author={Thomas, Llewellyn H},
  booktitle={Mathematical Proceedings of the Cambridge Philosophical Society},
  volume={23},
  number={5},
  pages={542--548},
  year={1927},
  organization={Cambridge University Press}
}

@article{Fermi1928,
  title={Eine statistische {M}ethode zur {B}estimmung einiger {E}igenschaften des {A}toms und ihre {A}nwendung auf die {T}heorie des periodischen {S}ystems der {E}lemente},
  author={Fermi, Enrico},
  journal={Zeitschrift f{\"u}r Physik},
  volume={48},
  number={1-2},
  pages={73--79},
  year={1928},
  publisher={Springer}
}

@article{vanderWaals1894,
  title={Thermodynamische {T}heorie der {K}apillarit{\"a}t unter {V}oraussetzung stetiger {D}ichte{\"a}nderung},
  author={van der Waals, JD},
  journal={Zeitschrift f{\"u}r Physikalische Chemie},
  volume={13},
  number={1},
  pages={657--725},
  year={1894},
  publisher={De Gruyter Oldenbourg}
}

@ARTICLE{MarconiTCM2008,
  author       = {{Marini Bettolo Marconi}, Umberto and Tarazona, P. and Cecconi, F. and Melchionna, S.},
  title        = {Beyond dynamic density functional theory: the role of inertia},
  journal      = JoP:CM,
  publisher    = IoPP,
  year         = {2008},
  volume       = {20},
  number       = {49},
  pages        = {494233},
  month        = DEC
}
% doi = {10.1088/0953-8984/20/49/494233}
% url = {http://adsabs.harvard.edu/abs/2008JPCM...20W4233M}

@ARTICLE{MarconiM2009,
  author       = {{Marini Bettolo Marconi}, Umberto and Melchionna, S.},
  title        = {Kinetic theory of correlated fluids: from dynamic density functional to {L}attice {B}oltzmann methods},
  journal      = JoChemP,
  publisher    = AIoP,
  year         = {2009},
  volume       = {131},
  number       = {1},
  pages        = {014105},
  month        = JUL
}
% doi = {10.1063/1.3166865}
% url = {http://adsabs.harvard.edu/abs/2009JChPh.131a4105M}

@article{MamasakhlisovNP2008,
  title={Partially annealed disorder and collapse of like-charged macroions},
  author={Mamasakhlisov, Yevgeni S and Naji, Ali and Podgornik, Rudolf},
  journal={Journal of Statistical Physics},
  volume={133},
  number={4},
  pages={659},
  year={2008},
  publisher={Springer}
}

@article{MarconiTC2007,
  title={Theory of thermostatted inhomogeneous granular fluids: A self-consistent density functional description},
  author={{Marini Bettolo Marconi}, Umberto and Tarazona, Pedro and Cecconi, Fabio},
  journal=JoChemP,
  volume={126},
  number={16},
  pages={164904},
  year={2007},
  publisher={AIP}
}

@article{HowardNP2017,
  title={Stratification dynamics in drying colloidal mixtures},
  author={Howard, Michael P and Nikoubashman, Arash and Panagiotopoulos, Athanassios Z},
  journal={Langmuir},
  volume={33},
  number={15},
  pages={3685--3693},
  year={2017},
  publisher={ACS Publications}
}

@article{GranasyPTTP2014,
  title={Heterogeneous nucleation of/on nanoparticles: a density functional study using the phase-field crystal model},
  author={Gr{\'a}n{\'a}sy, L{\'a}szl{\'o} and Podmaniczky, Frigyes and T{\'o}th, Gyula I and Tegze, Gy{\"o}rgy and Pusztai, Tam{\'a}s},
  journal={Chemical Society Reviews},
  volume={43},
  number={7},
  pages={2159-2173},
  year={2014},
  publisher={Royal Society of Chemistry}
}

@article{RamirezAACA2006,
  title={Theoretical description of the ion transport across nanopores with titratable fixed charges},
  author={Ramirez, P and Aguilella-Arzo, M and Alcaraz, A and Cervera, J and Aguilella, VM},
  journal={Cell Biochemistry and Biophysics},
  volume={44},
  number={2},
  pages={287--312},
  year={2006},
  publisher={Springer}
}

@article{RexLLD2006,
  title={Ultrasoft colloids in cavities of oscillating size or sharpness},
  author={Rex, M and Likos, CN and L{\"o}wen, H and Dzubiella, J},
  journal=MP,
  volume={104},
  number={4},
  pages={527--540},
  year={2006},
  publisher={Taylor \& Francis}
}

@article{Yoshimori2005,
  title={Microscopic derivation of time-dependent density functional methods},
  author={Yoshimori, Akira},
  journal=PRE,
  volume={71},
  number={3},
  pages={031203},
  year={2005},
  publisher={APS}
}

@article{Yoshimori2011,
  title={Time-dependent density functional theory formulated using the interaction-site model},
  author={Yoshimori, Akira},
  journal={Journal of the Physical Society of Japan},
  volume={80},
  number={3},
  pages={034801},
  year={2011},
  publisher={The Physical Society of Japan}
}

@article{Schmidt2011,
  title={Statics and dynamics of inhomogeneous liquids via the internal-energy functional},
  author={Schmidt, Matthias},
  journal=PRE,
  volume={84},
  number={5},
  pages={051203},
  year={2011},
  publisher={APS}
}

@article{DongFWWZ2018,
  title={Convergence analysis and numerical implementation of a second order numerical scheme for the three-dimensional phase field crystal equation},
  author={Dong, Lixiu and Feng, Wenqiang and Wang, Cheng and Wise, Steven M and Zhang, Zhengru},
  journal={Computers \& Mathematics with Applications},
  volume={75},
  number={6},
  pages={1912--1928},
  year={2018},
  publisher={Elsevier}
}

@article{BaskaranBL2014,
  title={Kinetic density functional theory of freezing},
  author={Baskaran, Arvind and Baskaran, Aparna and Lowengrub, John},
  journal=JoChemP,
  volume={141},
  number={17},
  pages={174506},
  year={2014},
  publisher={AIP}
}

@article{WitmanW2006,
  title={A lattice model of vitrification and gelation},
  author={Witman, Jennifer E and Wang, Zhen-Gang},
  journal={Journal of Physical Chemistry B},
  volume={110},
  number={12},
  pages={6312--6324},
  year={2006},
  publisher={ACS Publications}
}

@article{XuCMYL2014,
  title={Modeling and simulating asymmetrical conductance changes in gramicidin pores},
  author={Xu, Shixin and Chen, Minxin and Majd, Sheereen and Yue, Xingye and Liu, Chun},
  journal={Computational and Mathematical Biophysics},
  volume={2},
  number={1},
  year={2014},
  publisher={De Gruyter Open}
}

@article{ChauviereLC2012,
  title={Dynamic density functional theory of solid tumor growth: preliminary models},
  author={Chauviere, Arnaud and Hatzikirou, Haralambos and Kevrekidis, Ioannis G and Lowengrub, John S and Cristini, Vittorio},
  journal={AIP Advances},
  volume={2},
  number={1},
  pages={011210},
  year={2012},
  publisher={AIP}
}

@ARTICLE{MarconiM2010,
  author       = {{Marini Bettolo Marconi}, Umberto and Melchionna, S.},
  title        = {Dynamic density functional theory versus kinetic theory of simple fluids},
  journal      = JoP:CM,
  publisher    = IoPP,
  year         = {2010},
  volume       = {22},
  number       = {36},
  pages        = {364110},
  month        = SEP
}
% doi = {}
% url = {}

@ARTICLE{EspanolAZ2009,
  author       = {Espa{\~n}ol, P. and Anero, J.~G. and Z{\'u}{\~n}iga, I.},
  title        = {Microscopic derivation of discrete hydrodynamics},
  journal      = JoChemP,
  publisher    = AIoP,
  year         = {2009},
  volume       = {131},
  number       = {24},
  pages        = {244117},
  month        = DEC
}
% doi = {10.1063/1.3274222}
% url = {http://adsabs.harvard.edu/abs/2009JChPh.131x4117E}

@ARTICLE{Archer2006,
  author       = {Archer, A.~J.},
  title        = {Dynamical density functional theory for dense atomic liquids},
  journal      = JoP:CM,
  publisher    = IoPP,
  year         = {2006},
  volume       = {18},
  number       = {24},
  pages        = {5617-5628},
  month        = JUN
}
% doi = {10.1088/0953-8984/18/24/004}
% url = {http://adsabs.harvard.edu/abs/2006JPCM...18.5617A}

@ARTICLE{Archer2009,
  author       = {Archer, A.~J.},
  title        = {Dynamical density functional theory for molecular and colloidal fluids: a microscopic approach to fluid mechanics},
  journal      = JoChemP,
  publisher    = AIoP,
  year         = {2009},
  volume       = {130},
  number       = {1},
  pages        = {014509},
  month        = JAN
}
% doi = {10.1063/1.3054633}
% url = {http://adsabs.harvard.edu/abs/2009JChPh.130a4509A}

@ARTICLE{RexL2008,
  author       = {Rex, M. and L{\"o}wen, H.},
  title        = {Dynamical density functional theory with hydrodynamic interactions and colloids in unstable traps},
  journal      = PRL,
  publisher    = APS,
  year         = {2008},
  volume       = {101},
  number       = {14},
  pages        = {148302},
  month        = OCT
}
% doi = {10.1103/PhysRevLett.101.148302}
% url = {http://adsabs.harvard.edu/abs/2008PhRvL.101n8302R}

@ARTICLE{AsaroT1972,
  author       = {Asaro, R.~J. and Tiller, W.~A.},
  title        = {Interface morphology development during stress corrosion cracking: Part I. Via surface diffusion},
  journal      = {Metallurgical Transactions},
  publisher    = SPR,
  year         = {1972},
  volume       = {3},
  number       = {7},
  pages        = {1789-1796},
  month        = JUL
}
% doi = {10.1007/BF02642562}
% url = {http://adsabs.harvard.edu/abs/1972MT......3.1789A}

@ARTICLE{Grinfeld1986,
  author       = {Grinfeld, M.~A.},
  title        = {Instability of the interface between a nonhydrostatically stressed elastic body and a melt},
  journal      = {Akademiia Nauk SSSR Doklady},
  publisher    = {USSR Academy of Sciences},
  year         = {1986},
  volume       = {290},
  number       = {6},
  pages        = {1358-1363},
  month        = JAN
}
% url = {http://adsabs.harvard.edu/abs/1986DoSSR.290.1358G}

@ARTICLE{SakaguchiB1997,
  author       = {Sakaguchi, H. and Brand, H.~R.},
  title        = {A modulation mediated hexa\-gon-rec\-tan\-gle transition in pattern forming systems},
  journal      = PLA,
  publisher    = ELS,
  year         = {1997},
  volume       = {227},
  number       = {3-4},
  pages        = {209-215},
  month        = FEB
}
% doi = {10.1016/S0375-9601(97)00032-7}
% url = {http://adsabs.harvard.edu/abs/1997PhLA..227..209S}

@ARTICLE{WeyerichDACK1990,
  author       = {Weyerich, Bertram and D{'}Aguanno, Bruno and Canessa, Enrique and Klein, Rudolf},
  title        = {Structure and dynamics of suspensions of charged rod-like particles},
  journal      = FDotCS,
  publisher    = RSoC,
  year         = {1990},
  volume       = {90},
  number       = {1},
  pages        = {245-259},
  month        = JUL
}
% doi = {10.1039/DC9909000245}
% url = {http://dx.doi.org/10.1039/DC9909000245}

@ARTICLE{vanderBeekDWVL2008,
  author       = {van der Beek, D. and Davidson, P. and Wensink, H.~H. and Vroege, G.~J. and Lekkerkerker, H.~N.~W.},
  title        = {Influence of a magnetic field on the nematic phase of hard colloidal platelets},
  journal      = PRE,
  publisher    = APS,
  year         = {2008},
  volume       = {77},
  number       = {3},
  pages        = {031708},
  month        = MAR
}
% doi = {10.1103/PhysRevE.77.031708}
% url = {http://adsabs.harvard.edu/abs/2008PhRvE..77c1708V}

@ARTICLE{WatanabeT2007,
  author       = {Watanabe, G. and Tabe, Y.},
  title        = {Tilted and non-tilted liquid crystalline Langmuir monolayers: analogy to bulk smectic phases},
  journal      = JotPSoJ,
  publisher    = PSoJ,
  year         = {2007},
  volume       = {76},
  number       = {9},
  pages        = {094602},
  month        = SEP
}
% doi = {10.1143/JPSJ.76.094602}
% url = {http://adsabs.harvard.edu/abs/2007JPSJ...76i4602W}

@ARTICLE{TavaresHTdG2009,
  author       = {Tavares, J.~M. and Holder, B. and {Telo da Gama}, M.~M.},
  title        = {Structure and phase diagram of self-assembled rigid rods: equilibrium polydispersity and nematic ordering in two dimensions},
  journal      = PRE,
  publisher    = APS,
  year         = {2009},
  volume       = {79},
  number       = {2},
  pages        = {021505},
  month        = FEB
}
% doi = {10.1103/PhysRevE.79.021505}
% url = {http://adsabs.harvard.edu/abs/2009PhRvE..79b1505T}

@ARTICLE{Vink2007,
  author       = {Vink, R.~L.~C.},
  title        = {Liquid crystals in two dimensions: first-order phase transitions and nonuniversal critical behavior},
  journal      = PRL,
  publisher    = APS,
  year         = {2007},
  volume       = {98},
  number       = {21},
  pages        = {217801},
  month        = MAY
}
% doi = {10.1103/PhysRevLett.98.217801}
% url = {http://adsabs.harvard.edu/abs/2007PhRvL..98u7801V}

@ARTICLE{TothDY2002,
  author       = {T{\'o}th, G.~I. and Denniston, C. and Yeomans, J.~M.},
  title        = {Hydrodynamics of topological defects in nematic liquid crystals},
  journal      = PRL,
  publisher    = APS,
  year         = {2002},
  volume       = {88},
  number       = {10},
  pages        = {105504},
  month        = MAR
}
% doi = {10.1103/PhysRevLett.88.105504}
% url = {http://adsabs.harvard.edu/abs/2002PhRvL..88j5504T}

@ARTICLE{LopezLRPC2010,
  author       = {L{\'o}pez, L.~G. and Linares, D.~H. and {Ramirez-Pastor}, A.~J. and Cannas, S.~A.},
  title        = {Phase diagram of self-assembled rigid rods on two-dimensional lattices: theory and Monte Carlo simulations},
  journal      = JoChemP,
  publisher    = AIoP,
  year         = {2010},
  volume       = {133},
  number       = {13},
  pages        = {134706},
  month        = SEP
}
% doi = {doi:10.1063/1.3496482}
% url = {http://jcp.aip.org/resource/1/jcpsa6/v133/i13/p134706_s1}

@ARTICLE{GrafLS1997,
  author       = {Graf, H. and L{\"o}wen, H. and Schmidt, M.},
  title        = {Cell theory for the phase diagram of hard spherocylinders},
  journal      = PiCaPS,
  publisher    = SPR,
  year         = {1997},
  volume       = {104},
  number       = {1},
  pages        = {177-179},
  month        = JAN
}
% isbn = {3-7985-1085-7}
% doi = {10.1007/BFb0110773}
% url = {http://www.springerlink.com/content/n012485282110118/}

@ARTICLE{GrafL1999b,
  author       = {Graf, H. and L{\"o}wen, H.},
  title        = {Phase diagram of tobacco mosaic virus solutions},
  journal      = PRE,
  publisher    = APS,
  year         = {1999},
  volume       = {59},
  number       = {2},
  pages        = {1932-1942},
  month        = FEB
}
% doi = {10.1103/PhysRevE.59.1932}
% url = {http://adsabs.harvard.edu/abs/1999PhRvE..59.1932G}

@ARTICLE{FrenkelM1985,
  author       = {Frenkel, D. and Mulder, B.~M.},
  title        = {The hard ellipsoid-of-revolution fluid},
  journal      = {Molecular Physics},
  publisher    = {Taylor \& Francis},
  year         = {1985},
  volume       = {55},
  number       = {5},
  pages        = {1171-1192},
  month        = JAN
} 
% doi = {10.1080/00268978500101971}
% url = {http://adsabs.harvard.edu/abs/1985MolPh..55.1171F}

@ARTICLE{deMiguelMdRBA1996,
  author       = {de~Miguel, E. and Mart{\'{\i}}n~del~Rio, E. and Brown, J.~T. and Allen, M.~P.},
  title        = {Effect of the attractive interactions on the phase behavior of the Gay-Berne liquid crystal model},
  journal      = JoChemP,
  publisher    = AIoP,
  year         = {1996},
  volume       = {105},
  number       = {10},
  pages        = {4234-4249},
  month        = SEP
}
% doi = {10.1063/1.472292}
% url = {http://adsabs.harvard.edu/abs/1996JChPh.105.4234D}

@ARTICLE{KondratBH2010,
  author       = {Kondrat, S. and Bier, M. and Harnau, L.},
  title        = {Phase behavior of ionic liquid crystals},
  journal      = JoChemP,
  publisher    = AIoP,
  year         = {2010},
  volume       = {132},
  number       = {18},
  pages        = {184901},
  month        = MAY
}
% doi = {10.1063/1.3417384}
% url = {http://adsabs.harvard.edu/abs/2010JChPh.132r4901K}

@ARTICLE{McDonaldAS2000,
  author       = {McDonald, A.~J. and Allen, M.~P. and Schmid, F.},
  title        = {Surface tension of the isotropic-nematic interface},
  journal      = PRE,
  publisher    = APS,
  year         = {2000},
  volume       = {63},
  number       = {1},
  pages        = {010701(R)},
  month        = JAN
}
% doi = {10.1103/PhysRevE.63.010701}
% url = {http://adsabs.harvard.edu/abs/2001PhRvE..63a0701M}

@ARTICLE{vanderBeekEtAl2006,
  author       = {van~der~Beek, D. and Reich, H. and van~der~Schoot, P. and Dijkstra, M. and Schilling, T. and Vink, R. and Schmidt, M. and van~Roij, R. and Lekkerkerker, H.},
  title        = {Isotropic-nematic interface and wetting in suspensions of colloidal platelets},
  journal      = PRL,
  publisher    = APS,
  year         = {2006},
  volume       = {97},
  number       = {8},
  pages        = {087801},
  month        = AUG
}
% doi = {10.1103/PhysRevLett.97.087801}
% url = {http://adsabs.harvard.edu/abs/2006PhRvL..97h7801V}

@ARTICLE{HaertelBL2010,
  author       = {H{\"a}rtel, A. and Blaak, R. and L{\"o}wen, H.},
  title        = {Towing, breathing, splitting, and overtaking in driven colloidal liquid crystals},
  journal      = PRE,
  publisher    = APS,
  year         = {2010},
  volume       = {81},
  number       = {5},
  pages        = {051703},
  month        = MAY
}
% doi = {10.1103/PhysRevE.81.051703}
% url = {http://adsabs.harvard.edu/abs/2010PhRvE..81e1703H}

@ARTICLE{vanTeeffelenL2008,
  author       = {van~Teeffelen, S. and L{\"o}wen, H.},
  title        = {Dynamics of a {B}rownian circle swimmer},
  journal      = PRE,
  publisher    = APS,
  year         = {2008},
  volume       = {78},
  number       = {2},
  pages        = {020101},
  month        = AUG
}
% doi = {10.1103/PhysRevE.78.020101}
% url = {http://adsabs.harvard.edu/abs/2008PhRvE..78b0101V}

@ARTICLE{MkhontaEG2010,
  author       = {Mkhonta, S. and Elder, K.~R. and Grant, M.},
  title        = {Novel mechanical properties in lamellar phases of liquid-crystalline diblock copolymers},
  journal      = EPJE,
  publisher    = EDPS,
  year         = {2010},
  volume       = {32},
  number       = {4},
  pages        = {349-355},
  month        = AUG
}
% doi = {10.1140/epje/i2010-10636-9}
% url = {http://dx.doi.org/10.1140/epje/i2010-10636-9}

@INPROCEEDINGS{Oxtoby1991,
  author       = {Oxtoby, D.},
  title        = {Course 3: Crystallization of liquids: a density functional approach},
  booktitle    = {Liquids, Freezing and Glass Transition},
  publisher    = {North {H}olland, {E}lsevier {S}cience {P}ublishers {B}. {V}.},
  year         = {1991},
  editor       = {Hansen, J.-P. and Levesque, D. and {Zinn-Justin}, J.},
  volume       = {1},
  series       = {Proceedings of the {L}es {H}ouches {S}ummer {S}chool, {C}ourse {LI}, 3-28 {J}uly 1989},
  pages        = {145-192},
  address      = AMST,
  month        = JUL,
  organization = {{USMG}, {NATO} {A}dvanced {S}tudy {I}nstitute}
}
% isbn         = {0-444-88928-0}

@INPROCEEDINGS{Goetze1991,
  author       = {G{\"o}tze, Wolfgang},
  title        = {Course 5: Aspects of structural glass transitions},
  booktitle    = {Liquids, Freezing and Glass Transition},
  publisher    = {North {H}olland, {E}lsevier {S}cience {P}ublishers {B}. {V}.},
  year         = {1991},
  editor       = {Hansen, J.-P. and Levesque, D. and {Zinn-Justin}, J.},
  volume       = {2},
  series       = {Proceedings of the {L}es {H}ouches {S}ummer {S}chool, {C}ourse {LI}, 3-28 {J}uly 1989},
  pages        = {287-503},
  address      = AMST,
  month        = JUL,
  organization = {{USMG}, {NATO} {A}dvanced {S}tudy {I}nstitute}
}
% isbn         = {0-444-88928-0}

@INPROCEEDINGS{Frenkel1991,
  author       = {Frenkel, D.},
  title        = {Course 9: Statistical Mechanics of Liquid Crystals},
  booktitle    = {Liquids, Freezing and Glass Transition},
  publisher    = {North {H}olland, {E}lsevier {S}cience {P}ublishers {B}. {V}.},
  year         = {1991},
  editor       = {Hansen, J.-P. and Levesque, D. and {Zinn-Justin}, J.},
  volume       = {2},
  series       = {Proceedings of the {L}es {H}ouches {S}ummer {S}chool, {C}ourse {LI}, 3-28 {J}uly 1989},
  pages        = {689-762},
  address      = AMST,
  month        = JUL,
  organization = {{USMG}, {NATO} {A}dvanced {S}tudy {I}nstitute}
}
% isbn         = {0-444-88928-0}

@INPROCEEDINGS{Pusey1991,
  author       = {Pusey, P.~N.},
  title        = {Course 10: Colloidal suspensions},
  booktitle    = {Liquids, Freezing and Glass Transition},
  publisher    = {North {H}olland, {E}lsevier {S}cience {P}ublishers {B}. {V}.},
  year         = {1991},
  editor       = {Hansen, J.-P. and Levesque, D. and {Zinn-Justin}, J.},
  volume       = {2},
  series       = {Proceedings of the {L}es {H}ouches {S}ummer {S}chool, {C}ourse {LI}, 3-28 {J}uly 1989},
  pages        = {763-942},
  address      = AMST,
  month        = JUL,
  organization = {{USMG}, {NATO} {A}dvanced {S}tudy {I}nstitute}
}
% isbn         = {0-444-88928-0}

@ARTICLE{MarechalD2008,
  author       = {Marechal, M. and Dijkstra, M.},
  title        = {Stability of orientationally disordered crystal structures of colloidal hard dumbbells},
  journal      = PRE,
  publisher    = APS,
  year         = {2008},
  volume       = {77},
  number       = {6},
  pages        = {061405},
  month        = JUN
}
% doi = {10.1103/PhysRevE.77.061405}
% url = {http://adsabs.harvard.edu/abs/2008PhRvE..77f1405M}

@ARTICLE{DemiroersJvKvBI2010,
  author       = {Demir{\"o}rs, Ahmet~Faik and Johnson, Patrick~M. and Kats, Carlos~M.~van and van~Blaaderen, Alfons and Imhof, Arnout},
  title        = {Directed self-assembly of colloidal dumbbells with an electric field},
  journal      = LANG,
  publisher    = ACS,
  year         = {2010},
  volume       = {26},
  number       = {18},
  pages        = {14466-14471},
  month        = SEP
}
% doi = {10.1021/la102134w}
% url = {http://pubs.acs.org/doi/abs/10.1021/la102134w}

@ARTICLE{GerbodeAOLEC2010,
  author       = {Gerbode, S.~J. and Agarwal, U. and Ong, D.~C. and Liddell, C.~M. and Escobedo, F. and Cohen, I.},
  title        = {Glassy dislocation dynamics in 2D colloidal dimer crystals},
  journal      = PRL,
  publisher    = APS,
  year         = {2010},
  volume       = {105},
  number       = {7},
  pages        = {078301},
  month        = AUG
}
% doi = {10.1103/PhysRevLett.105.078301}
% url = {http://adsabs.harvard.edu/abs/2010PhRvL.105g8301G}

@article{Mermin1965,
  title={Thermal properties of the inhomogeneous electron gas},
  author={Mermin, N David},
  journal=PR,
  volume={137},
  number={5A},
  pages={A1441},
  year={1965},
  publisher={APS}
}

@ARTICLE{Mermin1979,
  author       = {Mermin, N.~D.},
  title        = {The topological theory of defects in ordered media},
  journal      = RoMP,
  publisher    = APS,
  year         = {1979},
  volume       = {51},
  number       = {3},
  pages        = {591-648},
  month        = JUL
}
% doi = {10.1103/RevModPhys.51.591}
% url = {http://adsabs.harvard.edu/abs/1979RvMP...51..591M}

@ARTICLE{TabeYNYY2003,
  author       = {Tabe, Y. and Yamamoto, T. and Nishiyama, I. and Yoneya, M. and Yokoyama, H.},
  title        = {Ferroelectric nematic monolayer},
  journal      = JJoAP,
  publisher    = {Japan Society of Applied Physics},
  year         = {2003},
  volume       = {42},
  number       = {4B},
  pages        = {L406-L409},
  month        = APR
}
% doi = {10.1143/JJAP.42.L406}Part 2
% url = {http://adsabs.harvard.edu/abs/2003JaJAP..42L.406T}

@ARTICLE{BarratHP1987,
  author       = {Barrat, J.-L. and Hansen, J.-P. and Pastore, G.},
  title        = {Factorization of the triplet direct correlation function in dense fluids},
  journal      = PRL,
  publisher    = APS,
  year         = {1987},
  volume       = {58},
  number       = {20},
  pages        = {2075-2078},
  month        = MAY
}
% doi = {10.1103/PhysRevLett.58.2075}
% url = {http://adsabs.harvard.edu/abs/1987PhRvL..58.2075B}

@ARTICLE{ZhuEtAl2010,
  author       = {Zhu, Yanwu and Murali, Shanthi and Cai, Weiwei and Li, Xuesong and Suk, Ji~Won and Potts, Jeffrey~R. and Ruoff, Rodney~S.},
  title        = {Graphene and Graphene Oxide: Synthesis, Properties, and Applications},
  journal      = {Advanced Materials},
  publisher    = WILEYVCH,
  year         = {2010},
  volume       = {22},
  number       = {35},
  pages        = {1521-4095},
  month        = SEP
}
% doi = {10.1002/adma.201001068}
% url = {http://dx.doi.org/10.1002/adma.201001068}

@ARTICLE{NovoselovEtAl2004,
  author       = {Novoselov, K.~S. and Geim, A.~K. and Morozov, S.~V. and Jiang, D. and Zhang, Y. and Dubonos, S.~V. and Grigorieva, I.~V. and Firsov, A.~A.},
  title        = {Electric Field Effect in Atomically Thin Carbon Films},
  journal      = SCI,
  publisher    = AAftAoS,
  year         = {2004},
  volume       = {306},
  number       = {5696},
  pages        = {666-669},
  month        = OCT
}
% doi = {10.1126/science.1102896}
% url = {http://adsabs.harvard.edu/abs/2004Sci...306..666N}

@ARTICLE{GeimNovoselov2007,
  author       = {Geim, A.~K. and Novoselov, K.~S.},
  title        = {The rise of graphene},
  journal      = NAT,
  publisher    = NPG,
  year         = {2007},
  volume       = {6},
  number       = {???},
  pages        = {183-191},
  month        = FEB
}
% ISSN = {1476-1122}
% doi = {doi:10.1038/nmat1849}
% url = {http://adsabs.harvard.edu/abs/2007cond.mat..2595G}

@ARTICLE{NovoselovEtAl2005,
  author       = {Novoselov, K.~S. and Jiang, D. and Schedin, F. and Booth, T.~J. and Khotkevich, V.~V. and Morozov, S.~V. and Geim, A.~K.},
  title        = {Two-dimensional atomic crystals},
  journal      = PotNAoS,
  publisher    = NAoS,
  year         = {2005},
  volume       = {102},
  number       = {30},
  pages        = {10451-10453},
  month        = JUL
}
% doi = {10.1073/pnas.0502848102}
% url = {http://adsabs.harvard.edu/abs/2005PNAS..10210451N}

@ARTICLE{RecherT2010,
  author       = {Recher, Patrik and Trauzettel, Bj{\"o}rn},
  title        = {Mit Tesafilm nach Stockholm},
  journal      = PJ,
  publisher    = WILEYVCH,
  year         = {2010},
  volume       = {9},
  number       = {12},
  pages        = {22-24},
  month        = DEC
}
% url = {http://www.pro-physik.de/Phy/leadArticle.do?laid=13568}

@ARTICLE{Kuzmany2009,
  author       = {Kuzmany, Hans},
  title        = {Der flachste Kohlenstoff},
  journal      = {Physik in unserer Zeit},
  publisher    = WILEYVCH,
  year         = {2009},
  volume       = {40},
  number       = {3},
  pages        = {115},
  month        = MAY
}
% doi = {10.1002/piuz.200990042}
% url = {http://onlinelibrary.wiley.com/doi/10.1002/piuz.200990042/abstract}

@ARTICLE{GrafL1998,
  author       = {Graf, H. and L{\"o}wen, H.},
  title        = {Density jumps across phase transitions in soft-matter systems},
  journal      = PRE,
  publisher    = APS,
  year         = {1998},
  volume       = {57},
  number       = {5},
  pages        = {5744-5753},
  month        = MAY
}
% doi = {10.1103/PhysRevE.57.5744}
% url = {http://adsabs.harvard.edu/abs/1998PhRvE..57.5744G}

@ARTICLE{AlvarezMW2008,
  author       = {Alvarez, C. and Mazars, M. and Weis, J.~J.},
  title        = {Structure and thermodynamics of a ferrofluid bilayer},
  journal      = PRE,
  publisher    = APS,
  year         = {2008},
  volume       = {77},
  number       = {5},
  pages        = {051501},
  month        = MAY
}
% doi = {10.1103/PhysRevE.77.051501}
% url = {http://adsabs.harvard.edu/abs/2008PhRvE..77e1501A}

@ARTICLE{LombaLW2000,
  author       = {Lomba, E. and Lado, F. and Weis, J.~J.},
  title        = {Structure and thermodynamics of a ferrofluid monolayer},
  journal      = PRE,
  publisher    = APS,
  year         = {2000},
  volume       = {61},
  number       = {4},
  pages        = {3838-3849},
  month        = APR
}
% doi = {10.1103/PhysRevE.61.3838}
% url = {http://adsabs.harvard.edu/abs/2000PhRvE..61.3838L}

@ARTICLE{FroltsovBLL2003,
  author       = {Froltsov, V.~A. and Blaak, R. and Likos, C.~N. and L{\"o}wen, H.},
  title        = {Crystal structures of two-dimensional magnetic colloids in tilted external magnetic fields},
  journal      = PRE,
  publisher    = APS,
  year         = {2003},
  volume       = {68},
  number       = {6},
  pages        = {061406},
  month        = DEC
}
% doi = {10.1103/PhysRevE.68.061406}
% url = {http://adsabs.harvard.edu/abs/2003PhRvE..68f1406F}

@ARTICLE{vanTeeffelenLHL2006,
  author       = {van~Teeffelen, S. and Likos, C.~N. and Hoffmann, N. and L{\"o}wen, H.},
  title        = {Density functional theory of freezing for soft interactions in two dimensions},
  journal      = EPL,
  publisher    = EPS,
  year         = {2006},
  volume       = {75},
  number       = {4},
  pages        = {583-589},
  month        = AUG
}
% doi = {10.1209/epl/i2006-10140-7}
% url = {http://adsabs.harvard.edu/abs/2006EL.....75..583V}

@ARTICLE{Barrat1987,
  author       = {Barrat, J.-L.},
  title        = {Role of triple correlations in the freezing of the one-component plasma},
  journal      = EPL,
  publisher    = EPS,
  year         = {1987},
  volume       = {3},
  number       = {5},
  pages        = {523-526},
  month        = MAR
}
% doi = {10.1209/0295-5075/3/5/002}
% url = {http://adsabs.harvard.edu/abs/1987EL......3..523B}

@ARTICLE{ElgetiG2009,
  author       = {Elgeti, J. and Gompper, G.},
  title        = {Self-propelled rods near surfaces},
  journal      = EPL,
  publisher    = EPS,
  year         = {2009},
  volume       = {85},
  number       = {3},
  pages        = {38002},
  month        = FEB
}
% doi = {10.1209/0295-5075/85/38002}
% url = {http://adsabs.harvard.edu/abs/2009EL.....8538002E}

@ARTICLE{BarratHP1988,
  author       = {Barrat, J.-L. and Hansen, J.-P. and Pastore, G.},
  title        = {On the equilibrium structure of dense fluids},
  journal      = MP,
  publisher    = TF,
  year         = {1988},
  volume       = {63},
  number       = {5},
  pages        = {747-767},
  month        = APR
}
% doi = {10.1080/00268978800100541}
% url = {http://adsabs.harvard.edu/abs/1988MolPh..63..747B}

@ARTICLE{HoseinJLEL2009,
  author       = {Hosein, Ian~D. and John, Bettina~S. and Lee, Stephanie~H. and Escobedo, Fernando~A. and Liddell, Chekesha~M.},
  title        = {Rotator and crystalline films via self-assembly of short-bond-length colloidal dimers},
  journal      = JoMC,
  publisher    = RSoC,
  year         = {2009},
  volume       = {19},
  number       = {3},
  pages        = {344-349},
  month        = JAN
}
% doi = {10.1039/B818613H}
% url = {http://dx.doi.org/10.1039/B818613H}

@ARTICLE{HongCLG2006,
  author       = {Hong, L. and Cacciuto, A. and Luijten, E. and Granick, S.},
  title        = {Clusters of charged {J}anus spheres},
  journal      = NL,
  publisher    = ACS,
  year         = {2006},
  volume       = {6},
  number       = {11},
  pages        = {2510-2514},
  month        = NOV
}
% doi = {10.1021/nl061857i}
% url = {http://adsabs.harvard.edu/abs/2006NanoL...6.2510H}

@ARTICLE{HoCSLK2008,
  author       = {Ho, {Chi-Chih} and Chen, {Wen-Shiang} and Shie, {Tzung-Ying} and Lin, {Jiun-Nan} and Kuo, Changshu},
  title        = {Novel fabrication of {J}anus particles from the surfaces of electrospun polymer fibers},
  journal      = LANG,
  publisher    = ACS,
  year         = {2008},
  volume       = {24},
  number       = {11},
  pages        = {5663-5666},
  month        = MAY
}
% doi = {10.1021/la800282j}
% url = {http://pubs.acs.org/doi/abs/10.1021/la800282j}

@ARTICLE{KegelBEP2006,
  author       = {Kegel, Willem~K. and Breed, Dana and Elsesser, Mark and Pine, David~J.},
  title        = {Formation of anisotropic polymer colloids by disparate relaxation times},
  journal      = LANG,
  publisher    = ACS,
  year         = {2006},
  volume       = {22},
  number       = {17},
  pages        = {7135-7136},
  month        = JUL
}
% doi = {10.1021/la061792z}
% url = {http://pubs.acs.org/doi/abs/10.1021/la061792z}

@article{PelaezUPDD2018,
  title={Hydrodynamic fluctuations in quasi-two dimensional diffusion},
  author={Pel{\'a}ez, Ra{\'u}l P and Usabiaga, Florencio Balboa and Panzuela, Sergio and Xiao, Qiyu and Delgado-Buscalioni, Rafael and Donev, Aleksandar},
  journal={Journal of Statistical Mechanics: Theory and Experiment},
  volume={2018},
  number={6},
  pages={063207},
  year={2018},
  publisher={IOP Publishing}
}

@article{StillingerB1962,
  title={Equilibrium statistical mechanics of inhomogeneous fluids},
  author={Stillinger Jr, Frank H and Buff, Frank P},
  journal=JoChemP,
  volume={37},
  number={1},
  pages={1--12},
  year={1962},
  publisher={AIP}
}

@article{MarquesG2004,
  title={Time-dependent density functional theory},
  author={Marques, Miguel AL and Gross, Eberhard KU},
  journal={Annual Review of Physical Chemistry},
  volume={55},
  pages={427--455},
  year={2004},
  publisher={Annual Reviews}
}


@article{KapralC1999,
  title={Mixed quantum-classical dynamics},
  author={Kapral, Raymond and Ciccotti, Giovanni},
  journal=JoChemP,
  volume={110},
  number={18},
  pages={8919--8929},
  year={1999},
  publisher={AIP}
}

@article{NielsenKC2001,
  title={Statistical mechanics of quantum-classical systems},
  author={Nielsen, Steve and Kapral, Raymond and Ciccotti, Giovanni},
  journal={The Journal of Chemical Physics},
  volume={115},
  number={13},
  pages={5805--5815},
  year={2001},
  publisher={AIP}
}


@article{LebowitzP1963,
  title={Asymptotic behavior of the radial distribution function},
  author={Lebowitz, JL and Percus, JK},
  journal={Journal of Mathematical Physics},
  volume={4},
  number={2},
  pages={248--254},
  year={1963},
  publisher={AIP}
}


@article{LebowitzP1963b,
  title={Statistical thermodynamics of nonuniform fluids},
  author={Lebowitz, JL and Percus, JK},
  journal={Journal of Mathematical Physics},
  volume={4},
  number={1},
  pages={116--123},
  year={1963},
  publisher={AIP}
}

@article{GalenkoDL2009,
  title={Phase-field-crystal and {S}wift-{H}ohenberg equations with fast dynamics},
  author={Galenko, Peter and Danilov, Denis and Lebedev, Vladimir},
  journal=PRE,
  volume={79},
  number={5},
  pages={051110},
  year={2009},
  publisher={APS}
}

@article{YangZ2019,
  title={Efficient linear schemes for the nonlocal {C}ahn--{H}illiard equation of phase field models},
  author={Yang, Xiaofeng and Zhao, Jia},
  journal={Computer Physics Communications},
  volume={235},
  pages={234--245},
  year={2019},
  publisher={Elsevier}
}

@article{FehrmanG2019,
  title={Well-posedness of nonlinear diffusion equations with nonlinear, conservative noise},
  author={Fehrman, Benjamin and Gess, Benjamin},
  journal={Archive for Rational Mechanics and Analysis},
  volume={233},
  number={1},
  pages={249--322},
  year={2019},
  publisher={Springer}
}

@article{BabelEL2018,
  title={Impedance resonance in narrow confinement},
  author={Babel, Sonja and Eikerling, Michael and L\"owen, Hartmut},
  journal={Journal of Physical Chemistry C},
  volume={122},
  number={38},
  pages={21724--21734},
  year={2018},
  publisher={ACS Publications}
}

@inproceedings{GoddardNSPK2013,
  title={Inertia and Hydrodynamic Interactions in Dynamical Density Functional Theory},
  author={Goddard, Benjamin D and Nold, Andreas and Savva, Nikos and Pavliotis, Grigorios A and Kalliadasis, Serafim},
  booktitle={Proceedings of the European Conference on Complex Systems 2012},
  pages={999--1004},
  year={2013},
  organization={Springer}
}

@article{JaatinenAEA2009,
  title={Thermodynamics of bcc metals in phase-field-crystal models},
  author={Jaatinen, A and Achim, CV and Elder, KR and Ala-Nissila, T},
  journal=PRE,
  volume={80},
  number={3},
  pages={031602},
  year={2009},
  publisher={APS}
}

@article{Wu2006,
  title={Density functional theory for chemical engineering: From capillarity to soft materials},
  author={Wu, Jianzhong},
  journal={AIChE journal},
  volume={52},
  number={3},
  pages={1169--1193},
  year={2006},
  publisher={Wiley Online Library}
}

@article{SuFYL2019,
  title={Numerical simulation of {S}wift--{H}ohenberg equation by the fourth-order compact scheme},
  author={Su, Jian and Fang, Weiwei and Yu, Qian and Li, Yibao},
  journal={Computational and Applied Mathematics},
  volume={38},
  number={2},
  pages={54},
  year={2019},
  publisher={Springer}
}

@article{YuZWD2019,
  title={From thermodynamics to kinetics: Theoretical study of CO$_2$ dissolving in poly (lactic acid) melt},
  author={Yu, Kesong and Zhou, Hongfu and Wang, Xiangdong and Du, Zhongjie and Mi, Jianguo},
  journal={Journal of Molecular Liquids},
  volume={280},
  pages={97--103},
  year={2019},
  publisher={Elsevier}
}

@article{LipsRM2019,
  title={Single-file transport in periodic potentials: The {B}rownian asymmetric simple exclusion process},
  author={Lips, Dominik and Ryabov, Artem and Maass, Philipp},
  journal={Physical Review E},
  volume={100},
  number={5},
  pages={052121},
  year={2019},
  publisher={APS}
}

@article{Duque2018,
  title={Particle method for phase separation on membranes},
  author={Duque, Daniel},
  journal={Microfluidics and Nanofluidics},
  volume={22},
  number={9},
  pages={95},
  year={2018},
  publisher={Springer}
}

@article{Loewen2015,
  title={History and perspective of the {C}ar--{P}arrinello approach applied to classical systems},
  author={L{\"o}wen, H},
  journal=MP,
  volume={113},
  number={17-18},
  pages={2385--2386},
  year={2015},
  publisher={Taylor \& Francis}
}

@incollection{Rauscher2013,
  title={Dynamics of nanodroplets on structured surfaces},
  author={Rauscher, Markus},
  booktitle={Nanodroplets},
  pages={143--167},
  year={2013},
  publisher={Springer}
}

@article{KocherP2019,
  title={Thermodensity coupling in phase-field-crystal-type models for the study of rapid crystallization},
  author={Kocher, Gabriel and Provatas, Nikolas},
  journal={Physical Review Materials},
  volume={3},
  number={5},
  pages={053804},
  year={2019},
  publisher={APS}
}

@inbook{TothPTG2012,
  title={Phase-Field Crystal Modeling of Homogeneous and Heterogeneous Crystal Nucleation},
  author={T{\'o}th, Gyula I and Pusztai, Tam{\'a}s and Tegze, Gy{\"o}rgy and Gr{\'a}n{\'a}sy, L{\'a}szl{\'o}},
  booktitle={Solidification of Containerless Undercooled Melts},
  pages={113--138},
  year={2012},
  publisher={Wiley Online Library}
}

@article{GaoX2018,
  title={Viscosity effects for ion transport in electrochemical systems},
  author={Gao, Haixia and Xiao, Changming},
  journal={Europhysics Letters},
  volume={124},
  number={5},
  pages={58002},
  year={2018},
  publisher={IOP Publishing}
}

@article{BruttingTdlHS2019,
  title={Superadiabatic Forces via the Acceleration Gradient in Quantum Many-Body Dynamics},
  author={Br{\"u}tting, Moritz and Trepl, Thomas and {de las Heras}, Daniel and Schmidt, Matthias},
  journal={Molecules},
  volume={24},
  number={20},
  pages={3660},
  year={2019},
  publisher={Multidisciplinary Digital Publishing Institute}
}

@Inbook{Rauscher2015,
author={Rauscher, Markus},
editor={Li, Dongqing},
title={Dynamic Density Functional Theory ({DDFT})},
bookTitle={Encyclopedia of Microfluidics and Nanofluidics},
year={2015},
publisher={Springer New York},
address={New York},
pages={693--699}
}

@article{InoueY2017,
  title={Method for Studying Many-Particle Effects on Nonequilibrium Steady States},
  author={Inoue, Masao and Yoshimori, Akira},
  journal={Journal of the Physical Society of Japan},
  volume={86},
  number={7},
  pages={074604},
  year={2017},
  publisher={The Physical Society of Japan}
}

@article{TaramaEL2019,
  title = {Traveling band formation in feedback-driven colloids},
  author = {Tarama, Sonja and Egelhaaf, Stefan U. and L\"owen, Hartmut},
  journal = PRE,
  volume = {100},
  issue = {2},
  pages = {022609},
  numpages = {14},
  year = {2019},
  month = {Aug},
  publisher = {American Physical Society}
}
%   doi = {10.1103/PhysRevE.100.022609},
%  url = {https://link.aps.org/doi/10.1103/PhysRevE.100.022609}


@article{SunKK2019,
  title={Hydrodynamic Stress Tensor in Inhomogeneous Colloidal Suspensions: an {Irving-Kirkwood} Extension},
  author={Sun, Zong-Li and Kang, Yan-Shuang and Kang, Yan-Mei},
  journal={Communications in Theoretical Physics},
  volume={71},
  number={7},
  pages={876},
  year={2019},
  publisher={IOP Publishing}
}

@article{RussoPDYCK2019,
  title={A Finite-Volume Method for Fluctuating Dynamical Density Functional Theory},
  author={Russo, Antonio and Perez, Sergio P and Dur{\'a}n-Olivencia, Miguel A and Yatsyshin, Peter and Carrillo, Jos{\'e} A and Kalliadasis, Serafim},
  journal={preprint},
  volume={arXiv:1910.05067},
  year={2019}
}

@article{MijalkovMWV2016,
  title={Engineering sensorial delay to control phototaxis and emergent collective behaviors},
  author={Mijalkov, Mite and McDaniel, Austin and Wehr, Jan and Volpe, Giovanni},
  journal={Physical Review X},
  volume={6},
  number={1},
  pages={011008},
  year={2016},
  publisher={APS}
}

@article{KhadkaHVC2018,
  title={Active particles bound by information flows},
  author={Khadka, Utsab and Holubec, Viktor and Yang, Haw and Cichos, Frank},
  journal={Nature Communications},
  volume={9},
  number={1},
  pages={3864},
  year={2018},
  publisher={Nature Publishing Group}
}

@article{LeymanOWV2018,
  title={Tuning phototactic robots with sensorial delays},
  author={Leyman, Maximilian and Ogemark, Freddie and Wehr, Jan and Volpe, Giovanni},
  journal=PRE,
  volume={98},
  number={5},
  pages={052606},
  year={2018},
  publisher={APS}
}

@article{MonchoGAAD2019,
  title={Nonequilibrium Uptake Kinetics of Molecular Cargo into Hollow Hydrogels Tuned by Electrosteric Interactions},
  author={Moncho-Jord{\'a}, Arturo and Germ{\'a}n-Bellod, Alicia and Angioletti-Uberti, Stefano and Adroher-Ben{\'\i}tez, Irene and Dzubiella, Joachim},
  journal={ACS Nano},
  volume={13},
  number={2},
  pages={1603--1616},
  year={2019},
  publisher={ACS Publications}
}

@article{Frusawa2014,
  title={Emerging quasi-0D states at vanishing total entropy of the 1D hard sphere system: A coarse-grained similarity to the car parking problem},
  author={Frusawa, Hiroshi},
  journal={Physics Letters A},
  volume={378},
  number={26-27},
  pages={1780--1786},
  year={2014},
  publisher={Elsevier}
}

@article{LiLXK2019,
  title={An efficient linear second order unconditionally stable direct discretization method for the phase-field crystal equation on surfaces},
  author={Li, Yibao and Luo, Chaojun and Xia, Binhu and Kim, Junseok},
  journal={Applied Mathematical Modelling},
  volume={67},
  pages={477--490},
  year={2019},
  publisher={Elsevier}
}

@article{VuijkBS2019,
  title={Effect of anisotropic diffusion on spinodal decomposition},
  author={Vuijk, Hidde Derk and Brader, Joseph Michael and Sharma, Abhinav},
  journal=SM,
  volume={15},
  number={6},
  pages={1319--1326},
  year={2019},
  publisher={Royal Society of Chemistry}
}

@article{AraujoZKT2017,
  title={Dynamic design of spatial patterns of colloidal suspensions},
  author={Ara{\'u}jo, NAM and Zezyulin, DA and Konotop, VV and Telo da Gama, MM},
  journal={Langmuir},
  volume={33},
  number={42},
  pages={11698--11702},
  year={2017},
  publisher={ACS Publications}
}

@article{GonzalezMV2017,
  title={Dynamical properties of heterogeneous nucleation of parallel hard squares},
  author={Gonz{\'a}lez-Pinto, Miguel and Mart{\'\i}nez-Rat{\'o}n, Yuri and Velasco, Enrique},
  journal=SM,
  volume={13},
  number={48},
  pages={9246--9258},
  year={2017},
  publisher={Royal Society of Chemistry}
}

@article{CamargodlTDBCE2019,
  title={Boundary conditions derived from a microscopic theory of hydrodynamics near solids},
  author={Camargo, Diego and de la Torre, JA and Delgado-Buscalioni, Rafael and Chejne, Farid and Espa{\~n}ol, Pep},
  journal=JoChemP,
  volume={150},
  number={14},
  pages={144104},
  year={2019},
  publisher={AIP Publishing}
}

@article{KawasakiG1973,
  title={Theory of nonlinear transport processes: nonlinear shear viscosity and normal stress effects},
  author={Kawasaki, Kyozi and Gunton, James D},
  journal=PRA,
  volume={8},
  number={4},
  pages={2048},
  year={1973},
  publisher={APS}
}

@article{KonarovskyiLvR2019,
  title={{Dean-Kawasaki} dynamics: ill-posedness vs. triviality},
  author={Konarovskyi, Vitalii and Lehmann, Tobias and von Renesse, Max-K and others},
  journal={Electronic Communications in Probability},
  volume={24},
  year={2019},
  publisher={The Institute of Mathematical Statistics and the Bernoulli Society}
}

@incollection{Loewen2017,
  title={Dynamical density functional theory for {B}rownian dynamics of colloidal particles},
  author={L{\"o}wen, Hartmut},
  booktitle={Variational Methods in Molecular Modeling},
  pages={255--284},
  year={2017},
  publisher={Springer}
}

@INBOOK{Loewen2010b,
  author       = {L{\"o}wen, H.},
  title        = {Understanding Soft Condensed Matter via Modeling and Computation},
  chapter      = {2: Applications of Density Functional Theory in Soft Condensed Matter},
  publisher    = WSP,
  year         = {2010},
  volume       = {3},
  series       = {Series in Soft Condensed Matter},
  address      = SIN,
  editor       = {Wenbing Hu and An-Chang Shi},
  month        = DEC,
  pages        = {9-45},
  isbn         = {9-814-29558-2}
}

@INCOLLECTION{PleinerB1996,
  author       = {Pleiner, Harald and Brand, Helmut~R.},
  title        = {Hydrodynamics and Electrohydrodynamics of Liquid Crystals},
  booktitle    = {Pattern Formation in Liquid Crystals},
  publisher    = SPR,
  year         = {1996},
  editor       = {Buka, Agnes and Kramer, Lorenz},
  volume       = {9},
  series       = {Partially Ordered Systems},
  type         = {chapter},
  chapter      = {2},
  pages        = {15-67},
  address      = NY,
  edition      = {1},
  month        = MAY,
  isbn         = {0-387-94604-7}
}

@article{JaatinenA2010,
  title={Eighth-order phase-field-crystal model for two-dimensional crystallization},
  author={Jaatinen, A and Ala-Nissila, T},
  journal=PRE,
  volume={82},
  number={6},
  pages={061602},
  year={2010},
  publisher={APS}
}

@article{RamirezGonzalezM2010,
  title={General nonequilibrium theory of colloid dynamics},
  author={Ram{\'\i}rez-Gonz{\'a}lez, Pedro and Medina-Noyola, Magdaleno},
  journal=PRE,
  volume={82},
  number={6},
  pages={061503},
  year={2010},
  publisher={APS}
}

@article{EdisonM2009,
  title={Modeling relaxation processes for fluids in porous materials using dynamic mean field theory: an application to partial wetting},
  author={Edison, John R and Monson, Peter A},
  journal={Journal of Low Temperature Physics},
  volume={157},
  number={3-4},
  pages={395--409},
  year={2009},
  publisher={Springer}
}

@article{Rauscher2010,
  title={{DDFT for B}rownian particles and hydrodynamics},
  author={Rauscher, Markus},
  journal=JoP:CM,
  volume={22},
  number={36},
  pages={364109},
  year={2010},
  publisher={IOP Publishing}
}

@article{DeDominicisM1964,
  title={Stationary entropy principle and renormalization in normal and superfluid systems. I. Algebraic formulation},
  author={De Dominicis, Cyrano and Martin, Paul C},
  journal={Journal of Mathematical Physics},
  volume={5},
  number={1},
  pages={14--30},
  year={1964},
  publisher={AIP}
}

@article{EvansORK2016,
  title={New developments in classical density functional theory},
  author={Evans, Robert and Oettel, Martin and Roth, Roland and Kahl, Gerhard},
  journal=JoP:CM,
  volume={28},
  number={24},
  pages={240401},
  year={2016},
  publisher={IOP Publishing}
}

@article{MenzelOL2014,
  title={Active crystals and their stability},
  author={Menzel, Andreas M and Ohta, Takao and L{\"o}wen, Hartmut},
  journal=PRE,
  volume={89},
  number={2},
  pages={022301},
  year={2014},
  publisher={APS}
}

@article{RexLL2005,
  title={Soft colloids driven and sheared by traveling wave fields},
  author={Rex, M and L{\"o}wen, H and Likos, CN},
  journal=PRE,
  volume={72},
  number={2},
  pages={021404},
  year={2005},
  publisher={APS}
}

@article{BrownUM2009,
	volume = {40},
	title = {Boltzmann's {H-}Theorem, its Discontents, and the Birth of Statistical Mechanics},
	author = {Brown, Harvey R. and Myrvold, Wayne and Uffink, Jos},
	number = {2},
	pages = {174--191},
	year = {2009},
	journal = {Studies in History and Philosophy of Modern Physics}
}

@article{Chavanis2011,
  title={Brownian particles with long-and short-range interactions},
  author={Chavanis, Pierre-Henri},
  journal={Physica A: Statistical Mechanics and its Applications},
  volume={390},
  number={9},
  pages={1546-1574},
  year={2011},
  publisher={Elsevier}
}

@ARTICLE{JaatinenAN2010,
  author       = {Jaatinen, A. and {Ala-Nissila}, T.},
  title        = {Extended phase diagram of the three-dimensional phase field crystal model},
  journal      = JoP:CM,
  publisher    = IoPP,
  year         = {2010},
  volume       = {22},
  number       = {20},
  pages        = {205402},
  month        = MAY
}
% doi = {10.1088/0953-8984/22/20/205402}
% url = {http://adsabs.harvard.edu/abs/2010JPCM...22t5402J}

@ARTICLE{BierHD2005,
  author       = {Bier, M. and Harnau, L. and Dietrich, S.},
  title        = {Free isotropic-nematic interfaces in fluids of charged platelike colloids},
  journal      = JoChemP,
  publisher    = AIoP,
  year         = {2005},
  volume       = {123},
  number       = {11},
  pages        = {114906},
  month        = SEP
}
% doi = {10.1063/1.2036975}
% url = {http://adsabs.harvard.edu/abs/2005JChPh.123k4906B}

@ARTICLE{PeruaniDB2006,
  author       = {Peruani, F. and Deutsch, A. and B{\"a}r, M.},
  title        = {Nonequilibrium clustering of self-propelled rods},
  journal      = PRE,
  publisher    = APS,
  year         = {2006},
  volume       = {74},
  number       = {3},
  pages        = {030904(R)},
  month        = SEP
}
% doi = {10.1103/PhysRevE.74.030904}
% url = {http://adsabs.harvard.edu/abs/2006PhRvE..74c0904P}

@ARTICLE{RennerLB1995,
  author       = {Renner, C. and L{\"o}wen, H. and Barrat, J.-L.},
  title        = {Orientational glass transition in a rotator model},
  journal      = PRE,
  publisher    = APS,
  year         = {1995},
  volume       = {52},
  number       = {5},
  pages        = {5091-5099},
  month        = NOV
}
% doi = {10.1103/PhysRevE.52.5091}
% url = {http://adsabs.harvard.edu/abs/1995PhRvE..52.5091R}

@ARTICLE{deGennes1973,
  author       = {de~Gennes, P.-G.},
  title        = {Some remarks on polymorphism of smectics},
  journal      = MCaLC,
  publisher    = TF,
  year         = {1973},
  volume       = {21},
  number       = {1-2},
  pages        = {49-76},
  month        = JAN
}

@BOOK{ChaikinL1995,
  author       = {Chaikin, P.~M. and Lubensky, T.~C.},
  title        = {Principles of Condensed Matter Physics},
  publisher    = CUP,
  year         = {1995},
  address      = CAM,
  edition      = {1st},
  month        = OCT,
  pages        = {699},
  isbn         = {0-521-79450-1}
}
% ISBN13 = {978-0521794503}

@ARTICLE{LoewenBW1990,
  author       = {L{\"o}wen, H. and Beier, T. and Wagner, H.},
  title        = {Multiple order parameter theory of surface melting: a van der {W}aals approach},
  journal      = ZfPB:CM,
  publisher    = SPR,
  year         = {1990},
  volume       = {79},
  number       = {1},
  pages        = {109-118},
  month        = FEB
}
% doi = {10.1007/BF01387832}
% url = {http://adsabs.harvard.edu/abs/1990ZPhyB..79..109L}

@ARTICLE{Emmerich2008,
  author       = {Emmerich, H.},
  title        = {Advances of and by phase-field modeling in condensed-matter physics},
  journal      = AiP,
  publisher    = TF,
  year         = {2008},
  volume       = {57},
  number       = {1},
  pages        = {1-87},
  month        = JAN
}
% doi = {10.1080/00018730701822522}
% url = {http://adsabs.harvard.edu/abs/2008AdPhy..57....1E}

@ARTICLE{Roth2010,
  author       = {Roth, R.},
  title        = {Fundamental measure theory for hard-sphere mixtures: a review},
  journal      = JoP:CM,
  publisher    = IoPP,
  year         = {2010},
  volume       = {22},
  number       = {6},
  pages        = {063102},
  month        = FEB
}
% doi = {10.1088/0953-8984/22/6/063102}
% url = {http://adsabs.harvard.edu/abs/2010JPCM...22f3102R}

@ARTICLE{MukherjeePB2002,
  author       = {Mukherjee, P.~K. and Pleiner, H. and Brand, H.~R.},
  title        = {Landau model of the smectic {C}-isotropic phase transition},
  journal      = JoChemP,
  publisher    = AIoP,
  year         = {2002},
  volume       = {117},
  number       = {16},
  pages        = {7788-7792},
  month        = OCT
}
% doi = {10.1063/1.1509055}
% url = {http://adsabs.harvard.edu/abs/2002JChPh.117.7788M}

@ARTICLE{MukherjeePB2005,
  author       = {Mukherjee, P.~K. and Pleiner, H. and Brand, H.~R.},
  title        = {A phenomenological theory of the isotropic to chiral smectic-C phase transition},
  journal      = EPJE,
  publisher    = EDPS,
  year         = {2005},
  volume       = {17},
  number       = {4},
  pages        = {501-506},
  month        = AUG
}
% doi = {10.1140/epje/i2004-10153-6}
% url = {http://adsabs.harvard.edu/abs/2005EPJE...17..501M}

@ARTICLE{PleinerB1989,
  author       = {Pleiner, H. and Brand, H.~R.},
  title        = {Spontaneous splay phases in polar nematic liquid crystals},
  journal      = EPL,
  publisher    = EPS,
  year         = {1989},
  volume       = {9},
  number       = {3},
  pages        = {243-249},
  month        = JUN
}
% doi = {10.1209/0295-5075/9/3/010}
% url = {http://adsabs.harvard.edu/abs/1989EL......9..243P}

@ARTICLE{WatanabeHTWM1998,
  author       = {Watanabe, J. and Hirose, Y. and Tokita, M. and Watanabe, T. and Miyata, S.},
  title        = {Polar structure in polypeptide cholesteric liquid crystals evidenced from observation of second-harmonic generation due to the helicoidal cavity effect},
  journal      = MAC,
  publisher    = ACS,
  year         = {1998},
  volume       = {31},
  number       = {17},
  pages        = {5937-5939},
  month        = AUG
}
% doi = {10.1021/ma980302+}
% url = {http://adsabs.harvard.edu/abs/1998MaMol..31.5937W}

@ARTICLE{TaguchiYKTW2009,
  author       = {Taguchi, Yoshiaki and Yen, {Chu-Chun} and Kang, Sungmin and Tokita, Masatoshi and Watanabe, Junji},
  title        = {Unusual Transformation of the Mechanically Induced Monodomain State to the Polydomain One in Polar Nematic Liquid Crystals of Aromatic Polyesters},
  journal      = JoPChemB,
  publisher    = ACS,
  year         = {2009},
  volume       = {113},
  number       = {16},
  pages        = {5341-5344},
  month        = MAR
}
% doi = {10.1021/jp900817v}
% url = {http://pubs.acs.org/doi/abs/10.1021/jp900817v}

@ARTICLE{ZimmermannPLB1985,
  author       = {Zimmermann, H. and Poupko, R. and Luz, Z. and Billard, J.},
  title        = {Pyramidic Mesophases},
  journal      = ZfNTA,
  publisher    = VdZfN,
  year         = {1985},
  volume       = {40},
  number       = {2},
  pages        = {149-160},
  month        = FEB
}
% doi = {}
% url = {}

@ARTICLE{HsiungRSSCK1987,
  author       = {Hsiung, H. and Rasing, T. and Shen, Y.~R. and Shvartsman, F.~P. and Cabrera, I.~R. and Krongauz, V.~A.},
  title        = {Polar ordering of quasiliquid crystals--An optical second harmonic generation study},
  journal      = JoChemP,

  publisher    = AIoP,
  year         = {1987},
  volume       = {87},
  number       = {5},
  pages        = {3127-3130},
  month        = SEP
}
% doi = {10.1063/1.453050}
% url = {http://adsabs.harvard.edu/abs/1987JChPh..87.3127H}

@ARTICLE{BrandCP1992,
  author       = {Brand, H.~R. and Cladis, P.~E. and Pleiner, H.},
  title        = {Symmetry and defects in the CM phase of polymeric liquid crystals},
  journal      = MAC,
  publisher    = ACS,
  year         = {1992},
  volume       = {25},
  number       = {26},
  pages        = {7223-7226},
  month        = DEC
}
% doi = {10.1021/ma00052a025}
% url = {http://adsabs.harvard.edu/abs/1992MaMol..25.7223B}

@ARTICLE{NioriSWFT1996,
  author       = {Niori, T. and Sekine, T. and Watanabe, J. and Furukawa, T. and Takezoe, H.},
  title        = {Distinct ferroelectric smectic liquid crystals consisting of banana shaped achiral molecules},
  journal      = JoMC,
  publisher    = RSCP,
  year         = {1996},
  volume       = {6},
  number       = {7},
  pages        = {1231-1233},
  month        = JAN
}
% doi = {10.1039/JM9960601231}
% url = {http://pubs.rsc.org/en/Content/ArticleLanding/1996/JM/jm9960601231}

@ARTICLE{LinkNSMCKW1997,
  author       = {Link, D.~R. and Natale, G. and Shao, R. and Maclennan, J.~E. and Clark, N.~A. and Korblova, E. and Walba, D.~M.},
  title        = {Spontaneous Formation of Macroscopic Chiral Domains in a Fluid Smectic Phase of Achiral Molecules},
  journal      = SCI,
  publisher    = AAftAoS,
  year         = {1997},
  volume       = {278},
  number       = {5345},
  pages        = {1924-1927},
  month        = DEC
}
% doi = {10.1126/science.278.5345.1924}
% url = {http://adsabs.harvard.edu/abs/1997Sci...278.1924L}

@ARTICLE{BrandCP1998,
  author       = {Brand, H.~R. and Cladis, P.~E. and Pleiner, H.},
  title        = {Macroscopic properties of smectic liquid crystals},
  journal      = EPJB,
  publisher    = EDPS,
  year         = {1998},
  volume       = {6},
  number       = {3},
  pages        = {347-353},
  month        = NOV
}
% doi = {10.1007/s100510050560}
% url = {http://adsabs.harvard.edu/abs/1998EPJB....6..347B}

@ARTICLE{PleinerCB2006,
  author       = {Pleiner, H. and Cladis, P.~E. and Brand, H.~R.},
  title        = {Splay-bend textures involving tetrahedratic order},
  journal      = EPJE,
  publisher    = EDPS,
  year         = {2006},
  volume       = {20},
  number       = {3},
  pages        = {257-266},
  month        = JUL
}
% doi = {10.1140/epje/i2005-10129-0}
% url = {http://adsabs.harvard.edu/abs/2006EPJE...20..257P}

@ARTICLE{BrandCP2003,
  author       = {Brand, H.~R. and Cladis, P.~E. and Pleiner, H.},
  title        = {Macroscopic properties of smectic C-G liquid crystals},
  journal      = EPJB,
  publisher    = EDPS,
  year         = {2003},
  volume       = {31},
  number       = {1},
  pages        = {147-147},
  month        = JAN
}
% doi = {10.1140/epjb/e2003-0018-6}
% url = {}

@ARTICLE{PelzlDW1999,
  author       = {Pelzl, Gerhard and Diele, Siegmar and Weissflog, Wolfgang},
  title        = {Banana-Shaped Compounds--A New Field of Liquid Crystals},
  journal      = AdvM,
  publisher    = WILEYVCH,
  year         = {1999},
  volume       = {11},
  number       = {9},
  pages        = {707-724},
  month        = JUN
}
% doi = {10.1002/(SICI)1521-4095(199906)11:9<707::AID-ADMA707>3.0.CO;2-D}
% url = {http://onlinelibrary.wiley.com/doi/10.1002/%28SICI%291521-4095%28199906%2911:9%3C707::AID-ADMA707%3E3.0.CO;2-D/abstract}

@ARTICLE{BrandCP2000,
  author       = {Brand, H.~R. and Cladis, P.~E. and Pleiner, H.},
  title        = {Polar biaxial liquid crystalline phases with fluidity in two and three spatial dimensions},
  journal      = IJoES,
  publisher    = ELS,
  year         = {2000},
  volume       = {38},
  number       = {9-10},
  pages        = {1099-1112},
  month        = JUN
}
% doi = {doi:10.1016/S0020-7225(99)00110-X}
% url = {http://www.sciencedirect.com/science?_ob=ArticleURL&_udi=B6V32-4037BS9-V&_user=10&_coverDate=06%2F01%2F2000&_rdoc=1&_fmt=high&_orig=gateway&_origin=gateway&_sort=d&_docanchor=&view=c&_rerunOrigin=scholar.google&_acct=C000050221&_version=1&_urlVersion=0&_userid=10&md5=39174f71b24c883997c4f6e175bd3700&searchtype=a}

@ARTICLE{NioriYY2004,
  author       = {Niori, T. and Yamamoto, J. and Yokoyama, H.},
  title        = {Dynamics of the nematic phase formed by achiral banana-shaped materials},
  journal      = MCaLC,
  publisher    = TF,
  year         = {2004},
  volume       = {409},
  number       = {1},
  pages        = {475-482},
  month        = FEB
}
% doi = {10.1080/15421400490434072}
% url = {}

@ARTICLE{ShenDPWT1999,
  author       = {Shen, Dong and Diele, Siegmar and Pelzl, Gerhard and Wirth, Ina and Tschierske, Carsten},
  title        = {Designing banana-shaped liquid crystals without Schiff's base units: m-terphenyls, 2,6-diphenylpyridines and V-shaped tolane derivatives},
  journal      = JoMC,
  publisher    = RSCP,
  year         = {1999},
  volume       = {9},
  number       = {3},
  pages        = {661-672},
  month        = MAR
}
% doi = {10.1039/A808275H}
% url = {http://pubs.rsc.org/en/Content/ArticleLanding/1999/JM/a808275h}

@ARTICLE{WeissflogNDPDEK2001,
  author       = {Weissflog, Wolfgang and N{\'a}dasi, Hajnalka and Dunemann, Ulrike and Pelzl, Gerhard and Diele, Siegmar and Eremin, Alexei and Kresse, Horst},
  title        = {Influence of lateral substituents on the mesophase behaviour of banana-shaped mesogens},
  journal      = JoMC,
  publisher    = RSCP,
  year         = {2001},
  volume       = {11},
  number       = {11},
  pages        = {2748-2758},
  month        = NOV
}
% doi = {10.1039/B104098G}
% url = {http://pubs.rsc.org/en/Content/ArticleLanding/2001/JM/b104098g}

@article{Kawasaki1995,
  title={Mode coupling in critical phenomena and supercooled liquids},
  author={Kawasaki, Kyozi},
  journal={Transport Theory and Statistical Physics},
  volume={24},
  number={6-8},
  pages={755--779},
  year={1995},
  publisher={Taylor \& Francis}
}

@article{Kawasaki1994,
  title={Stochastic model of slow dynamics in supercooled liquids and dense colloidal suspensions},
  author={Kawasaki, Kyozi},
  journal={Physica A: Statistical Mechanics and its Applications},
  volume={208},
  number={1},
  pages={35--64},
  year={1994},
  publisher={Elsevier}
}

@article{KawasakiK1993,
  title={Relaxation and growth of concentration fluctuations in binary fluids and polymer blends},
  author={Kawasaki, Kyozi and Koga, Tsuyoshi},
  journal={Physica A: Statistical Mechanics and its Applications},
  volume={201},
  number={1-3},
  pages={115--128},
  year={1993},
  publisher={Elsevier}
}

@article{KogaK1993,
  title={Late stage dynamics of spinodal decomposition in binary fluid mixtures},
  author={Koga, Tsuyoshi and Kawasaki, Kyozi},
  journal={Physica A: Statistical Mechanics and its Applications},
  volume={196},
  number={3},
  pages={389--415},
  year={1993},
  publisher={Elsevier}
}

@article{AlaimoV2018,
  title={Microscopic field-theoretical approach for mixtures of active and passive particles},
  author={Alaimo, Francesco and Voigt, Axel},
  journal=PRE,
  volume={98},
  number={3},
  pages={032605},
  year={2018},
  publisher={APS}
}

@article{AlaimoPV2016,
  title={A microscopic field theoretical approach for active systems},
  author={Alaimo, Francesco and Praetorius, Simon and Voigt, Axel},
  journal=NJoP,
  volume={18},
  number={8},
  pages={083008},
  year={2016},
  publisher={IOP Publishing}
}

@article{KirkpatrickT1989,
  title={Random solutions from a regular density functional {H}amiltonian: a static and dynamical theory for the structural glass transition},
  author={Kirkpatrick, TR and Thirumalai, D},
  journal={Journal of Physics A: Mathematical and General},
  volume={22},
  number={5},
  pages={L149},
  year={1989},
  publisher={IOP Publishing}
}

@article{Bagchi1987,
  title={Stability of a supercooled liquid to periodic density waves and dynamics of freezing},
  author={Bagchi, Biman},
  journal={Physica A: Statistical Mechanics and its Applications},
  volume={145},
  number={1-2},
  pages={273-289},
  year={1987},
  publisher={Elsevier}
}

@article{Munakata1977,
  title={Liquid Instability and Freezing—Reductive Perturbation Approach—},
  author={Munakata, Toyonori},
  journal={Journal of the Physical Society of Japan},
  volume={43},
  number={5},
  pages={1723--1728},
  year={1977},
  publisher={The Physical Society of Japan}
}

@article{Munakata1989,
author = {Munakata, Toyonori},
title = {A Dynamical Extension of the Density Functional Theory},
journal = {Journal of the Physical Society of Japan},
volume = {58},
number = {7},
pages = {2434-2438},
year = {1989},
}
%doi={https://doi.org/10.1143/JPSJ.58.2434},

@article{Boltzmann1872,
  author={Boltzmann, Ludwig},
  title ={Weitere {S}tudien \"uber das {W}\"armegleichgewicht unter {G}asmolek\"ulen},
  journal = {Sitzungberichte der Akademie der Wissenschaften zu Wien, mathematisch-naturwissenschaftliche Klasse, 66},
  year={1872},
  pages={275--370}
}

@article{Kawasaki1998,
  title={Microscopic analyses of the dynamical density functional equation of dense fluids},
  author={Kawasaki, Kyozi},
  journal={Journal of Statistical Physics},
  volume={93},
  number={3-4},
  pages={527--546},
  year={1998},
  publisher={Springer}
}

@article{KawasakiM1997,
  title={Path integral formulation of dynamical density functional equation for dense fluids},
  author={Kawasaki, Kyozi and Miyazima, Sasuke},
  journal={Zeitschrift f{\"u}r Physik B Condensed Matter},
  volume={103},
  number={3},
  pages={423-431},
  year={1997},
  publisher={Springer}
}

@ARTICLE{BrandK1986,
  author       = {Brand, H.~R. and Kawasaki, K.},
  title        = {Gradient free energy of nematic liquid crystals with topological defects},
  journal      = JoPC,
  publisher    = IoPP,
  year         = {1986},
  volume       = {19},
  number       = {7},
  pages        = {937-942},
  month        = MAR
}
% doi = {10.1088/0022-3719/19/7/006}
% url = {http://adsabs.harvard.edu/abs/1986JPhC...19..937B}

@ARTICLE{BrandK1994,
  author       = {{Brand}, H.~R. and {Kawasaki}, K.},
  title        = {Mode coupling theory of the isotropic-nematic transition in sidechain liquid crystalline polymers},
  journal      = JdPII,
  publisher    = EDPS,
  year         = {1994},
  volume       = {4},
  number       = {4},
  pages        = {543-548},
  month        = APR
}
% doi = {10.1051/jp2:1994144}
% url = {http://adsabs.harvard.edu/abs/1994JPhy2...4..543B}

@ARTICLE{KawasakiB1985,
  author       = {Kawasaki, K. and Brand, H.~R.},
  title        = {Gauge theory of continuous media with topological defects: uniaxial nematic liquid crystals and superfluid $^{4}$He},
  journal      = AdPD,
  publisher    = WILEYVCH,
  year         = {1985},
  volume       = {160},
  number       = {2},
  pages        = {420-440},
  month        = APR
}
% doi = {10.1016/0003-4916(85)90151-4}
% url = {http://adsabs.harvard.edu/abs/1985AnPhy.160..420K}

@ARTICLE{KawasakiB1994,
  author       = {{Kawasaki}, K. and {Brand}, H.~R.},
  title        = {Life-time renormalization in critical dynamics as applied to the isotropic-nematic transition in sidechain liquid crystalline polymers},
  journal      = PA,
  publisher    = ELS,
  year         = {1994},
  volume       = {208},
  number       = {3-4},
  pages        = {407-422},
  month        = AUG
}
% doi = {10.1016/0378-4371(94)00059-X}
% url = {http://adsabs.harvard.edu/abs/1994PhyA..208..407K}

@ARTICLE{Kawasaki1970,
  author       = {{Kawasaki}, K.},
  title        = {Kinetic equations and time correlation functions of critical fluctuations},
  journal      = AoP,
  publisher    = ELS,
  year         = {1970},
  volume       = {61},
  number       = {1},
  pages        = {1-56},
  month        = NOV
}
% doi = {10.1016/0003-4916(70)90375-1}
% url = {http://adsabs.harvard.edu/abs/1970AnPhy..61....1K}

@ARTICLE{Kawasaki1974,
  author       = {{Kawasaki}, K.},
  title        = {A macroscopic equation of motion for the {TDGL} model},
  journal      = PoTP,
  publisher    = YHaKU,
  year         = {1974},
  volume       = {52},
  number       = {1},
  pages        = {359-360},
  month        = JUL
}
% doi = {10.1143/PTP.52.359}
% url = {http://adsabs.harvard.edu/abs/1974PThPh..52..359K}

@ARTICLE{Kawasaki1975,
  author       = {{Kawasaki}, K.},
  title        = {Multiplicative renormalization of life time},
  journal      = PLA,
  publisher    = ELS,
  year         = {1975},
  volume       = {54},
  number       = {2},
  pages        = {131-132},
  month        = AUG
}
% doi = {10.1016/0375-9601(75)90836-1}
% url = {http://adsabs.harvard.edu/abs/1975PhLA...54..131K}

@ARTICLE{ChoiKPTNW1998,
  author       = {Choi, S.-W. and Kinoshita, Y. and Park, B. and Takezoe, H. and Niori, T. and Watanabe, J.},
  title        = {Second-Harmonic Generation in Achiral Bent-Shaped Liquid Crystals},
  journal      = JJoAP,
  publisher    = JSoAP,
  year         = {1998},
  volume       = {37},
  number       = {6A},
  pages        = {3408-3411},
  month        = JUN
}
% doi = {10.1143/JJAP.37.3408}
% url = {http://adsabs.harvard.edu/abs/1998JaJAP..37.3408C}

@ARTICLE{YenTPTW2006,
  author       = {Yen, C.-C. and Tokita, M. and Park, B. and Takezoe, H. and Watanabe, J.},
  title        = {Spontaneous Organization of Helical Polypeptide Molecules into Polar Packing Structure},
  journal      = MAC,
  publisher    = ACS,
  year         = {2006},
  volume       = {39},
  number       = {4},
  pages        = {1313-1315},
  month        = FEB
}
% doi = {10.1021/ma052424p}
% url = {http://adsabs.harvard.edu/abs/2006MaMol..39.1313Y}

@ARTICLE{KoikeEtAl2007,
  author       = {Koike, M. and Yen, C.-C. and Yuqing, L. and Tsuchiya, H. and Tokita, M. and Kawauchi, S. and Takezoe, H. and Watanabe, J.},
  title        = {Unusual Nematic Liquid Crystal with PolarCsSymmetry Formed from Aromatic Polyesters with Head-Tail Character},
  journal      = MAC,
  publisher    = ACS,
  year         = {2007},
  volume       = {40},
  number       = {7},
  pages        = {2524-2531},
  month        = APR
}
% doi = {10.1021/ma0626214}
% url = {http://adsabs.harvard.edu/abs/2007MaMol..40.2524K}

@ARTICLE{FrancescangeliS2010,
  author       = {Francescangeli, Oriano and Samulski, Edward~T.},
  title        = {Insights into the cybotactic nematic phase of bent-core molecules},
  journal      = SM,
  publisher    = RSoC,
  year         = {2010},
  volume       = {6},
  number       = {11},
  pages        = {2413-2420},
  month        = MAR
}
% doi = {10.1039/C003310C}
% url = {http://dx.doi.org/10.1039/C003310C}

@ARTICLE{FrancescangeliEtAl2009,
  author       = {Francescangeli, Oriano and Stanic, Vesna and Torgova, {Sofia I.} and Strigazzi, Alfredo and Scaramuzza, Nicola and 
Ferrero, Claudio and Dolbnya, Igor~P. and Weiss, Thomas~M. and Berardi, Roberto and Muccioli, Luca and Orlandi, Silvia and Zannoni, Claudio},
  title        = {Ferroelectric response and induced biaxiality in the nematic phase of bent-core mesogens},
  journal      = AFM,
  publisher    = WILEYVCH,
  year         = {2009},
  volume       = {19},
  number       = {16},
  pages        = {2592-2600},
  month        = AUG
}
% doi = {10.1002/adfm.200801865}
% url = {http://dx.doi.org/10.1002/adfm.200801865}

@ARTICLE{BrandPZ2006,
  author       = {Brand, H.~R. and Pleiner, H. and Ziebert, F.},
  title        = {Macroscopic dynamics of polar nematic liquid crystals},
  journal      = PRE,
  publisher    = APS,
  year         = {2006},
  volume       = {74},
  number       = {2},
  pages        = {021713},
  month        = AUG
}
% doi = {10.1103/PhysRevE.74.021713}
% url = {http://adsabs.harvard.edu/abs/2006PhRvE..74b1713B}

@ARTICLE{BrandCP2009,
  author       = {Brand, H.~R. and Cladis, P.~E. and Pleiner, H.},
  title        = {Reversible macroscopic dynamics of polar nematic liquid crystals: reversible currents and their experimental consequences},
  journal      = PRE,
  publisher    = APS,
  year         = {2009},
  volume       = {79},
  number       = {3},
  pages        = {032701},
  month        = MAR
}
% doi = {10.1103/PhysRevE.79.032701}
% url = {http://adsabs.harvard.edu/abs/2009PhRvE..79c2701B}

@article{ArcherRRS2019,
  title={Deriving phase field crystal theory from dynamical density functional theory: consequences of the approximations},
  author={Archer, Andrew J and Ratliff, Daniel J and Rucklidge, Alastair M and Subramanian, Priya},
  journal=PRE,
  volume={100},
  number={2},
  pages={022140},
  year={2019},
  publisher={APS}
}

@ARTICLE{PleinerB1980,
  author       = {Pleiner, H. and Brand, H.~R.},
  title        = {On the influence of higher order gradient terms on the hydrodynamics of liquid crystals},
  journal      = JdPL,
  publisher    = EDPS,
  year         = {1980},
  volume       = {41},
  number       = {20},
  pages        = {491-493},
  month        = OCT
}
% doi = {10.1051/jphyslet:019800041020049100}
% url = {http://jphyslet.journaldephysique.org/index.php?option=com_article&access=standard&Itemid=129&url=/articles/jphyslet/abs/1980/20/jphyslet_1980__41_20_491_0/jphyslet_1980__41_20_491_0.html}

@BOOK{Kittel1995,
  author       = {Kittel, Charles},
  title        = {Introduction to Solid State Physics},
  publisher    = WIL,
  year         = {1995},
  volume       = {1},
  address      = NY,
  edition      = {7},
  month        = OCT,
  pages        = {688},
  isbn         = {0-471-11181-3}
}
%ISBN-13: 978-0471111818

@article{FangSS2005,
  title={Kinetics of protein adsorption and desorption on surfaces with grafted polymers},
  author={Fang, Fang and Satulovsky, Javier and Szleifer, Igal},
  journal=BJ,
  volume={89},
  number={3},
  pages={1516--1533},
  year={2005},
  publisher={Elsevier}
}

@article{Gillespie2008,
  title={Energetics of divalent selectivity in a calcium channel: the ryanodine receptor case study},
  author={Gillespie, Dirk},
  journal=BJ,
  volume={94},
  number={4},
  pages={1169--1184},
  year={2008},
  publisher={Elsevier}
}


@ARTICLE{FelixMH1986,
  author       = {Felix, J.~W. and Mukamel, D. and Hornreich, R.~M.},
  title        = {Novel class of continuous phase transitions to incommensurate structures},
  journal      = PRL,
  publisher    = APS,
  year         = {1986},
  volume       = {57},
  number       = {17},
  pages        = {2180-2183},
  month        = OCT
}
% doi = {10.1103/PhysRevLett.57.2180}
% url = {http://adsabs.harvard.edu/abs/1986PhRvL..57.2180F}

@article{HowardNP2017b,
  title={Stratification in Drying Polymer--Polymer and Colloid--Polymer Mixtures},
  author={Howard, Michael P and Nikoubashman, Arash and Panagiotopoulos, Athanassios Z},
  journal={Langmuir},
  volume={33},
  number={42},
  pages={11390--11398},
  year={2017},
  publisher={ACS Publications}
}

@ARTICLE{AslanyanL1978,
  author       = {Aslanyan, T.~A. and Levanyuk, A.~P.},
  title        = {Lifshits critical points},
  journal      = SPSS,
  publisher    = AIoP,
  year         = {1978},
  volume       = {20},
  number       = {3},
  pages        = {466-469},
  month        = MAR
}
% doi = {}
% url = {}

@Article{Grabert1978,
  author  = {Grabert, Hermann},
  title   = {Nonlinear transport and dynamics of fluctuations},
  journal = {Journal of Statistical Physics},
  volume  = {19},
  number  = {5},
  pages   = {479-497},
  year    = {1978},
  month   = NOV
}

@article{HermannKdlHS2019,
  title = {Phase coexistence of active {B}rownian particles},
  author = {Hermann, Sophie and Krinninger, Philip and {de las Heras}, Daniel and Schmidt, Matthias},
  journal = PRE,
  volume = {100},
  issue = {5},
  pages = {052604},
  numpages = {15},
  year = {2019},
  month = {Nov},
  publisher = {American Physical Society}
}

%   doi = {10.1103/PhysRevE.100.052604},
%  url = {https://link.aps.org/doi/10.1103/PhysRevE.100.052604}


@ARTICLE{Brand1986,
  author       = {Brand, H.~R.},
  title        = {Static Effects of Electric-Fields near the Nematic and Cholesteric to Isotropic Transitions},
  journal      = MCaLCL,
  publisher    = TF,
  year         = {1986},
  volume       = {3},
  number       = {5},
  pages        = {147-153},
  month        = MAR
}
% doi = {}
% url = {http://www.informaworld.com/smpp/title~db=all~content=t713644168~tab=issueslist}

@ARTICLE{Brand1986b,
  author       = {Brand, H.~R.},
  title        = {Dynamic Effects of Electric-Fields near the Nematic and Cholesteric to Isotropic-Phase Transitions},
  journal      = MCaLC,
  publisher    = TF,
  year         = {1986},
  volume       = {4},
  number       = {1},
  pages        = {23-30},
  month        = APR
}
% doi = {}
% url = {http://cat.inist.fr/?aModele=afficheN&cpsidt=7600171}

@ARTICLE{WatanabeEtAl2002,
  author       = {Watanabe, K. and Wakita, J.-I. and Itoh, H. and Shimada, H. and Kurosu, S. and Ikeda, T. and Yamazaki, Y. 
and Matsuyama, T. and Matsushita, M.},
  title        = {Dynamical properties of transient spatio-temporal patterns in bacterial colony of Proteus mirabilis},
  journal      = JotPSoJ,
  publisher    = PSoJ,
  year         = {2002},
  volume       = {71},
  number       = {2},
  pages        = {650-656},
  month        = FEB
}
% doi = {10.1143/JPSJ.71.650}
% url = {http://adsabs.harvard.edu/abs/2002JPSJ...71..650W}

@ARTICLE{VerkhovskySB1999,
  author       = {Verkhovsky, Alexander~B. and Svitkina, Tatyana~M. and Borisy, Gary~G.},
  title        = {Self-polarization and directional motility of cytoplasm},
  journal      = CB,
  publisher    = ES,
  year         = {1999},
  volume       = {9},
  number       = {1},
  pages        = {11-20},
  month        = DEC
}
% doi = {10.1016/S0960-9822(99)80042-6}
% url = {http://biomednet.com/elecref/0960982200900011}

@ARTICLE{CisnerosDGK2006,
  author       = {Cisneros, L. and Dombrowski, C. and Goldstein, R.~E. and Kessler, J.~O.},
  title        = {Reversal of bacterial locomotion at an obstacle},
  journal      = PRE,
  publisher    = APS,
  year         = {2006},
  volume       = {73},
  number       = {3},
  pages        = {030901},
  month        = MAR
}
% doi = {10.1103/PhysRevE.73.030901}
% url = {http://adsabs.harvard.edu/abs/2006PhRvE..73c0901C}

@ARTICLE{ChwangW1974,
  author       = {Chwang, A.~T. and Wu, T.~Y.},
  title        = {Hydromechanics of low-{R}eynolds-number flow. Part 1. Rotation of axisymmetric prolate bodies},
  journal      = JoFM,
  publisher    = CUP,
  year         = {1974},
  volume       = {63},
  number       = {3},
  pages        = {607-622},
  month        = FEB
}
% doi = {10.1017/S0022112074001819}
% url = {http://adsabs.harvard.edu/abs/1974JFM....63..607C}

@ARTICLE{ChwangW1975,
  author       = {Chwang, A.~T. and Wu, T.~Y.},
  title        = {Hydromechanics of low-{R}eynolds-number flow. Part 2. Singularity method for Stokes flows},
  journal      = JoFM,
  publisher    = CUP,
  year         = {1975},
  volume       = {67},
  number       = {4},
  pages        = {787-815},
  month        = FEB
}
% doi = {10.1017/S0022112075000614}
% url = {http://adsabs.harvard.edu/abs/1975JFM....67..787C}

@ARTICLE{Chwang1975,
  author       = {Chwang, A.~T.},
  title        = {Hydromechanics of low-{R}eynolds-number flow. Part 3. Motion of a spheroidal particle in quadratic flows},
  journal      = JoFM,
  publisher    = CUP,
  year         = {1975},
  volume       = {72},
  number       = {1},
  pages        = {17-34},
  month        = NOV
}
% doi = {10.1017/S0022112075002911}
% url = {http://adsabs.harvard.edu/abs/1975JFM....72...17C}

@ARTICLE{ChwangW1976,
  author       = {Chwang, A.~T. and Wu, T.~Y.},
  title        = {Hydromechanics of low-{R}eynolds-number flow. Part 4. Translation of spheroids},
  journal      = JoFM,
  publisher    = CUP,
  year         = {1976},
  volume       = {75},
  number       = {4},
  pages        = {677-689},
  month        = FEB
}
% doi = {10.1017/S0022112076000451}
% url = {http://adsabs.harvard.edu/abs/1976JFM....75..677C}

@ARTICLE{JohnsonW1979,
  author       = {Johnson, R.~E. and Wu, T.~Y.},
  title        = {Hydromechanics of low-{R}eynolds-number flow. Part 5. Motion of a slender torus},
  journal      = JoFM,
  publisher    = CUP,
  year         = {1979},
  volume       = {95},
  number       = {2},
  pages        = {263-277},
  month        = NOV
}
% doi = {10.1017/S0022112079001464}
% url = {http://adsabs.harvard.edu/abs/1979JFM....95..263J}

@ARTICLE{HuangC1986,
  author       = {Huang, L.~H. and Chwang, A.~T.},
  title        = {Hydromechanics of low-{R}eynolds-number flow. Part 6. Rotation of oblate bodies},
  journal      = JoEM,
  publisher    = SPR,
  year         = {1986},
  volume       = {20},
  number       = {4},
  pages        = {307-322},
  month        = DEC
}
% doi = {10.1007/BF00044608}
% url = {http://www.springerlink.com/content/r510652783173448/}

@ARTICLE{Freeman1985,
  author       = {Freeman, Brett},
  title        = {The motion of rigid ellipsoidal particles in slow flows},
  journal      = {Tectonophysics},
  publisher    = ELS,
  year         = {1985},
  volume       = {113},
  number       = {1-2},
  pages        = {163-183},
  month        = MAR
}
% doi = {10.1016/0040-1951(85)90115-5}
% url = {http://www.sciencedirect.com/science?_ob=ArticleURL&_udi=B6V72-48BDM7W-8C&_user=10&_coverDate=03%2F10%2F1985&_rdoc=1&_fmt=high&_orig=gateway&_origin=gateway&_sort=d&_docanchor=&view=c&_searchStrId=1683175430&_rerunOrigin=google&_acct=C000050221&_version=1&_urlVersion=0&_userid=10&md5=1cc1d5f7e4aafe7ec058d638db9acbfb&searchtype=a}

@ARTICLE{Jeffery1922,
  author       = {Jeffery, G.~B.},
  title        = {The motion of ellipsoidal particles immersed in a viscous fluid},
  journal      = PotRSoLSACPoaMaPC,
  publisher    = RS,
  year         = {1922},
  volume       = {102},
  number       = {715},
  pages        = {161-179},
  month        = NOV
}
% url = {http://adsabs.harvard.edu/abs/1922RSPSA.102..161J}

@ARTICLE{MazurB1974,
  author       = {Mazur, P. and Bedeaux, D.},
  title        = {A generalization of Fax{\'e}n's theorem to nonsteady motion of a sphere through an incompressible fluid in arbitrary flow},
  journal      = {Physica},
  publisher    = ELS,
  year         = {1974},
  volume       = {76},
  number       = {2},
  pages        = {235-246},
  month        = SEP
}
% doi = {10.1016/0031-8914(74)90197-9}
% url = {http://adsabs.harvard.edu/abs/1974Phy....76..235M}

@ARTICLE{Hills1975,
  author       = {Hills, B.~P.},
  title        = {A generalized {Langevin} equation for the angular velocity of a spherical Brownian particle from fluctuating hydrodynamics},
  journal      = PA,
  publisher    = ELS,
  year         = {1975},
  volume       = {80},
  number       = {4},
  pages        = {360-368},
  month        = FEB
}
% doi = {10.1016/0378-4371(75)90126-0}
% url = {http://adsabs.harvard.edu/abs/1975PhyA...80..360H}

@BOOK{HappelB1991,
  author       = {Happel, John and Brenner, Howard},
  title        = {Low {R}eynolds Number Hydrodynamics: With Special Applications to Particulate Media},
  publisher    = KAP,
  year         = {1991},
  volume       = {1},
  series       = {Mechanics of Fluids and Transport Processes},
  address      = DOR,
  edition      = {2},
  month        = SEP,
  pages        = {553},
  isbn         = {9-024-72877-0}
}

@BOOK{DoiE2007,
  author       = {Doi, M. and Edwards, S.~F.},
  title        = {The Theory of Polymer Dynamics},
  publisher    = OUP,
  year         = {2007},
  volume       = {73},
  series       = {International Series of Monographs on Physics},
  address      = OX,
  edition      = {1},
  month        = NOV,
  pages        = {391},
  isbn         = {978-0-198-52033-7}
}

@BOOK{Acheson1990,
  author       = {Acheson, J.~D.},
  title        = {Elementary Fluid Dynamics},
  publisher    = OUP,
  year         = {1990},
  volume       = {22},
  series       = {Oxford Applied Mathematics and Computing Science Series},
  address      = OX,
  edition      = {1},
  month        = MAR,
  pages        = {397},
  isbn         = {978-0-198-59679-0}
}

@BOOK{Batchelor2000,
  author       = {Batchelor, G.~K.},
  title        = {An Introduction to Fluid Dynamics},
  publisher    = CUP,
  year         = {2000},
  volume       = {5},
  series       = {Cambridge Mathematical Library},
  address      = NY,
  edition      = {1},
  month        = FEB,
  pages        = {615},
  isbn         = {978-0-521-66396-0}
}

@BOOK{Schutte1976,
  author       = {Schutte, Casper~Jan~Hendrik},
  title        = {The Quantum Mechanics and Group Theory of Vibrating and Rotating Molecules},
  publisher    = NH,
  year         = {1976},
  volume       = {1},
  series       = {The Theory of Molecular Spectroscopy},
  address      = AMST,
  edition      = {1},
  pages        = {512},
  isbn         = {0-720-40291-3}
}

@BOOK{Lamb1993,
  author       = {Lamb, Sir~Horace},
  title        = {Hydrodynamics},
  publisher    = CUP,
  year         = {1993},
  volume       = {6},
  series       = {Cambridge Mathematical Library},
  address      = NY,
  edition      = {6},
  month        = NOV,
  pages        = {768},
  isbn         = {978-0-521-45868-9}
}

@ARTICLE{Brenner1963a,
  author       = {Brenner, Howard},
  title        = {The Stokes resistance of an arbitrary particle},
  journal      = CES,
  publisher    = ES,
  year         = {1963},
  volume       = {18},
  number       = {1},
  pages        = {1-25},
  month        = JAN
}
% doi = {10.1016/0009-2509(63)80001-9}

@ARTICLE{Brenner1963b,
  author       = {Brenner, Howard},
  title        = {The Stokes resistance of an arbitrary particle - comment},
  journal      = CES,
  publisher    = ES,
  year         = {1963},
  volume       = {18},
  number       = {8},
  pages        = {557},
  month        = AUG
}

@ARTICLE{Brenner1964a,
  author       = {Brenner, Howard},
  title        = {The Stokes resistance of a slightly deformed sphere},
  journal      = CES,
  publisher    = ES,
  year         = {1964},
  volume       = {19},
  number       = {8},
  pages        = {519-539},
  month        = AUG
}
% doi = {10.1016/0009-2509(64)85045-4}

@ARTICLE{Brenner1964b,
  author       = {Brenner, Howard},
  title        = {The Stokes resistance of an arbitrary particle II: an extension},
  journal      = CES,
  publisher    = ES,
  year         = {1964},
  volume       = {19},
  number       = {9},
  pages        = {599-629},
  month        = SEP
}
% doi = {10.1016/0009-2509(64)85051-X}

@ARTICLE{Brenner1964c,
  author       = {Brenner, Howard},
  title        = {The Stokes resistance of an arbitrary particle III: shear fields},
  journal      = CES,
  publisher    = ES,
  year         = {1964},
  volume       = {19},
  number       = {9},
  pages        = {631-651},
  month        = SEP
}
% doi = {10.1016/0009-2509(64)85052-1}

@ARTICLE{Brenner1964d,
  author       = {Brenner, Howard},
  title        = {The Stokes resistance of an arbitrary particle IV: arbitrary fields of flow},
  journal      = CES,
  publisher    = ES,
  year         = {1964},
  volume       = {19},
  number       = {10},
  pages        = {703-727},
  month        = OCT
}
% doi = {10.1016/0009-2509(64)85084-3}

@ARTICLE{Brenner1965,
  author       = {Brenner, Howard},
  title        = {Coupling between the translational and rotational Brownian motions of rigid particles of arbitrary shape I: helicoidally isotropic particles},
  journal      = JoCS,
  publisher    = ES,
  year         = {1965},
  volume       = {20},
  number       = {2},
  pages        = {104-122},
  month        = FEB
}
% doi = {10.1016/0095-8522(65)90002-4}

@ARTICLE{Brenner1967,
  author       = {Brenner, Howard},
  title        = {Coupling between the translational and rotational Brownian motions of rigid particles of arbitrary shape II: general theory},
  journal      = JoCaIS,
  publisher    = ES,
  year         = {1967},
  volume       = {23},
  number       = {3},
  pages        = {407-436},
  month        = MAR
}
% doi = {10.1016/0021-9797(67)90185-3}

@INCOLLECTION{Brenner1966,
  author       = {Brenner, Howard},
  title        = {Hydrodynamic resistance of particles at small {R}eynolds numbers},
  booktitle    = {Advances in Chemical Engineering},
  publisher    = AP,
  year         = {1966},
  editor       = {Thomas~B. Drew and John~W. {Hoopes, Jr.} and Theodore Vermeulen},
  volume       = {6},
  series       = {Technology \& Engineering},
  type         = {chapter},
  chapter      = {5},
  pages        = {287-438},
  address      = NY,
  edition      = {1},
  isbn         = {978-0-120-08506-4}
}
% url = {http://www.sciencedirect.com/science/book/9780120085064}

@ARTICLE{TemleitnerP2010,
  author       = {Temleitner, L. and Pusztai, L.},
  title        = {Local order and orientational correlations in liquid and crystalline phases of carbon tetrabromide from neutron powder diffraction measurements},
  journal      = PRB,
  publisher    = APS,
  year         = {2010},
  volume       = {81},
  number       = {13},
  pages        = {134101},
  month        = APR
}
% doi = {10.1103/PhysRevB.81.134101}
% url = {http://adsabs.harvard.edu/abs/2010PhRvB..81m4101T}

@ARTICLE{SuzukiIGK1999,
  author       = {Suzuki, T. and Iiyama, T. and Gubbins, K.~E. and Kaneko, K.},
  title        = {Quasi-symmetry structure of {C}{C}l$_{4}$ molecular assemblies in a graphitic nanopore: a grand canonical {M}onte {C}arlo simulation},
  journal      = LANG,
  publisher    = ACS,
  year         = {1999},
  volume       = {15},
  number       = {18},
  pages        = {5870-5875},
  month        = AUG
}
% doi = {10.1021/la980960n}
% url = {http://pubs.acs.org/doi/abs/10.1021/la980960n}

@BOOK{KloedenP2006,
  author       = {Kloeden, Peter~E. and Platen, Eckhard},
  title        = {Numerical Solution of Stochastic Differential Equations},
  publisher    = SPR,
  year         = {2006},
  volume       = {23},
  series       = {Applications of Mathematics: Stochastic Modelling and Applied Probability},
  address      = BER,
  edition      = {1},
  month        = AUG,
  pages        = {636},
  isbn         = {3-540-54062-8}
}

@BOOK{Hassler2007,
  author       = {Hassler, Uwe},
  title        = {Stochastische Integration und Zeitreihenmodellierung: Eine Einf{\"u}hrung mit Anwendungen aus Finanzierung und {\"O}konometrie},
  publisher    = SPR,
  year         = {2007},
  volume       = {16},
  series       = {Statistik und ihre Anwendungen},
  address      = BER,
  edition      = {1},
  month        = SEP,
  pages        = {325},
  isbn         = {3-540-73567-4}
}

@ARTICLE{tenHagenvTL2009,
  author       = {ten~Hagen, B. and van~Teeffelen, S. and L{\"o}wen, H.},
  title        = {Non-Gaussian behaviour of a self-propelled particle on a substrate},
  journal      = CMP,
  publisher    = IfCMP,
  year         = {2009},
  volume       = {12},
  number       = {4},
  pages        = {725-738},
  month        = JUN
}
% url = {http://adsabs.harvard.edu/abs/2009arXiv0906.3418T}

@ARTICLE{ImperioRZ2008,
  author       = {Imperio, A. and Reatto, L. and Zapperi, S.},
  title        = {Rheology of colloidal microphases in a model with competing interactions},
  journal      = PRE,
  publisher    = APS,
  year         = {2008},
  volume       = {78},
  number       = {2},
  pages        = {021402},
  month        = AUG
}
% doi = {10.1103/PhysRevE.78.021402}
% url = {http://adsabs.harvard.edu/abs/2008PhRvE..78b1402I}

@ARTICLE{vanTeeffelenZL2009,
  author       = {van~Teeffelen, Sven and Zimmermann, Urs and L{\"o}wen, Hartmut},
  title        = {Clockwise-directional circle swimmer moves counter-clockwise in Petri dish- and ring-like confinements},
  journal      = SM,
  publisher    = RSoC,
  year         = {2009},
  volume       = {5},
  number       = {22},
  pages        = {4510-4519},
  month        = SEP
}
% doi = {10.1039/b911365g}

@article{MeyerVS2019,
  title={On the dynamics of reaction coordinates in classical, time-dependent, many-body processes},
  author={Meyer, Hugues and Voigtmann, Thomas and Schilling, Tanja},
  journal=JoChemP,
  volume={150},
  number={17},
  pages={174118},
  year={2019},
  publisher={AIP Publishing}
}

@article{LebyodkinBLE2020,
  title={Scaling in the local strain-rate field during jerky flow in an {Al}-{3\%} {Mg} alloy},
  author={Lebyodkin, Mikhail and Bougherira, Youcef and Lebedkina, Tatiana and Entemeyer, Denis},
  journal={Metals},
  volume={10},
  number={1},
  pages={134},
  year={2020},
  publisher={Multidisciplinary Digital Publishing Institute}
}

@article{AroldS2020,
  title={Active phase field crystal systems with inertial delay and underdamped dynamics},
  author={Arold, Dominic and Schmiedeberg, Michael},
  journal={European Physical Journal E},
  volume={43},
  number={7},
  pages={47},
  year={2020},
  publisher={Springer}
}

@article{AroldS2020b,
  title={Mean field approach of dynamical pattern formation in underdamped active matter with short-ranged alignment and distant anti-alignment interactions},
  author={Arold, Dominic and Schmiedeberg, Michael},
  journal=JoP:CM,
  volume={32},
  number={31},
  pages={315403},
  year={2020},
  publisher={IOP Publishing}
}

@article{DauchotD2019,
  title={Dynamics of a self-propelled particle in a harmonic trap},
  author={Dauchot, Olivier and D{\'e}mery, Vincent},
  journal=PRL,
  volume={122},
  number={6},
  pages={068002},
  year={2019},
  publisher={APS}
}

@article{MeyerPS2020,
  title={Non-{M}arkovian out-of-equilibrium dynamics: a general numerical procedure to construct time-dependent memory kernels for coarse-grained observables},
  author={Meyer, Hugues and Pelagejcev, Philipp and Schilling, Tanja},
  journal={Europhysics Letters},
  volume={128},
  number={4},
  pages={40001},
  year={2020},
  publisher={IOP Publishing}
}

@article{AmatiMS2019,
  title={Memory effects in the {F}ermi-{P}asta-{U}lam model},
  author={Amati, Graziano and Meyer, Hugues and Schilling, Tanja},
  journal={Journal of Statistical Physics},
  volume={174},
  number={1},
  pages={219--257},
  year={2019},
  publisher={Springer}
}

@article{MeyerVS2017,
  title={On the non-stationary generalized {L}angevin equation},
  author={Meyer, Hugues and Voigtmann, Thomas and Schilling, Tanja},
  journal=JoChemP,
  volume={147},
  number={21},
  pages={214110},
  year={2017},
  publisher={AIP Publishing LLC}
}

@ARTICLE{FradenMCM1989,
  author       = {Fraden, S. and Maret, G. and Caspar, D.~L.~D. and Meyer, R.~B.},
  title        = {Isotropic-nematic phase transition and angular correlations in isotropic suspensions of tobacco mosaic virus},
  journal      = PRL,
  publisher    = APS,
  year         = {1989},
  volume       = {63},
  number       = {19},
  pages        = {2068-2071},
  month        = NOV
}
% doi = {10.1103/PhysRevLett.63.2068}
% url = {http://adsabs.harvard.edu/abs/1989PhRvL..63.2068F}

@ARTICLE{WhiteCH2001,
  author       = {White, T.~O. and Ciccotti, G. and Hansen, J.-P.},
  title        = {Brownian dynamics with constraints},
  journal      = MP,
  publisher    = TF,
  year         = {2001},
  volume       = {99},
  number       = {24},
  pages        = {2023-2036},
  month        = DEC
}
% doi = {10.1080/00268970110090854}
% url = {http://adsabs.harvard.edu/abs/2001MolPh..99.2023W}

@ARTICLE{QuillietZRvBI2008,
  author       = {Quilliet, C. and Zoldesi, C. and Riera, C. and van~Blaaderen, A. and Imhof, A.},
  title        = {Anisotropic colloids through non-trivial buckling},
  journal      = EPJE,
  publisher    = EDPS,
  year         = {2008},
  volume       = {27},
  number       = {1},
  pages        = {13-20},
  month        = SEP
}
% doi = {10.1140/epje/i2007-10365-2}
% url = {http://adsabs.harvard.edu/abs/2008EPJE...27...13Q}

@ARTICLE{RoordavDPGvBK2004,
  author       = {Roorda, S. and van~Dillen, T. and Polman, A. and Graf, C. and van~Blaaderen, A. and Kooi, B.~J.},
  title        = {Aligned gold nanorods in silica made by ion irradiation of core-shell colloidal particles},
  journal      = AdvM,
  publisher    = WILEYVCH,
  year         = {2004},
  volume       = {16},
  number       = {3},
  pages        = {235-237},
  month        = FEB
}
% doi = {10.1002/adma.200305742}
% url = {http://onlinelibrary.wiley.com/doi/10.1002/adma.200305742/abstract}

@ARTICLE{GlotzerS2007,
  author       = {Glotzer, Sharon~C. and Solomon, Michael~J.},
  title        = {Anisotropy of building blocks and their assembly into complex structures},
  journal      = NM,
  publisher    = NPG,
  year         = {2007},
  volume       = {6},
  number       = {8},
  pages        = {557-562},
  month        = AUG
}
% doi = {10.1038/nmat1949}
% url = {http://www.nature.com/nmat/journal/v6/n8/full/nmat1949.html}

@ARTICLE{Loewen2001,
  author       = {L{\"o}wen, H.},
  title        = {Colloidal soft matter under external control},
  journal      = JoP:CM,
  publisher    = IoPP,
  year         = {2001},
  volume       = {13},
  number       = {24},
  pages        = {R415-R432},
  month        = JUN
}
% doi = {10.1088/0953-8984/13/24/201}
% url = {http://adsabs.harvard.edu/abs/2001JPCM...13R.415L}

@ARTICLE{ManoharanEP2003,
  author       = {Manoharan, V.~N. and Elsesser, M.~T. and Pine, D.~J.},
  title        = {Dense packing and symmetry in small clusters of microspheres},
  journal      = SCI,
  publisher    = AAftAoS,
  year         = {2003},
  volume       = {301},
  number       = {5632},
  pages        = {483-487},
  month        = JUL
}
% doi = {10.1126/science.1086189}
% url = {http://adsabs.harvard.edu/abs/2003Sci...301..483M}

@ARTICLE{SolomonZOSDSBGM2010,
  author       = {Solomon, Michael~J. and Zeitoun, Ramsey and Ortiz, Daniel and Sung, Kyung~Eun and Deng, Di and Shah, Aayush and Burns, Mark~A. and Glotzer, Sharon~C. and Millunchick, Joanna~M.},
  title        = {Toward assembly of non-close-packed colloidal structures from anisotropic pentamer particles},
  journal      = MRC,
  publisher    = WILEYVCH,
  year         = {2010},
  volume       = {31},
  number       = {2},
  pages        = {196-201},
  month        = JAN
}
% doi = {10.1002/marc.200900586}
% url = {http://onlinelibrary.wiley.com/doi/10.1002/marc.200900586/abstract}

@ARTICLE{JohnsonvKvB2005,
  author       = {Johnson, Patrick~M. and Kats, Carlos~M.~van and van~Blaaderen, Alfons},
  title        = {Synthesis of colloidal silica dumbbells},
  journal      = LANG,
  publisher    = ACS,
  year         = {2005},
  volume       = {21},
  number       = {24},
  pages        = {11510-11517},
  month        = OCT
}
% doi = {10.1021/la0518750}
% url = {http://pubs.acs.org/doi/abs/10.1021/la0518750}

@ARTICLE{KraftVvKvBIK2009,
  author       = {Kraft, Daniela~J. and Vlug, Wessel~S. and Kats, Carlos~M.~van and van~Blaaderen, Alfons and Imhof, Arnout and Kegel, Willem~K.},
  title        = {Self-assembly of colloids with liquid protrusions},
  journal      = JotACS,
  publisher    = ACS,
  year         = {2009},
  volume       = {131},
  number       = {3},
  pages        = {1182-1186},
  month        = JAN
}
% doi = {10.1021/ja8079803}
% url = {http://pubs.acs.org/doi/abs/10.1021/ja8079803}

@ARTICLE{WierengaLP1998,
  author       = {Wierenga, Anieke~M. and Lenstra, Tjerk~A.~J. and Philipse, Albert~P.},
  title        = {Aqueous dispersions of colloidal gibbsite platelets: synthesis, characterisation and intrinsic viscosity measurements},
  journal      = CaSA:PaEA,
  publisher    = ELS,
  year         = {1998},
  volume       = {134},
  number       = {3},
  pages        = {359-371},
  month        = MAR
}
% doi = {10.1016/S0927-7757(97)00224-0}

@ARTICLE{GaugerS2006,
  author       = {Gauger, E. and Stark, H.},
  title        = {Numerical study of a microscopic artificial swimmer},
  journal      = PRE,
  publisher    = APS,
  year         = {2006},
  volume       = {74},
  number       = {2},
  pages        = {021907},
  month        = AUG
}
% doi = {10.1103/PhysRevE.74.021907}
% url = {http://adsabs.harvard.edu/abs/2006PhRvE..74b1907G}

@ARTICLE{DharFWMPS2006,
  author       = {Dhar, P. and Fischer, T.~M. and Wang, Y. and Mallouk, T.~E. and Paxton, W.~F. and Sen, A.},
  title        = {Autonomously moving nanorods at a viscous interface},
  journal      = NL,
  publisher    = ACS,
  year         = {2006},
  volume       = {6},
  number       = {1},
  pages        = {66-72},
  month        = JAN
}
% doi = {10.1021/nl052027s}
% url = {http://adsabs.harvard.edu/abs/2006NanoL...6...66D}

@ARTICLE{WaltherM2008,
  author       = {Walther, Andreas and M{\"u}ller, Axel~H.~E.},
  title        = {Janus particles},
  journal      = SM,
  publisher    = RSoC,
  year         = {2008},
  volume       = {4},
  number       = {4},
  pages        = {663-668},
  month        = APR
}
% doi = {10.1039/B718131K}
% url = {http://pubs.rsc.org/en/Content/ArticleLanding/2008/SM/b718131k}

@ARTICLE{ErbeZBKL2008,
  author       = {Erbe, A. and Zientara, M. and Baraban, L. and Kreidler, C. and Leiderer, P.},
  title        = {Various driving mechanisms for generating motion of colloidal particles},
  journal      = JoP:CM,
  publisher    = IoPP,
  year         = {2008},
  volume       = {20},
  number       = {40},
  pages        = {4215},
  month        = OCT
}
% doi = {10.1088/0953-8984/20/40/404215}
% url = {http://adsabs.harvard.edu/abs/2008JPCM...20N4215E}

@ARTICLE{DreyfusBRFSB2005,
  author       = {Dreyfus, R. and Baudry, J. and Roper, M.~L. and Fermigier, M. and Stone, H.~A. and Bibette, J.},
  title        = {Microscopic artificial swimmers},
  journal      = NAT,
  publisher    = NPG,
  year         = {2005},
  volume       = {437},
  number       = {7060},
  pages        = {862-865},
  month        = OCT
}
% doi = {10.1038/nature04090}
% url = {http://adsabs.harvard.edu/abs/2005Natur.437..862D}

@ARTICLE{PopescuDO2009,
  author       = {Popescu, M.~N. and Dietrich, S. and Oshanin, G.},
  title        = {Confinement effects on diffusiophoretic self-propellers},
  journal      = JoChemP,
  publisher    = AIoP,
  year         = {2009},
  volume       = {130},
  number       = {19},
  pages        = {194702},
  month        = MAY
}
% doi = {10.1063/1.3133239}
% url = {http://adsabs.harvard.edu/abs/2009JChPh.130s4702P}

@ARTICLE{BergT1990,
  author       = {Berg, H.~C. and Turner, L.},
  title        = {Chemotaxis of bacteria in glass capillary arrays. Escherichia coli, motility, microchannel plate, and light scattering},
  journal      = BJ,
  publisher    = ELS,
  year         = {1990},
  volume       = {58},
  number       = {4},
  pages        = {919-930},
  month        = OCT
}
% doi = {10.1016/S0006-3495(90)82436-X}
% url = {http://adsabs.harvard.edu/abs/1990BpJ....58..919B}

@ARTICLE{DiluzioTMGWBW2005,
  author       = {Diluzio, W.~R. and Turner, L. and Mayer, M. and Garstecki, P. and Weibel, D.~B. and Berg, H.~C. and Whitesides, G.~M.},
  title        = {Escherichia coli swim on the right-hand side},
  journal      = NAT,
  publisher    = NPG,
  year         = {2005},
  volume       = {435},
  number       = {7046},
  pages        = {1271-1274},
  month        = JUN
}
% doi = {10.1038/nature03660}
% url = {http://adsabs.harvard.edu/abs/2005Natur.435.1271D}

@ARTICLE{LaugaDWS2006,
  author       = {Lauga, E. and Diluzio, W. and Whitesides, G. and Stone, H.},
  title        = {Swimming in circles: motion of bacteria near solid boundaries},
  journal      = BJ,
  publisher    = ELS,
  year         = {2006},
  volume       = {90},
  number       = {2},
  pages        = {400-412},
  month        = JAN
}
% doi = {10.1529/biophysj.105.069401}
% url = {http://adsabs.harvard.edu/abs/2006BpJ....90..400L}

@ARTICLE{HillKMK2007,
  author       = {Hill, J. and Kalkanci, O. and McMurry, J.~L. and Koser, H.},
  title        = {Hydrodynamic surface interactions enable Escherichia coli to seek efficient routes to swim upstream},
  journal      = PRL,
  publisher    = APS,
  year         = {2007},
  volume       = {98},
  number       = {6},
  pages        = {068101},
  month        = FEB
}
% doi = {10.1103/PhysRevLett.98.068101}
% url = {http://adsabs.harvard.edu/abs/2007PhRvL..98f8101H}

@ARTICLE{ShenoyTPT2007,
  author       = {Shenoy, V.~B. and Tambe, D.~T. and Prasad, A. and Theriot, J.~A.},
  title        = {A kinematic description of the trajectories of Listeria monocytogenes propelled by actin comet tails},
  journal      = PotNAoS,
  publisher    = NAoS,
  year         = {2007},
  volume       = {104},
  number       = {20},
  pages        = {8229-8234},
  month        = MAY
}
% doi = {10.1073/pnas.0702454104}
% url = {http://adsabs.harvard.edu/abs/2007PNAS..104.8229S}

@ARTICLE{SchmidtvdGBWHF2008,
  author       = {Schmidt, Stephan and van~der~Gucht, Jasper and Biesheuvel, P.~Maarten and Weinkamer, Richard and Helfer, Emmanu{\`e}le and Fery, Andreas},
  title        = {Non-Gaussian curvature distribution of actin-propelled biomimetic colloid trajectories},
  journal      = EBJ,
  publisher    = SPR,
  year         = {2008},
  volume       = {37},
  number       = {8},
  pages        = {1361-1366},
  month        = OCT
}
% doi = {10.1007/s00249-008-0340-x}
% url = {http://www.springerlink.com/content/j72n0u6884u4xl2g/}

@ARTICLE{RiedelKH2005,
  author       = {Riedel, I.~H. and Kruse, K. and Howard, J.},
  title        = {A Self-Organized Vortex Array of Hydrodynamically Entrained Sperm Cells},
  journal      = SCI,
  publisher    = AAftAoS,
  year         = {2005},
  volume       = {309},
  number       = {5732},
  pages        = {300-303},
  month        = JUL
}
% doi = {10.1126/science.1110329}
% url = {http://adsabs.harvard.edu/abs/2005Sci...309..300R}

@ARTICLE{Woolley2003,
  author       = {Woolley, D.~M.},
  title        = {Motility of spermatozoa at surfaces},
  journal      = {Reproduction},
  publisher    = {Society for Reproduction and Fertility},
  year         = {2003},
  volume       = {126},
  number       = {2},
  pages        = {259-270},
  month        = AUG
}
% doi = {10.1530/rep.0.1260259}
% url = {http://www.reproduction-online.org/cgi/content/abstract/126/2/259}

@ARTICLE{FriedrichJ2008,
  author       = {Friedrich, B.~M. and J{\"u}licher, F.},
  title        = {The stochastic dance of circling sperm cells: sperm chemotaxis in the plane},
  journal      = NJoP,
  publisher    = IoPP,
  year         = {2008},
  volume       = {10},
  number       = {12},
  pages        = {123025},
  month        = DEC
}
% doi = {10.1088/1367-2630/10/12/123025}
% url = {http://adsabs.harvard.edu/abs/2008NJPh...10l3025F}

@ARTICLE{CaldererFW2004,
  author       = {Calderer, M.~Carme and Forest, M.~Gregory and Wang, Qi},
  title        = {Kinetic theories and mesoscopic models for solutions of nonhomogeneous liquid crystal polymers},
  journal      = JoNNFM,
  publisher    = ELS,
  year         = {2004},
  volume       = {120},
  number       = {1-3},
  pages        = {69-78},
  month        = JUL
}
% doi = {10.1016/j.jnnfm.2004.01.015}
% url = {http://www.sciencedirect.com/science?_ob=ArticleURL&_udi=B6TGV-4CSG50T-7&_user=2665780&_coverDate=07%2F01%2F2004&_rdoc=1&_fmt=high&_orig=gateway&_origin=gateway&_sort=d&_docanchor=&view=c&_rerunOrigin=scholar.google&_acct=C000053885&_version=1&_urlVersion=0&_userid=2665780&md5=81f47a2f882db0f0f83b59fba0f9f492&searchtype=a}

@ARTICLE{Lubensky2000,
  author       = {Lubensky, T.~C.},
  title        = {Liquid crystals - New banana phases},
  journal      = SCI,
  publisher    = AAftAoS,
  year         = {2000},
  volume       = {288},
  number       = {5474},
  pages        = {2146-2147},
  month        = JUN
}

@ARTICLE{vandenPolPTWBV2009,
  author       = {van~den~Pol, E. and Petukhov, A.~V. and {Thies-Weesie}, D.~M.~E. and Byelov, D.~V. and Vroege, G.~J.},
  title        = {Experimental realization of biaxial liquid crystal phases in colloidal dispersions of boardlike particles},
  journal      = PRL,
  publisher    = APS,
  year         = {2009},
  volume       = {103},
  number       = {25},
  pages        = {258301},
  month        = DEC
}
% doi = {10.1103/PhysRevLett.103.258301}
% url = {http://adsabs.harvard.edu/abs/2009PhRvL.103y8301V}

@ARTICLE{HowseJRGVG2007,
  author       = {Howse, J.~R. and Jones, R.~A.~L. and Ryan, A.~J. and Gough, T. and Vafabakhsh, R. and Golestanian, R.},
  title        = {Self-motile colloidal particles: from directed propulsion to random walk},
  journal      = PRL,
  publisher    = APS,
  year         = {2007},
  volume       = {99},
  number       = {4},
  pages        = {048102},
  month        = JUL
}
% doi = {10.1103/PhysRevLett.99.048102}
% url = {http://adsabs.harvard.edu/abs/2007PhRvL..99d8102H}

@ARTICLE{JahnV1972,
  author       = {Jahn, T.~L. and Votta, J.~J.},
  title        = {Locomotion of protozoa},
  journal      = {Annual Review of Fluid Mechanics},
  publisher    = AR,
  year         = {1972},
  volume       = {4},
  number       = {1},
  pages        = {93-116},
  month        = JAN
}
% doi = {10.1146/annurev.fl.04.010172.000521}
% url = {http://adsabs.harvard.edu/abs/1972AnRFM...4...93J}

@ARTICLE{SchulzJ2001,
  author       = {Schulz, Heide~N. and J{\o{}}rgensen, Bo~Barker},
  title        = {Big bacteria},
  journal      = {Annual Review of Microbiology},
  publisher    = AR,
  year         = {2001},
  volume       = {55},
  number       = {1},
  pages        = {105-137},
  month        = OCT
}
% doi = {10.1146/annurev.micro.55.1.105}
% url = {http://www.annualreviews.org/doi/abs/10.1146/annurev.micro.55.1.105}

@ARTICLE{NewtonW2007,
  author       = {Newton, Marcus~C. and Warburton, Paul~A.},
  title        = {ZnO tetrapod nanocrystals},
  journal      = MT,
  publisher    = ELS,
  year         = {2007},
  volume       = {10},
  number       = {5},
  pages        = {50-54},
  month        = MAY
}
% doi = {10.1016/S1369-7021(07)70079-2}
% url = {http://www.sciencedirect.com/science?_ob=ArticleURL&_udi=B6X1J-4NMDD7T-S&_user=10&_coverDate=05%2F31%2F2007&_rdoc=1&_fmt=high&_orig=gateway&_origin=gateway&_sort=d&_docanchor=&view=c&_rerunOrigin=scholar.google&_acct=C000050221&_version=1&_urlVersion=0&_userid=10&md5=653527d84f8e1cc0f9f3b6eebf823419&searchtype=a}

@ARTICLE{WuCH2009,
  author       = {Wu, {Hsin-Lun} and Chen, {Chiu-Hua} and Huang, Michael~H.},
  title        = {Seed-mediated synthesis of branched gold nanocrystals derived from the side growth of pentagonal bipyramids and the formation of gold nanostars},
  journal      = CoM,
  publisher    = ACS,
  year         = {2009},
  volume       = {21},
  number       = {1},
  pages        = {110-114},
  month        = JAN
}
% doi = {10.1021/cm802257e}
% url = {http://pubs.acs.org/doi/abs/10.1021/cm802257e}

@ARTICLE{ZhouLXWZZW2006,
  author       = {Zhou, Guangjun and L{\"u}, Mengkai and Xiu, Zhiliang and Wang, Shufen and Zhang, Haiping and Zhou, Yuanyuan and Wang, Shumei},
  title        = {Controlled synthesis of high-quality PbS star-shaped dendrites, multipods, truncated nanocubes, and nanocubes and their shape evolution process},
  journal      = JoPChemB,
  publisher    = ACS,
  year         = {2006},
  volume       = {110},
  number       = {13},
  pages        = {6543-6548},
  month        = FEB
}
% doi = {10.1021/jp0549881}
% url = {http://pubs.acs.org/doi/abs/10.1021/jp0549881}

@ARTICLE{DekaMDGBM2010,
  author       = {Deka, S. and Miszta, K. and Dorfs, D. and Genovese, A. and Bertoni, G. and Manna, L.},
  title        = {Octapod-shaped colloidal nanocrystals of cadmium chalcogenides via ``one-pot'' cation exchange and seeded growth},
  journal      = NL,
  publisher    = ACS,
  year         = {2010},
  volume       = {10},
  number       = {9},
  pages        = {3770-3776},
  month        = SEP
}
% doi = {10.1021/nl102539a}
% url = {http://adsabs.harvard.edu/abs/2010NanoL..10.3770D}

@ARTICLE{Dean1996,
  author       = {Dean, D.~S.},
  title        = {Langevin equation for the density of a system of interacting {Langevin} processes},
  journal      = JoPA:MaG,
  publisher    = IoPP,
  year         = {1996},
  volume       = {29},
  number       = {24},
  pages        = {L613-L617},
  month        = DEC
}
% doi = {10.1088/0305-4470/29/24/001}
% url = {http://adsabs.harvard.edu/abs/1996JPhA...29L.613D}

@ARTICLE{AcharyaPK2004,
  author       = {Acharya, B.~R. and Primak, A. and Kumar, S.},
  title        = {Biaxial Nematic Phase in Bent-Core Thermotropic Mesogens},
  journal      = PRL,
  publisher    = APS,
  year         = {2004},
  volume       = {92},
  number       = {14},
  pages        = {145506},
  month        = APR
}
% doi = {10.1103/PhysRevLett.92.145506}
% url = {http://adsabs.harvard.edu/abs/2004PhRvL..92n5506A}

@ARTICLE{Vorlaender1929,
  author       = {Vorl{\"a}nder, D.},
  title        = {Die Richtung der Kohlenstoff-Valenzen in Benzol-Abk{\"o}mmlingen},
  journal      = BddcG,
  publisher    = WILEYVCH,
  year         = {1929},
  volume       = {62},
  number       = {10},
  pages        = {2831-2835},
  month        = NOV
}
% doi = {10.1002/cber.19290621026}
% url = {http://onlinelibrary.wiley.com/doi/10.1002/cber.19290621026/abstract?systemMessage=Wiley+Online+Library+will+be+disrupted+14+May+from+10-12+BST+for+monthly+maintenance}

@ARTICLE{VorlaenderA1932,
  author       = {Vorl{\"a}nder, D. and Apel, Alfons},
  title        = {Die Richtung der Kohlenstoff-Valenzen in Benzolabk??mlingen II},
  journal      = BddcG,
  publisher    = WILEYVCH,
  year         = {1932},
  volume       = {65},
  number       = {7},
  pages        = {1101-1109},
  month        = JUL
}
% doi = {10.1002/cber.19320650710}
% url = {http://onlinelibrary.wiley.com/doi/10.1002/cber.19320650710/abstract}

@ARTICLE{Gubbins1980,
  author       = {Gubbins, K.~E.},
  title        = {Structure of non-uniform molecular fluids: integrodifferential equations for the density-orientation profile},
  journal      = CPL,
  publisher    = ELS,
  year         = {1980},
  volume       = {76},
  number       = {2},
  pages        = {329-332},
  month        = DEC
}
% doi = {10.1016/0009-2614(80)87034-5}
% url = {http://adsabs.harvard.edu/abs/1980CPL....76..329G}

@ARTICLE{YanLWLZ2006,
  author       = {Yan, Qingfeng and Liu, Fang and Wang, Likui and Lee, Jim~Yang and Zhao, X.~S.},
  title        = {Drilling nanoholes in colloidal spheres by selective etching},
  journal      = JoMC,
  publisher    = RSoC,
  year         = {2006},
  volume       = {16},
  number       = {22},
  pages        = {2132-2134},
  month        = MAY
}
% doi = {10.1039/B604463H}
% url = {http://pubs.rsc.org/en/Content/ArticleLanding/2006/JM/b604463h}

@article{HuWWL2009,
  title={Stable and efficient finite-difference nonlinear-multigrid schemes for the phase field crystal equation},
  author={Hu, Zhengzheng and Wise, Steven M and Wang, Cheng and Lowengrub, John S},
  journal=JoCompP,
  volume={228},
  number={15},
  pages={5323--5339},
  year={2009},
  publisher={Elsevier}
}

@article{WhiteGRV2000,
  title={Density-functional theory of inhomogeneous fluids in the canonical ensemble},
  author={White, JA and Gonz{\'a}lez, A and Rom{\'a}n, FL and Velasco, S},
  journal=PRL,
  volume={84},
  number={6},
  pages={1220},
  year={2000},
  publisher={APS}
}

@article{BaskaranHLWWZ2013,
  title={Energy stable and efficient finite-difference nonlinear multigrid schemes for the modified phase field crystal equation},
  author={Baskaran, Arvind and Hu, Zhengzheng and Lowengrub, John S and Wang, Cheng and Wise, Steven M and Zhou, Peng},
  journal=JoCompP,
  volume={250},
  pages={270--292},
  year={2013},
  publisher={Elsevier}
}

@ARTICLE{RexL2009,
  author       = {Rex, M. and L{\"o}wen, H.},
  title        = {Dynamical density functional theory for colloidal dispersions including hydrodynamic interactions},
  journal      = EPJE,
  publisher    = EDPS,
  year         = {2009},
  volume       = {28},
  number       = {2},
  pages        = {139-146},
  month        = FEB
}
% doi = {10.1140/epje/i2008-10363-x}
% url = {http://adsabs.harvard.edu/abs/2009EPJE...28..139R}

@ARTICLE{HansenGoosM2010,
  author       = {{Hansen-Goos}, Hendrik and Mecke, Klaus},
  title        = {Tensorial density functional theory for non-spherical hard-body fluids},
  journal      = JoP:CM,
  publisher    = IoPP,
  year         = {2010},
  volume       = {22},
  number       = {36},
  pages        = {364107},
  month        = AUG
}
% doi = {10.1088/0953-8984/22/36/364107}
% url = {http://iopscience.iop.org/0953-8984/22/36/364107}

@ARTICLE{Loewen1999,
  author       = {L{\"o}wen, H.},
  title        = {Anisotropic self-diffusion in colloidal nematic phases},
  journal      = PRE,
  publisher    = APS,
  year         = {1999},
  volume       = {59},
  number       = {2},
  pages        = {1989-1995},
  month        = FEB
}
% doi = {10.1103/PhysRevE.59.1989}
% url = {http://adsabs.harvard.edu/abs/1999PhRvE..59.1989L}

@ARTICLE{SchillingF2004,
  author       = {Schilling, T. and Frenkel, D.},
  title        = {Self-poisoning of crystal nuclei in hard-rod liquids},
  journal      = JoP:CM,
  publisher    = IoPP,
  year         = {2004},
  volume       = {16},
  number       = {19},
  pages        = {2029},
  month        = MAY 
}
% doi = {10.1088/0953-8984/16/19/014}
% url = {http://adsabs.harvard.edu/abs/2004JPCM...16S2029S}

@ARTICLE{MillerC2009,
  author       = {Miller, W.~L. and Cacciuto, A.},
  title        = {Hierarchical self-assembly of asymmetric amphiphatic spherical colloidal particles},
  journal      = PRE,
  publisher    = APS,
  year         = {2009},
  volume       = {80},
  number       = {2},
  pages        = {021404},
  month        = AUG
}
% doi = {10.1103/PhysRevE.80.021404}
% url = {http://adsabs.harvard.edu/abs/2009PhRvE..80b1404M}

@ARTICLE{MazzaGVKS2010,
  author       = {Mazza, M.~G. and Greschek, M. and Valiullin, R. and K{\"a}rger, J. and Schoen, M.},
  title        = {Entropy-Driven Enhanced Self-Diffusion in Confined Reentrant Supernematics},
  journal      = PRL,
  publisher    = APS,
  year         = {2010},
  volume       = {105},
  number       = {22},
  pages        = {227802},
  month        = 2010
}
% doi = {10.1103/PhysRevLett.105.227802}
% url = {http://adsabs.harvard.edu/abs/2010PhRvL.105v7802M}

@ARTICLE{PiniTPR2003,
  author       = {Pini, D. and Tau, M. and Parola, A. and Reatto, L.},
  title        = {Phase diagram of symmetric binary mixtures at equimolar and nonequimolar concentrations: a systematic investigation},
  journal      = PRE,
  publisher    = APS,
  year         = {2003},
  volume       = {67},
  number       = {4},
  pages        = {046116},
  month        = APR
}
% doi = {10.1103/PhysRevE.67.046116}
% url = {http://adsabs.harvard.edu/abs/2003PhRvE..67d6116P}

@ARTICLE{HanANZLY2006,
  author       = {Han, Y. and Alsayed, A.~M. and Nobili, M. and Zhang, J. and Lubensky, T.~C. and Yodh, A.~G.},
  title        = {Brownian motion of an ellipsoid},
  journal      = SCI,
  publisher    = AAftAoS,
  year         = {2006},
  volume       = {314},
  number       = {5799},
  pages        = {626-630},
  month        = OCT
}
% doi = {10.1126/science.1130146}
% url = {http://www.sciencemag.org/content/314/5799/626.short}

@ARTICLE{SircarLW2010,
  author       = {Sircar, Sarthok and Li, Jun and Wang, Qi},
  title        = {Biaxial phases of bent-core liquid crystal polymers in shear flows},
  journal      = CiMS,
  publisher    = IP,
  year         = {2010},
  volume       = {8},
  number       = {3},
  pages        = {697-720},
  month        = SEP
}
% url = {http://projecteuclid.org/euclid.cms/1282747136}

@ARTICLE{Thaokar2008,
  author       = {Thaokar, Rochish~M.},
  title        = {Brownian motion of a torus},
  journal      = CaSA:PaEA,
  publisher    = ELS,
  year         = {2008},
  volume       = {317},
  number       = {1-3},
  pages        = {650-657},
  month        = MAR
}
% doi = {doi:10.1016/j.colsurfa.2007.11.058}
% url = {http://www.sciencedirect.com/science/article/pii/S0927775707010217}

@ARTICLE{RavnikZ2009,
  author       = {Ravnik, Miha and \u{Z}umer, Slobodan},
  title        = {Nematic colloids entangled by topological defects},
  journal      = SM,
  publisher    = RSoC,
  year         = {2009},
  volume       = {5},
  number       = {2},
  pages        = {269-274},
  month        = JAN
}
% doi = {10.1039/B810933H}
% url = {http://pubs.rsc.org/en/Content/ArticleLanding/2009/SM/b810933h}

@ARTICLE{ZhangvD2006,
  author       = {Zhang, Z.~X. and van~Duijneveldt, J.~S.},
  title        = {Isotropic-nematic phase transition of nonaqueous suspensions of natural clay rods},
  journal      = JoChemP,
  publisher    = AIoP,
  year         = {2006},
  volume       = {124},
  number       = {15},
  pages        = {154910},
  month        = APR
}
% doi = {10.1063/1.2185642}
% url = {http://adsabs.harvard.edu/abs/2006JChPh.124o4910Z}

@ARTICLE{VerhoeffBOvdSL2011,
  author       = {Verhoeff, A.~A. and Bakelaar, I.~A. and Otten, R.~H.~J. and van~der~Schoot, P. and Lekkerkerker, H.~N.~W.},
  title        = {Tactoids of Plate-Like Particles: Size, Shape, and Director Field},
  journal      = LANG,
  publisher    = ACS,
  year         = {2011},
  volume       = {27},
  number       = {1},
  pages        = {116-125},
  month        = JAN
}
% doi = {10.1021/la104128m}
% url = {http://pubs.acs.org/doi/abs/10.1021/la104128m}

@article{Yoshimori2004,
  title={Time dependent density functional methods and their application to chemical physics},
  author={Yoshimori, Akira},
  journal={Journal of Theoretical and Computational Chemistry},
  volume={3},
  number={01},
  pages={117--144},
  year={2004},
  publisher={World Scientific}
}

@article{GoddardPK2012,
  title={The overdamped limit of dynamic density functional theory: Rigorous results},
  author={Goddard, Benjamin D and Pavliotis, Grigorios A and Kalliadasis, Serafim},
  journal={Multiscale Modeling \& Simulation},
  volume={10},
  number={2},
  pages={633--663},
  year={2012},
  publisher={SIAM}
}

@article{CalefW1983,
  title={Smoluchowski--{V}lasov theory of charge solvation dynamics},
  author={Calef, Daniel F and Wolynes, Peter G},
  journal=JoChemP,
  volume={78},
  number={6},
  pages={4145--4153},
  year={1983},
  publisher={AIP}
}

@article{MarechalL2013,
  title={Density functional theory for hard polyhedra},
  author={Marechal, Matthieu and L{\"o}wen, Hartmut},
  journal=PRL,
  volume={110},
  number={13},
  pages={137801},
  year={2013},
  publisher={APS}
}

@article{YatsyshinSK2012,
  title={Spectral methods for the equations of classical density-functional theory: Relaxation dynamics of microscopic films},
  author={Yatsyshin, Petr and Savva, Nikos and Kalliadasis, Serafim},
  journal=JoChemP,
  volume={136},
  number={12},
  pages={124113},
  year={2012},
  publisher={AIP}
}

@ARTICLE{CasagrandeV1988,
  author       = {Casagrande, C. and Veyssi{\'e}, M.},
  title        = {Janus beads - realization and 1st observation of interfacial properties},
  journal      = CRdlAdS:SII,
  publisher    = FAoS,
  year         = {1988},
  volume       = {306},
  number       = {20},
  pages        = {1423-1425},
  month        = MAY
}

@ARTICLE{CasagrandeFRV1989,
  author       = {Casagrande, C. and Fabre, P. and Rapha{\"e}l, E. and Veyssi{\'e}, M.},
  title        = {Janus beads: realization and behaviour at water/oil interfaces},
  journal      = EPL,
  publisher    = EPS,
  year         = {1989},
  volume       = {9},
  number       = {3},
  pages        = {251},
  month        = JUN
}
% doi = {10.1209/0295-5075/9/3/011}
% url = {http://adsabs.harvard.edu/abs/1989EL......9..251C}

@ARTICLE{JaszczakRDG2003,
  author       = {Jaszczak, John~A. and Robinson, George~W. and Dimovski, Svetlana and Gogotsi, Yury},
  title        = {Naturally occurring graphite cones},
  journal      = {Carbon},
  publisher    = ELS,
  year         = {2003},
  volume       = {41},
  number       = {11},
  pages        = {2085-2092},
  month        = AUG
}
% doi = {10.1016/S0008-6223(03)00214-8}
% url = {http://www.sciencedirect.com/science/article/pii/S0008622303002148}

@ARTICLE{KrishnanDTHLE1997,
  author       = {Krishnan, A. and Dujardin, E. and Treacy, M.~M.~J. and Hugdahl, J. and Lynum, S. and Ebbesen, T.~W.},
  title        = {Graphitic cones and the nucleation of curved carbon surfaces},
  journal      = NAT,
  publisher    = NPG,
  year         = {1997},
  volume       = {388},
  number       = {6641},
  pages        = {451-454},
  month        = JUL
}
% doi = {10.1038/41284}
% url = {http://adsabs.harvard.edu/abs/1997Natur.388..451K}

@ARTICLE{NaessEHK2009,
  author       = {Naess, S.~Nalum and Elgsaeter, A. and Helgesen, G. and Knudsen, K.~D.},
  title        = {Carbon nanocones: wall structure and morphology},
  journal      = SaToAM,
  publisher    = ELS,
  year         = {2009},
  volume       = {10},
  number       = {6},
  pages        = {065002},
  month        = DEC
}
% doi = {10.1088/1468-6996/10/6/065002}
% url = {http://adsabs.harvard.edu/abs/2009STAdM..10f5002N}

@ARTICLE{MourchidDLLL1995,
  author       = {Mourchid, A. and Delville, A. and Lambard, J. and L{\'e}Colier, E. and Levitz, P.},
  title        = {Phase diagram of colloidal dispersions of anisotropic charged particles: equilibrium properties, structure, and rheology of laponite suspensions},
  journal      = LANG,
  publisher    = ACS,
  year         = {1995},
  volume       = {11},
  number       = {6},
  pages        = {1942-1950},
  month        = JUN
}
% doi = {10.1021/la00006a020}
% url = {http://pubs.acs.org/doi/abs/10.1021/la00006a020}

@ARTICLE{DijkstraHM1995,
  author       = {Dijkstra, M. and Hansen, J.-P. and Madden, P.~A.},
  title        = {Gelation of a clay colloid suspension},
  journal      = PRL,
  publisher    = APS,
  year         = {1995},
  volume       = {75},
  number       = {11},
  pages        = {2236-2239},
  month        = SEP
}
% doi = {10.1103/PhysRevLett.75.2236}
% url = {http://adsabs.harvard.edu/abs/1995PhRvL..75.2236D}

@ARTICLE{Helseth2005,
  author       = {Helseth, L.~E.},
  title        = {Self-assembly of colloidal pyramids in magnetic fields},
  journal      = LANG,
  publisher    = ACS,
  year         = {2005},
  volume       = {21},
  number       = {16},
  pages        = {7276-7279},
  month        = AUG
}
% doi = {10.1021/la051140v}
% url = {http://pubs.acs.org/doi/abs/10.1021/la051140v}

@ARTICLE{FanTOB2008,
  author       = {Fan, Dongbo and Thomas, P.~John and O'Brien, Paul},
  title        = {Pyramidal lead sulfide crystallites with high energy \{113\} facets},
  journal      = JotACS,
  publisher    = ACS,
  year         = {2008},
  volume       = {130},
  number       = {33},
  pages        = {10892-10894},
  month        = JUL
}
% doi = {10.1021/ja804516q}
% url = {http://pubs.acs.org/doi/abs/10.1021/ja804516q}

@ARTICLE{GuhaKC2004,
  author       = {Guha, P. and Kar, S. and Chaudhuri, S.},
  title        = {Direct synthesis of single crystalline In$_{2}$O$_{3}$ nanopyramids and nanocolumns and their photoluminescence properties},
  journal      = APL,
  publisher    = AIoP,
  year         = {2004},
  volume       = {85},
  number       = {17},
  pages        = {3851},
  month        = OCT
}
% doi = {10.1063/1.1808886}
% url = {http://adsabs.harvard.edu/abs/2004ApPhL..85.3851G}

@ARTICLE{MouradPVL2010,
  author       = {Mourad, Maurice~C.~D. and Petukhov, Andrei~V. and Vroege, Gert~Jan and Lekkerkerker, Henk~N.~W.},
  title        = {Lyotropic hexagonal columnar liquid crystals of large colloidal gibbsite platelets},
  journal      = LANG,
  publisher    = ACS,
  year         = {2010},
  volume       = {26},
  number       = {17},
  pages        = {14182-14187},
  month        = AUG
}
% doi = {10.1021/la100797x}
% url = {http://pubs.acs.org/doi/abs/10.1021/la100797x}

@ARTICLE{LapointeHMS2010,
  author       = {Lapointe, C.~P. and Hopkins, S. and Mason, T.~G. and Smalyukh, I.~I.},
  title        = {Electrically driven multiaxis rotational dynamics of colloidal platelets in nematic liquid crystals},
  journal      = PRL,
  publisher    = APS,
  year         = {2010},
  volume       = {105},
  number       = {17},
  pages        = {178301},
  month        = OCT
}
% doi = {10.1103/PhysRevLett.105.178301}
% url = {http://adsabs.harvard.edu/abs/2010PhRvL.105q8301L}

@ARTICLE{DijkstraHM1997,
  author       = {Dijkstra, M. and Hansen, J.-P. and Madden, P.~A.},
  title        = {Statistical model for the structure and gelation of smectite clay suspensions},
  journal      = PRE,
  publisher    = APS,
  year         = {1997},
  volume       = {55},
  number       = {3},
  pages        = {3044-3053},
  month        = MAR
}
% doi = {10.1103/PhysRevE.55.3044}
% url = {http://adsabs.harvard.edu/abs/1997PhRvE..55.3044D}

@ARTICLE{MarechalKDID2010,
  author       = {Marechal, Matthieu and Kortschot, Rob~J. and Demir{\"o}rs, Ahmet~Faik and Imhof, Arnout and Dijkstra, Marjolein},
  title        = {Phase behavior and structure of a new colloidal model system of bowl-shaped particles},
  journal      = NL,
  publisher    = ACS,
  year         = {2010},
  volume       = {10},
  number       = {5},
  pages        = {1907-1911},
  month        = APR
}
% doi = {10.1021/nl100783g}
% url = {http://pubs.acs.org/doi/abs/10.1021/nl100783g}

@ARTICLE{MarechalD2010,
  author       = {Marechal, M. and Dijkstra, M.},
  title        = {Phase behavior and structure of colloidal bowl-shaped particles: simulations},
  journal      = PRE,
  publisher    = APS,
  year         = {2010},
  volume       = {82},
  number       = {3},
  pages        = {031405},
  month        = SEP
}
% doi = {10.1103/PhysRevE.82.031405}
% url = {http://adsabs.harvard.edu/abs/2010PhRvE..82c1405M}

@ARTICLE{JeongICKX2007,
  author       = {Jeong, Unyong and Im, Sang~Hyuk and Camargo, Pedro~H.~C. and Kim, Jung~Hyun and Xia, Younan}, 
  title        = {Microscale fish bowls: a new class of latex particles with hollow interiors and engineered porous structures in their surfaces},
  journal      = LANG,
  publisher    = ACS,
  year         = {2007},
  volume       = {23},
  number       = {22},
  pages        = {10968-10975},
  month        = OCT
}
% doi = {10.1021/la702324q}
% url = {http://pubs.acs.org/doi/abs/10.1021/la702324q}

@ARTICLE{SevonkaevGM2008,
  author       = {Sevonkaev, Igor and Goia, Dan~V. and Matijevi{\'c}, Egon},
  title        = {Formation and structure of cubic particles of sodium magnesium fluoride (neighborite)},
  journal      = JoCaIS,
  publisher    = ES,
  year         = {2008},
  volume       = {317},
  number       = {1},
  pages        = {130-136},
  month        = JAN
}
% doi = {10.1016/j.jcis.2007.09.036}
% url = {http://www.sciencedirect.com/science/article/pii/S0021979707013872}

@ARTICLE{CuestaMR1997,
  author       = {Cuesta, J.~A. and {Mart{\'{\i}}nez-Rat{\'o}n}, Y.},
  title        = {Fundamental measure theory for mixtures of parallel hard cubes: I. General formalism},
  journal      = JoChemP,
  publisher    = AIoP,
  year         = {1997},
  volume       = {107},
  number       = {16},
  pages        = {6379-6389},
  month        = OCT
}
% doi = {10.1063/1.474298}
% url = {http://adsabs.harvard.edu/abs/1997JChPh.107.6379C}

@ARTICLE{MartinezRatonC1999,
  author       = {{Mart{\'{\i}}nez-Rat{\'o}n}, Y. and Cuesta, J.~A.},
  title        = {Fundamental measure theory for mixtures of parallel hard cubes: II. Phase behavior of the one-component fluid and of the binary mixture},
  journal      = JoChemP,
  publisher    = AIoP,
  year         = {1999},
  volume       = {111},
  number       = {1},
  pages        = {317-327},
  month        = JUL
}
% doi = {10.1063/1.479273}
% url = {http://adsabs.harvard.edu/abs/1999JChPh.111..317M}

@ARTICLE{HajiAkbariEKZPPMG2009,
  author       = {{Haji-Akbari}, A. and Engel, M. and Keys, A.~S. and Zheng, X. and Petschek, R.~G. and {Palffy-Muhoray}, P. and Glotzer, S.~C.},
  title        = {Disordered, quasicrystalline and crystalline phases of densely packed tetrahedra},
  journal      = NAT,
  publisher    = NPG,
  year         = {2009},
  volume       = {462},
  number       = {7274},
  pages        = {773-777},
  month        = DEC
}
% doi = {10.1038/nature08641}
% url = {http://adsabs.harvard.edu/abs/2009Natur.462..773H}

@ARTICLE{YinW1997,
  author       = {Yin, J.~S. and Wang, Z.~L.},
  title        = {Ordered self-assembling of tetrahedral oxide nanocrystals},
  journal      = PRL,
  publisher    = APS,
  year         = {1997},
  volume       = {79},
  number       = {13},
  pages        = {2570-2573},
  month        = SEP
}
% doi = {10.1103/PhysRevLett.79.2570}
% url = {http://adsabs.harvard.edu/abs/1997PhRvL..79.2570Y}

@ARTICLE{ZhangG2004,
  author       = {Zhang, Zhenli and Glotzer, Sharon~C.},
  title        = {Self-assembly of patchy particles},
  journal      = NL,
  publisher    = ACS,
  year         = {2004},
  volume       = {4},
  number       = {8},
  pages        = {1407-1413},
  month        = JUL
}
% doi = {10.1021/nl0493500}
% url = {http://pubs.acs.org/doi/abs/10.1021/nl0493500}

@ARTICLE{ChoYKJEYYP2007,
  author       = {Cho, {Young-Sang} and Yi, {Gi-Ra} and Kim, {Shin-Hyun} and Jeon, {Seog-Jin} and Elsesser, Mark~T. and Yu, Hyung~Kyun and Yang, {Seung-Man} and Pine, David~J.}, 
  title        = {Particles with coordinated patches or windows from oil-in-water emulsions},
  journal      = CoM,
  publisher    = ACS,
  year         = {2007},
  volume       = {19},
  number       = {13},
  pages        = {3183-3193},
  month        = MAY
}
% doi = {10.1021/cm070051w}
% url = {http://pubs.acs.org/doi/abs/10.1021/cm070051w}

@ARTICLE{Sciortino2008,
  author       = {Sciortino, F.},
  title        = {Gel-forming patchy colloids and network glass formers: thermodynamic and dynamic analogies},
  journal      = EPJB,
  publisher    = EDPS,
  year         = {2008},
  volume       = {64},
  number       = {3-4},
  pages        = {505-509},
  month        = AUG
}
% doi = {10.1140/epjb/e2008-00034-0}
% url = {http://adsabs.harvard.edu/abs/2008EPJB...64..505S}

@ARTICLE{BianchiLTZS2006,
  author       = {Bianchi, Emanuela and Largo, Julio and Tartaglia, Piero and Zaccarelli, Emanuela and Sciortino, Francesco},
  title        = {Phase diagram of patchy colloids: towards empty liquids},
  journal      = PRL,
  publisher    = APS,
  year         = {2006},
  volume       = {97},
  number       = {16},
  pages        = {168301},
  month        = OCT
}
% doi = {10.1103/PhysRevLett.97.168301}
% url = {http://prl.aps.org/abstract/PRL/v97/i16/e168301}

@ARTICLE{LiddellS2003,
  author       = {Liddell, C.~M. and Summers, C.~J.},
  title        = {Monodispersed ZnS dimers, trimers, and tetramers for lower symmetry photonic crystal lattices},
  journal      = AdvM,
  publisher    = WILEYVCH,
  year         = {2003},
  volume       = {15},
  number       = {20},
  pages        = {1715-1719},
  month        = OCT
}
% doi = {10.1002/adma.200305283}
% url = {http://onlinelibrary.wiley.com/doi/10.1002/adma.200305283/abstract}

@ARTICLE{ZerroukiBPCB2008,
  author       = {Zerrouki, D. and Baudry, J. and Pine, D. and Chaikin, P. and Bibette, J.},
  title        = {Chiral colloidal clusters},
  journal      = NAT,
  publisher    = NPG,
  year         = {2008},
  volume       = {455},
  number       = {7211},
  pages        = {380-382},
  month        = SEP
}
% doi = {10.1038/nature07237}
% url = {http://adsabs.harvard.edu/abs/2008Natur.455..380Z}

@ARTICLE{WensinkJ2009,
  author       = {Wensink, H.~H. and Jackson, G.},
  title        = {Generalized van der {W}aals theory for the twist elastic modulus and helical pitch of cholesterics},
  journal      = JoChemP,
  publisher    = AIoP,
  year         = {2009},
  volume       = {130},
  number       = {23},
  pages        = {234911},
  month        = JUN
}
% doi = {10.1063/1.3153348}
% url = {http://adsabs.harvard.edu/abs/2009JChPh.130w4911W}

@ARTICLE{SacannaICP2010,
  author       = {Sacanna, S. and Irvine, W.~T.~M. and Chaikin, P.~M. and Pine, D.~J.},
  title        = {Lock and key colloids},
  journal      = NAT,
  publisher    = NPG,
  year         = {2010},
  volume       = {464},
  number       = {7288},
  pages        = {575-578},
  month        = MAR
}
% doi = {10.1038/nature08906}
% url = {http://adsabs.harvard.edu/abs/2010Natur.464..575S}

@ARTICLE{CurtinA1986,
  author       = {Curtin, W.~A. and Ashcroft, N.~W.},
  title        = {Density-functional theory and freezing of simple liquids},
  journal      = PRL,
  publisher    = APS,
  year         = {1986},
  volume       = {56},
  number       = {26},
  pages        = {2775-2778},
  month        = JUN
}
% doi = {10.1103/PhysRevLett.56.2775}
% url = {http://adsabs.harvard.edu/abs/1986PhRvL..56.2775C}

@ARTICLE{LikosHLL2002,
  author       = {Likos, C.~N. and Hoffmann, N. and L{\"o}wen, H. and Louis, A.~A.},
  title        = {Exotic fluids and crystals of soft polymeric colloids},
  journal      = JoP:CM,
  publisher    = IoPP,
  year         = {2002},
  volume       = {14},
  number       = {33},
  pages        = {7681-7698},
  month        = AUG
}
% doi = {10.1088/0953-8984/14/33/309}
% url = {http://adsabs.harvard.edu/abs/2002JPCM...14.7681L}

@ARTICLE{LikosNL1994,
  author       = {Likos, C.~N. and Nemeth, Z.~T. and L{\"o}wen, H.},
  title        = {Density-functional theory of solid-to-solid isostructural transitions},
  journal      = JoP:CM,
  publisher    = IoPP,
  year         = {1994},
  volume       = {6},
  number       = {50},
  pages        = {10965-10975},
  month        = DEC
}
% doi = {10.1088/0953-8984/6/50/007}
% url = {http://adsabs.harvard.edu/abs/1994JPCM....610965L}

@INBOOK{Boyd2008K56,
  author       = {Boyd, Robert~W.},
  title        = {Nonlinear Optics},
  chapter      = {5.6: Nonlinear Optics of Liquid Crystals},
  publisher    = AP,
  year         = {2008},
  volume       = {1},
  series       = {},
  address      = AMST,
  edition      = {3},
  month        = APR,
  pages        = {271-276},
  isbn         = {978-0-123-69470-6}
}
% 640 Seiten

@ARTICLE{Rapaport2007,
  author       = {Rapaport, D.~C.},
  title        = {Microscale swimming: the molecular dynamics approach},
  journal      = PRL,
  publisher    = APS,
  year         = {2007},
  volume       = {99},
  number       = {23},
  pages        = {238101},
  month        = DEC
}
% doi = {10.1103/PhysRevLett.99.238101}
% url = {http://prl.aps.org/abstract/PRL/v99/i23/e238101}

@ARTICLE{AlmenarR2011,
  author       = {Almenar, L. and Rauscher, M.},
  title        = {Dynamics of colloids in confined geometries},
  journal      = JoP:CM,
  publisher    = IoPP,
  year         = {2011},
  volume       = {23},
  number       = {18},
  pages        = {184115},
  month        = APR
}
% doi = {10.1088/0953-8984/23/18/184115}
% url = {http://iopscience.iop.org/0953-8984/23/18/184115}

@BOOK{deGrootM1984,
  author       = {Groot, Sybren~Ruurds~de and Mazur, Peter},
  title        = {Non-Equilibrium Thermodynamics},
  publisher    = DP,
  year         = {1984},
  volume       = {},
  series       = {Dover Books on Physics and Chemistry},
  address      = NY,
  edition      = {1},
  month        = OCT,
  pages        = {544},
  isbn         = {0-486-64741-2}
}

@BOOK{Khalatnikov1989,
  author       = {Khalatnikov, I.~M.},
  title        = {An Introduction to the Theory of Superfluidity},
  publisher    = AW,
  year         = {1989},
  volume       = {23},
  series       = {Frontiers in Physics},
  address      = {Redwood City},
  edition      = {2},
  month        = MAR,
  pages        = {206},
  isbn         = {0-201-09505-X}
}

@BOOK{Reichl1998,
  author       = {Reichl, Linda~E.},
  title        = {A Modern Course in Statistical Physics},
  publisher    = WIL,
  year         = {1998},
  volume       = {},
  series       = {},
  address      = NY,
  edition      = {2nd},
  month        = MAR,
  pages        = {842},
  isbn         = {978-0-471-59520-5}
}

@ARTICLE{Risken1972,
  author       = {Risken, H.},
  title        = {Solutions of the {F}okker-{P}lanck equation in detailed balance},
  journal      = ZfP,
  publisher    = SPR,
  year         = {1972},
  volume       = {251},
  number       = {3},
  pages        = {231-243},
  month        = JUN
}
% doi = {10.1007/BF01379601}
% url = {http://adsabs.harvard.edu/abs/1972ZPhy..251..231R}

@ARTICLE{MartinPP1972,
  author       = {Martin, P.~C. and Parodi, O. and Pershan, P.~S.},
  title        = {Unified hydrodynamic theory for crystals, liquid crystals, and normal fluids},
  journal      = PRA,
  publisher    = APS,
  year         = {1972},
  volume       = {6},
  number       = {6},
  pages        = {2401-2420},
  month        = DEC
}
% doi = {10.1103/PhysRevA.6.2401}
% url = {http://adsabs.harvard.edu/abs/1972PhRvA...6.2401M}

@ARTICLE{GrahamH1971a,
  author       = {Graham, R. and Haken, H.},
  title        = {Fluctuations and stability of stationary non-equilibrium systems in detailed balance},
  journal      = ZfP,
  publisher    = SPR,
  year         = {1971},
  volume       = {245},
  number       = {2},
  pages        = {141-153},
  month        = APR
}
% doi = {10.1007/BF01402335}
% url = {http://adsabs.harvard.edu/abs/1971ZPhy..245..141G}

@ARTICLE{GrahamH1971b,
  author       = {Graham, R. and Haken, H.},
  title        = {Generalized thermodynamic potential for {M}ar\-koff systems in detailed balance and far from thermal equilibrium},
  journal      = ZfP,
  publisher    = SPR,
  year         = {1971},
  volume       = {243},
  number       = {3},
  pages        = {289-302},
  month        = JUN
}
% doi = {10.1007/BF01394858}
% url = {http://adsabs.harvard.edu/abs/1971ZPhy..243..289G}

@ARTICLE{Graham1974,
  author       = {Graham, R.},
  title        = {Hydrodynamic fluctuations near the convection instability},
  journal      = PRA,
  publisher    = APS,
  year         = {1974},
  volume       = {10},
  number       = {5},
  pages        = {1762-1784},
  month        = NOV
}
% doi = {10.1103/PhysRevA.10.1762}
% url = {http://adsabs.harvard.edu/abs/1974PhRvA..10.1762G}

@ARTICLE{tenHagenvTL2011,
  author       = {ten~Hagen, Borge and van~Teeffelen, Sven and L{\"o}wen, Hartmut},
  title        = {Brownian motion of a self-propelled particle},
  journal      = JoP:CM,
  publisher    = IoPP,
  year         = {2011},
  volume       = {23},
  number       = {19},
  pages        = {194119},
  month        = APR
}
% doi = {10.1088/0953-8984/23/19/194119}
% url = {http://iopscience.iop.org/0953-8984/23/19/194119}

@article{JouG2018,
  title={Coarse-graining for fast dynamics of order parameters in the phase-field model},
  author={Jou, D and Galenko, P K},
  journal={Philosophical Transactions of the Royal Society A: Mathematical, Physical and Engineering Sciences},
  volume={376},
  number={2113},
  pages={20170203},
  year={2018},
  publisher={The Royal Society Publishing}
}

@ARTICLE{WittkowskiLB2011,
  author       = {Wittkowski, Raphael  and L{\"o}wen, Hartmut and Brand, Helmut~R.},
  title        = {Polar liquid crystals in two spatial dimensions: the bridge from microscopic to macroscopic modeling},
  journal      = PRE,
  publisher    = APS,
  year         = {2011},
  volume       = {83},
  number       = {6},
  pages        = {061706},
  month        = JUN
}
% doi = {10.1103/PhysRevE.83.061706}
% url = {http://pre.aps.org/abstract/PRE/v83/i6/e061706}

@BOOK{Risken1996,
  author       = {Risken, Hannes},
  title        = {The Fokker-Planck Equation: Methods of Solution and Applications},
  publisher    = SPR,
  year         = {1996},
  volume       = {18},
  series       = {Springer Series in Synergetics},
  address      = BER,
  edition      = {3},
  month        = SEP,
  pages        = {474},
  isbn         = {3-540-61530-X}
}

@ARTICLE{HaertelL2010,
  author       = {H{\"a}rtel, A. and L{\"o}wen, H.},
  title        = {Fundamental measure density functional theory for hard spherocylinders in static and time-dependent aligning fields},
  journal      = JoP:CM,
  publisher    = IoPP,
  year         = {2010},
  volume       = {22},
  number       = {10},
  pages        = {104112},
  month        = MAR
}
% doi = {10.1088/0953-8984/22/10/104112}
% url = {http://adsabs.harvard.edu/abs/2010JPCM...22j4112H}

@ARTICLE{Wilson2007,
  author       = {Wilson, Mark~Richard},
  title        = {Molecular simulation of liquid crystals: progress towards a better understanding of bulk structure and the prediction of material properties},
  journal      = CSR,
  publisher    = RSCP,
  year         = {2007},
  volume       = {36},
  number       = {12},
  pages        = {1881-1888},
  month        = JUN
}
% doi = {10.1039/B612799C}
% url = {http://pubs.rsc.org/en/Content/ArticleLanding/2007/CS/b612799c}

@ARTICLE{CareC2005,
  author       = {Care, C.~M. and Cleaver, D.~J.},
  title        = {Computer simulation of liquid crystals},
  journal      = RoPiP,
  publisher    = IoPP,
  year         = {2005},
  volume       = {68},
  number       = {11},
  pages        = {2665-2700},
  month        = NOV
}
% doi = {10.1088/0034-4885/68/11/R04}
% url = {http://adsabs.harvard.edu/abs/2005RPPh...68.2665C}

@ARTICLE{NguyenW2005,
  author       = {Nguyen, Nam-Trung and Wu, Zhigang},
  title        = {Micromixers-a review},
  journal      = JoMaM,
  publisher    = IoPP,
  year         = {2005},
  volume       = {15},
  number       = {2},
  pages        = {R1-R16},
  month        = FEB
}
% doi = {10.1088/0960-1317/15/2/R01}
% url = {http://iopscience.iop.org/0960-1317/15/2/R01}

@ARTICLE{KimYY2005,
  author       = {Kim, J.~H. and Yoneya, M. and Yokoyama, H.},
  title        = {Fine bistable device of nematic liquid crystal realized on orientational surface patterns},
  journal      = MCaLC,
  publisher    = TF,
  year         = {2005},
  volume       = {433},
  number       = {1},
  pages        = {41-49},
  month        = JUN
}
% doi = {10.1080/15421400590955208}
% url = {http://www.informaworld.com/smpp/content~db=all~content=a725639728}

@ARTICLE{HoogboomRRN2006,
  author       = {Hoogboom, Johan and Rasing, Theo and Rowan, Alan~E. and Nolte, Roeland~J.~M.},
  title        = {LCD alignment layers. Controlling nematic domain properties},
  journal      = JoMC,
  publisher    = RSoC,
  year         = {2006},
  volume       = {16},
  number       = {14},
  pages        = {1305-1314},
  month        = APR
}
% doi = {10.1039/B510579J}
% url = {http://pubs.rsc.org/en/Content/ArticleLanding/2006/JM/b510579j}

@ARTICLE{BerardiMZ2008,
  author       = {Berardi, R. and Muccioli, L. and Zannoni, C.},
  title        = {Field response and switching times in biaxial nematics},
  journal      = JoChemP,
  publisher    = AIoP,
  year         = {2008},
  volume       = {128},
  number       = {2},
  pages        = {024905},
  month        = JAN
}
% doi = {10.1063/1.2815804}
% url = {http://adsabs.harvard.edu/abs/2008JChPh.128b4905B}

@ARTICLE{BraderK2011,
  author       = {Brader, J.~M. and Kr{\"u}ger, M.},
  title        = {Density profiles of a colloidal liquid at a wall under shear flow},
  journal      = MP,
  publisher    = TF,
  year         = {2011},
  volume       = {7},
  number       = {7},
  pages        = {1029-1041},
  month        = APR
}
% doi = {10.1080/00268976.2010.541889}
% url = {http://adsabs.harvard.edu/abs/2011MolPh.109.1029B}

@ARTICLE{LopezLeonKDVFN2011,
  author       = {{Lopez-Leon}, T. and Koning, V. and Devaiah, K.~B.~S. and Vitelli, V. and {Fernandez-Nieves}, A.},
  title        = {Frustrated nematic order in spherical geometries},
  journal      = NAT,
  publisher    = NPG,
  year         = {2011},
  volume       = {7},
  number       = {5},
  pages        = {1745-2473},
  month        = MAY
}
% doi = {10.1038/nphys1920}
% url = {http://dx.doi.org/10.1038/nphys1920}

@ARTICLE{Boyadzhiev1999,
  author       = {Boyadzhiev, Khristo~N.},
  title        = {Spirals and conchospirals in the flight of insects},
  journal      = CMJ,
  publisher    = MAoA,
  year         = {1999},
  volume       = {30},
  number       = {1},
  pages        = {23-31},
  month        = JAN
}
% url = {http://xahlee.info/SpecialPlaneCurves_dir/EquiangularSpiral_dir/_p/Khristo_Boyadzhiev_CMJ-99.pdf}

@ARTICLE{ChelakkotWG2010,
  author       = {Chelakkot, R. and Winkler, R.~G. and Gompper, G.},
  title        = {Migration of semiflexible polymers in microcapillary flow},
  journal      = EPL,
  publisher    = EPS,
  year         = {2010},
  volume       = {91},
  number       = {1},
  pages        = {14001},
  month        = JUL
}
% doi = {10.1209/0295-5075/91/14001}
% url = {http://adsabs.harvard.edu/abs/2010EL.....9114001C}

@ARTICLE{DunkelPZY2010,
  author       = {Dunkel, J. and Putz, V.~B. and Zaid, I.~M. and Yeomans, J.~M.},
  title        = {Swimmer-tracer scattering at low {R}eynolds number},
  journal      = SM,
  publisher    = RSoC,
  year         = {2010},
  volume       = {6},
  number       = {17},
  pages        = {4268-4276},
  month        = JUL
}
% doi = {10.1039/C0SM00164C}
% url = {http://pubs.rsc.org/en/Content/ArticleLanding/2010/SM/c0sm00164c}

@ARTICLE{LeptosGGPG2009,
  author       = {Leptos, K.~C. and Guasto, J.~S. and Gollub, J.~P. and Pesci, A.~I. and Goldstein, R.~E.},
  title        = {Dynamics of enhanced tracer diffusion in suspensions of swimming eukaryotic microorganisms},
  journal      = PRL,
  publisher    = APS,
  year         = {2009},
  volume       = {103},
  number       = {19},
  pages        = {198103},
  month        = NOV
}
% doi = {10.1103/PhysRevLett.103.198103}
% url = {http://adsabs.harvard.edu/abs/2009PhRvL.103s8103L}

@ARTICLE{RafaiJP2010,
  author       = {Rafa{\"i}, S. and Jibuti, L. and Peyla, P.},
  title        = {Effective viscosity of microswimmer suspensions},
  journal      = PRL,
  publisher    = APS,
  year         = {2010},
  volume       = {104},
  number       = {9},
  pages        = {098102},
  month        = MAR
}
% doi = {10.1103/PhysRevLett.104.098102}
% url = {http://adsabs.harvard.edu/abs/2010PhRvL.104i8102R}

@ARTICLE{BrandP1981a,
  author       = {Brand, H. and Pleiner, H.},
  title        = {Hydrodynamics of biaxial discotics},
  journal      = PRA,
  publisher    = APS,
  year         = {1981},
  volume       = {24},
  number       = {5},
  pages        = {2777-2788},
  month        = NOV
}
% doi = {10.1103/PhysRevA.24.2777}
% url = {http://adsabs.harvard.edu/abs/1981PhRvA..24.2777B}

@ARTICLE{BrandP1981b,
  author       = {Brand, H. and Pleiner, H.},
  title        = {Broken symmetries and hydrodynamics of superfluid ^{3}P_{2}-neutron-star matter},
  journal      = PRD,
  publisher    = APS,
  year         = {1981},
  volume       = {24},
  number       = {12},
  pages        = {3048-3057},
  month        = DEC
}
% doi = {10.1103/PhysRevD.24.3048}
% url = {http://adsabs.harvard.edu/abs/1981PhRvD..24.3048B}

@ARTICLE{Forster1974,
  author       = {Forster, D.},
  title        = {Hydrodynamics and correlation functions in ordered systems: nematic liquid crystals},
  journal      = AoP,
  publisher    = ELS,
  year         = {1974},
  volume       = {84},
  number       = {1-2},
  pages        = {505-534},
  month        = MAY
}
% doi = {10.1016/0003-4916(74)90314-5}
% url = {http://adsabs.harvard.edu/abs/1974AnPhy..84..505F}

@ARTICLE{KadanoffM1963,
  author       = {{Kadanoff}, L.~P. and {Martin}, P.~C.},
  title        = {Hydrodynamic equations and correlation functions},
  journal      = AoP,
  publisher    = ELS,
  year         = {1963},
  volume       = {24},
  number       = {1},
  pages        = {419-469},
  month        = OCT
}
% doi = {10.1016/0003-4916(63)90078-2}
% url = {http://adsabs.harvard.edu/abs/1963AnPhy..24..419K}

@ARTICLE{HohenbergM1965,
  author       = {{Hohenberg}, P.~C. and {Martin}, P.~C.},
  title        = {Microscopic theory of superfluid helium},
  journal      = AoP,
  publisher    = ELS,
  year         = {1965},
  volume       = {34},
  number       = {2},
  pages        = {291-359},
  month        = SEP
}
% doi = {10.1016/0003-4916(65)90280-0}
% url = {http://adsabs.harvard.edu/abs/1965AnPhy..34..291H}

@ARTICLE{BrandP1982b,
  author       = {Brand, H. and Pleiner, H.},
  title        = {Linearized hydrodynamics of $^{3}$He-A$_{1}$: correlation functions and hydrodynamic parameters},
  journal      = JdP,
  publisher    = EDPS,
  year         = {1982},
  volume       = {43},
  number       = {2},
  pages        = {369-380},
  month        = FEB
}
% doi = {10.1051/jphys:01982004302036900}
% url = {http://jphys.journaldephysique.org}

@ARTICLE{BohliusBP2004,
  author       = {{Bohlius}, S. and {Brand}, H.~R. and {Pleiner}, H.},
  title        = {Macroscopic dynamics of uniaxial magnetic gels},
  journal      = PRE,
  publisher    = APS,
  year         = {2004},
  volume       = {70},
  number       = {6},
  pages        = {061411},
  month        = DEC
}
% doi = {10.1103/PhysRevE.70.061411}
% url = {http://adsabs.harvard.edu/abs/2004PhRvE..70f1411B}

@ARTICLE{BrandDG1979,
  author       = {{Brand}, H. and {D{\"o}rfle}, M. and {Graham}, R.},
  title        = {Hydrodynamic parameters and correlation functions of superfluid $^{3}$He},
  journal      = AoP,
  publisher    = ELS,
  year         = {1979},
  volume       = {119},
  number       = {2},
  pages        = {434-479},
  month        = JUN
}
% doi = {10.1016/0003-4916(79)90194-5}
% url = {http://adsabs.harvard.edu/abs/1979AnPhy.119..434B}

@ARTICLE{Liu1981,
  author       = {Liu, M.},
  title        = {Hydrodynamic theory of biaxial nematics},
  journal      = PRA,
  publisher    = APS,
  year         = {1981},
  volume       = {24},
  number       = {5},
  pages        = {2720-2726},
  month        = NOV
}
% doi = {10.1103/PhysRevA.24.2720}
% url = {http://adsabs.harvard.edu/abs/1981PhRvA..24.2720L}

@ARTICLE{Liu1979,
  author       = {Liu, M.},
  title        = {Hydrodynamic theory near the nematic-smectic-A transition},
  journal      = PRA,
  publisher    = APS,
  year         = {1979},
  volume       = {19},
  number       = {5},
  pages        = {2090-2094},
  month        = MAY
}
% doi = {10.1103/PhysRevA.19.2090}
% url = {http://adsabs.harvard.edu/abs/1979PhRvA..19.2090L}

@ARTICLE{Brand1986c,
  author       = {Brand, H.~R.},
  title        = {Macroscopic dynamics near the nematic-columnar transition in liquid crystals},
  journal      = PRA,
  publisher    = APS,
  year         = {1986},
  volume       = {33},
  number       = {1},
  pages        = {643-646},
  month        = JAN
}
% doi = {10.1103/PhysRevA.33.643}
% url = {http://adsabs.harvard.edu/abs/1986PhRvA..33..643B}

@ARTICLE{BrandP1982,
  author       = {Brand, H. and Pleiner, H.},
  title        = {Theory of flow alignment in biaxial nematics and nematic discotics},
  journal      = JdP,
  publisher    = EDPS,
  year         = {1982},
  volume       = {43},
  number       = {6},
  pages        = {853-858},
  month        = JUN
}
% doi = {10.1051/jphys:01982004306085300}
% url = {http://jphys.journaldephysique.org/index.php?option=com_article&access=standard&Itemid=129&url=/articles/jphys/abs/1982/06/jphys_1982__43_6_853_0/jphys_1982__43_6_853_0.html}

@ARTICLE{PleinerB1985,
  author       = {Pleiner, H. and Brand, H.~R.},
  title        = {Macroscopic behaviour of a biaxial mixture of uniaxial nematics versus hydrodynamics of a biaxial nematic liquid crystal},
  journal      = JdP,
  publisher    = EDPS,
  year         = {1985},
  volume       = {46},
  number       = {4},
  pages        = {615-620},
  month        = APR
}
% doi = {10.1051/jphys:01985004604061500}
% url = {http://jphys.journaldephysique.org/index.php?option=com_article&access=standard&Itemid=129&url=/articles/jphys/abs/1985/04/jphys_1985__46_4_615_0/jphys_1985__46_4_615_0.html}

@ARTICLE{CladisCB1985,
  author       = {Cladis, P.~E. and Couder, Y. and Brand, H.~R.},
  title        = {Phase winding and flow alignment in freely suspended films of smectic-C liquid crystals},
  journal      = PRL,
  publisher    = APS,
  year         = {1985},
  volume       = {55},
  number       = {27},
  pages        = {2945-2948},
  month        = DEC
}
% doi = {10.1103/PhysRevLett.55.2945}
% url = {http://adsabs.harvard.edu/abs/1985PhRvL..55.2945C}

@ARTICLE{PleinerLB2002,
  author       = {Pleiner, Harald and Liu, Mario and Brand, Helmut~R.},
  title        = {Convective nonlinearities for the orientational tensor order parameter in polymeric systems},
  journal      = RA,
  publisher    = SPR,
  year         = {2002},
  volume       = {41},
  number       = {4},
  pages        = {375-382},
  month        = MAY
}
% doi = {10.1007/s00397-002-0237-z}
% url = {http://www.springerlink.com/content/4er6lg1bh78m2guh}

@ARTICLE{GuerraPMC2009,
  author       = {Guerra, M. and Pinto, R.~M. and Martins, R. and Casquilho, J.~P.},
  title        = {On the Effect of a Rotating Magnetic Field or Sample on the Bend Freedericksz Critical Field in Nematic Slabs},
  journal      = MCaLC,
  publisher    = TF,
  year         = {2009},
  volume       = {508},
  number       = {1},
  pages        = {337-347},
  month        = AUG
}
% doi = {10.1080/15421400903065853}
% url = {http://www.tandfonline.com/toc/gmcl20/508/1}

@ARTICLE{LeeCS2008,
  author       = {Lee, Wei and Chen, H.~Y. and Shih, Y.~C.},
  title        = {Reduced dc offset and faster dynamic response in a carbon-nanotube-impregnated liquid-crystal display},
  journal      = JotSfID,
  publisher    = SfID,
  year         = {2008},
  volume       = {16},
  number       = {7},
  pages        = {733-741},
  month        = JUL
}
% doi = {10.1889/1.2953480}
% url = {http://dx.doi.org/10.1889/1.2953480}

@ARTICLE{AchimWL2011,
  author       = {Achim, Cristian~V. and Wittkowski, Raphael and L{\"o}wen, Hartmut},
  title        = {Stability of liquid crystalline phases in the phase-field-crystal model},
  journal      = PRE,
  publisher    = APS,
  year         = {2011},
  volume       = {83},
  number       = {6},
  pages        = {061712},
  month        = JUN
}
% doi = {http://link.aps.org/doi/10.1103/PhysRevE.83.061712}
% url = {http://pre.aps.org/abstract/PRE/v83/i6/e061712}

@Article{Lutsko2011,
  author  = {Lutsko, James F.},
  title   = {Communication: A dynamical theory of homogeneous nucleation for colloids and macromolecules},
  journal = JoChemP,
  year    = {2011},
  volume  = {135},
  number  = {16},
  pages   = {161101}
}
% doi     = {10.1063/1.3657400},
%  eprint  = {https://doi.org/10.1063/1.3657400},
% url     = { https://doi.org/10.1063/1.3657400},
}

@article{OnsagerM1953,
  title={Fluctuations and irreversible processes},
  author={Onsager, Lars and Machlup, Stefan},
  journal=PR,
  volume={91},
  number={6},
  pages={1505},
  year={1953},
  publisher={APS}
}


@ARTICLE{AokiOMYT2007,
  author       = {Aoki, M. and Ozaki, R. and Moritake, H. and Yoshino, K. and Toda, K.},
  title        = {Viscosity Response Measurement of Nematic Liquid Crystal under Application of Electric Field Using Shear Horizontal Wave},
  journal      = MCaLC,
  publisher    = TF,
  year         = {2007},
  volume       = {477},
  number       = {1},
  pages        = {607-618},
  month        = OCT
}
% doi = {10.1080/15421400701686785}
% url = {http://www.tandfonline.com/toc/gmcl20/477/1}

@ARTICLE{AbrahamEtAl2007,
  author       = {{Abraham, J. et~al. [The Pierre Auger collaboration]}},
  title        = {Correlation of the Highest-Energy Cosmic Rays with Nearby Extragalactic Objects},
  journal      = SCI,
  publisher    = AAftAoS,
  year         = {2007},
  volume       = {318},
  number       = {938},
  pages        = {938-943},
  month        = NOV
}
% doi = {10.1126/science.1151124}
% url = {} Abreu, P. and Aglietta, M. and Aguirre, C. and Allard, D. and Allekotte, I. and Allen, J. and Allison, P. and Alvarez, C. and et~al.

@ARTICLE{Naegele1996,
  author       = {N{\"a}gele, G.},
  title        = {On the dynamics and structure of charge-stabilized suspensions},
  journal      = PRS,
  publisher    = ELS,
  year         = {1996},
  volume       = {272},
  number       = {1},
  pages        = {215-372},
  month        = JUL
}
% doi = {10.1016/0370-1573(95)00078-X}
% url = {http://adsabs.harvard.edu/abs/1996PhR...272..215N}

@BOOK{Gardiner2009,
  author       = {Gardiner, Crispin~W.},
  title        = {Handbook of Stochastic Methods: for Physics, Chemistry and the Natural Sciences},
  publisher    = SPR,
  year         = {2009},
  volume       = {13},
  series       = {Springer Series in Synergetics},
  address      = BER,
  edition      = {4},
  month        = JAN,
  pages        = {447},
  isbn         = {978-3-540-70712-7}
}

@BOOK{Adomian2001,
  author       = {Adomian, George},
  title        = {Nonlinear Stochastic Systems Theory and Application to Physics},
  publisher    = KAP,
  year         = {2001},
  volume       = {46},
  series       = {Mathematics and its Applications},
  address      = DOR,
  edition      = {1},
  month        = NOV,
  pages        = {248},
  isbn         = {978-1-402-00338-7}
}

@ARTICLE{Langevin1908,
  author       = {Langevin, Paul},
  title        = {Sur la th{\'e}orie du mouvement brownien},
  journal      = CRdlAdS,
  publisher    = FAoS,
  year         = {1908},
  volume       = {146},
  number       = {1},
  pages        = {530-533},
  month        = JAN
}

@BOOK{Protter2010,
  author       = {Protter, Philip~E.},
  title        = {Stochastic Integration and Differential Equations},
  publisher    = SPR,
  year         = {2010},
  volume       = {21},
  series       = {Stochastic Modelling and Applied Probability},
  address      = BER,
  edition      = {2},
  month        = DEC,
  pages        = {419},
  isbn         = {978-3-642-05560-7}
}

@BOOK{CoffeyKW2004,
  author       = {Coffey, William~T. and Kalmykov, Yuri~P. and Waldron, John~T.},
  title        = {The Langevin Equation: With Applications to Stochastic Problems in Physics, Chemistry and Electrical Engineering},
  publisher    = WSP,
  year         = {2004},
  volume       = {14},
  series       = {World Scientific Series in Contemporary Chemical Physics},
  address      = SIN,
  edition      = {2},
  month        = MAR,
  pages        = {678},
  isbn         = {978-9-812-38462-1}
}

@BOOK{HoldenOUZ2009,
  author       = {Holden, Helge and {\O}ksendal, Bernt~Karsten and Uboe, Jan and Zhang, Tusheng},
  title        = {Stochastic Partial Differential Equations: A Modeling, White Noise Approach},
  publisher    = SPR,
  year         = {2009},
  volume       = {},
  series       = {Universitext},
  address      = BER,
  edition      = {2},
  month        = DEC,
  pages        = {305},
  isbn         = {978-0-387-89487-4}
}

@BOOK{Oksendal2003,
  author       = {{\O}ksendal, Bernt~Karsten},
  title        = {Stochastic Differential Equations: An Introduction with Applications},
  publisher    = SPR,
  year         = {2003},
  volume       = {},
  series       = {Universitext},
  address      = BER,
  edition      = {6},
  month        = JUL,
  pages        = {374},
  isbn         = {978-3-540-04758-2}
}

@ARTICLE{Maruyama1955,
  author       = {Maruyama, Gisiro},
  title        = {Continuous Markov processes and stochastic equations},
  journal      = {Rendiconti del Circolo Matematico di Palermo},
  publisher    = {Circolo Matematico di Palermo},
  year         = {1955},
  volume       = {4},
  number       = {1},
  pages        = {48-90},
  month        = JAN
}
% doi = {10.1007/BF02846028}
% url = {http://www.springerlink.com/content/g31773v1nk758282/}

@ARTICLE{Milstein1975,
  author       = {Milstein, G.~N.},
  title        = {Approximate integration of stochastic differential equations},
  journal      = {Theory of Probability and Its Applications},
  publisher    = {SIAM},
  year         = {1975},
  volume       = {19},
  number       = {3},
  pages        = {557-562},
  month        = JUL
}

@INBOOK{FaettiFFG1985,
  author       = {Faetti, S. and Festa, C. and Fronzoni, L. and Grigolini, P.},
  title        = {Memory Function Approaches to Stochastic Problems in Condensed Matter},
  chapter      = {10: Experimental Investigation on the Effect of Multiplicative Noise by Means of Electric Circuits},
  publisher    = WIL,
  year         = {1985},
  volume       = {62},
  series       = {Advances in Chemical Physics},
  address      = NY,
  editor       = {Myron~W. Evans and Paolo Grigolini and Giuseppe~Pastori Parravicini},
  edition      = {1},
  month        = JAN,
  pages        = {445-475},
  isbn         = {978-0-470-14331-5}
}
% doi = {10.1002/9780470142868.ch10}
% url = {http://dx.doi.org/10.1002/9780470142868.ch10}

@ARTICLE{PalacciCBYB2010,
  author       = {Palacci, J. and {Cottin-Bizonne}, C. and Ybert, C. and Bocquet, L.},
  title        = {Sedimentation and effective temperature of active colloidal suspensions},
  journal      = PRL,
  publisher    = APS,
  year         = {2010},
  volume       = {105},
  number       = {8},
  pages        = {088304},
  month        = AUG
}
% doi = {10.1103/PhysRevLett.105.088304}
% url = {http://adsabs.harvard.edu/abs/2010PhRvL.105h8304P}

@ARTICLE{MatsumotoN1998,
  author       = {Matsumoto, Makoto and Nishimura, Takuji},
  title        = {Mersenne twister: a 623-di\-men\-sio\-nal\-ly equidistributed uniform pseudorandom number generator},
  journal      = {ACM Transactions on Modeling and Computer Simulation},
  publisher    = {Association for Computing Machinery},
  year         = {1998},
  volume       = {8},
  number       = {1},
  pages        = {3-30},
  month        = JAN
}
% doi = {10.1145/272991.272995}
% url = {http://portal.acm.org/citation.cfm?id=272995}
 
@ARTICLE{MonaghanL1985,
  author       = {Monaghan, J.~J. and Lattanzio, J.~C.},
  title        = {A refined particle method for astrophysical problems},
  journal      = AaA,
  publisher    = EDPS,
  year         = {1985},
  volume       = {149},
  number       = {1},
  pages        = {135-143},
  month        = AUG
}
% url = {http://adsabs.harvard.edu/abs/1985A%26A...149..135M}

@ARTICLE{HernquistK1989,
  author       = {Hernquist, L. and Katz, N.},
  title        = {TREESPH - A unification of SPH with the hierarchical tree method},
  journal      = AJSS,
  publisher    = IoPP,
  year         = {1989},
  volume       = {70},
  number       = {1},
  pages        = {419-446},
  month        = JUN
}
% doi = {10.1086/191344}
% url = {http://adsabs.harvard.edu/abs/1989ApJS...70..419H}

@ARTICLE{Lucy1977,
  author       = {Lucy, L.~B.},
  title        = {A numerical approach to the testing of the fission hypothesis},
  journal      = AJ,
  publisher    = IoPP,
  year         = {1977},
  volume       = {82},
  number       = {1},
  pages        = {1013-1024},
  month        = DEC
}
% doi = {10.1086/112164}
% url = {http://adsabs.harvard.edu/abs/1977AJ.....82.1013L}

@ARTICLE{GingoldM1977,
  author       = {Gingold, R.~A. and Monaghan, J.~J.},
  title        = {Smoothed particle hydrodynamics - Theory and application to non-spherical stars},
  journal      = MNotRAS,
  publisher    = WIL,
  year         = {1977},
  volume       = {181},
  number       = {1},
  pages        = {375-389},
  month        = NOV
}
% url = {http://adsabs.harvard.edu/abs/1977MNRAS.181..375G}

@BOOK{LiuL2003,
  author       = {Liu, G.~R. and Liu, M.~B.},
  title        = {Smoothed Particle Hydrodynamics: A Meshfree Particle Method},
  publisher    = WSP,
  year         = {2003},
  volume       = {},
  series       = {},
  address      = SIN,
  edition      = {1},
  month        = OCT,
  pages        = {449},
  isbn         = {981-238-456-1}
}

@ARTICLE{MakinoD2004,
  author       = {Makino, M. and Doi, M.},
  title        = {Brownian motion of a particle of general shape in {N}ewtonian fluid},
  journal      = JotPSoJ,
  publisher    = PSoJ,
  year         = {2004},
  volume       = {73},
  number       = {10},
  pages        = {2739},
  month        = OCT
}
% doi = {10.1143/JPSJ.73.2739}
% url = {http://adsabs.harvard.edu/abs/2004JPSJ...73.2739M}

@article{delaHozBKLS2014,
  title={Unpolarized states and hidden polarization},
  author={de la Hoz, P and Bj{\"o}rk, G and Klimov, Andrei B and Leuchs, G and S{\'a}nchez-Soto, LL},
  journal={Physical Review A},
  volume={90},
  number={4},
  pages={043826},
  year={2014},
  publisher={APS}
}

@article{StuhlmullerNEdlHS2018,
  title={Structural nonequilibrium forces in driven colloidal systems},
  author={Stuhlm{\"u}ller, Nico CX and Eckert, Tobias and {de las Heras}, Daniel and Schmidt, Matthias},
  journal=PRL,
  volume={121},
  number={9},
  pages={098002},
  year={2018},
  publisher={APS}
}

@article{delasHerasS2018,
  title={Velocity Gradient Power Functional for {Brownian} Dynamics},
  author={{de las Heras}, Daniel and Schmidt, Matthias},
  journal=PRL,
  volume={120},
  number={2},
  pages={028001},
  year={2018},
  publisher={APS}
}

@article{SchmidtB2013,
  title={Power functional theory for {B}rownian dynamics},
  author={Schmidt, Matthias and Brader, Joseph~M},
  journal=JoChemP,
  volume={138},
  number={21},
  pages={214101},
  year={2013},
  publisher={AIP}
}

@article{BraderS2015,
  title={Power functional theory for the dynamic test particle limit},
  author={Brader, Joseph M and Schmidt, Matthias},
  journal=JoP:CM,
  volume={27},
  number={19},
  pages={194106},
  year={2015},
  publisher={IOP Publishing}
}

@article{KrinningerS2019,
  title={Power functional theory for active {Brownian} particles: general formulation and power sum rules},
  author={Krinninger, Philip and Schmidt, Matthias},
  journal=JoChemP,
  volume={150},
  number={7},
  pages={074112},
  year={2019},
  publisher={AIP Publishing}
}

@book{Blum2012,
  title={Density matrix theory and applications},
  author={Blum, Karl},
  series={Springer Series on Atomic, Optical, and Plasma Physics},
  volume={64},
  year={2012},
  publisher={Springer},
  address=BER
}
% DOI: 10.1007/978-3-642-20561-3

@ARTICLE{FernandesdlT2002,
  author       = {Fernandes, M.~X. and {Garc\'{\i}a de la Torre}, J.},
  title        = {Brownian dynamics simulation of rigid particles of arbitrary shape in external fields},
  journal      = BJ,
  publisher    = ELS,
  year         = {2002},
  volume       = {83},
  number       = {6},
  pages        = {3039-3048},
  month        = DEC
}
% doi = {10.1016/S0006-3495(02)75309-5}
% url = {http://adsabs.harvard.edu/abs/2002BpJ....83.3039F}

@ARTICLE{MontgomeryB1977,
  author       = {Montgomery, J.~A. and Berne, B.~J.},
  title        = {The effects of hydrodynamic interactions on translational and rotational relaxation},
  journal      = JoChemP,
  publisher    = AIoP,
  year         = {1977},
  volume       = {67},
  number       = {10},
  pages        = {4589-4596},
  month        = NOV
}
% doi = {10.1063/1.434619}
% url = {http://adsabs.harvard.edu/abs/1977JChPh..67.4589M}

@ARTICLE{DickinsonAMC1985,
  author       = {Dickinson, Eric and Allison, Stuart~A. and McCammon, J.~Andrew},
  title        = {Brownian dynamics with rotation-translation coupling},
  journal      = {Journal of the Chemical Society, Faraday Transactions 2: Molecular and Chemical Physics},
  publisher    = RSCP,
  year         = {1985},
  volume       = {81},
  number       = {4},
  pages        = {591-601},
  month        = APR
}
% doi = {10.1039/F29858100591}
% url = {http://pubs.rsc.org/en/Content/ArticleLanding/1985/F2/F29858100591}

@ARTICLE{CondiffD1966,
  author       = {Condiff, D.~W. and Dahler, J.~S.},
  title        = {Brownian Motion of Polyatomic Molecules: The Coupling of Rotational and Translational Motions},
  journal      = JoChemP,
  publisher    = AIoP,
  year         = {1966},
  volume       = {44},
  number       = {1},
  pages        = {3988-4004},
  month        = MAY
}
% doi = {10.1063/1.1726561}
% url = {http://adsabs.harvard.edu/abs/1966JChPh..44.3988C}

@BOOK{GoldsteinPS2006,
  author       = {Goldstein, Herbert and {Poole Jr.}, Charles~P. and {Safko Sr.}, John~L.},
  title        = {Klassische Mechanik},
  publisher    = WILEYVCH,
  year         = {2006},
  series       = {Lehrbuch Physik},
  address      = BER,
  edition      = {3},
  month        = JUL,
  pages        = {704},
  isbn         = {978-3-527-40589-3}
}

@BOOK{Butcher2008,
  author       = {Butcher, John~Charles},
  title        = {Numerical Methods for Ordinary Differential Equations},
  publisher    = WIL,
  series       = {},
  volume       = {},
  year         = {2008},
  address      = {Chichester},
  edition      = {2},
  month        = MAY,
  pages        = {482},
  isbn         = {0-470-72335-1}
}
% EAN: 978-0-470-72335-7

@MISC{Brenner1963a_1964d,
  author       = {Brenner, Howard},
  howpublished = {Chemical Engineering Science \textbf{18}, 1 (Jan. 1963); -- \textbf{18}, 557 (Aug. 1963); -- \textbf{19}, 519 (Aug. 1964); -- \textbf{19}, 599 (Sep. 1964); -- \textbf{19}, 631 (Sep. 1964); -- \textbf{19}, 703 (Oct. 1964)}
}

@MISC{ChwangWu1974_1976,
  author       = {Chwang, A.~T. and Wu, T.~Y.},
  howpublished = {Journal of Fluid Mechanics \textbf{63}, 607 (Feb. 1974); -- \textbf{67}, 787 (Feb. 1975); -- \textbf{75}, 677 (Feb. 1976)}
}

@ARTICLE{HoffmannWHW2009,
  author       = {Hoffmann, Martin and Wagner, Claudia~S. and Harnau, Ludger and Wittemann, Alexander},
  title        = {3D Brownian diffusion of submicron-sized particle clusters},
  journal      = NANO,
  publisher    = ACS,
  year         = {2009},
  volume       = {3},
  number       = {10},
  pages        = {3326-3334},
  month        = SEP
}
% doi = {10.1021/nn900902b}
% url = {http://pubs.acs.org/doi/abs/10.1021/nn900902b}

@ARTICLE{tenHagenWL2011,
  author       = {ten~Hagen, Borge and Wittkowski, Raphael and L{\"o}wen, Hartmut},
  title        = {Brownian dynamics of a self-propelled particle in shear flow},
  journal      = PRE,
  publisher    = APS,
  year         = {2011},
  volume       = {84},
  number       = {3},
  pages        = {031105},
  month        = SEP
}
% doi = {10.1103/PhysRevE.84.031105}
% url = {http://link.aps.org/doi/10.1103/PhysRevE.84.031105}

@ARTICLE{LauL2007,
  author       = {{Lau}, A.~W.~C. and {Lubensky}, T.~C.},
  title        = {State-dependent diffusion: thermodynamic consistency and its path integral formulation},
  journal      = PRE,
  publisher    = APS,
  year         = {2007},
  volume       = {76},
  number       = {1},
  pages        = {011123},
  month        = JUL
}
% doi = {10.1103/PhysRevE.76.011123}
% url = {http://adsabs.harvard.edu/abs/2007PhRvE..76a1123L}

@ARTICLE{GrohD1996,
  author       = {Groh, B. and Dietrich, S.},
  title        = {Density-functional theory for the freezing of {S}tockmayer fluids},
  journal      = PRE,
  publisher    = APS,
  year         = {1996},
  volume       = {54},
  number       = {2},
  pages        = {1687-1697},
  month        = AUG
}
% doi = {10.1103/PhysRevE.54.168}
% url = {http://link.aps.org/doi/10.1103/PhysRevE.54.1687}

@article{SuJH2021,
  title={Inertia-induced nucleation-like motility-induced phase separation},
  author={Su, Jie and Jiang, Huijun and Hou, Zhonghuai},
  journal={New Journal of Physics},
  volume={23},
  number={1},
  pages={013005},
  year={2021},
  publisher={IOP Publishing}
}

@ARTICLE{GrohD1994,
  author       = {Groh, B. and Dietrich, S.},
  title        = {Ferroelectric phase in {S}tockmayer fluids},
  journal      = PRE,
  publisher    = APS,
  year         = {1994},
  volume       = {50},
  number       = {5},
  pages        = {3814-3833},
  month        = NOV
}
% doi = {10.1103/PhysRevE.50.3814}
% url = {http://link.aps.org/doi/10.1103/PhysRevE.50.3814}

@ARTICLE{VolpeBVKB2011,
  author       = {Volpe, Giovanni and Buttinoni, Ivo and Vogt, Dominik and K{\"u}mmerer, {Hans-J{\"u}rgen} and Bechinger, Clemens},
  title        = {Microswimmers in patterned environments},
  journal      = SM,
  publisher    = RSoC,
  year         = {2011},
  volume       = {7},
  number       = {19},
  pages        = {8810-8815},
  month        = AUG
}
% doi = {10.1039/C1SM05960B}
% url = {http://pubs.rsc.org/en/content/articlelanding/2011/sm/c1sm05960b}

@ARTICLE{LanconBLO2001,
  author       = {{Lan{\c c}on}, P. and {Batrouni}, G. and {Lobry}, L. and {Ostrowsky}, N.},
  title        = {Drift without flux: Brownian walker with a space-dependent diffusion coefficient},
  journal      = EPL,
  publisher    = EPS,
  year         = {2001},
  volume       = {54},
  number       = {1},
  pages        = {28-34},
  month        = APR
}
% doi = {10.1209/epl/i2001-00103-6}
% url = {http://adsabs.harvard.edu/abs/2001EL.....54...28L}

@ARTICLE{LanconBLO2002,
  author       = {{Lan{\c c}on}, P. and {Batrouni}, G. and {Lobry}, L. and {Ostrowsky}, N.},
  title        = {Brownian walker in a confined geometry leading to a space-dependent diffusion coefficient},
  journal      = PA,
  publisher    = ELS,
  year         = {2002},
  volume       = {304},
  number       = {1-2},
  pages        = {65-76},
  month        = FEB
}
% doi = {10.1016/S0378-4371(01)00510-6}
% url = {http://adsabs.harvard.edu/abs/2002PhyA..304...65L}

@ARTICLE{ChoudharyLEL2011,
  author       = {{Choudhary}, M.~Ajmal and {Li}, D. and {Emmerich}, H. and {L{\"o}wen}, H.},
  title        = {{DDFT} calibration and investigation of an anisotropic phase-field crystal model},
  journal      = JoP:CM,
  publisher    = IoPP,
  year         = {2011},
  volume       = {23},
  number       = {26},
  pages        = {265005},
  month        = JUL
}
% doi = {10.1088/0953-8984/23/26/265005}
% url = {http://adsabs.harvard.edu/abs/2011JPCM...23z5005A}

@ARTICLE{WittkowskiLB2011b,
  author       = {Wittkowski, Raphael and L{\"o}wen, Hartmut and Brand, Helmut~R.},
  title        = {Microscopic and macroscopic theories for the dynamics of polar liquid crystals},
  journal      = PRE,
  publisher    = APS,
  year         = {2011},
  volume       = {84},
  number       = {4},
  pages        = {041708},
  month        = OCT
}
% doi = {10.1103/PhysRevE.84.041708}
% url = {http://link.aps.org/doi/10.1103/PhysRevE.84.041708}

@ARTICLE{BawdenPBF1936,
  author       = {{Bawden}, F.~C. and {Pirie}, N.~W. and {Bernal}, J.~D. and {Fankuchen}, I.},
  title        = {Liquid crystalline substances from virus-infected plants},
  journal      = NAT,
  publisher    = NPG,
  year         = {1936},
  volume       = {138},
  number       = {3503},
  pages        = {1051-1052},
  month        = DEC
}
% doi = {10.1038/1381051a0}
% url = {http://adsabs.harvard.edu/abs/1936Natur.138.1051B}

@ARTICLE{OhnesorgeLW1993,
  author       = {{Ohnesorge}, R. and {L{\"o}wen}, H. and {Wagner}, H.},
  title        = {Density distribution in a hard-sphere crystal},
  journal      = EPL,
  publisher    = EPS,
  year         = {1993},
  volume       = {22},
  number       = {4},
  pages        = {245-249},
  month        = MAY
}
% doi = {10.1209/0295-5075/22/4/002}
% url = {http://adsabs.harvard.edu/abs/1993EL.....22..245O}

@ARTICLE{CerdavdV1983,
  author       = {Cerda, C.~M. and Ven, T.~G.~M.~van~de},
  title        = {Translational diffusion of axisymmetrical particles in shear flow},
  journal      = JoCaIS,
  publisher    = ES,
  year         = {1983},
  volume       = {93},
  number       = {1},
  pages        = {54-62},
  month        = MAY
}
% doi = {10.1016/0021-9797(83)90383-1}
% url = {http://www.sciencedirect.com/science/article/pii/0021979783903831}

@ARTICLE{HolzerBRZ2010,
  author       = {Holzer, Lukas and Bammert, Jochen and Rzehak, Roland and Zimmermann, Walter},
  title        = {Dynamics of a trapped Brownian particle in shear flows},
  journal      = PRE,
  publisher    = APS,
  year         = {2010},
  volume       = {81},
  number       = {4},
  pages        = {041124},
  month        = APR
}
% doi = {10.1103/PhysRevE.81.041124}
% url = {http://link.aps.org/doi/10.1103/PhysRevE.81.041124}

@ARTICLE{OhtaO2009,
  author       = {{Ohta}, T. and {Ohkuma}, T.},
  title        = {Deformable self-propelled particles},
  journal      = PRL,
  publisher    = APS,
  year         = {2009},
  volume       = {102},
  number       = {15},
  pages        = {154101},
  month        = APR
}
% doi = {10.1103/PhysRevLett.102.154101}
% url = {http://adsabs.harvard.edu/abs/2009PhRvL.102o4101O}

@ARTICLE{MitchellK2006,
  author       = {Mitchell, James~G. and Kogure, Kazuhiro},
  title        = {Bacterial motility: links to the environment and a driving force for microbial physics},
  journal      = {FEMS Microbiology Ecology},
  publisher    = WIL,
  year         = {2006},
  volume       = {55},
  number       = {1},
  pages        = {3-16},
  month        = JAN
}
% doi = {10.1111/j.1574-6941.2005.00003.x}
% url = {http://dx.doi.org/10.1111/j.1574-6941.2005.00003.x}

@ARTICLE{KochS2011,
  author       = {{Koch}, D.~L. and {Subramanian}, G.},
  title        = {Collective hydrodynamics of swimming microorganisms: living fluids},
  journal      = ARoFM,
  publisher    = AR,
  year         = {2011},
  volume       = {43},
  number       = {1},
  pages        = {637-659},
  month        = JAN
}
% doi = {10.1146/annurev-fluid-121108-145434}
% url = {http://adsabs.harvard.edu/abs/2011AnRFM..43..637K}

@ARTICLE{MuhuriRR2007,
  author       = {{Muhuri}, S. and {Rao}, M. and {Ramaswamy}, S.},
  title        = {Shear-flow induced isotropic-to-nematic transition in a suspension of active filaments},
  journal      = EPL,
  publisher    = EPS,
  year         = {2007},
  volume       = {78},
  number       = {4},
  pages        = {48002},
  month        = MAY
}
% doi = {10.1209/0295-5075/78/48002}
% url = {http://adsabs.harvard.edu/abs/2007EL.....7848002M}

@article{PraetoriusV2011,
  title={A phase field crystal approach for particles in a flowing solvent},
  author={Praetorius, Simon and Voigt, Axel},
  journal={Macromolecular Theory and Simulations},
  volume={20},
  number={7},
  pages={541--547},
  year={2011},
  publisher={Wiley Online Library}
}

@article{CapriniM2021,
  title={Inertial self-propelled particles},
  author={Caprini, Lorenzo and {Marini Bettolo Marconi}, Umberto},
  journal=JoChemP,
  volume={154},
  number={2},
  pages={024902},
  year={2021},
  publisher={AIP Publishing LLC}
}

@article{AzizG2004,
  title={Causal diffusion and the survival of charge fluctuations in nuclear collisions},
  author={Aziz, Mohamed Abdel and Gavin, Sean},
  journal={Physical Review C},
  volume={70},
  number={3},
  pages={034905},
  year={2004},
  publisher={APS}
}

@ARTICLE{CatesFMOY2008,
  author       = {{Cates}, M.~E. and {Fielding}, S.~M. and {Marenduzzo}, D. and {Orlandini}, E. and {Yeomans}, J.~M.},
  title        = {Shearing active gels close to the isotropic-nematic transition},
  journal      = PRL,
  publisher    = APS,
  year         = {2008},
  volume       = {101},
  number       = {6},
  pages        = {068102},
  month        = AUG
}
% doi = {10.1103/PhysRevLett.101.068102}
% url = {http://adsabs.harvard.edu/abs/2008PhRvL.101f8102C}

@ARTICLE{LobaskinLK2008,
  author       = {{Lobaskin}, V. and {Lobaskin}, D. and {Kuli{\'c}}, I.~M.},
  title        = {Brownian dynamics of a microswimmer},
  journal      = EPJST,
  publisher    = EDPS,
  year         = {2008},
  volume       = {157},
  number       = {1},
  pages        = {149-156},
  month        = APR
}
% doi = {10.1140/epjst/e2008-00637-7}
% url = {http://adsabs.harvard.edu/abs/2008EPJST.157..149L}

@ARTICLE{DunkelZ2009,
  author       = {{Dunkel}, J. and {Zaid}, I.~M.},
  title        = {Noisy swimming at low {R}eynolds numbers},
  journal      = PRE,
  publisher    = APS,
  year         = {2009},
  volume       = {80},
  number       = {2},
  pages        = {021903},
  month        = AUG
}
% doi = {10.1103/PhysRevE.80.021903}
% url = {http://adsabs.harvard.edu/abs/2009PhRvE..80b1903D}

@ARTICLE{PeruaniM2007, 
  author       = {{Peruani}, F. and {Morelli}, L.~G.},
  title        = {Self-propelled particles with fluctuating speed and direction of motion in two dimensions},
  journal      = PRL,
  publisher    = APS,
  year         = {2007},
  volume       = {99},
  number       = {1},
  pages        = {010602},
  month        = JUL
}
% doi = {10.1103/PhysRevLett.99.010602}
% url = {http://adsabs.harvard.edu/abs/2007PhRvL..99a0602P}

@article{XuTQ2015,
  title={A variational approach to thin film hydrodynamics of binary mixtures},
  author={Xu, Xinpeng and Thiele, Uwe and Qian, Tiezheng},
  journal=JoP:CM,
  volume={27},
  number={8},
  pages={085005},
  year={2015},
  publisher={IOP Publishing}
}

@article{Kawasaki2006,
  title={Principle of maximum entropy and reduced dynamics},
  author={Kawasaki, Kyozi},
  journal={Journal of Statistical Physics},
  volume={123},
  number={4},
  pages={711--740},
  year={2006},
  publisher={Springer}
}


@article{AlandHLV2015,
  title={A mechanistic collective cell model for epithelial colony growth and contact inhibition},
  author={Aland, Sebastian and Hatzikirou, Haralambos and Lowengrub, John and Voigt, Axel},
  journal=BJ,
  volume={109},
  number={7},
  pages={1347--1357},
  year={2015},
  publisher={Elsevier}
}


@article{vanTeeffelenAL2013,
  title={Vacancy diffusion in colloidal crystals as determined by dynamical density-functional theory and the phase-field-crystal model},
  author={van Teeffelen, Sven and Achim, Cristian Vasile and L{\"o}wen, Hartmut},
  journal=PRE,
  volume={87},
  number={2},
  pages={022306},
  year={2013},
  publisher={APS}
}


@ARTICLE{PahlavanS2011, 
  author       = {{Pahlavan}, A.~A. and {Saintillan}, D.},
  title        = {Instability regimes in flowing suspensions of swimming micro-organisms},
  journal      = PoF,
  publisher    = AIoP,
  year         = {2011},
  volume       = {23},
  number       = {1},
  pages        = {011901},
  month        = JAN
}
% doi = {10.1063/1.3529411}
% url = {http://adsabs.harvard.edu/abs/2011PhFl...23a1901P}

@BOOK{Balescu1997,
  author       = {Balescu, Radu},
  title        = {Statistical Dynamics: Matter out of Equilibrium},
  publisher    = ICP,
  year         = {1997},
  address      = {London},
  edition      = {1},
  month        = APR,
  pages        = {330},
  isbn         = {1-860-94045-5}
}

@BOOK{WaleckaF2003,
  author       = {Walecka, John~Dirk and Fetter, Alexander~L.},
  title        = {Quantum Theory of Many-Particle Systems},
  publisher    = DP,
  year         = {2003},
  address      = {Mineola},
  edition      = {1},
  month        = JUN,
  pages        = {640},
  isbn         = {0-486-42827-3}
}

@BOOK{Somov2000,
  author       = {Somov, Boris~V.},
  title        = {Cosmic Plasma Physics},
  publisher    = KAP,
  year         = {2000},
  volume       = {251},
  series       = {Astrophysics and Space Science Library},
  address      = DOR,
  edition      = {1},
  month        = SEP,
  pages        = {672},
  isbn         = {0-792-36512-7}
}

@BOOK{Kleinert2009,
  author       = {Kleinert, Hagen},
  title        = {Path Integrals in Quantum Mechanics, Statistics, Polymer Physics, and Financial Markets},
  publisher    = WSP,
  year         = {2009},
  address      = SIN,
  edition      = {5},
  month        = JUL,
  pages        = {1579},
  isbn         = {9-814-27356-2}
}
% EAN: 9789814273565

@BOOK{HinrichsenW2004,
  author       = {Hinrichsen, Haye and Wolf, Dietrich~E.},
  title        = {The Physics of Granular Media},
  publisher    = WIL,
  year         = {2004},
  volume       = {7},
  series       = {Nonlinear \& Complex Systems},
  address      = WEI,
  edition      = {1},
  month        = OCT,
  pages        = {500},
  isbn         = {3-527-40373-6}
}
% EAN: 9783527403738

@BOOK{Bouchoule1999,
  author       = {Bouchoule, Andre},
  title        = {Dusty Plasmas: Physics, Chemistry, and Technological Impact in Plasma Processing},
  publisher    = WIL,
  year         = {1999},
  address      = CHI,
  edition      = {1},
  month        = JUL,
  pages        = {408},
  isbn         = {0-471-97386-6}
}
% EAN: 9780471973867

@ARTICLE{Archer2005,
  author       = {{Archer}, A.~J.},
  title        = {Dynamical density functional theory: binary phase-separating colloidal fluid in a cavity},
  journal      = JoP:CM,
  publisher    = IoPP,
  year         = {2005},
  volume       = {17},
  number       = {10},
  pages        = {1405-1427},
  month        = MAR
}
% doi = {10.1088/0953-8984/17/10/001}
% url = {http://adsabs.harvard.edu/abs/2005JPCM...17.1405A}

@ARTICLE{LaugaS2005,
  author       = {{Lauga}, E. and {Squires}, T.~M.},
  title        = {Brownian motion near a partial-slip boundary: a local probe of the no-slip condition},
  journal      = PoF,
  publisher    = AIoP,
  year         = {2005},
  volume       = {17},
  number       = {10},
  pages        = {103102},
  month        = OCT
}
% doi = {10.1063/1.2083748}
% url = {http://adsabs.harvard.edu/abs/2005PhFl...17j3102L}

@BOOK{Satoh2010,
  author       = {Satoh, Akira},
  title        = {Introduction to Practice of Molecular Simulation: Molecular Dynamics, {M}onte {C}arlo, {B}rownian Dynamics, Lattice {B}oltzmann and Dissipative Particle Dynamics},
  publisher    = ELS,
  year         = {2010},
  volume       = {7},
  series       = EI,
  address      = LON,
  edition      = {1},
  month        = DEC,
  pages        = {322},
  isbn         = {0-123-85148-3}
}
% EAN: 9780123851482

@ARTICLE{BowickCSS1994,
  author       = {{Bowick}, M.~J. and {Chandar}, L. and {Schiff}, E.~A. and {Srivastava}, A.~M.},
  title        = {The cosmological Kibble mechanism in the laboratory: string formation in liquid crystals},
  journal      = SCI,
  publisher    = AAftAoS,
  year         = {1994},
  volume       = {263},
  number       = {5149},
  pages        = {943-945},
  month        = FEB
}
% doi = {10.1126/science.263.5149.943}
% url = {http://adsabs.harvard.edu/abs/1994Sci...263..943B}

@ARTICLE{TiernoGPS2008b,
  author       = {Tierno, Pietro and Golestanian, Ramin and Pagonabarraga, Ignacio and Sagu{\'e}s, Francesc},
  title        = {Magnetically actuated colloidal microswimmers},
  journal      = JoPChemB,
  publisher    = ACS,
  year         = {2008},
  volume       = {112},
  number       = {51},
  pages        = {16525-16528},
  month        = NOV
}
% doi = {10.1021/jp808354n}
% url = {http://pubs.acs.org/doi/abs/10.1021/jp808354n}

@ARTICLE{Purcell1977,
  author       = {Purcell, E.~M.},
  title        = {Life at low {R}eynolds number},
  journal      = AJoP,
  publisher    = AAoPT,
  year         = {1977},
  volume       = {45},
  number       = {1},
  pages        = {3-11},
  month        = JAN
}
% doi = {10.1119/1.10903}
% url = {http://www.citeulike.org/group/2000/article/1342642}

@ARTICLE{BelovsC2009,
  author       = {{Belovs}, M. and {C{\v e}bers}, A.},
  title        = {Ferromagnetic microswimmer},
  journal      = PRE,
  publisher    = APS,
  year         = {2009},
  volume       = {79},
  number       = {5},
  pages        = {051503},
  month        = MAY
}
% doi = {10.1103/PhysRevE.79.051503}
% url = {http://adsabs.harvard.edu/abs/2009PhRvE..79e1503B}

@ARTICLE{QinBXHM2007,
  author       = {Qin, Lidong and Banholzer, Matthew~J. and Xu, Xiaoyang and Huang, Ling and Mirkin, Chad~A.},
  title        = {Rational design and synthesis of catalytically driven nanorotors},
  journal      = JotACS,
  publisher    = ACS,
  year         = {2007},
  volume       = {129},
  number       = {48},
  pages        = {14870-14871},
  month        = NOV
}
% doi = {10.1021/ja0772391}
% url = {http://pubs.acs.org/doi/abs/10.1021/ja0772391}

@ARTICLE{PaxtonEtAl2004,
  author       = {Paxton, Walter~F. and Kistler, Kevin~C. and Olmeda, Christine~C. and Sen, Ayusman and St.~Angelo, Sarah~K. and Cao, Yanyan and Mallouk, Thomas~E. and Lammert, Paul~E. and Crespi, Vincent~H.},
  title        = {Catalytic nanomotors: autonomous movement of striped nanorods},
  journal      = JotACS,
  publisher    = ACS,
  year         = {2004},
  volume       = {126},
  number       = {41},
  pages        = {13424-13431},
  month        = SEP
}
% doi = {10.1021/ja047697z}
% url = {http://pubs.acs.org/doi/abs/10.1021/ja047697z}

@ARTICLE{LammertPB1996,
  author       = {Lammert, P.~E. and Prost, J. and Bruinsma, R.},
  title        = {Ion drive for vesicles and cells},
  journal      = JoTB,
  publisher    = ELS,
  year         = {1996},
  volume       = {178},
  number       = {4},
  pages        = {387-391},
  month        = FEB
}
% doi = {10.1006/jtbi.1996.0035}
% url = {http://www.sciencedirect.com/science/article/pii/S0022519396900351}

@ARTICLE{SaintillanS2008,
  author       = {Saintillan, David and Shelley, Michael~J.},
  title        = {Instabilities and pattern formation in active particle suspensions: kinetic theory and continuum simulations},
  journal      = PRL,
  publisher    = APS,
  year         = {2008},
  volume       = {100},
  number       = {17},
  pages        = {178103},
  month        = APR
}
% doi = {10.1103/PhysRevLett.100.178103}
% url = {http://link.aps.org/doi/10.1103/PhysRevLett.100.178103}

@ARTICLE{KimB2004,
  author       = {{Kim}, M.~J. and {Breuer}, K.~S.},
  title        = {Enhanced diffusion due to motile bacteria},
  journal      = PoF,
  publisher    = AIoP,
  year         = {2004},
  volume       = {16},
  number       = {9},
  pages        = {L78-L81},
  month        = SEP
}
% doi = {10.1063/1.1787527}
% url = {http://adsabs.harvard.edu/abs/2004PhFl...16L..78K}

@ARTICLE{HatwalneRRS2004,
  author       = {{Hatwalne}, Y. and {Ramaswamy}, S. and {Rao}, M. and {Simha}, R.~A.},
  title        = {Rheology of active-particle suspensions},
  journal      = PRL,
  publisher    = APS,
  year         = {2004},
  volume       = {92},
  number       = {11},
  pages        = {118101},
  month        = MAR
}
% doi = {10.1103/PhysRevLett.92.118101}
% url = {http://adsabs.harvard.edu/abs/2004PhRvL..92k8101H}

@article{Kawasaki1966,
  title={Diffusion constants near the critical point for time-dependent {I}sing models. I},
  author={Kawasaki, Kyozi},
  journal=PR,
  volume={145},
  number={1},
  pages={224},
  year={1966},
  publisher={APS}
}

@article{GouyetPDM2003,
  title={Description of far-from-equilibrium processes by mean-field lattice gas models},
  author={Gouyet, J-F and Plapp, Mathis and Dieterich, W and Maass, P},
  journal=AiP,
  volume={52},
  number={6},
  pages={523--638},
  year={2003},
  publisher={Taylor \& Francis}
}

@ARTICLE{EnculescuS2011,
  author       = {{Enculescu}, M. and {Stark}, H.},
  title        = {Active colloidal suspensions exhibit polar order under gravity},
  journal      = PRL,
  publisher    = APS,
  year         = {2011},
  volume       = {107},
  number       = {5},
  pages        = {058301},
  month        = JUL
}
% doi = {10.1103/PhysRevLett.107.058301}
% url = {http://adsabs.harvard.edu/abs/2011PhRvL.107e8301E}

@ARTICLE{MarconiT2000,
  author       = {{Marini Bettolo Marconi}, Umberto and Tarazona, Pedro},
  title        = {Dynamic density functional theory of fluids},
  journal      = JoP:CM,
  publisher    = IoPP,
  year         = {2000},
  volume       = {12},
  number       = {8A},
  pages        = {413-418},
  month        = FEB
}
% doi = {10.1088/0953-8984/12/8A/356}
% url = {http://adsabs.harvard.edu/abs/2000JPCM...12..413M}

@ARTICLE{Rauscher2010,
  author       = {{Rauscher}, M.},
  title        = {{DDFT} for Brownian particles and hydrodynamics},
  journal      = JoP:CM,
  publisher    = IoPP,
  year         = {2010},
  volume       = {22},
  number       = {36},
  pages        = {J4109},
  month        = SEP
}
% doi = {10.1088/0953-8984/22/36/364109}
% url = {http://adsabs.harvard.edu/abs/2010JPCM...22J4109R}

@ARTICLE{RauscherDKP2007,
  author       = {{Rauscher}, M. and {Dom{\'{\i}}nguez}, A. and {Kr{\"u}ger}, M. and {Penna}, F.},
  title        = {A dynamic density functional theory for particles in a flowing solvent},
  journal      = JoChemP,
  publisher    = AIoP,
  year         = {2007},
  volume       = {127},
  number       = {24},
  pages        = {244906},
  month        = DEC
}
% doi = {10.1063/1.2806094}
% url = {http://adsabs.harvard.edu/abs/2007JChPh.127x4906R}

@article{WilczekTGKCT2015,
  title={Modelling pattern formation in dip-coating experiments},
  author={Wilczek, Markus and Tewes, Walter BH and Gurevich, Svetlana V and K{\"o}pf, Michael H and Chi, LF and Thiele, Uwe},
  journal={Mathematical Modelling of Natural Phenomena},
  volume={10},
  number={4},
  pages={44--60},
  year={2015},
  publisher={EDP Sciences}
}

@article{FrusawaH2000,
  title={On the controversy over the stochastic density functional equations},
  author={Frusawa, H and Hayakawa, R},
  journal={Journal of Physics A: Mathematical and General},
  volume={33},
  number={15},
  pages={L155},
  year={2000},
  publisher={IOP Publishing}
}

@incollection{Boda2014,
  title={Monte {C}arlo simulation of electrolyte solutions in biology: in and out of equilibrium},
  author={Boda, Dezs{\H{o}}},
  booktitle={Annual Reports in Computational Chemistry},
  volume={10},
  pages={127-163},
  year={2014},
  publisher={Elsevier}
}

@article{ChavanisS2011b,
  title={Exact analytical solution of the collapse of self-gravitating {B}rownian particles and bacterial populations at zero temperature},
  author={Chavanis, Pierre-Henri and Sire, Cl{\'e}ment},
  journal=PRE,
  volume={83},
  number={3},
  pages={031131},
  year={2011},
  publisher={APS}
}

@article{GaoCX2017,
  title={Dielectric Decrement in a Charging Microcapacitor},
  author={Gao, Haixia and Chang, Yanmei and Xiao, Changming},
  journal={Chemistry Letters},
  volume={46},
  number={8},
  pages={1258--1261},
  year={2017},
  publisher={The Chemical Society of Japan}
}


@article{GiavazziSVC2016,
  title={Structure and dynamics of concentration fluctuations in a non-equilibrium dense colloidal suspension},
  author={Giavazzi, Fabio and Savorana, Giovanni and Vailati, Alberto and Cerbino, Roberto},
  journal=SM,
  volume={12},
  number={31},
  pages={6588-6600},
  year={2016},
  publisher={Royal Society of Chemistry}
}

@article{DiawM2017,
  title={A viscous quantum hydrodynamics model based on dynamic density functional theory},
  author={Diaw, Abdourahmane and Murillo, Michael S},
  journal={Scientific Reports},
  volume={7},
  number={1},
  pages={15352},
  year={2017},
  publisher={Nature Publishing Group}
}

@article{EvertsSEvdHvdBvdR2017,
  title={Colloid--oil--water-interface interactions in the presence of multiple salts: charge regulation and dynamics},
  author={Everts, Jeffrey C and Samin, Sela and Elbers, NA and van der Hoeven, Jessi ES and van Blaaderen, Alfons and van Roij, Ren{\'e}},
  journal={Physical Chemistry Chemical Physics},
  volume={19},
  number={22},
  pages={14345--14357},
  year={2017},
  publisher={Royal Society of Chemistry}
}


@article{DiawM2016,
  title={A Dynamic Density Functional Theory Approach to Diffusion in White Dwarfs and Neutron Star Envelopes},
  author={Diaw, Abdourahmane and Murillo, Michael Sean},
  journal={Astrophysical Journal},
  volume={829},
  number={1},
  pages={16},
  year={2016},
  publisher={IOP Publishing}
}

@article{DiawM2015,
  title={Generalized hydrodynamics model for strongly coupled plasmas},
  author={Diaw, Abdourahmane and Murillo, Michael Sean},
  journal=PRE,
  volume={92},
  number={1},
  pages={013107},
  year={2015},
  publisher={APS}
}

@article{EdisonM2010,
  title={Dynamic mean field theory of condensation and evaporation processes for fluids in porous materials: Application to partial drying and drying},
  author={Edison, JR and Monson, PA},
  journal={Faraday Discussions},
  volume={146},
  pages={167--184},
  year={2010},
  publisher={Royal Society of Chemistry}
}

@article{KrinningerSB2017,
  title={Erratum: Nonequilibrium Phase Behavior from Minimization of Free Power Dissipation [{Phys. Rev. Lett}. 117, 208003 (2016)]},
  author={Krinninger, Philip and Schmidt, Matthias and Brader, Joseph M},
  journal=PRL,
  volume={119},
  number={2},
  pages={029902},
  year={2017},
  publisher={APS}
}

@article{KruegerR2007,
  title={Colloid-colloid and colloid-wall interactions in driven suspensions},
  author={Kr{\"u}ger, Matthias and Rauscher, Markus},
  journal=JoChemP,
  volume={127},
  number={3},
  pages={034905},
  year={2007},
  publisher={AIP}
}

@article{Kawasaki2000,
  title={Theoretical methods dealing with slow dynamics},
  author={Kawasaki, Kyozi},
  journal=JoP:CM,
  volume={12},
  number={29},
  pages={6343},
  year={2000},
  publisher={IOP Publishing}
}

@article{AerovK2014,
  title={Driven colloidal suspensions in confinement and density functional theory: Microstructure and wall-slip},
  author={Aerov, Artem A and Kr{\"u}ger, Matthias},
  journal=JoChemP,
  volume={140},
  number={9},
  pages={094701},
  year={2014},
  publisher={AIP}
}

@article{ZimmermannSL2016,
  title={Flow of colloidal solids and fluids through constrictions: dynamical density functional theory versus simulation},
  author={Zimmermann, Urs and Smallenburg, Frank and L{\"o}wen, Hartmut},
  journal=JoP:CM,
  volume={28},
  number={24},
  pages={244019},
  year={2016},
  publisher={IOP Publishing}
}

@article{PhillipsD2005,
  title={Thermal disorder, fluctuations, growth and fragmentation of finite one-dimensional atomic chains},
  author={Phillips, James M and Dash, JG},
  journal={Journal of Statistical Physics},
  volume={120},
  number={3-4},
  pages={721--735},
  year={2005},
  publisher={Springer}
}

@article{MickelJB2011,
  title={Transport, phase transitions, and wetting in micro/nanochannels: A phase field/{DDFT} approach},
  author={Mickel, Walter and Joly, Laurent and Biben, Thierry},
  journal=JoChemP,
  volume={134},
  number={9},
  pages={094105},
  year={2011},
  publisher={AIP}
}

@article{vanTeeffelenLL2008,
  title={Crystallization of magnetic dipolar monolayers: a density functional approach},
  author={van Teeffelen, Sven and L{\"o}wen, Hartmut and Likos, Christos N},
  journal=JoP:CM,
  volume={20},
  number={40},
  pages={404217},
  year={2008},
  publisher={IOP Publishing}
}

@article{KohlIBML2012,
  title={Microscopic theory for anisotropic pair correlations in driven binary mixtures},
  author={Kohl, Matthias and Ivlev, Alexei V and Brandt, Philip and Morfill, Gregor E and L{\"o}wen, Hartmut},
  journal=JoP:CM,
  volume={24},
  number={46},
  pages={464115},
  year={2012},
  publisher={IOP Publishing}
}

@article{Iwamatsu2009,
  title={Minimum free-energy path of homogenous nucleation from the phase-field equation},
  author={Iwamatsu, Masao},
  journal=JoChemP,
  volume={130},
  number={24},
  pages={244507},
  year={2009},
  publisher={AIP}
}

@article{MarconiM2011,
  title={Dynamics of fluid mixtures in nanospaces},
  author={{Marini Bettolo Marconi}, Umberto and Melchionna, Simone},
  journal=JoChemP,
  volume={134},
  number={6},
  pages={064118},
  year={2011},
  publisher={AIP}
}

@article{Monson2011,
  title={Fluids confined in porous materials: Towards a unified understanding of thermodynamics and dynamics},
  author={Monson, Peter A},
  journal={Chemie Ingenieur Technik},
  volume={83},
  number={1-2},
  pages={143--151},
  year={2011},
  publisher={Wiley Online Library}
}

@article{NoldGYSK2017,
  title={Pseudospectral methods for density functional theory in bounded and unbounded domains},
  author={Nold, Andreas and Goddard, Benjamin D and Yatsyshin, Peter and Savva, Nikos and Kalliadasis, Serafim},
  journal={Journal of Computational Physics},
  volume={334},
  pages={639--664},
  year={2017},
  publisher={Elsevier}
}

@article{Marconi2011,
  title={Non-local kinetic theory of inhomogeneous liquid mixtures},
  author={{Marini Bettolo Marconi}, Umberto},
  journal={Molecular Physics},
  volume={109},
  number={7-10},
  pages={1265--1274},
  year={2011},
  publisher={Taylor \& Francis}
}

@incollection{Monson2009,
  title={Dynamic Mean Field Theory for Fluids Confined in Porous Materials: Application to an Inkbottle Pore Geometry},
  author={Monson, PA},
  booktitle={Characterisation of Porous Solids VIII: Proceedings of the 8th International Symposium on the Characterisation of Porous Solids},
  editor = {Seaton, Nigel and Reinoso, Francisco R and Llewellyn, Philip and Kaskel, Stefan},
  number={318},
  year={2009}
}


@article{LichtnerK2013,
  title={Spinodal decomposition of a binary magnetic fluid confined to a surface},
  author={Lichtner, Ken and Klapp, Sabine HL},
  journal=PRE,
  volume={88},
  number={3},
  pages={032301},
  year={2013},
  publisher={APS}
}

@article{Ram2014,
  title={Equilibrium theory of molecular fluids: Structure and freezing transitions},
  author={Ram, Jokhan},
  journal={Physics Reports},
  volume={538},
  number={4},
  pages={121--185},
  year={2014},
  publisher={Elsevier}
}


@article{KierlikLRT2011,
  title={Spontaneous imbibition in a slit pore: a lattice--gas dynamic mean field study},
  author={Kierlik, Edouard and Leoni, Fabio and Rosinberg, Martin Luc and Tarjus, Gilles},
  journal=MP,
  volume={109},
  number={7-10},
  pages={1143--1157},
  year={2011},
  publisher={Taylor \& Francis}
}

@article{NeuhausSL2013b,
  title={Crystallization induced by multiple seeds: Dynamical density functional approach},
  author={Neuhaus, T and Schmiedeberg, M and L{\"o}wen, H},
  journal=PRE,
  volume={88},
  number={6},
  pages={062316},
  year={2013},
  publisher={APS}
}

@article{GrzeticWS2014,
  title={Statistical dynamics of classical systems: A self-consistent field approach},
  author={Grzetic, Douglas J and Wickham, Robert A and Shi, An-Chang},
  journal=JoChemP,
  volume={140},
  number={24},
  pages={244907},
  year={2014},
  publisher={AIP}
}

@article{OlaisLM2015,
  title={Non-equilibrium theory of arrested spinodal decomposition},
  author={Olais-Govea, Jos{\'e} Manuel and L{\'o}pez-Flores, Leticia and Medina-Noyola, Magdaleno},
  journal=JoChemP,
  volume={143},
  number={17},
  pages={174505},
  year={2015},
  publisher={AIP Publishing}
}

@article{BleibelDOD2014,
  title={Capillary attraction induced collapse of colloidal monolayers at fluid interfaces},
  author={Bleibel, J and Dominguez, A and Oettel, M and Dietrich, S},
  journal=SM,
  volume={10},
  number={23},
  pages={4091--4109},
  year={2014},
  publisher={Royal Society of Chemistry}
}

@article{BleibelDO2015,
  title={{3D hydrodynamic interactions lead to divergences in 2D diffusion}},
  author={Bleibel, Johannes and Dom{\'\i}nguez, Alvaro and Oettel, Martin},
  journal=JoP:CM,
  volume={27},
  number={19},
  pages={194113},
  year={2015},
  publisher={IOP Publishing}
}

@article{AerovK2015,
  title={Theory of rheology in confinement},
  author={Aerov, Artem A and Kr{\"u}ger, Matthias},
  journal=PRE,
  volume={92},
  number={4},
  pages={042301},
  year={2015},
  publisher={APS}
}

@article{GrawitterS2018b,
  title={Correction: Feedback control of photoresponsive fluid interfaces},
  author={Grawitter, Josua and Stark, Holger},
  journal=SM,
  volume={14},
  number={35},
  pages={7287--7287},
  year={2018},
  publisher={Royal Society of Chemistry}
}

@article{LutskoN2016,
  title={Mechanism for the stabilization of protein clusters above the solubility curve},
  author={Lutsko, James F and Nicolis, Gr{\'e}goire},
  journal=SM,
  volume={12},
  number={1},
  pages={93--98},
  year={2016},
  publisher={Royal Society of Chemistry}
}

@article{XuHY2007,
  title={Dynamic density functional theory based on equation of state},
  author={Xu, Hui and Honglai, Liu and Ying, Hu},
  journal={Chemical Engineering Science},
  volume={62},
  number={13},
  pages={3494--3501},
  year={2007},
  publisher={Elsevier}
}

@article{HeinrichsDMF2004,
  title={Static and time dependent density functional theory with internal degrees of freedom: Merits and limitations demonstrated for the {P}otts model},
  author={Heinrichs, S and Dieterich, W and Maass, P and Frisch, HL},
  journal={Journal of Statistical Physics},
  volume={114},
  number={3-4},
  pages={1115--1125},
  year={2004},
  publisher={Springer}
}

@article{Chavanis2015,
  title={Generalized stochastic {Fokker-Planck} equations},
  author={Chavanis, Pierre-Henri},
  journal={Entropy},
  volume={17},
  number={5},
  pages={3205--3252},
  year={2015},
  publisher={Multidisciplinary Digital Publishing Institute}
}

@article{Titulaer1978,
  title={A systematic solution procedure for the {F}okker-{P}lanck equation of a {B}rownian particle in the high-friction case},
  author={Titulaer, Urbanus M},
  journal={Physica A: Statistical Mechanics and its Applications},
  volume={91},
  number={3-4},
  pages={321--344},
  year={1978},
  publisher={Elsevier}
}

@article{Titulaer1980,
  title={The {C}hapman-{E}nskog procedure as a form of degenerate perturbation theory},
  author={Titulaer, UM},
  journal={Physica A: Statistical Mechanics and its Applications},
  volume={100},
  number={2},
  pages={234--250},
  year={1980},
  publisher={Elsevier}
}

@article{XuALDB2018,
  title={Interaction of proteins with polyelectrolytes: Comparison of theory to experiment},
  author={Xu, Xiao and Angioletti-Uberti, Stefano and Lu, Yan and Dzubiella, Joachim and Ballauff, Matthias},
  journal={Langmuir},
  volume={35},
  number={16},
  pages={5373--5391},
  year={2018},
  publisher={ACS Publications}
}

@article{WittmannSSL2017,
  title={Phase diagram of two-dimensional hard rods from fundamental mixed measure density functional theory},
  author={Wittmann, Ren{\'e} and Sitta, Christoph E and Smallenburg, Frank and L{\"o}wen, Hartmut},
  journal=JoChemP,
  volume={147},
  number={13},
  pages={134908},
  year={2017},
  publisher={AIP Publishing}
}

@article{BaskaranGL2016,
  title={Energy stable multigrid method for local and non-local hydrodynamic models for freezing},
  author={Baskaran, Arvind and Guan, Zhen and Lowengrub, John},
  journal={Computer Methods in Applied Mechanics and Engineering},
  volume={299},
  pages={22--56},
  year={2016},
  publisher={Elsevier}
}

@article{LeoniKRT2011,
  title={Spontaneous imbibition in disordered porous solids: A theoretical study of helium in silica aerogels},
  author={Leoni, F and Kierlik, E and Rosinberg, ML and Tarjus, G},
  journal={Langmuir},
  volume={27},
  number={13},
  pages={8160-8170},
  year={2011},
  publisher={ACS Publications}
}

@article{BiervR2007,
  title={Relaxation dynamics in fluids of platelike colloidal particles},
  author={Bier, Markus and van Roij, Ren{\'e}},
  journal=PRE,
  volume={76},
  number={2},
  pages={021405},
  year={2007},
  publisher={APS}
}

@article{DasY2013,
  title={Coarse-grained forms for equations describing the microscopic motion of particles in a fluid},
  author={Das, Shankar P and Yoshimori, Akira},
  journal=PRE,
  volume={88},
  number={4},
  pages={043008},
  year={2013},
  publisher={APS}
}

@article{KelkarFC2014,
  title={Nonideal diffusion effects and short-range ordering lead to higher aggregation rates in concentrated hard-sphere dispersions},
  author={Kelkar, Aniruddha~V and Franses, Elias~I and Corti, David~S},
  journal={Langmuir},
  volume={30},
  number={13},
  pages={3647-3657},
  year={2014},
  publisher={ACS Publications}
}

@article{LichtnerK2010,
  title={Feedback-controlled transport in an interacting colloidal system},
  author={Lichtner, Ken and Klapp, Sabine HL},
  journal={Europhysics Letters},
  volume={92},
  number={4},
  pages={40007},
  year={2010},
  publisher={IOP Publishing}
}

@article{MendozaLSRPM2017,
  title={Crossover from equilibration to aging: Nonequilibrium theory versus simulations},
  author={Mendoza-M{\'e}ndez, P and L{\'a}zaro-L{\'a}zaro, E and S{\'a}nchez-D{\'\i}az, LE and Ram{\'\i}rez-Gonz{\'a}lez, PE and P{\'e}rez-{\'A}ngel, G and Medina-Noyola, M},
  journal=PRE,
  volume={96},
  number={2},
  pages={022608},
  year={2017},
  publisher={APS}
}


@article{MarconiM2011b,
  title={Multicomponent diffusion in nanosystems},
  author={{Marini Bettolo Marconi}, Umberto and Melchionna, Simone},
  journal=JoChemP,
  volume={135},
  number={4},
  pages={044104},
  year={2011},
  publisher={AIP}
}

@article{ZhaoWZDTM2014,
  title={Role of polymer conformation and hydrodynamics on nanoparticle deposits on a substrate},
  author={Zhao, Qiangli and Wang, Xiangdong and Zhang, Chen and Du, Zhongjie and Tian, Ming and Mi, Jianguo},
  journal={Journal of Physical Chemistry C},
  volume={118},
  number={46},
  pages={26808--26815},
  year={2014},
  publisher={ACS Publications}
}


@article{WangS2016,
  title={Large amplitude oscillatory shear of hard-sphere colloidal dispersions: {B}rownian dynamics simulation and {F}ourier-transform rheology},
  author={Wang, Gang and Swan, James W},
  journal={Journal of Rheology},
  volume={60},
  number={6},
  pages={1041--1053},
  year={2016},
  publisher={SOR}
}


@article{PiazzaDDDF2013,
  title={Diffusion-limited reactions in crowded environments: a local density approximation},
  author={Piazza, F and Dorsaz, N and De Michele, C and De Los Rios, P and Foffi, G},
  journal=JoP:CM,
  volume={25},
  number={37},
  pages={375104},
  year={2013},
  publisher={IOP Publishing}
}


@article{StopperR2017,
  title={Massively parallel {GPU}-accelerated minimization of classical density functional theory},
  author={Stopper, Daniel and Roth, Roland},
  journal=JoChemP,
  volume={147},
  number={6},
  pages={064508},
  year={2017},
  publisher={AIP Publishing}
}

@article{HuetterB2009,
  title={Nonlocal effects in nonisothermal hydrodynamics from the perspective of beyond-equilibrium thermodynamics},
  author={H{\"u}tter, Markus and Brader, Joseph M},
  journal=JoChemP,
  volume={130},
  number={21},
  pages={214908},
  year={2009},
  publisher={AIP}
}

@article{Cahn1965,
  title={Phase separation by spinodal decomposition in isotropic systems},
  author={Cahn, John W},
  journal=JoChemP,
  volume={42},
  number={1},
  pages={93--99},
  year={1965},
  publisher={AIP}
}


@article{GuanLW2017,
  title={{Convergence analysis for second-order accurate schemes for the periodic nonlocal {A}llen-{C}ahn and {C}ahn-{H}illiard equations}},
  author={Guan, Zhen and Lowengrub, John and Wang, Cheng},
  journal={Mathematical Methods in the Applied Sciences},
  volume={40},
  number={18},
  pages={6836--6863},
  year={2017},
  publisher={Wiley Online Library}
}

@article{Wilemski1976,
  title={On the derivation of {S}moluchowski equations with corrections in the classical theory of {B}rownian motion},
  author={Wilemski, Gerald},
  journal={Journal of Statistical Physics},
  volume={14},
  number={2},
  pages={153--169},
  year={1976},
  publisher={Springer}
}

@article{Lutsko2016,
  title={Mechanism for the stabilization of protein clusters above the solubility curve: the role of non-ideal chemical reactions},
  author={Lutsko, James F},
  journal=JoP:CM,
  volume={28},
  number={24},
  pages={244020},
  year={2016},
  publisher={IOP Publishing}
}


@article{Bier2015,
  title={Nonequilibrium interfacial tension during relaxation},
  author={Bier, Markus},
  journal=PRE,
  volume={92},
  number={4},
  pages={042128},
  year={2015},
  publisher={APS}
}


@article{AngiolettiBD2018,
  title={Competitive adsorption of multiple proteins to nanoparticles: the {V}roman effect revisited},
  author={Angioletti-Uberti, Stefano and Ballauff, Matthias and Dzubiella, Joachim},
  journal=MP,
  volume={116},
  number={21-22},
  pages={3154--3163},
  year={2018},
  publisher={Taylor \& Francis}
}


@article{SanchezLOGMN2014,
  title={Non-equilibrium dynamics of glass-forming liquid mixtures},
  author={S{\'a}nchez-D{\'\i}az, Luis Enrique and L{\'a}zaro-L{\'a}zaro, Edilio and Olais-Govea, Jos{\'e} Manuel and Medina-Noyola, Magdaleno},
  journal=JoChemP,
  volume={140},
  number={23},
  pages={234501},
  year={2014},
  publisher={AIP}
}


@article{BurghardtB2006,
  title={On the non-adiabatic dynamics of solvation: A molecular hydrodynamic formulation},
  author={Burghardt, Irene and Bagchi, Biman},
  journal={Chemical Physics},
  volume={329},
  number={1-3},
  pages={343--356},
  year={2006},
  publisher={Elsevier}
}


@article{KlopotekHGDSSO2017,
  title={Monolayers of hard rods on planar substrates. II. Growth},
  author={Klopotek, Miriam and Hansen-Goos, Hendrik and Dixit, Mohit and Schilling, Tanja and Schreiber, Frank and Oettel, Martin},
  journal=JoChemP,
  volume={146},
  number={8},
  pages={084903},
  year={2017},
  publisher={AIP Publishing}
}


@article{Haertel2017,
  title={Structure of electric double layers in capacitive systems and to what extent (classical) density functional theory describes it},
  author={Haertel, Andreas},
  journal={JoP:CM},
  volume={29},
  number={42},
  pages={423002},
  year={2017},
  publisher={IOP Publishing}
}


@article{TothGT2013,
  title={Nonlinear hydrodynamic theory of crystallization},
  author={T{\'o}th, Gyula I and Gr{\'a}n{\'a}sy, L{\'a}szl{\'o} and Tegze, Gy{\"o}rgy},
  journal=JoP:CM,
  volume={26},
  number={5},
  pages={055001},
  year={2014},
  publisher={IOP Publishing}
}

@article{NeuhausHMSL2014,
  title={Density functional theory of heterogeneous crystallization},
  author={Neuhaus, T and H{\"a}rtel, Andreas and Marechal, M and Schmiedeberg, M and L{\"o}wen, Hartmut},
  journal={European Physical Journal Special Topics},
  volume={223},
  number={3},
  pages={373--387},
  year={2014},
  publisher={Springer}
}

@article{NeuhausSL2013,
  title={Compatibility waves drive crystal growth on patterned substrates},
  author={Neuhaus, Tim and Schmiedeberg, Michael and L{\"o}wen, Hartmut},
  journal=NJoP,
  volume={15},
  number={7},
  pages={073013},
  year={2013},
  publisher={IOP Publishing}
}

@article{MendezFKYSCA2018,
  title={Density-functional fluctuation theory of crowds},
  author={M{\'e}ndez-Valderrama, J Felipe and Kinkhabwala, Yunus A and Silver, Jeffrey and Cohen, Itai and Arias, TA},
  journal=NC,
  volume={9},
  number={1},
  pages={3538},
  year={2018},
  publisher={Nature Publishing Group}
}

@article{MartinezCMOL2013,
  title={Optimizing the search for resources by sharing information: {M}ongolian gazelles as a case study},
  author={Mart{\'\i}nez-Garc{\'\i}a, Ricardo and Calabrese, Justin M and Mueller, Thomas and Olson, Kirk A and L{\'o}pez, Crist{\'o}bal},
  journal=PRL,
  volume={110},
  number={24},
  pages={248106},
  year={2013},
  publisher={APS}
}

@article{FuchizakiK2002,
  title={Dynamical density functional theory for glassy behaviour},
  author={Fuchizaki, Kazuhiro and Kawasaki, Kyozi},
  journal=JoP:CM,
  volume={14},
  number={46},
  pages={12203},
  year={2002},
  publisher={IOP Publishing}
}

@article{Angioletti2017,
  title={Theory, simulations and the design of functionalized nanoparticles for biomedical applications: A Soft Matter Perspective},
  author={Angioletti-Uberti, Stefano},
  journal={npj Computational Materials},
  volume={3},
  number={1},
  pages={1--15},
  year={2017},
  publisher={Nature Publishing Group}
}

@article{SaltzmanYS2008,
  title={Anomalous diffusion, structural relaxation and shear thinning in glassy hard sphere fluids},
  author={Saltzman, Erica J and Yatsenko, Galina and Schweizer, Kenneth S},
  journal=JoP:CM,
  volume={20},
  number={24},
  pages={244129},
  year={2008},
  publisher={IOP Publishing}
}

@article{BousquetHMB2011,
  title={Extended hydrodynamic approach to quantum-classical nonequilibrium evolution. I. Theory},
  author={Bousquet, David and Hughes, Keith H and Micha, David A and Burghardt, Irene},
  journal=JoChemP,
  volume={134},
  number={6},
  pages={064116},
  year={2011},
  publisher={AIP}
}


@article{BackofenGPPVW2011,
  title={A continuous approach to discrete ordering on $S^{2}$},
  author={Backofen, Rainer and Gr{\"a}f, Manuel and Potts, Daniel and Praetorius, Simon and Voigt, Axel and Witkowski, Thomas},
  journal={Multiscale Modeling \& Simulation},
  volume={9},
  number={1},
  pages={314--334},
  year={2011},
  publisher={SIAM}
}

@article{LopezM2007,
  title={Multiple time-scale approach for a system of {B}rownian particles in a nonuniform temperature field},
  author={L{\'o}pez, Crist{\'o}bal and {Marini Bettolo Marconi}, Umberto},
  journal={Physical Review E},
  volume={75},
  number={2},
  pages={021101},
  year={2007},
  publisher={APS}
}

@article{MarconiM2007,
  title={Phase-space approach to dynamical density functional theory},
  author={{Marini Bettolo Marconi}, Umberto and Melchionna, Simone},
  journal=JoChemP,
  volume={126},
  number={18},
  pages={184109},
  year={2007},
  publisher={AIP}
}

@INCOLLECTION{Graham1978,
  author       = {{Graham}, R.},
  title        = {Path-integral methods in Nonequilibrium Thermodynamics and Statistics},
  booktitle    = {Stochastic Processes in Nonequilibrium Systems},
  publisher    = SPR,
  year         = {1978},
  editor       = {L. Garrido and P. Seglar and P.~J. Shepherd},
  volume       = {84},
  series       = {Lecture Notes in Physics},
  type         = {chapter},
  chapter      = {4},
  pages        = {82-138},
  address      = BER,
  edition      = {1},
  month        = JUN,
  isbn         = {3-540-08942-X}
}
% doi = {10.1007/BFb0016715}
% url = {http://adsabs.harvard.edu/abs/1978LNP....84...82G}

@ARTICLE{Graham1977a,
  author       = {{Graham}, R.},
  title        = {Path integral formulation of general diffusion processes},
  journal      = ZfPB:CM,
  publisher    = SPR,
  year         = {1977},
  volume       = {26},
  number       = {3},
  pages        = {281-290},
  month        = SEP
}
% doi = {10.1007/BF01312935}
% url = {http://adsabs.harvard.edu/abs/1977ZPhyB..26..281G}

@ARTICLE{Graham1977b,
  author       = {{Graham}, R.},
  title        = {Covariant formulation of non-equilibrium statistical thermodynamics},
  journal      = ZfPB:CM,
  publisher    = SPR,
  year         = {1977},
  volume       = {26},
  number       = {4},
  pages        = {397-405},
  month        = DEC
}
% doi = {10.1007/BF01570750}
% url = {http://adsabs.harvard.edu/abs/1977ZPhyB..26..397G}

@ARTICLE{deGennes1969,
  author       = {de~Gennes, P.-G.},
  title        = {Conjectures sur l'{\'e}tat smectique},
  journal      = JdPC,
  publisher    = {Soci\'{e}t\'{e} fran\c{c}aise de physique},
  year         = {1969},
  volume       = {30},
  number       = {C4},
  pages        = {65-71},
  month        = NOV
}
% doi = {10.1051/jphyscol:1969415}
% url = {http://dx.doi.org/10.1051/jphyscol:1969415}

@ARTICLE{HohenbergK1964,
  author       = {{Hohenberg}, P. and {Kohn}, W.},
  title        = {Inhomogeneous electron gas},
  journal      = PR,
  publisher    = APS,
  year         = {1964},
  volume       = {136},
  number       = {3B},
  pages        = {864-871},
  month        = NOV
}
% doi = {10.1103/PhysRev.136.B864}
% url = {http://adsabs.harvard.edu/abs/1964PhRv..136..864H}

@article{LoewenLABvT2007,
  title={Critical nuclei and crystallization in colloidal suspensions},
  author={L{\"o}wen, H and Likos, CN and Assoud, L and Blaak, R and van Teefelen, S},
  journal={Philosophical Magazine},
  volume={87},
  number={11},
  pages={847--854},
  year={2007},
  publisher={Taylor \& Francis}
}

@article{LichtnerK2014,
  title={Novel structure formation of a phase separating colloidal fluid in a ratchet potential},
  author={Lichtner, Ken and Klapp, Sabine HL},
  journal={Europhysics Letters},
  volume={106},
  number={5},
  pages={56004},
  year={2014},
  publisher={IOP Publishing}
}

@article{CapriniM2021b,
  title={Spatial velocity correlations in inertial systems of Active {B}rownian Particles},
  author={Caprini, Lorenzo and {Marini Bettolo Marconi}, Umberto},
  journal=SM,
  note={{DOI}: 10.1039/D0SM02273J},
  year={2021},
  publisher={Royal Society of Chemistry}
}

@article{ChervanyovGT2016,
  title={Effect of the orientational relaxation on the collective motion of patterns formed by self-propelled particles},
  author={Chervanyov, A I and Gomez, H and Thiele, U},
  journal=EPL,
  volume={115},
  number={6},
  pages={68001},
  year={2016},
  publisher={IOP Publishing}
}

@article{SprengerJIL2021,
  title={Time-dependent inertia of self-propelled particles: the {L}angevin rocket},
  author={Sprenger, Alexander R and Jahanshahi, Soudeh and Ivlev, Alexei V and L{\"o}wen, Hartmut},
  journal={arXiv:2101.01608},
  year={2021}
}

@ARTICLE{HohenbergH1977,
  author       = {{Hohenberg}, P.~C. and {Halperin}, B.~I.},
  title        = {Theory of dynamic critical phenomena},
  journal      = RoMP,
  publisher    = APS,
  year         = {1977},
  volume       = {49},
  number       = {3},
  pages        = {435-479},
  month        = JUL
}
% doi = {10.1103/RevModPhys.49.435}
% url = {http://adsabs.harvard.edu/abs/1977RvMP...49..435H}

@article{Jones2015,
  title={Density functional theory: Its origins, rise to prominence, and future},
  author={Jones, Robert O},
  journal=RoMP,
  volume={87},
  number={3},
  pages={897},
  year={2015},
  publisher={APS}
}

@article{JohnsonN1981,
  title={Generalized van der {W}aals theory. VI. Application to adsorption},
  author={Johnson, Mark and Nordholm, Sture},
  journal=JoChemP,
  volume={75},
  number={4},
  pages={1953--1957},
  year={1981},
  publisher={AIP}
}

@article{NordholmH1980,
  title={Generalized van der {W}aals theory. I. Basic formulation and application to uniform fluids},
  author={Nordholm, S and Haymet, ADJ},
  journal={Australian Journal of Chemistry},
  volume={33},
  number={9},
  pages={2013--2027},
  year={1980},
  publisher={CSIRO}
}

@ARTICLE{KohnS1965,
  author       = {{Kohn}, W. and {Sham}, L.~J.},
  title        = {Self-consistent equations including exchange and correlation effects},
  journal      = PR,
  publisher    = APS,
  year         = {1965},
  volume       = {140},
  number       = {4A},
  pages        = {1133-1138},
  month        = NOV
}
% doi = {10.1103/PhysRev.140.A1133}
% url = {http://adsabs.harvard.edu/abs/1965PhRv..140.1133K}

@ARTICLE{EbnerSS1976,
  author       = {{Ebner}, C. and {Saam}, W.~F. and {Stroud}, D},
  title        = {Density-functional theory of simple classical fluids: {I.} {S}urfaces},
  journal      = PRA,
  publisher    = APS,
  year         = {1976},
  volume       = {14},
  number       = {6},
  pages        = {2264-2273},
  month        = DEC
}
% doi = {10.1103/PhysRevA.14.2264}
% url = {http://adsabs.harvard.edu/abs/1976PhRvA..14.2264E}

@ARTICLE{SaamE1977,
  author       = {Saam, W.~F. and Ebner, C.},
  title        = {Density-functional theory of classical systems},
  journal      = PRA,
  publisher    = APS,
  year         = {1977},
  volume       = {15},
  number       = {6},
  pages        = {2566-2568},
  month        = JUN
}
% doi = {10.1103/PhysRevA.15.2566}
% url = {http://link.aps.org/doi/10.1103/PhysRevA.15.2566}

@ARTICLE{YangFG1976,
  author       = {{Yang}, A.~J.~M. and {Fleming}, P.~D. and {Gibbs}, J.~H.},
  title        = {Molecular theory of surface tension},
  journal      = JoChemP,
  publisher    = AIoP,
  year         = {1976},
  volume       = {64},
  number       = {9},
  pages        = {3732-3747},
  month        = MAY
}
% doi = {10.1063/1.432687}
% url = {http://adsabs.harvard.edu/abs/1976JChPh..64.3732Y}

@article{MatuszakAD2006,
  title={Single-component permeation maximum with respect to temperature: a lattice density functional theory study},
  author={Matuszak, Daniel and Aranovich, Gregory L and Donohue, Marc D},
  journal={Industrial \& Engineering Chemistry Research},
  volume={45},
  number={16},
  pages={5501--5511},
  year={2006},
  publisher={ACS Publications}
}


@article{PierrezMadridRR2002,
	year = 2002,
	month = {feb},
	publisher = {{IOP} Publishing},
	volume = {14},
	number = {7},
	pages = {1651--1657},
	author = {A P{\'{e}}rez-Madrid and D Reguera and J M Rub{\'{\i}}},
	title = {A mesoscopic approach to the slow dynamics of supercooled liquids and colloidal systems},
	journal = JoP:CM
}

%	doi = {10.1088/0953-8984/14/7/321},
%	url = {https://doi.org/10.1088%2F0953-8984%2F14%2F7%2F321},

@article{LeoniPEENASA2020,
  title={Surfing and crawling macroscopic active particles under strong confinement: inertial dynamics},
  author={Leoni, Marco and Paoluzzi, Matteo and Eldeen, Sarah and Estrada, Anthony and Nguyen, Lauren and Alexandrescu, Maria and Sherb, Karin and Ahmed, Wylie W},
  journal={Physical Review Research},
  volume={2},
  number={4},
  pages={043299},
  year={2020},
  publisher={APS}
}

@article{CapriniMM2021,
  title={Collective effects in confined Active {B}rownian Particles},
  author={Caprini, Lorenzo and Maggi, Claudio and {Marini Bettolo Marconi}, Umberto},
  journal={arXiv:2103.13220},
  year={2021}
}

@INBOOK{Pleiner1994,
  author       = {Pleiner, Harald},
  title        = {Komplexe Systeme zwischen Atom und Festk{\"o}rper: Vorlesungsmanuskripte des 25.\ IFF-Ferienkurses vom 28. Februar 1994 bis 
11. M{\"a}rz 1994 im Forschungszentrum J{\"u}lich},
  chapter      = {20: Hydrodynamik komplexer Fluide},
  publisher    = {Forschungszentrum J{\"u}lich},
  year         = {1994},
  volume       = {25},
  series       = {IFF Spring School Lecture Notes},
  address      = {J{\"u}lich},
  editor       = {R. H{\"o}lzle},
  edition      = {1},
  month        = MAR,
  pages        = {20.1-20.32},
  isbn         = {3-893-36128-6}
}
% ISBN-13: 978-3893361281

@INBOOK{Pleiner1997,
  author       = {Pleiner, Harald},
  title        = {Dynamik und Strukturbildung in kondensierter Materie: Vorlesungsmanuskripte des 28.\ IFF-Ferienkurses vom 3. M{\"a}rz 1997 bis 
14. M{\"a}rz 1997 im Forschungszentrum J{\"u}lich},
  chapter      = {E1: Komplexe Fluide -- ein {\"U}berblick},
  publisher    = {Forschungszentrum J{\"u}lich},
  year         = {1997},
  volume       = {28},
  series       = {IFF Spring School Lecture Notes},
  address      = {J{\"u}lich},
  editor       = {R. H{\"o}lzle},
  edition      = {1},
  month        = MAR,
  pages        = {E1.1-E1.22},
  isbn         = {3-893-36204-5}
}
% ISBN-13: 978-3893362042

@BOOK{Onuki2002,
  author       = {Onuki, Akira},
  title        = {Phase Transition Dynamics},
  publisher    = CUP,
  year         = {2002},
  address      = CAM,
  edition      = {1st},
  month        = JUN,
  pages        = {726},
  isbn         = {0-521-57293-2}
}
% ISBN-13: 978-0521572934

@BOOK{MaW2013,
  author       = {Ma, Tian and Wang, Shouhong},
  title        = {Phase Transition Dynamics},
  publisher    = SPR,
  year         = {2013},
  address      = NY,
  edition      = {1st},
  month        = NOV,
  pages        = {555},
  isbn         = {978-1-461-48962-7}
}
% ISBN-10: 
% DOI:10.1007/978-1-4614-8963-4

@BOOK{ToledanoT1987,
  author       = {Toledano, Jean-Claude and Toledano, Pierre},
  title        = {The Landau Theory of Phase Transitions: Application to Structural, Incommensurate, Magnetic, and Liquid Crystal Systems},
  publisher    = WSP,
  year         = {1987},
  volume       = {3},
  series       = {World Scientific Lecture Notes in Physics},
  address      = SIN,
  edition      = {1},
  month        = AUG,
  pages        = {451},
  isbn         = {9-971-50025-6}
}
% EAN: 9789971500252

@ARTICLE{ChuangYDT1991,
  author       = {{Chuang}, I. and {Yurke}, B. and {Durrer}, R. and {Turok}, N.},
  title        = {Cosmology in the laboratory -- defect dynamics in liquid crystals},
  journal      = SCI,
  publisher    = AAftAoS,
  year         = {1991},
  volume       = {251},
  number       = {4999},
  pages        = {1336-1342},
  month        = MAR
}
% doi = {10.1126/science.251.4999.1336}
% url = {http://adsabs.harvard.edu/abs/1991Sci...251.1336C}

@BOOK{ProvatasE2010,
  author       = {Provatas, Nikolas and Elder, Ken},
  title        = {Phase-Field Methods in Materials Science and Engineering},
  publisher    = WILEYVCH,
  year         = {2010},
  address      = WEI,
  edition      = {1},
  month        = OCT,
  pages        = {312},
  isbn         = {3-527-40747-2}
}
% ISBN13: 978-3527407477

@ARTICLE{SwiftH1977,
  author       = {{Swift}, J. and {Hohenberg}, P.~C.},
  title        = {Hydrodynamic fluctuations at the convective instability},
  journal      = PRA,
  publisher    = APS,
  year         = {1977},
  volume       = {15},
  number       = {1},
  pages        = {319-328},
  month        = JAN
}
% doi = {10.1103/PhysRevA.15.319}
% url = {http://adsabs.harvard.edu/abs/1977PhRvA..15..319S}


@article{Lutsko2008,
  title={Density functional theory of inhomogeneous liquids. III. Liquid-vapor nucleation},
  author={Lutsko, James F},
  journal=JoChemP,
  volume={129},
  number={24},
  pages={244501},
  year={2008},
  publisher={AIP}
}

@article{CecconiDMP2004,
  title={Fluid-like behavior of a one-dimensional granular gas},
  author={Cecconi, Fabio and Diotallevi, Fabiana and {Marini Bettolo Marconi}, Umberto and Puglisi, Andrea},
  journal=JoChemP,
  volume={120},
  number={1},
  pages={35--42},
  year={2004},
  publisher={AIP}
}

@article{MarconiT2006,
  title={Nonequilibrium inertial dynamics of colloidal systems},
  author={{Marini Bettolo Marconi}, Umberto and Tarazona, Pedro},
  journal=JoChemP,
  volume={124},
  number={16},
  pages={164901},
  year={2006},
  publisher={AIP}
}

@article{BerryGE2006,
  title={Diffusive atomistic dynamics of edge dislocations in two dimensions},
  author={Berry, J and Grant, M and Elder, KR},
  journal=PRE,
  volume={73},
  number={3},
  pages={031609},
  year={2006},
  publisher={APS}
}

@ARTICLE{BerryG2011,
  author       = {{Berry}, J. and {Grant}, M.},
  title        = {Modeling multiple time scales during glass formation with phase-field crystals},
  journal      = PRL,
  publisher    = APS,
  year         = {2011},
  volume       = {106},
  number       = {17},
  pages        = {175702},
  month        = APR
}
% doi = {10.1103/PhysRevLett.106.175702}
% url = {http://adsabs.harvard.edu/abs/2011PhRvL.106q5702B}

@INBOOK{Evans1992,
  author       = {Evans, R.},
  title        = {Fundamentals of Inhomogeneous Fluids},
  chapter      = {3: Density Functionals in the Theory of Nonuniform Fluids},
  publisher    = MD,
  year         = {1992},
  address      = NY,
  editor       = {Douglas Henderson},
  edition      = {1},
  month        = AUG,
  pages        = {85-176},
  isbn         = {0-824-78711-0}
}
% ISBN-13: 978-0824787110

@article{Loewen2003,
  title={Density functional theory: from statics to dynamics},
  author={L{\"o}wen, H},
  journal=JoP:CM,
  volume={15},
  number={6},
  pages={V1},
  year={2003},
  publisher={Bristol, UK: IOP Pub., c1989-}
}

@article{YeNTZM2019,
  title={Shear-Induced Microscopic Structure Damage in Polymer Nanocomposites: A Dynamic Density Functional Theoretical Study},
  author={Ye, Yi and Ning, Nanying and Tian, Ming and Zhang, Liqun and Mi, Jianguo},
  journal={Journal of Physical Chemistry C},
  volume={123},
  number={36},
  pages={22529--22538},
  year={2019},
  publisher={ACS Publications}
}


@inproceedings{YatsyshinK2016,
  title={Classical Density-Functional Theory Studies of Fluid Adsorption on Nanopatterned Planar Surfaces},
  author={Yatsyshin, Peter and Kalliadasis, Serafim},
  booktitle={Workshop on Coupled Mathematical Models for Physical and Nanoscale Systems and their Applications},
  pages={171--185},
  year={2016},
  organization={Springer}
}


@ARTICLE{Loewen2002,
  author       = {L{\"o}wen, H.},
  title        = {Density functional theory of inhomogeneous classical fluids: recent developments and new perspectives},
  journal      = JoP:CM,
  publisher    = IoPP,
  year         = {2002},
  volume       = {14},
  number       = {46},
  pages        = {11897-11905},
  month        = NOV
}
% doi = {10.1088/0953-8984/14/46/301}
% url = {http://adsabs.harvard.edu/abs/2002JPCM...1411897L}

@BOOK{Kalikmanov2010,
  author       = {Kalikmanov, V.~I.},
  title        = {Statistical Physics of Fluids: Basic Concepts and Applications},
  publisher    = SPR,
  year         = {2010},
  volume       = {46},
  series       = {Theoretical and Mathematical Physics},
  address      = BER,
  edition      = {2},
  month        = DEC,
  pages        = {260},
  isbn         = {3-642-07511-8}
}
% EAN: 9783642075117

@ARTICLE{GinzburgL1950,
  author       = {Ginzburg, V.~L. and Landau, L.~D.},
  title        = {On the theory of superconductivity},
  journal      = ZEiTF,
  publisher    = {Maik Nauka/Interperiodica},
  year         = {1950},
  volume       = {20},
  number       = {},
  pages        = {1064-1082},
  month        = {}
}
% doi = {}
% url = {}

@article{DeeL1983,
  title={Propagating pattern selection},
  author={Dee, G and Langer, JS},
  journal=PRL,
  volume={50},
  number={6},
  pages={383},
  year={1983},
  publisher={APS}
}

@article{BenJacobDK1985,
  title={Pattern propagation in nonlinear dissipative systems},
  author={Ben-Jacob, E and Brand, H and Dee, G and Kramer, L and Langer, JS},
  journal={Physica D: Nonlinear Phenomena},
  volume={14},
  number={3},
  pages={348--364},
  year={1985},
  publisher={Elsevier}
}

@ARTICLE{BardeenCS1957b,
  author       = {{Bardeen}, J. and {Cooper}, L.~N. and {Schrieffer}, J.~R.},
  title        = {Theory of superconductivity},
  journal      = PR,
  publisher    = APS,
  year         = {1957},
  volume       = {108},
  number       = {5},
  pages        = {1175-1204},
  month        = DEC
}
% doi = {10.1103/PhysRev.108.1175}
% url = {http://adsabs.harvard.edu/abs/1957PhRv..108.1175B}

@ARTICLE{BardeenCS1957a,
  author       = {{Bardeen}, J. and {Cooper}, L.~N. and {Schrieffer}, J.~R.},
  title        = {Microscopic theory of superconductivity},
  journal      = PR,
  publisher    = APS,
  year         = {1957},
  volume       = {106},
  number       = {1},
  pages        = {162-164},
  month        = APR
}
% doi = {10.1103/PhysRev.106.162}
% url = {http://link.aps.org/doi/10.1103/PhysRev.106.162}

@ARTICLE{Rosenfeld1989,
  author       = {{Rosenfeld}, Y.},
  title        = {Free-energy model for the inhomogeneous hard-sphere fluid mixture and density-functional theory of freezing},
  journal      = PRL,
  publisher    = APS,
  year         = {1989},
  volume       = {63},
  number       = {9},
  pages        = {980-983},
  month        = AUG
}
% doi = {10.1103/PhysRevLett.63.980}
% url = {http://adsabs.harvard.edu/abs/1989PhRvL..63..980R}

@ARTICLE{RosenfeldSLT1996,
  author       = {{Rosenfeld}, Y. and {Schmidt}, M. and {L{\"o}wen}, H. and {Tarazona}, P.},
  title        = {Dimensional crossover and the freezing transition in density functional theory},
  journal      = JoP:CM,
  publisher    = IoPP,
  year         = {1996},
  volume       = {8},
  number       = {40},
  pages        = {L577-L581},
  month        = SEP
}
% doi = {10.1088/0953-8984/8/40/002}
% url = {http://adsabs.harvard.edu/abs/1996JPCM....8L.577R}

@ARTICLE{Rosenfeld1994,
  author       = {{Rosenfeld}, Y.},
  title        = {Density functional theory of molecular fluids: free-energy model for the inhomogeneous hard-body fluid},
  journal      = PRE,
  publisher    = APS,
  year         = {1994},
  volume       = {50},
  number       = {5},
  pages        = {R3318-R3321},
  month        = NOV
}
% doi = {10.1103/PhysRevE.50.R3318}
% url = {http://adsabs.harvard.edu/abs/1994PhRvE..50.3318R}

@ARTICLE{DieterichFM1990,
  author       = {{Dieterich}, W. and {Frisch}, H.~L. and {Majhofer}, A.},
  title        = {Nonlinear diffusion and density functional theory},
  journal      = ZfPB:CM,
  publisher    = SPR,
  year         = {1990},
  volume       = {78},
  number       = {2},
  pages        = {317-323},
  month        = JUN
}
% doi = {10.1007/BF01307852}
% url = {http://adsabs.harvard.edu/abs/1990ZPhyB..78..317D}

@ARTICLE{LovettMB1976,
  author       = {{Lovett}, R. and {Mou}, C.~Y. and {Buff}, F.~P.},
  title        = {The structure of the liquid-vapor interface},
  journal      = JoChemP,
  publisher    = AIoP,
  year         = {1976},
  volume       = {65},
  number       = {2},
  pages        = {570-572},
  month        = JUL
}
% doi = {10.1063/1.433110}
% url = {http://adsabs.harvard.edu/abs/1976JChPh..65..570L}

@ARTICLE{YabunakaA2011,
  author       = {{Yabunaka}, S. and {Araki}, T.},
  title        = {Polydomain growth at isotropic-nematic transitions in liquid crystalline polymers},
  journal      = PRE,
  publisher    = APS,
  year         = {2011},
  volume       = {83},
  number       = {6},
  pages        = {061711},
  month        = JUN
}
% doi = {10.1103/PhysRevE.83.061711}
% url = {http://adsabs.harvard.edu/abs/2011PhRvE..83f1711Y}

@ARTICLE{ZaburdaevUPEFS2011,
  author       = {Zaburdaev, Vasily and Uppaluri, Sravanti and Pfohl, Thomas and Engstler, Markus and Friedrich, Rudolf and Stark, Holger},
  title        = {Langevin dynamics deciphers the motility pattern of swimming parasites},
  journal      = PRL,
  publisher    = APS,
  year         = {2011},
  volume       = {106},
  number       = {20},
  pages        = {208103},
  month        = MAY
}
% doi = {10.1103/PhysRevLett.106.208103}
% url = {http://link.aps.org/doi/10.1103/PhysRevLett.106.208103}

@ARTICLE{KioyJM2004,
  author       = {Kioy, D. and Jannin, J. and Mattock, N.},
  title        = {Focus: Human African trypanosomiasis},
  journal      = {Nature Reviews Microbiology},
  publisher    = NPG,
  year         = {2004},
  volume       = {2},
  number       = {3},
  pages        = {186-187},
  month        = MAR
}
% doi = {doi:10.1038/nrmicro848}
% url = {http://www.nature.com/nrmicro/journal/v2/n3/full/nrmicro848.html}

@ARTICLE{DowntonS2009,
  author       = {{Downton}, M.~T. and {Stark}, H.},
  title        = {Beating kinematics of magnetically actuated cilia},
  journal      = EPL,
  publisher    = EPS,
  year         = {2009},
  volume       = {85},
  number       = {4},
  pages        = {44002},
  month        = FEB
}
% doi = {10.1209/0295-5075/85/44002}
% url = {http://adsabs.harvard.edu/abs/2009EL.....8544002D}

@ARTICLE{GaugerDS2009,
  author       = {{Gauger}, E.~M. and {Downton}, M.~T. and {Stark}, H.},
  title        = {Fluid transport at low {R}eynolds number with magnetically actuated artificial cilia},
  journal      = EPJE,
  publisher    = EDPS,
  year         = {2009},
  volume       = {28},
  number       = {2},
  pages        = {231-242},
  month        = FEB
}
% doi = {10.1140/epje/i2008-10388-1}
% url = {http://adsabs.harvard.edu/abs/2009EPJE...28..231G}

@MISC{StarkPhysikJournal2007bis2009,
  author       = {Stark, Holger},
  note = {Phys. J. \textbf{6}, 31 (2007); \textbf{8}, 16 (2009); \textbf{9}, 16 (2010)}
}

@ARTICLE{Stark2007,
  author       = {Stark, Holger},
  title        = {{I}mmer in {B}ewegung bleiben: {D}ie sonderbare {W}elt der kleinen {R}eynolds-{Z}ahlen},
  journal      = PJ,
  publisher    = WILEYVCH,
  year         = {2007},
  volume       = {6},
  number       = {11},
  pages        = {31-37},
  month        = NOV
}

@ARTICLE{Stark2009,
  author       = {Stark, Holger},
  title        = {{A}lgentanz im {W}alzertakt: {Z}ilien von {M}ikroorganismen erzeugen eine nichtlineare {D}ynamik in z{\"a}her {U}mgebung},
  journal      = PJ,
  publisher    = WILEYVCH,
  year         = {2009},
  volume       = {8},
  number       = {7},
  pages        = {16-19},
  month        = JUL
}

@ARTICLE{Stark2010,
  author       = {Stark, Holger},
  title        = {{Z}{\"a}he {S}chwimmer},
  journal      = PJ,
  publisher    = WILEYVCH,
  year         = {2010},
  volume       = {9},
  number       = {5},
  pages        = {16-17},
  month        = MAY
}

@ARTICLE{LiangSRL2011,
  author       = {{Liang}, H.-L. and {Schymura}, S. and {Rudquist}, P. and {Lagerwall}, J.},
  title        = {Nematic-Smectic Transition under Confinement in Liquid Crystalline Colloidal Shells},
  journal      = PRL,
  publisher    = APS,
  year         = {2011},
  volume       = {106},
  number       = {24},
  pages        = {247801},
  month        = JUN
}
% doi = {10.1103/PhysRevLett.106.247801}
% url = {http://adsabs.harvard.edu/abs/2011PhRvL.106x7801L}

@ARTICLE{Louis2001,
  author       = {{Louis}, A.~A.},
  title        = {Effective potentials for polymers and colloids: beyond the van der {W}aals picture of fluids?},
  journal      = RSoLPTSA,
  publisher    = RS,
  year         = {2001},
  volume       = {359},
  number       = {1782},
  pages        = {939-960},
  month        = MAY
}
% doi = {10.1098/rsta.2000.0804}
% url = {http://adsabs.harvard.edu/abs/2001RSPTA.359..939L}

@ARTICLE{LikosLWL2001,
  author       = {{Likos}, C.~N. and {Lang}, A. and {Watzlawek}, M. and {L{\"o}wen}, H.},
  title        = {Criterion for determining clustering versus reentrant melting behavior for bounded interaction potentials},
  journal      = PRE,
  publisher    = APS,
  year         = {2001},
  volume       = {63},
  number       = {3},
  pages        = {031206},
  month        = MAR
}
% doi = {10.1103/PhysRevE.63.031206}
% url = {http://adsabs.harvard.edu/abs/2001PhRvE..63c1206L}

@ARTICLE{Tarazona2000,
  author       = {{Tarazona}, P.},
  title        = {Density functional for hard sphere crystals: a fundamental measure approach},
  journal      = PRL,
  publisher    = APS,
  year         = {2000},
  volume       = {84},
  number       = {4},
  pages        = {694-697},
  month        = JAN
}
% doi = {10.1103/PhysRevLett.84.694}
% url = {http://adsabs.harvard.edu/abs/2000PhRvL..84..694T}

@ARTICLE{HansenL2000,
  author       = {{Hansen}, J.-P. and {L{\"o}wen}, H.},
  title        = {Effective interactions between electric double layers},
  journal      = ARoPC,
  publisher    = AR,
  year         = {2000},
  volume       = {51},
  number       = {1},
  pages        = {209-242},
  month        = OCT
}
% doi = {10.1146/annurev.physchem.51.1.209}
% url = {http://adsabs.harvard.edu/abs/2000ARPC...51..209H}

@ARTICLE{FreyK2005,
  author       = {{Frey}, E. and {Kroy}, K.},
  title        = {{B}rownian motion: a paradigm of soft matter and biological physics},
  journal      = AdPD,
  publisher    = WILEYVCH,
  year         = {2005},
  volume       = {517},
  number       = {1},
  pages        = {20-50},
  month        = FEB
}
% doi = {10.1002/andp.200410132}
% url = {http://adsabs.harvard.edu/abs/2005AnP...517...20F}

@INPROCEEDINGS{MorfillIBL2010,
  author       = {{Morfill}, G.~E. and {Ivlev}, A.~V. and {Brandt}, P. and {L{\"o}wen}, H.},
  title        = {Interdisciplinary research with complex plasmas},
  booktitle    = {Plasmas in the Laboratory and the Universe: Interactions, Patterns, and Turbulence},
  publisher    = AIoP,
  year         = {2010},
  editor       = {G.~Bertin and F.~de Luca and G.~Lodato and R.~Pozzoli and M.~Rom{\'e}},
  volume       = {1242},
  series       = {American Institute of Physics Conference Series},
  pages        = {67-79},
  address      = {Melville},
  month        = JUN,
  organization = AIoP
}
% 378 pages, AIP Press, 
% edition = {1},
% ISBN = {0-735-40787-8}
% EAN = {978-0-735-40787-9}
% doi = {10.1063/1.3460153}
% url = {http://adsabs.harvard.edu/abs/2010AIPC.1242...67M}

@ARTICLE{MorfillI2009,
  author       = {{Morfill}, G.~E. and {Ivlev}, A.~V.},
  title        = {Complex plasmas: an interdisciplinary research field},
  journal      = RoMP,
  publisher    = APS,
  year         = {2009},
  volume       = {81},
  number       = {4},
  pages        = {1353-1404},
  month        = OCT
}
% doi = {10.1103/RevModPhys.81.1353}
% url = {http://adsabs.harvard.edu/abs/2009RvMP...81.1353M}

@ARTICLE{OhnesorgeLW1994,
  author       = {{Ohnesorge}, R. and {L{\"o}wen}, H. and {Wagner}, H.},
  title        = {Density functional theory of crystal-fluid interfaces and surface melting},
  journal      = PRE,
  publisher    = APS,
  year         = {1994},
  volume       = {50},
  number       = {6},
  pages        = {4801-4809},
  month        = DEC
}
% doi = {10.1103/PhysRevE.50.4801}
% url = {http://adsabs.harvard.edu/abs/1994PhRvE..50.4801O}

@ARTICLE{DentonA1989,
  author       = {{Denton}, A.~R. and {Ashcroft}, N.~W.},
  title        = {Modified weighted-density-functional theory of nonuniform classical liquids},
  journal      = PRA,
  publisher    = APS,
  year         = {1989},
  volume       = {39},
  number       = {9},
  pages        = {4701-4708},
  month        = MAY
}
% doi = {10.1103/PhysRevA.39.4701}
% url = {http://adsabs.harvard.edu/abs/1989PhRvA..39.4701D}

@ARTICLE{CurtinA1985,
  author       = {{Curtin}, W.~A. and {Ashcroft}, N.~W.},
  title        = {Weighted-density-functional theory of inhomogeneous liquids and the freezing transition},
  journal      = PRA,
  publisher    = APS,
  year         = {1985},
  volume       = {32},
  number       = {5},
  pages        = {2909-2919},
  month        = NOV
}
% doi = {10.1103/PhysRevA.32.2909}
% url = {http://adsabs.harvard.edu/abs/1985PhRvA..32.2909C}

@ARTICLE{HowellEtAl2006,
  author       = {{Howell}, D.~A. and {Sullivan}, M. and {Nugent}, P.~E. and {Ellis}, R.~S. and {Conley}, A.~J. and {Le Borgne}, D. and {Carlberg}, R.~G. and {Guy}, J. and {Balam}, D. and {Basa}, S. and {Fouchez}, D. and {Hook}, I.~M. and {Hsiao}, E.~Y. and {Neill}, J.~D. and {Pain}, R. and {Perrett}, K.~M. and {Pritchet}, C.~J.},
  title        = {The type {Ia} supernova {SNLS}-{03D3bb} from a super-{C}handrasekhar-mass white dwarf star},
  journal      = NAT,
  publisher    = NPG,
  year         = {2006},
  volume       = {443},
  number       = {7109},
  pages        = {308-311},
  month        = SEP
}
% doi = {10.1038/nature05103}
% url = {http://adsabs.harvard.edu/abs/2006Natur.443..308H}

@ARTICLE{Perlmutter2003,
  author       = {{Perlmutter}, S.},
  title        = {Supernovae, Dark Energy and the Accelerating Universe},
  journal      = PT,
  publisher    = AIoP,
  year         = {2003},
  pages        = {53-60},
  month        = APR
}
% S-0031-9228-0304-030-4

@ARTICLE{AmanullahEtAl2010,
  author       = {{Amanullah}, R. and {Lidman}, C. and {Rubin}, D. and {Aldering}, G. and {Astier}, P. and {Barbary}, K. and {Burns}, M.~S. and {Conley}, A. and {Dawson}, K.~S. and {Deustua}, S.~E. and {Doi}, M. and {Fabbro}, S. and {Faccioli}, L. and {Fakhouri}, H.~K. and {Folatelli}, G. and {Fruchter}, A.~S. and {Furusawa}, H. and {Garavini}, G. and {Goldhaber}, G. and {Goobar}, A. and {Groom}, D.~E. and {Hook}, I. and {Howell}, D.~A. and {Kashikawa}, N. and {Kim}, A.~G. and {Knop}, R.~A. and {Kowalski}, M. and {Linder}, E. and {Meyers}, J. and {Morokuma}, T. and {Nobili}, S. and {Nordin}, J. and {Nugent}, P.~E. and {{\"O}stman}, L. and {Pain}, R. and {Panagia}, N. and {Perlmutter}, S. and {Raux}, J. and {Ruiz-Lapuente}, P. and {Spadafora}, A.~L. and {Strovink}, M. and {Suzuki}, N. and {Wang}, L. and {Wood-Vasey}, W.~M. and {Yasuda}, N. and {Supernova Cosmology Project}, T.},
  title        = {Spectra and Hubble Space Telescope Light Curves of Six Type Ia Supernovae at 0.511 {\lt} z {\lt} 1.12 and the Union2 Compilation},
  journal      = AphysJ,
  publisher    = IoPP,
  year         = {2010},
  volume       = {716},
  number       = {1},
  pages        = {712-738},
  month        = JUN
}
% doi = {10.1088/0004-637X/716/1/712}
% url = {http://adsabs.harvard.edu/abs/2010ApJ...716..712A}

@ARTICLE{PerlmutterEtAl1998,
  author       = {{Perlmutter}, S. and {Aldering}, G. and {della Valle}, M. and {Deustua}, S. and {Ellis}, R.~S. and {Fabbro}, S. and {Fruchter}, A. and {Goldhaber}, G. and {Groom}, D.~E. and {Hook}, I.~M. and {Kim}, A.~G. and {Kim}, M.~Y. and {Knop}, R.~A. and {Lidman}, C. and {McMahon}, R.~G. and {Nugent}, P. and {Pain}, R. and {Panagia}, N. and {Pennypacker}, C.~R. and {Ruiz-Lapuente}, P. and {Schaefer}, B. and {Walton}, N.},
  title        = {Discovery of a supernova explosion at half the age of the universe},
  journal      = NAT,
  publisher    = NPG,
  year         = {1998},
  volume       = {391},
  number       = {6662},
  pages        = {51-54},
  month        = JAN
}
% doi = {10.1038/34124}
% url = {http://adsabs.harvard.edu/abs/1998Natur.391...51P}

@ARTICLE{PerlmutterEtAl1999,
  author       = {{Perlmutter}, S. and {Aldering}, G. and {Goldhaber}, G. and {Knop}, R.~A. and {Nugent}, P. and {Castro}, P.~G. and {Deustua}, S. and {Fabbro}, S. and {Goobar}, A. and {Groom}, D.~E. and {Hook}, I.~M. and {Kim}, A.~G. and {Kim}, M.~Y. and {Lee}, J.~C. and {Nunes}, N.~J. and {Pain}, R. and {Pennypacker}, C.~R. and {Quimby}, R. and {Lidman}, C. and {Ellis}, R.~S. and {Irwin}, M. and {McMahon}, R.~G. and {Ruiz-Lapuente}, P. and {Walton}, N. and {Schaefer}, B. and {Boyle}, B.~J. and {Filippenko}, A.~V. and {Matheson}, T. and {Fruchter}, A.~S. and {Panagia}, N. and {Newberg}, H.~J.~M. and {Couch}, W.~J. and {The Supernova Cosmology Project}},
  title        = {Measurements of Omega and Lambda from 42 High-Redshift Supernovae},
  journal      = AphysJ,
  publisher    = IoPP,
  year         = {1999},
  volume       = {517},
  number       = {2},
  pages        = {565-586},
  month        = JUN
}
% doi = {10.1086/307221}
% url = {http://adsabs.harvard.edu/abs/1999ApJ...517..565P}

@MISC{Perlmutter2012,
  author       = {Saul Perlmutter},
  title        = {Supernova Cosmology Project},
  howpublished = {http://supernova.lbl.gov/},
  year         = {2012},
  month        = JAN
}

@ARTICLE{Leibundgut2011,
  author       = {Bruno Leibundgut},
  title        = {Eins-a-{V}ermessung des {U}niversums},
  journal      = PJ,
  publisher    = WILEYVCH,
  year         = {2011},
  volume       = {10},
  number       = {12},
  pages        = {27-29},
  month        = DEC
}

@article{GutscheKKRWH2008,
  title={Colloids dragged through a polymer solution: Experiment, theory, and simulation},
  author={Gutsche, Christof and Kremer, Friedrich and Kr{\"u}ger, Matthias and Rauscher, Markus and Weeber, Rudolf and Harting, Jens},
  journal=JoChemP,
  volume={129},
  number={8},
  pages={084902},
  year={2008},
  publisher={AIP}
}

@article{DuranYGK2017,
  title={General framework for fluctuating dynamic density functional theory},
  author={Dur{\'a}n-Olivencia, Miguel A and Yatsyshin, Peter and Goddard, Benjamin D and Kalliadasis, Serafim},
  journal=NJoP,
  volume={19},
  number={12},
  pages={123022},
  year={2017},
  publisher={IOP Publishing}
}

@article{LamorgeseM2016,
  title={Phase-field modeling of interfacial dynamics in emulsion flows: Nonequilibrium surface tension},
  author={Lamorgese, Andrea and Mauri, Roberto},
  journal={International Journal of Multiphase Flow},
  volume={85},
  pages={164--172},
  year={2016},
  publisher={Elsevier}
}


@article{SaltzmanS2006,
  title={Activated hopping and dynamical fluctuation effects in hard sphere suspensions and fluids},
  author={Saltzman, Erica J and Schweizer, Kenneth S},
  journal=JoChemP,
  volume={125},
  number={4},
  pages={044509},
  year={2006},
  publisher={AIP}
}

@article{CarrilloGPS2019,
  title={Long-time behaviour and phase transitions for the {M}c{K}ean--{V}lasov equation on the torus},
  author={Carrillo, JA and Gvalani, RS and Pavliotis, GA and Schlichting, A},
  journal={Archive for Rational Mechanics and Analysis},
  pages={1--56},
  year={2019},
  publisher={Springer}
}

@article{GomesKPY2019,
  title={Dynamics of the {D}esai-{Z}wanzig model in multiwell and random energy landscapes},
  author={Gomes, Susana N and Kalliadasis, Serafim and Pavliotis, Grigorios A and Yatsyshin, Petr},
  journal=PRE,
  volume={99},
  number={3},
  pages={032109},
  year={2019},
  publisher={APS}
}

@article{GoddardNSPK2012,
  title={General dynamical density functional theory for classical fluids},
  author={Goddard, Benjamin D and Nold, Andreas and Savva, Nikos and Pavliotis, Grigorios A and Kalliadasis, Serafim},
  journal=PRL,
  volume={109},
  number={12},
  pages={120603},
  year={2012},
  publisher={APS}
}

@article{DuranGK2016,
  title={Dynamical density functional theory for orientable colloids including inertia and hydrodynamic interactions},
  author={Dur{\'a}n-Olivencia, Miguel A and Goddard, Benjamin D and Kalliadasis, Serafim},
  journal={Journal of Statistical Physics},
  volume={164},
  number={4},
  pages={785--809},
  year={2016},
  publisher={Springer}
}


@article{WeiBADWNLBH2014b,
  title={Wechselwirkungen von {P}roteinen mit {P}olymerbeschichtungen und {B}iomaterialien},
  author={Wei, Qiang and Becherer, Tobias and Angioletti-Uberti, Stefano and Dzubiella, Joachim and Wischke, Christian and Neffe, Axel T and Lendlein, Andreas and Ballauff, Matthias and Haag, Rainer},
  journal={Angewandte Chemie},
  volume={126},
  number={31},
  pages={8138--8169},
  year={2014},
  publisher={Wiley Online Library}
}

@ARTICLE{vanTeeffelenLL2008,
  author       = {van~Teeffelen, Sven and Likos, Christos~N. and L\"owen, Hartmut},
  title        = {Colloidal Crystal Growth at Externally Imposed Nucleation Clusters},
  journal      = PRL,
  publisher    = APS,
  year         = {2008},
  volume       = {100},
  number       = {10},
  pages        = {108302},
  month        = MAR
}
% doi = {10.1103/PhysRevLett.100.108302}
% url = {http://link.aps.org/doi/10.1103/PhysRevLett.100.108302}

@ARTICLE{WittkowskiL2011,
  author       = {Wittkowski, Raphael and L{\"o}wen, Hartmut},
  title        = {Dynamical density functional theory for colloidal particles with arbitrary shape},
  journal      = MP,
  publisher    = TF,
  year         = {2011},
  volume       = {109},
  number       = {23-24},
  pages        = {2935-2943},
  month        = DEC
}
% doi = {10.1080/00268976.2011.609145}
% url = {http://www.tandfonline.com/doi/abs/10.1080/00268976.2011.609145}

@ARTICLE{HunterEEW2011,
  author       = {{Hunter}, G.~L. and {Edmond}, K.~V. and {Elsesser}, M.~T. and {Weeks}, E.~R.},
  title        = {Tracking rotational diffusion of colloidal clusters},
  journal      = OE,
  publisher    = OSoA,
  year         = {2011},
  volume       = {19},
  number       = {18},
  pages        = {17189-17202},
  month        = AUG
}
% doi = {10.1364/OE.19.017189}
% url = {http://adsabs.harvard.edu/abs/2011OExpr..1917189H}

@ARTICLE{WittkowskiL2012,
  author       = {Wittkowski, Raphael and L{\"o}wen, Hartmut},
  title        = {Self-propelled {B}rownian spinning top: dynamics of a biaxial swimmer at low {R}eynolds numbers},
  journal      = PRE,
  publisher    = APS,
  year         = {2012},
  volume       = {85},
  number       = {2},
  pages        = {021406},
  month        = FEB
}
% doi = {10.1103/PhysRevE.85.021406}
% url = {http://link.aps.org/doi/10.1103/PhysRevE.85.021406}

@ARTICLE{Crenshaw1996,
  author       = {Crenshaw, H.~C.},
  title        = {A new look at locomotion in microorganisms: rotating and translating},
  journal      = AZ,
  publisher    = ASoZ,
  year         = {1996},
  volume       = {36},
  number       = {6},
  pages        = {608-618},
  month        = DEC
}
% doi = {10.1093/icb/36.6.608}
% url = {http://icb.oxfordjournals.org/content/36/6/608.short}

@BOOK{Forster1989,
  author       = {Forster, Dieter},
  title        = {Hydrodynamic Fluctuations, Broken Symmetry, and Correlation Functions},
  publisher    = AW,
  year         = {1989},
  volume       = {47},
  series       = {Frontiers in Physics},
  address      = {Redwood City},
  edition      = {1},
  month        = OCT,
  pages        = {326},
  isbn         = {978-0-201-50882-6}
}

@BOOK{Forster1990,
  author       = {Forster, Dieter},
  title        = {Hydrodynamic Fluctuations, Broken Symmetry, and Correlation Functions},
  publisher    = PBP,
  year         = {1990},
  volume       = {10},
  series       = {Advanced Book Classics},
  address      = NY,
  edition      = {1},
  month        = JAN,
  pages        = {352},
  isbn         = {978-0-201-41049-5}
}
% ISBN-10: 0-201-41049-4

@BOOK{Zwanzig2001,
  author       = {Zwanzig, Robert},
  title        = {Nonequilibrium Statistical Mechanics},
  publisher    = OUP,
  year         = {2001},
  address      = NY,
  edition      = {3},
  month        = MAY,
  pages        = {240},
  isbn         = {978-0-195-14018-7}
}
% ISBN-10: 0-19-514018-4


@BOOK{Grabert1982,
  author       = {Grabert, Hermann},
  title        = {Projection Operator Techniques in Nonequilibrium Statistical Mechanics},
  publisher    = SPR,
  year         = {1982},
  volume       = {95},
  series       = {Springer Tracts in Modern Physics},
  address      = BER,
  edition      = {1},
  month        = AUG,
  pages        = {166},
  isbn         = {978-3-540-11635-6}
}
% ISBN-10: 3-540-11635-4

@ARTICLE{EbbensH2010,
  author       = {Ebbens, Stephen~J. and Howse, Jonathan~R.},
  title        = {In pursuit of propulsion at the nanoscale},
  journal      = SM,
  publisher    = RSoC,
  year         = {2010},
  volume       = {6},
  number       = {4},
  pages        = {726-738},
  month        = FEB
}
% doi = {10.1039/B918598D}
% url = {http://pubs.rsc.org/en/Content/ArticleLanding/2010/SM/b918598d}

@ARTICLE{GarciadelaTorreREO2007,
  author       = {{Garc\'{\i}a de la Torre}, J. and {del Rio Echenique}, G. and Ortega, A.},
  title        = {Improved calculation of rotational diffusion and intrinsic viscosity of bead models for macromolecules and nanoparticles},
  journal      = JoPChemB,
  publisher    = ACS,
  year         = {2007},
  volume       = {111},
  number       = {5},
  pages        = {955-961},
  month        = FEB
}
% doi = {10.1021/jp0647941}
% url = {http://pubs.acs.org/doi/abs/10.1021/jp0647941}

@ARTICLE{delaTorreNMDC1994,
  author       = {{Garc\'{\i}a de la Torre}, J. and Navarro, S. and Martinez, M.~C.~Lopez and Diaz, F.~G. and Cascales, J.~J.~Lopez},
  title        = {HYDRO: a computer program for the prediction of hydrodynamic properties of macromolecules},
  journal      = BJ,
  publisher    = ELS,
  year         = {1994},
  volume       = {67},
  number       = {2},
  pages        = {530-531},
  month        = AUG
}
% doi = {10.1016/S0006-3495(94)80512-0}
% url = {http://www.cell.com/biophysj/abstract/S0006-3495(94)80512-0}

@ARTICLE{HarveydlT1980,
  author       = {Harvey, S. and {Garc\'{\i}a de la Torre}, J.},
  title        = {Coordinate systems for modeling the hydrodynamic resistance and diffusion coefficients of irregularly shaped rigid macromolecules},
  journal      = MAC,
  publisher    = ACS,
  year         = {1980},
  volume       = {13},
  number       = {4},
  pages        = {960-964},
  month        = JUL
}
% doi = {10.1021/ma60076a037}
% url = {http://pubs.acs.org/doi/abs/10.1021/ma60076a037}

@ARTICLE{RotneP1969,
  author       = {Rotne, J. and Prager, S.},
  title        = {Variational treatment of hydrodynamic interaction in polymers},
  journal      = JoChemP,
  publisher    = AIoP,
  year         = {1969},
  volume       = {50},
  number       = {11},
  pages        = {4831-4837},
  month        = JUN
}
% doi = {10.1063/1.1670977}
% url = {http://jcp.aip.org/resource/1/jcpsa6/v50/i11/p4831_s1}

@ARTICLE{Mori1965,
  author       = {Mori, Hazime},
  title        = {Transport, collective motion, and {B}rownian motion},
  journal      = PoTP,
  publisher    = YHaKU,
  year         = {1965},
  volume       = {33},
  number       = {3},
  pages        = {423-455},
  month        = MAR
}
% doi = {10.1143/PTP.33.423}
% url = {http://ptp.ipap.jp/link?PTP/33/423}

@article{Popper1956,
  title={The arrow of time},
  author={Popper, Karl R.},
  journal={Nature},
  volume={177},
  number={4507},
  year={1956},
  pages={538},
  publisher={Nature Publishing Group}
}

@ARTICLE{ZwanzigM1965,
  author       = {Zwanzig, Robert and Mountain, Raymond~D.},
  title        = {High-frequency elastic moduli of simple fluids},
  journal      = JoChemP,
  publisher    = AIoP,
  year         = {1965},
  volume       = {43},
  number       = {12},
  pages        = {4464-4471},
  month        = DEC
}
% doi = {10.1063/1.1696718}
% url = {http://jcp.aip.org/resource/1/jcpsa6/v43/i12/p4464_s1?isAuthorized=no}

@article{Zwanzig1960,
author = {Zwanzig, Robert},
title = {Ensemble method in the theory of irreversibility},
journal = JoChemP,
volume = {33},
number = {5},
year = {1960},
pages = {1338-1341}
}

@ARTICLE{Zwanzig1974,
  author       = {{Zwanzig}, R.},
  title        = {Theoretical basis for the {R}ouse-{Z}imm model in polymer solution dynamics},
  journal      = JoChemP,
  publisher    = AIoP,
  year         = {1974},
  volume       = {60},
  number       = {7},
  pages        = {2717-2720},
  month        = APR
}
% doi = {10.1063/1.1681433}
% url = {http://adsabs.harvard.edu/abs/1974JChPh..60.2717Z}

@ARTICLE{BixonZ1978,
  author       = {{Bixon}, M. and {Zwanzig}, R.},
  title        = {Optimized {R}ouse-{Z}imm theory for stiff polymers},
  journal      = JoChemP,
  publisher    = AIoP,
  year         = {1978},
  volume       = {68},
  number       = {4},
  pages        = {1896-1902},
  month        = FEB
}
% doi = {10.1063/1.435916}
% url = {http://adsabs.harvard.edu/abs/1978JChPh..68.1896B}

@ARTICLE{ReysDGMLBD1988,
  author       = {{Reys}, V. and {Dormoy}, Y. and {Gallani}, J.~L. and {Martinoty}, P. and {Le Barny}, P. and {Dubois}, J.~C.},
  title        = {Short-range-order effects in the isotropic phase of a side-chain polymeric liquid crystal},
  journal      = PRL,
  publisher    = APS,
  year         = {1988},
  volume       = {61},
  number       = {20},
  pages        = {2340-2343},
  month        = NOV
}
% doi = {10.1103/PhysRevLett.61.2340}
% url = {http://adsabs.harvard.edu/abs/1988PhRvL..61.2340R}

@ARTICLE{ReysDCKM1992,
  author       = {{Reys}, V. and {Dormoy}, Y. and {Collin}, D. and {Keller}, P. and {Martinoty}, P.},
  title        = {Kerr effect in the isotropic phase of a side-chain polymeric liquid crystal},
  journal      = JdPII,
  publisher    = EDPS,
  year         = {1992},
  volume       = {2},
  number       = {2},
  pages        = {209-218},
  month        = FEB
}
% doi = {10.1051/jp2:1992124}
% url = {http://adsabs.harvard.edu/abs/1992JPhy2...2..209R}

@ARTICLE{SigelSSL1993,
  author       = {{Sigel}, R. and {Stille}, W. and {Strobl}, G. and {Lehnert}, R.},
  title        = {Near-critical behavior of the dynamic stress-optical coefficient in liquid crystalline elastomers},
  journal      = MAC,
  publisher    = ACS,
  year         = {1993},
  volume       = {26},
  number       = {16},
  pages        = {4226-4233},
  month        = AUG
}
% doi = {10.1021/ma00068a024}
% url = {http://adsabs.harvard.edu/abs/1993MaMol..26.4226S}

@ARTICLE{VerhoeffOvdSL2011,
  author       = {{Verhoeff}, A.~A. and {Otten}, R.~H.~J. and {van der Schoot}, P. and {Lekkerkerker}, H.~N.~W.},
  title        = {Magnetic field effects on tactoids of plate-like colloids},
  journal      = JoChemP,
  publisher    = AIoP,
  year         = {2011},
  volume       = {134},
  number       = {4},
  pages        = {044904},
  month        = JAN
}
% doi = {10.1063/1.3520389}
% url = {http://adsabs.harvard.edu/abs/2011JChPh.134d4904V}

@ARTICLE{UllrichASV2010,
  author       = {Ullrich, B. and Auernhammer, G.~K. and Sam, E.~M. and Vollmer, D.},
  title        = {Tracer colloids close to an isotropic-nematic domain interface with phase transition-induced solute transport},
  journal      = CaSA:PaEA,
  publisher    = ELS,
  year         = {2010},
  volume       = {354},
  number       = {1-3},
  pages        = {298-307},
  month        = FEB
}
% doi = {10.1016/j.colsurfa.2009.11.023}
% url = {http://www.sciencedirect.com/science/article/pii/S0927775709006736}

@ARTICLE{ReichDvRS2007,
  author       = {Reich, H. and Dijkstra, M. and {van Roij}, R. and Schmidt, M.},
  title        = {Entropic wetting and the free isotropic-nematic interface of hard colloidal platelets},
  journal      = JoPChemB,
  publisher    = ACS,
  year         = {2007},
  volume       = {111},
  number       = {27},
  pages        = {7825-7835},
  month        = JUL
}
% doi = {10.1021/jp068870b}
% url = {http://pubs.acs.org/doi/abs/10.1021/jp068870b}

@ARTICLE{WolfsheimerTSvRS2006,
  author       = {{Wolfsheimer}, S. and {Tanase}, C. and {Shundyak}, K. and {van Roij}, R. and {Schilling}, T.},
  title        = {Isotropic-nematic interface in suspensions of hard rods: mean-field properties and capillary waves},
  journal      = PRE,
  publisher    = APS,
  year         = {2006},
  volume       = {73},
  number       = {6},
  pages        = {061703},
  month        = JUN
}
% doi = {10.1103/PhysRevE.73.061703}
% url = {http://adsabs.harvard.edu/abs/2006PhRvE..73f1703W}

@ARTICLE{VinkS2005,
  author       = {{Vink}, R.~L.~C. and {Schilling}, T.},
  title        = {Interfacial tension of the isotropic-nematic interface in suspensions of soft spherocylinders},
  journal      = PRE,
  publisher    = APS,
  year         = {2005},
  volume       = {71},
  number       = {5},
  pages        = {051716},
  month        = MAY
}
% doi = {10.1103/PhysRevE.71.051716}
% url = {http://adsabs.harvard.edu/abs/2005PhRvE..71e1716V}

@ARTICLE{BierHD2004,
  author       = {{Bier}, M. and {Harnau}, L. and {Dietrich}, S.},
  title        = {Bulk and interfacial properties of binary hard-platelet fluids},
  journal      = PRE,
  publisher    = APS,
  year         = {2004},
  volume       = {69},
  number       = {2},
  pages        = {021506},
  month        = FEB
}
% doi = {10.1103/PhysRevE.69.021506}
% url = {http://adsabs.harvard.edu/abs/2004PhRvE..69b1506B}

@ARTICLE{VelascoMS2002,
  author       = {{Velasco}, E. and {Mederos}, L. and {Sullivan}, D.~E.},
  title        = {Density-functional study of the nematic-isotropic interface of hard spherocylinders},
  journal      = PRE,
  publisher    = APS,
  year         = {2002},
  volume       = {66},
  number       = {2},
  pages        = {021708},
  month        = AUG
}
% doi = {10.1103/PhysRevE.66.021708}
% url = {http://adsabs.harvard.edu/abs/2002PhRvE..66b1708V}

@ARTICLE{Blanc2001,
  author       = {{Blanc}, C.},
  title        = {Measurement of the anisotropy of the energy of an isotropic-smectic interface in a smectic cylinder: application to the {L}$_{3}$-{L}$_{\alpha}$ interface},
  journal      = PRE,
  publisher    = APS,
  year         = {2001},
  volume       = {64},
  number       = {1},
  pages        = {011702},
  month        = JUL
}
% doi = {10.1103/PhysRevE.64.011702}
% url = {http://adsabs.harvard.edu/abs/2001PhRvE..64a1702B}

@ARTICLE{OsipovSR2003,
  author       = {Osipov, M.~A. and Sambles, J.~R. and Ruan, L.},
  title        = {Theory of nematic-smectic phase separation in thin twisted liquid crystal cells},
  journal      = LC,
  publisher    = TF,
  year         = {2003},
  volume       = {30},
  number       = {7},
  pages        = {823-830},
  month        = JUL
}
% doi = {10.1080/0267829031000111450}
% url = {http://www.tandfonline.com/doi/abs/10.1080/0267829031000111450}

@ARTICLE{MederosS1992,
  author       = {{Mederos}, L. and {Sullivan}, D.~E.},
  title        = {Smectic-{A} ordering at a liquid-vapor interface},
  journal      = PRA,
  publisher    = APS,
  year         = {1992},
  volume       = {46},
  number       = {12},
  pages        = {7700-7707},
  month        = DEC
}
% doi = {10.1103/PhysRevA.46.7700}
% url = {http://adsabs.harvard.edu/abs/1992PhRvA..46.7700M}

@ARTICLE{BrownAdRM1998,
  author       = {{Brown}, J.~T. and {Allen}, M.~P. and {del R{\'{\i}}o}, E.~M. and {Miguel}, E.~D.},
  title        = {Effects of elongation on the phase behavior of the Gay-Berne fluid},
  journal      = PRE,
  publisher    = APS,
  year         = {1998},
  volume       = {57},
  number       = {6},
  pages        = {6685-6699},
  month        = JUN
}
% doi = {10.1103/PhysRevE.57.6685}
% url = {http://adsabs.harvard.edu/abs/1998PhRvE..57.6685B}

@ARTICLE{SomozaMS1995,
  author       = {{Somoza}, A.~M. and {Mederos}, L. and {Sullivan}, D.~E.},
  title        = {Wetting and layering transitions in liquid crystals},
  journal      = PRE,
  publisher    = APS,
  year         = {1995},
  volume       = {52},
  number       = {5},
  pages        = {5017-5027},
  month        = NOV
}
% doi = {10.1103/PhysRevE.52.5017}
% url = {http://adsabs.harvard.edu/abs/1995PhRvE..52.5017S}

@ARTICLE{SchillingF2004b,
  author       = {Schilling, Tanja and Frenkel, Daan},
  title        = {Self-poisoning of crystal nuclei in hard-rod liquids},
  journal      = PRL,
  publisher    = APS,
  year         = {2004},
  volume       = {92},
  number       = {8},
  pages        = {085505},
  month        = FEB
}
% doi = {10.1103/PhysRevLett.92.085505}
% url = {http://link.aps.org/doi/10.1103/PhysRevLett.92.085505}

@ARTICLE{DogicF2001,
  author       = {Dogic, Z. and Fraden, S.},
  title        = {Development of model colloidal liquid crystals and the kinetics of the isotropic-smectic transition},
  journal      = PTotRSA:MPaES,
  publisher    = RS,
  year         = {2001},
  volume       = {359},
  number       = {1782},
  pages        = {997-1014},
  month        = MAY
}
% doi = {10.1098/rsta.2000.0814 }
% url = {http://rsta.royalsocietypublishing.org/content/359/1782/997.full.pdf+html}

@ARTICLE{VerhoeffL2012,
  author       = {Verhoeff, A.~A. and Lekkerkerker, H.~N.~W.},
  title        = {Direct observation of columnar liquid crystal droplets},
  journal      = SM,
  publisher    = RSoC,
  year         = {2012},
  volume       = {8},
  number       = {18},
  pages        = {4865-4868},
  month        = APR
}
% doi = {10.1039/C2SM25208B}
% url = {http://pubs.rsc.org/en/content/articlelanding/2012/sm/c2sm25208b}

@ARTICLE{Palberg1999,
  author       = {{Palberg}, T.},
  title        = {Crystallization kinetics of repulsive colloidal spheres},
  journal      = JoP:CM,
  publisher    = IoPP,
  year         = {1999},
  volume       = {11},
  number       = {28},
  pages        = {R323-R360},
  month        = JUL
}
% doi = {10.1088/0953-8984/11/28/201}
% url = {http://adsabs.harvard.edu/abs/1999JPCM...11R.323P}

@BOOK{Woodruff1980,
  author       = {Woodruff, D.~P.},
  title        = {The Solid-Liquid Interface},
  publisher    = CUP,
  year         = {1980},
  volume       = {1},
  series       = {Cambridge Solid State Science Series},
  address      = LON,
  edition      = {1},
  month        = NOV,
  pages        = {192},
  isbn         = {0-521-29971-3}
}
% EAN: 9780521299718

@ARTICLE{MarrG1993,
  author       = {{Marr}, D.~W. and {Gast}, A.~P.},
  title        = {Planar density-functional approach to the solid-fluid interface of simple liquids},
  journal      = PRE,
  publisher    = APS,
  year         = {1993},
  volume       = {47},
  number       = {2},
  pages        = {1212-1221},
  month        = FEB
}
% doi = {10.1103/PhysRevE.47.1212}
% url = {http://adsabs.harvard.edu/abs/1993PhRvE..47.1212M}

@ARTICLE{Curtin1987,
  author       = {{Curtin}, W.~A.},
  title        = {Density-functional theory of the solid-liquid interface},
  journal      = PRL,
  publisher    = APS,
  year         = {1987},
  volume       = {59},
  number       = {11},
  pages        = {1228-1231},
  month        = SEP
}
% doi = {10.1103/PhysRevLett.59.1228}
% url = {http://adsabs.harvard.edu/abs/1987PhRvL..59.1228C}

@ARTICLE{Curtin1989,
  author       = {{Curtin}, W.~A.},
  title        = {Density-functional theory of crystal-melt interfaces},
  journal      = PRB,
  publisher    = APS,
  year         = {1989},
  volume       = {39},
  number       = {10},
  pages        = {6775-6791},
  month        = APR
}
% doi = {10.1103/PhysRevB.39.6775}
% url = {http://adsabs.harvard.edu/abs/1989PhRvB..39.6775C}

@ARTICLE{HaertelOREHL2012,
  author       = {H{\"a}rtel, A. and Oettel, M. and Rozas, R.~E. and Egelhaaf, S.~U. and Horbach, J. and L{\"o}wen, H.},
  title        = {Tension and stiffness of the hard sphere crystal-fluid interface},
  journal      = PRL,
  publisher    = APS,
  year         = {2012},
  volume       = {108},
  number       = {22},
  pages        = {226101},
  month        = MAY
}
% doi = {10.1103/PhysRevLett.108.226101}
% url = {http://link.aps.org/doi/10.1103/PhysRevLett.108.226101}

@ARTICLE{HoytAK2001,
  author       = {{Hoyt}, J.~J. and {Asta}, M. and {Karma}, A.},
  title        = {Method for computing the anisotropy of the solid-liquid interfacial free energy},
  journal      = PRL,
  publisher    = APS,
  year         = {2001},
  volume       = {86},
  number       = {24},
  pages        = {5530-5533},
  month        = JUN
}
% doi = {10.1103/PhysRevLett.86.5530}
% url = {http://adsabs.harvard.edu/abs/2001PhRvL..86.5530H}

@ARTICLE{BinderM2000,
  author       = {{Binder}, K. and {M{\"u}ller}, M.},
  title        = {Computer simulation of profiles of interfaces between coexisting phases},
  journal      = IJoMPC,
  publisher    = WSP,
  year         = {2000},
  volume       = {11},
  number       = {6},
  pages        = {1093-1113},
  month        = SEP
}
% doi = {10.1142/S012918310000095X}
% url = {http://adsabs.harvard.edu/abs/2000IJMPC..11.1093B}

@ARTICLE{DavidchackML2006,
  author       = {{Davidchack}, R.~L. and {Morris}, J.~R. and {Laird}, B.~B.},
  title        = {The anisotropic hard-sphere crystal-melt interfacial free energy from fluctuations},
  journal      = JoChemP,
  publisher    = AIoP,
  year         = {2006},
  volume       = {125},
  number       = {9},
  pages        = {094710},
  month        = SEP
}
% doi = {10.1063/1.2338303}
% url = {http://adsabs.harvard.edu/abs/2006JChPh.125i4710D}

@ARTICLE{ZykovaTimanRHB2009,
  author       = {{Zykova-Timan}, T. and {Rozas}, R.~E. and {Horbach}, J. and {Binder}, K.},
  title        = {Computer simulation studies of finite-size broadening of solid-liquid interfaces: from hard spheres to nickel},
  journal      = JoChemP,
  publisher    = AIoP,
  year         = {2009},
  volume       = {21},
  number       = {46},
  pages        = {464102},
  month        = NOV
}
% doi = {10.1088/0953-8984/21/46/464102}
% url = {http://adsabs.harvard.edu/abs/2009JPCM...21T4102Z}

@ARTICLE{ZykovaTimanHB2010,
  author       = {{Zykova-Timan}, T. and {Horbach}, J. and {Binder}, K.},
  title        = {{M}onte {C}arlo simulations of the solid-liquid transition in hard spheres and colloid-polymer mixtures},
  journal      = JoP:CM,
  publisher    = IoPP,
  year         = {2010},
  volume       = {133},
  number       = {1},
  pages        = {014705},
  month        = JUL
}
% doi = {10.1063/1.3455504}
% url = {http://adsabs.harvard.edu/abs/2010JChPh.133a4705Z}

@ARTICLE{RozasH2011,
  author       = {{Rozas}, R.~E. and {Horbach}, J.},
  title        = {Capillary wave analysis of rough solid-liquid interfaces in nickel},
  journal      = EPL,
  publisher    = EPS,
  year         = {2011},
  volume       = {93},
  number       = {2},
  pages        = {26006},
  month        = JAN
}
% doi = {10.1209/0295-5075/93/26006}
% url = {http://adsabs.harvard.edu/abs/2011EL.....9326006R}

@ARTICLE{ClunieB1999,
  author       = {Clunie, John~C. and Baird, James~K.},
  title        = {Interdiffusion coefficient and dynamic viscosity for the mixture 2,6-lutidine + water near the lower consolute point},
  journal      = PaCoL,
  publisher    = TF,
  year         = {1999},
  volume       = {37},
  number       = {4},
  pages        = {357-371},
  month        = JUL
}
% doi = {10.1080/00319109908031441}
% url = {http://www.tandfonline.com/doi/abs/10.1080/00319109908031441}

@ARTICLE{Wiegand2004,
  author       = {{Wiegand}, S.},
  title        = {Thermal diffusion in liquid mixtures and polymer solutions},
  journal      = JoP:CM,
  publisher    = IoPP,
  year         = {2004},
  volume       = {16},
  number       = {10},
  pages        = {R357-R379},
  month        = MAR
}
% doi = {10.1088/0953-8984/16/10/R02}
% url = {http://adsabs.harvard.edu/abs/2004JPCM...16R.357W}

@ARTICLE{Wuerger2010,
  author       = {{W{\"u}rger}, A.},
  title        = {Thermal non-equilibrium transport in colloids},
  journal      = RoPiP,
  publisher    = IoPP,
  year         = {2010},
  volume       = {73},
  number       = {12},
  pages        = {126601},
  month        = DEC
}
% doi = {10.1088/0034-4885/73/12/126601}
% url = {http://adsabs.harvard.edu/abs/2010RPPh...73l6601W}

@ARTICLE{PiazzaP2008,
  author       = {{Piazza}, R. and {Parola}, A.},
  title        = {Thermophoresis in colloidal suspensions},
  journal      = JoP:CM,
  publisher    = IoPP,
  year         = {2008},
  volume       = {20},
  number       = {15},
  pages        = {153102},
  month        = APR
}
% doi = {10.1088/0953-8984/20/15/153102}
% url = {http://adsabs.harvard.edu/abs/2008JPCM...20o3102P}

@ARTICLE{Piazza2004,
  author       = {{Piazza}, R.},
  title        = {'Thermal forces': colloids in temperature gradients},
  journal      = JoP:CM,
  publisher    = IoPP,
  year         = {2004},
  volume       = {16},
  number       = {38},
  pages        = {S4195-S4211},
  month        = SEP
}
% doi = {10.1088/0953-8984/16/38/032}
% url = {http://adsabs.harvard.edu/abs/2004JPCM...16S4195P}

@ARTICLE{ArcherHS2007,
  author       = {{Archer}, A.~J. and {Hopkins}, P. and {Schmidt}, M.},
  title        = {Dynamics in inhomogeneous liquids and glasses via the test particle limit},
  journal      = PRE,
  publisher    = APS,
  year         = {2007},
  volume       = {75},
  number       = {4},
  pages        = {040501},
  month        = APR
}
% doi = {10.1103/PhysRevE.75.040501}
% url = {http://adsabs.harvard.edu/abs/2007PhRvE..75d0501A}

@ARTICLE{RoyallDSvB2007,
  author       = {{Royall}, C.~P. and {Dzubiella}, J. and {Schmidt}, M. and {van Blaaderen}, A.},
  title        = {Nonequilibrium sedimentation of colloids on the particle scale},
  journal      = PRL,
  publisher    = APS,
  year         = {2007},
  volume       = {98},
  number       = {18},
  pages        = {188304},
  month        = MAY
}
% doi = {10.1103/PhysRevLett.98.188304}
% url = {http://adsabs.harvard.edu/abs/2007PhRvL..98r8304R}

@ARTICLE{BiervR2007,
  author       = {{Bier}, M. and {van Roij}, R.},
  title        = {Relaxation dynamics in fluids of platelike colloidal particles},
  journal      = PRE,
  publisher    = APS,
  year         = {2007},
  volume       = {76},
  number       = {2},
  pages        = {021405},
  month        = AUG
}
% doi = {10.1103/PhysRevE.76.021405}
% url = {http://adsabs.harvard.edu/abs/2007PhRvE..76b1405B}

@ARTICLE{BiervRDvdS2008,
  author       = {{Bier}, M. and {van Roij}, R. and {Dijkstra}, M. and {van der Schoot}, P.},
  title        = {Self-diffusion of particles in complex fluids: temporary cages and permanent barriers},
  journal      = PRL,
  publisher    = APS,
  year         = {2008},
  volume       = {101},
  number       = {21},
  pages        = {215901},
  month        = NOV
}
% doi = {10.1103/PhysRevLett.101.215901}
% url = {http://adsabs.harvard.edu/abs/2008PhRvL.101u5901B}

@ARTICLE{RothRA2009,
  author       = {{Roth}, R. and {Rauscher}, M. and {Archer}, A.~J.},
  title        = {Selectivity in binary fluid mixtures: static and dynamical properties},
  journal      = PRE,
  publisher    = APS,
  year         = {2009},
  volume       = {80},
  number       = {2},
  pages        = {021409},
  month        = AUG
}
% doi = {10.1103/PhysRevE.80.021409}
% url = {http://adsabs.harvard.edu/abs/2009PhRvE..80b1409R}

@ARTICLE{KahlL2009,
  author       = {{Kahl}, G. and {L{\"o}wen}, H.},
  title        = {Classical density functional theory: an ideal tool to study heterogeneous crystal nucleation},
  journal      = JoP:CM,
  publisher    = IoPP,
  year         = {2009},
  volume       = {21},
  number       = {46},
  pages        = {464101},
  month        = NOV
}
% doi = {10.1088/0953-8984/21/46/464101}
% url = {http://adsabs.harvard.edu/abs/2009JPCM...21T4101K}

@article{ThieleAP2012,
  title={Thermodynamically consistent description of the hydrodynamics of free surfaces covered by insoluble surfactants of high concentration},
  author={Thiele, Uwe and Archer, Andrew J and Plapp, Mathis},
  journal=PoF,
  volume={24},
  number={10},
  pages={102107},
  year={2012},
  publisher={AIP}
}

@article{LeeKVG2015,
  title={Dynamics of ion transport in ionic liquids},
  author={Lee, Alpha A and Kondrat, Svyatoslav and Vella, Dominic and Goriely, Alain},
  journal=PRL,
  volume={115},
  number={10},
  pages={106101},
  year={2015},
  publisher={APS}
}

@article{GuanWW2014,
  title={A convergent convex splitting scheme for the periodic nonlocal {Cahn-Hilliard} equation},
  author={Guan, Zhen and Wang, Cheng and Wise, Steven M},
  journal={Numerische Mathematik},
  volume={128},
  number={2},
  pages={377--406},
  year={2014},
  publisher={Springer}
}

@article{DonevVE2014,
  title={Dynamic density functional theory with hydrodynamic interactions and fluctuations},
  author={Donev, Aleksandar and Vanden-Eijnden, Eric},
  journal=JoChemP,
  volume={140},
  number={23},
  pages={234115},
  year={2014},
  publisher={AIP}
}

@article{ArcherR2004,
  title={Dynamical density functional theory for interacting {B}rownian particles: stochastic or deterministic?},
  author={Archer, Andrew~J. and Rauscher, Markus},
  journal={Journal of Physics A: Mathematical and General},
  volume={37},
  number={40},
  pages={9325},
  year={2004},
  publisher={IOP Publishing}
}

@ARTICLE{ArcherRT2010,
  author       = {{Archer}, A.~J. and {Robbins}, M.~J. and {Thiele}, U.},
  title        = {Dynamical density functional theory for the dewetting of evaporating thin films of nanoparticle suspensions exhibiting pattern formation},
  journal      = PRE,
  publisher    = APS,
  year         = {2010},
  volume       = {81},
  number       = {2},
  pages        = {021602},
  month        = FEB
}
% doi = {10.1103/PhysRevE.81.021602}
% url = {http://adsabs.harvard.edu/abs/2010PhRvE..81b1602A}

@article{KinjoH2007,
  title={Equation of motion for coarse-grained simulation based on microscopic description},
  author={Kinjo, Tomoyuki and Hyodo, Shi-aki},
  journal=PRE,
  volume={75},
  number={5},
  pages={051109},
  year={2007},
  publisher={APS}
}

@ARTICLE{HuangEP2010,
  author       = {{Huang}, Z.-F. and {Elder}, K.~R. and {Provatas}, N.},
  title        = {Phase-field-crystal dynamics for binary systems: derivation from dynamical density functional theory, amplitude equation formalism, and applications to alloy heterostructures},
  journal      = PRE,
  publisher    = APS,
  year         = {2010},
  volume       = {82},
  number       = {2},
  pages        = {021605},
  month        = AUG
}
% doi = {10.1103/PhysRevE.82.021605}
% url = {http://adsabs.harvard.edu/abs/2010PhRvE..82b1605H}

@ARTICLE{LichtnerAK2012,
  author       = {Lichtner, K. and Archer, A.~J. and Klapp, S.~H.~L.},
  title        = {Phase separation dynamics in a two-dimensional magnetic mixture},
  journal      = JoChemP,
  publisher    = AIoP,
  year         = {2012},
  volume       = {136},
  number       = {2},
  pages        = {024502},
  month        = JAN
}
% doi = {http://dx.doi.org/10.1063/1.3674270}
% url = {http://jcp.aip.org/resource/1/jcpsa6/v136/i2/p024502_s1?isAuthorized=no}

@ARTICLE{VissersWRLRIvB2011,
  author       = {Vissers, Teun and Wysocki, Adam and Rex, Martin and L{\"o}wen, Hartmut and Royall, C.~Patrick and Imhof, Arnout and {van Blaaderen}, Alfons},
  title        = {Lane formation in driven mixtures of oppositely charged colloids},
  journal      = SM,
  publisher    = RSoC,
  year         = {2011},
  volume       = {7},
  number       = {6},
  pages        = {2352-2356},
  month        = FEB
}
% doi = {10.1039/C0SM01343A}
% url = {http://dx.doi.org/10.1039/C0SM01343A}

@ARTICLE{MilinkovicPD2011,
  author       = {Milinkovic, Kristina and Padding, Johan~T. and Dijkstra, Marjolein},
  title        = {Hydrodynamic {R}ayleigh-{T}aylor-like instabilities in sedimenting colloidal mixtures},
  journal      = SM,
  publisher    = RSoC,
  year         = {2011},
  volume       = {7},
  number       = {23},
  pages        = {11177-11186},
  month        = FEB
}
% doi = {10.1039/C1SM05930K}
% url = {http://pubs.rsc.org/en/Content/ArticleLanding/2011/SM/c1sm05930k}

@article{OettelGHLRS2010,
  title={Free energies, vacancy concentrations, and density distribution anisotropies in hard-sphere crystals: A combined density functional and simulation study},
  author={Oettel, M and G{\"o}rig, S and H{\"a}rtel, A and L{\"o}wen, H and Radu, Marc and Schilling, Tanja},
  journal=PRE,
  volume={82},
  number={5},
  pages={051404},
  year={2010},
  publisher={APS}
}

@article{TagliazucchiRS2011,
  title={Ion transport and molecular organization are coupled in polyelectrolyte-modified nanopores},
  author={Tagliazucchi, Mario and Rabin, Yitzhak and Szleifer, Igal},
  journal={Journal of the American Chemical Society},
  volume={133},
  number={44},
  pages={17753--17763},
  year={2011},
  publisher={ACS Publications}
}

@ARTICLE{Loewen2010c,
  author       = {L{\"o}wen, Hartmut},
  title        = {Particle-resolved instabilities in colloidal dispersions},
  journal      = SM,
  publisher    = RSoC,
  year         = {2010},
  volume       = {6},
  number       = {14},
  pages        = {3133-3142},
  month        = APR
}
% doi = {10.1039/B923685F}
% url = {http://pubs.rsc.org/en/Content/ArticleLanding/2010/SM/b923685f}

@ARTICLE{Lionberger2002,
  author       = {{Lionberger}, R.~A.},
  title        = {Viscosity of bimodal and polydisperse colloidal suspensions},
  journal      = PRE,
  publisher    = APS,
  year         = {2002},
  volume       = {65},
  number       = {6},
  pages        = {061408},
  month        = JUN
}
% doi = {10.1103/PhysRevE.65.061408}
% url = {http://adsabs.harvard.edu/abs/2002PhRvE..65f1408L}

@ARTICLE{NaegeleD1998,
  author       = {N{\"a}gele, G. and Dhont, J.~K.~G.},
  title        = {Tracer-diffusion in colloidal mixtures: a mode-coupling scheme with hydrodynamic interactions},
  journal      = JoChemP,
  publisher    = AIoP,
  year         = {1998},
  volume       = {108},
  number       = {22},
  pages        = {9566-9576},
  month        = JUN
}
% doi = {10.1063/1.476405}
% url = {http://jcp.aip.org/resource/1/jcpsa6/v108/i22/p9566_s1?isAuthorized=no}

@ARTICLE{RexL2008b,
  author       = {{Rex}, M. and {L{\"o}wen}, H.},
  title        = {Influence of hydrodynamic interactions on lane formation in oppositely charged driven colloids},
  journal      = EPJE,
  publisher    = EDPS,
  year         = {2008},
  volume       = {26},
  number       = {1-2},
  pages        = {143-150},
  month        = MAY
}
% doi = {10.1140/epje/i2007-10274-4}
% url = {http://adsabs.harvard.edu/abs/2008EPJE...26..143R}

@ARTICLE{KleinN2010,
  author       = {Klein, Rudolf and N{\"a}gele, Gerhard},
  title        = {Static and dynamic scattering by concentrated systems: theory},
  journal      = COiCaIS,
  publisher    = ELS,
  year         = {2010},
  volume       = {1},
  number       = {1},
  pages        = {4-10},
  month        = FEB
}
% doi = {10.1016/S1359-0294(96)80037-9}
% url = {http://www.sciencedirect.com/science/article/pii/S1359029496800379}

@ARTICLE{KaplanYP1992,
  author       = {{Kaplan}, P.~D. and {Yodh}, A.~G. and {Pine}, D.~J.},
  title        = {Diffusion and structure in dense binary suspensions},
  journal      = PRL,
  publisher    = APS,
  year         = {1992},
  volume       = {68},
  number       = {3},
  pages        = {393-396},
  month        = JAN
}
% doi = {10.1103/PhysRevLett.68.393}
% url = {http://adsabs.harvard.edu/abs/1992PhRvL..68..393K}

@ARTICLE{WagnerK1991,
  author       = {Wagner, N.~J. and Klein, R.},
  title        = {The rheology and microstructure of charged colloidal suspensions},
  journal      = CaPS,
  publisher    = SPR,
  year         = {1991},
  volume       = {269},
  number       = {4},
  pages        = {295-319},
  month        = APR
}
% doi = {10.1007/BF00654577}
% url = {http://www.springerlink.com/content/t3374113443gn256/}

@ARTICLE{WuL2007,
  author       = {{Wu}, J. and {Li}, Z.},
  title        = {Density-functional theory for complex fluids},
  journal      = ARoPC,
  publisher    = AR,
  year         = {2007},
  volume       = {58},
  number       = {1},
  pages        = {85-112},
  month        = MAY
}
% doi = {10.1146/annurev.physchem.58.032806.104650}
% url = {http://adsabs.harvard.edu/abs/2007ARPC...58...85W}

@INCOLLECTION{TarazonaCMR2008,
  author       = {{Tarazona}, P. and {Cuesta}, J.~A. and {Mart{\'{\i}}nez-Rat{\'o}n}, Y.},
  title        = {Density Functional Theories of Hard Particle Systems},
  booktitle    = {Theory and Simulation of Hard-Sphere Fluids and Related Systems},
  publisher    = SPR,
  year         = {2008},
  editor       = {{\'A}. {Mulero}},
  volume       = {753},
  series       = {Lecture Notes in Physics},
  type         = {chapter},
  chapter      = {7},
  pages        = {247-341},
  address      = BER,
  edition      = {1},
  month        = AUG,
  isbn         = {978-3-540-78766-2}
}
% doi = {10.1007/978-3-540-78767-9_7}
% url = {http://adsabs.harvard.edu/abs/2008LNP...753..247T}

@ARTICLE{HessK1983,
  author       = {{Hess}, W. and {Klein}, R.},
  title        = {Generalized hydrodynamics of systems of {B}rownian particles},
  journal      = AiP,
  publisher    = TF,
  year         = {1983},
  volume       = {32},
  number       = {2},
  pages        = {247-341},
  month        = NOV
}
% doi = {10.1080/00018738300101551}
% url = {http://adsabs.harvard.edu/abs/1983AdPhy..32..173H}

@ARTICLE{CichockiH1987,
  author       = {{Cichocki}, B. and {Hess}, W.},
  title        = {On the memory function for the dynamic structure factor of interacting {B}rownian particles},
  journal      = PA,
  publisher    = ELS,
  year         = {1987},
  volume       = {141},
  number       = {2-3},
  pages        = {475-488},
  month        = MAR
}
% doi = {10.1016/0378-4371(87)90176-2}
% url = {http://adsabs.harvard.edu/abs/1987PhyA..141..475C}

@ARTICLE{SzamelL1991,
  author       = {{Szamel}, G. and {L{\"o}wen}, H.},
  title        = {Mode-coupling theory of the glass transition in colloidal systems},
  journal      = PRA,
  publisher    = APS,
