## Supplementary figures and images for "Analytical method for reconstructing the stress on a spherical particle from its surface deformation"

### fig1.pdf

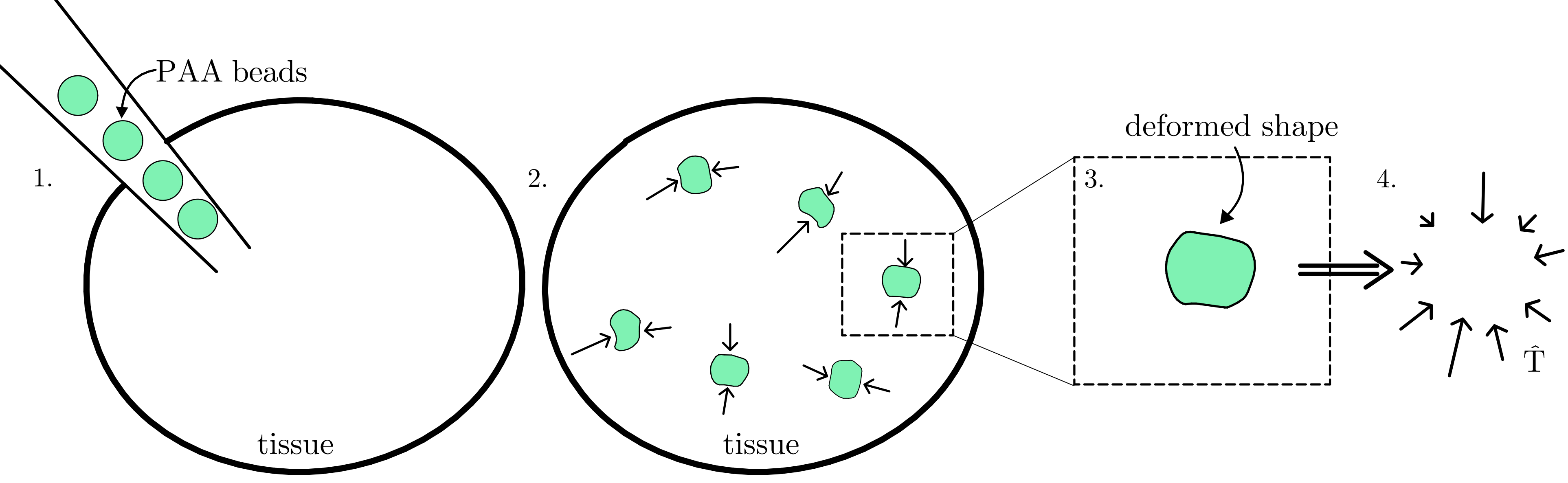

### fig2.pdf

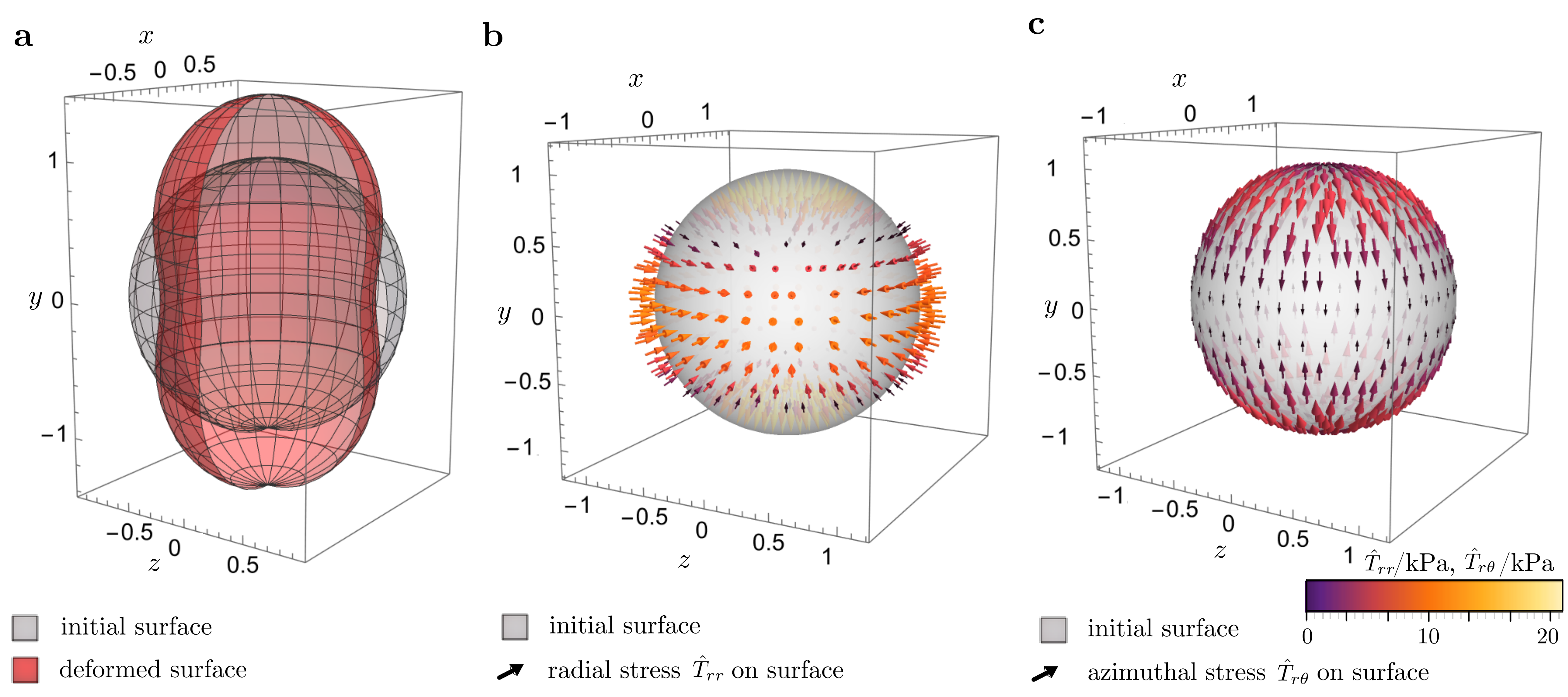

### fig3.pdf

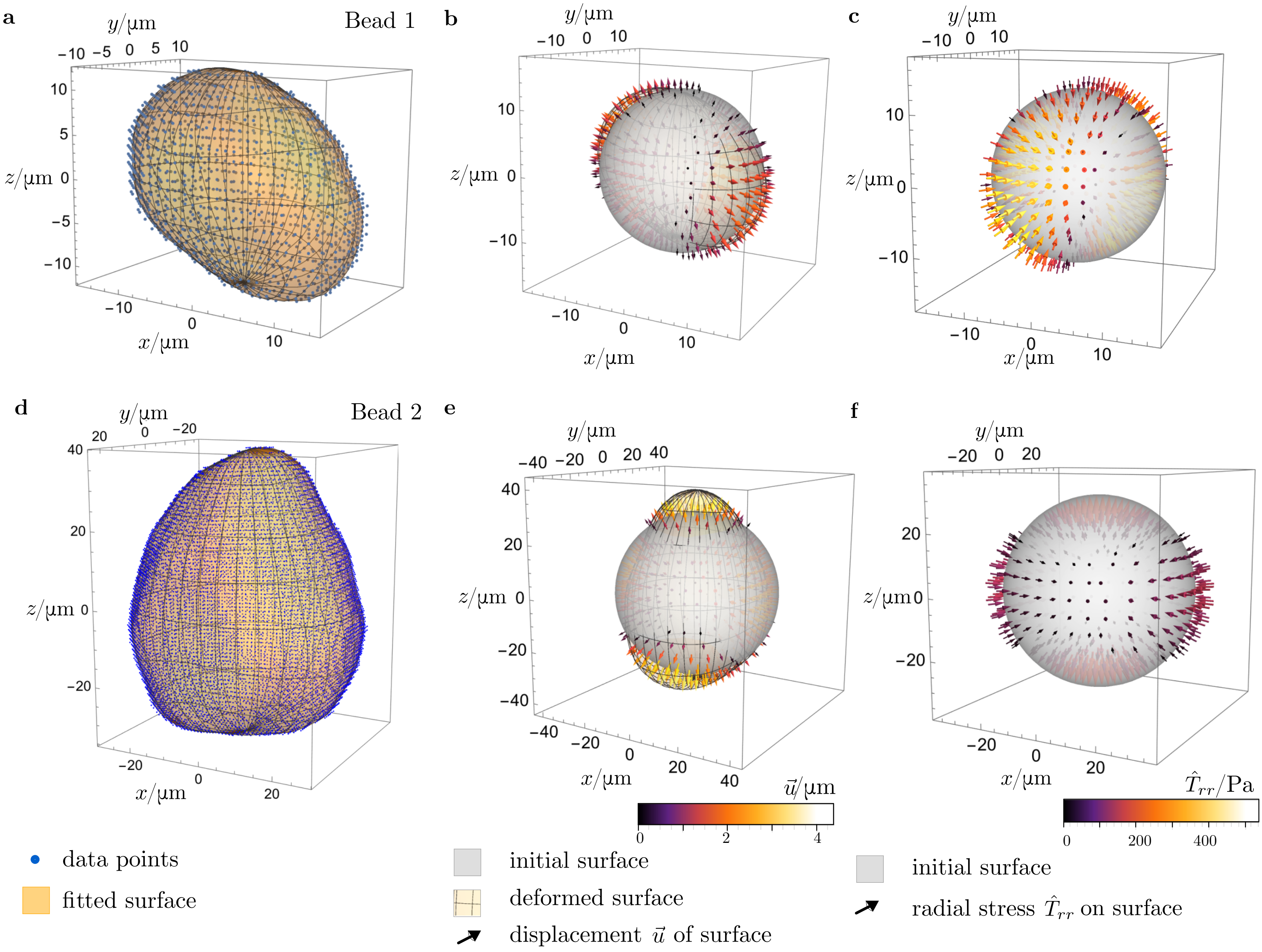
